# LSD Modulates Proteins Involved in Cell Proteostasis, Energy Metabolism and Neuroplasticity in Human Cerebral Organoids

**DOI:** 10.1101/2024.01.30.577659

**Authors:** Marcelo N. Costa, Livia Goto-Silva, Juliana M. Nascimento, Ivan Domith, Karina Karmirian, Amanda Feilding, Pablo Trindade, Daniel Martins-de-Souza, Stevens K. Rehen

## Abstract

Proteomic analysis of human cerebral organoids may reveal how psychedelics regulate biological processes, shedding light on drug-induced changes in the brain. This study elucidates the proteomic alterations induced by lysergic acid diethylamide (LSD) in human cerebral organoids. By employing high-resolution mass spectrometry-based proteomics, we quantitatively analyzed the differential abundance of proteins in cerebral organoids exposed to LSD. Our findings indicate changes in proteostasis, energy metabolism, and neuroplasticity-related pathways. Specifically, LSD exposure led to alterations in protein synthesis, folding, autophagy, and proteasomal degradation, suggesting a complex interplay in the regulation of neural cell function. Additionally, we observed modulation in glycolysis and oxidative phosphorylation, crucial for cellular energy management and synaptic function. In support of the proteomic data, complementary experiments demonstrated LSD’s potential to enhance neurite outgrowth in vitro, confirming its impact on neuroplasticity. Collectively, our results provide a comprehensive insight into the molecular mechanisms through which LSD may affect neuroplasticity and potentially contribute to therapeutic effects for neuropsychiatric disorders.

## INTRODUCTION

Lysergic acid diethylamide (LSD) is a psychedelic substance that induces altered states of consciousness characterized by changes in sensory perception, mood, and thought patterns (1). LSD’s psychedelic effects are primarily attributed to its agonist actions at brain serotonin 2A receptors (5-HT2ARs) (2,3). LSD also binds to dopaminergic, adrenergic, and other subtypes of serotonergic receptors (4). Studies have also shown that psychedelics can penetrate cellular membranes, allowing interaction with intracellular 5-HT2A receptors (5). LSD exhibits allosteric binding to the tropomyosin receptor kinase B (TrkB), enhancing TrkB’s interaction with brain-derived neurotrophic factor (BDNF) (6). The interaction with multiple receptors underscores the complex pharmacology of psychedelics.

LSD exhibited potential therapeutic effects for anxiety, depression, and addiction (7–9), conditions associated with impaired neuroplasticity (10–12). It is hypothesized that psychedelics like LSD exert long-term psychotropic effects by acutely inducing heightened plasticity in brain circuits (8). Emerging evidence suggests that LSD could cause lasting changes in neuronal plasticity and brain function (13). In rodents, LSD can stimulate structural remodeling, such as increased dendritic arborization and spinogenesis (14).

Despite important progress, the molecular mechanisms underlying LSD’s pro-plasticity properties remain incompletely defined, especially in human neural cells (15–17). Advances in stem cell technology enabled the generation of cerebral organoids that model aspects of human neurobiology (18,19). Additionally, analyzing the proteomic profiling of these organoids has provided insights into drug-induced functional changes in the human brain (20–22).

In a previous study (23), our group utilized proteomics data from human cerebral organoids exposed to 10 nM LSD to gain insights into the molecular mechanisms underlying improved cognitive performance in both rats and humans. Here, we exposed cerebral organoids to a higher concentration of 100 nM LSD and conducted a more comprehensive and in-depth proteomic analysis. Given the range of doses used in clinical trials, typically from 20 to 200 µg (24–26), understanding the impact of varying concentrations is relevant. To further investigate the neuroplastic changes identified through proteomics, we performed a functional neurite outgrowth assay on human brain spheroids.

## RESULTS AND DISCUSSION

### 45-day old human cerebral organoids present characteristic cytoarchitecture, cell diversity, and express serotonin 2A receptors

Human cerebral organoids partially reproduce the cytoarchitecture of the cortex and can develop multiple brain regions and cell types (19,27,28). In a previous study, we used 45-day-old organoids to study the effects of the psychedelic 5-MeO-DMT through proteomics, showing anti-inflammatory and neuroplastic effects (29). Thus, forty-five days of cultivation provide sufficient cellular and structural complexity, ensuring experimental consistency and reproducibility.

Here, we show the characterization of the brain organoids, conducted through immunofluorescence. The expression of serotonin receptors was evidenced by positive labeling of 5-HT2A receptors, known for their key role in mediating psychedelic effects (2,3). This labeling demonstrated a clear colocalization with the neuronal marker MAP2, indicating the presence of these receptors within neurons (Figure 1A). The analysis also revealed the presence of young neurons and neural progenitors, as evidenced by positive immunostaining for β-tubulin III (TUJ1) and PAX6, respectively. The distribution of TUJ1 was widespread throughout the organoids, whereas the expression of PAX6 was concentrated near the ventricles, delineating the ventricle-like zone (Figure 1B). GFAP-positive cells, representing radial glia and astrocytes, were present throughout the organoids, including within the ventricular zones (Figure 1C). Therefore, our cerebral organoids present characteristic cytoarchitecture, cell diversity, and express the primary receptor for psychedelic action, exhibiting the necessary elements to investigate a response to LSD exposure.

**Figure 1.**
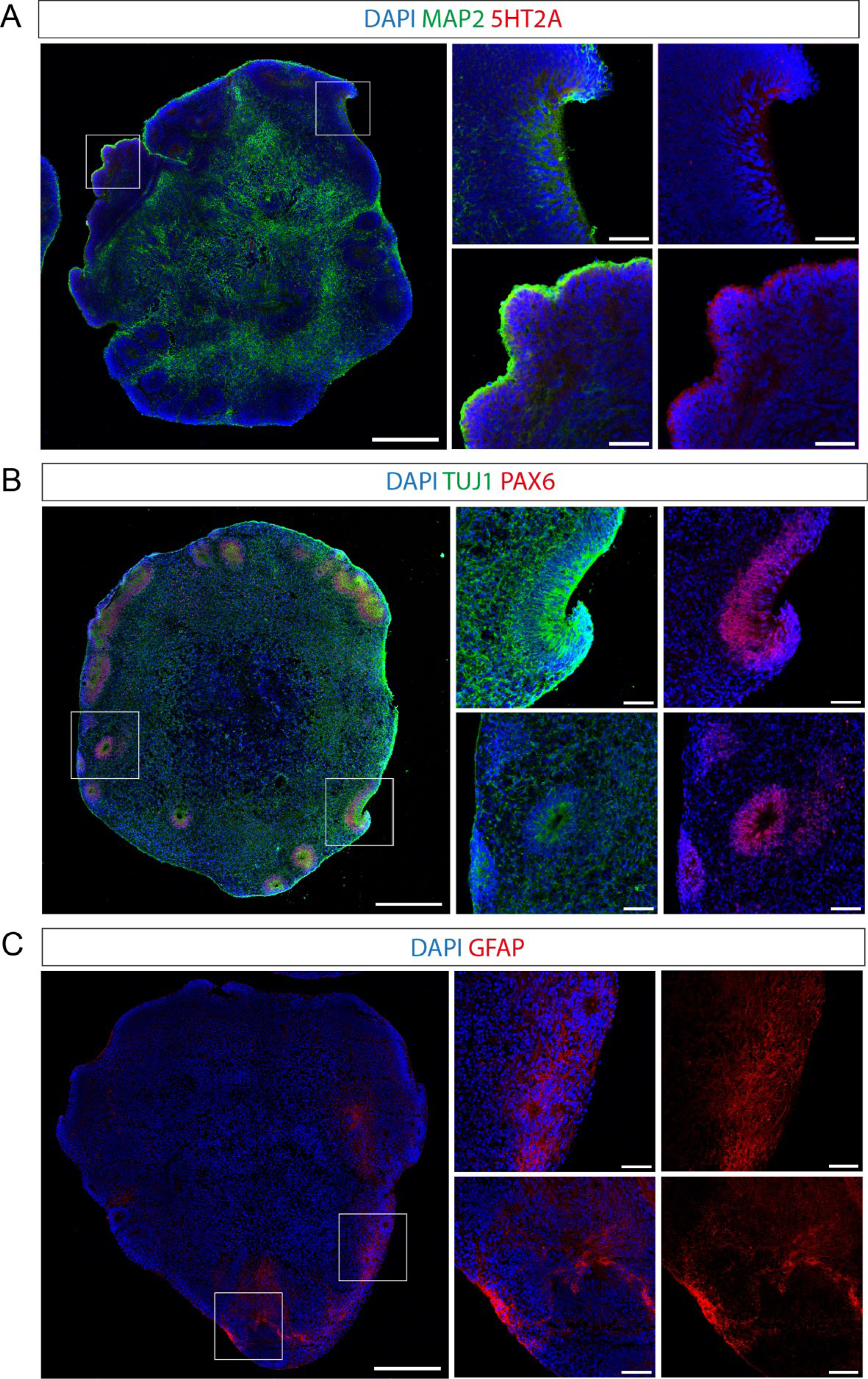
Characterization of the human cerebral organoids. Nuclei are stained blue with 4’,6-Diamidino-2-Phenylindole, Dihydrochloride (DAPI) in all images. (**A**) Immunostaining for 5-HT2A receptors, which colocalize with the neuronal marker MAP2; (**B**) TUJ1 immunostaining (green), a neuronal marker, accompanied by PAX6 immunostaining in red, which highlights neural progenitors; (**C**) GFAP immunostaining identifies radial glia and astrocytes. Scale bars represent 400 μm for whole organoid images and 60 μm for zoomed-in images.

### LSD changes the proteome of cerebral organoids

Determining LSD concentrations in human neural tissue remains challenging, as no pharmacokinetic study has directly measured LSD levels in the human brain. A postmortem study (30) indicated that LSD concentrations in the brain following typical recreational doses fall within the nanomolar range (up to 36 nM). This study also found that brain concentrations of LSD exceeded those in the blood in all three cases (up to 4 nM). In contrast, a pharmacokinetic study (31) reported even higher blood concentrations; with an average maximum plasma concentration of 9.6 nM following the administration of 200 µg of LSD. This discrepancy may be attributed to a smaller ingested amount or the timing of measurements taken after peak plasma concentration in the postmortem study.

Therefore, it is plausible that administering a dose of 200 µg could yield a concentration close to 100 nM in brain tissue, highlighting the importance of studying the effects of this higher concentration. Additionally, the exposure period was chosen considering the pharmacokinetic study mentioned above, which showed that LSD remained in the plasma of most human participants for up to 24 hours after a 200 μg oral dose (31).

After exposing the organoids to 100 nM LSD for 24 hours, protein extracts were subjected to liquid chromatography-tandem mass spectrometry (LC-MS/MS)-based shotgun proteomic analysis (Figure 2A). When comparing LSD-exposed and control organoids proteomes, we identified and quantified a total of 3195 proteins (false discovery rate [FDR] = 1%) (Table S1), with 239 showing significant differences in their abundance (p-value <0.05).

**Figure 2.**
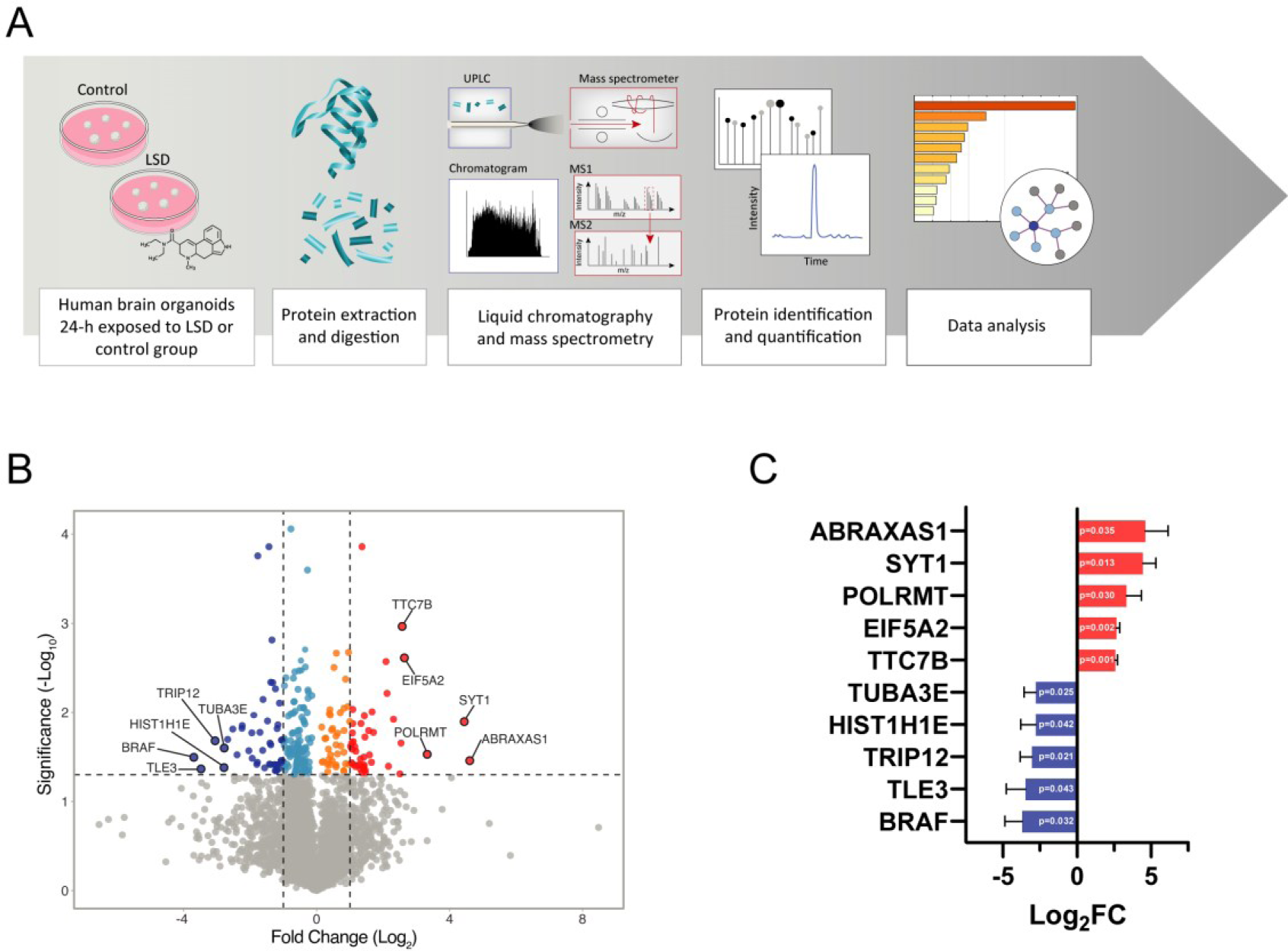
LSD alters the proteome of human cerebral organoids. In the experimental setup (**A**), 45-day-old organoids were exposed to either LSD or vehicle for 24 hours. Subsequently, label-free quantitative proteomic analysis was performed on the samples. The volcano plot (**B**) presents proteins identified by mass spectrometry-based shotgun proteomics in the organoids, highlighting DAPs induced by LSD. The significance threshold is marked by a horizontal line (p-value = 0.05), while two vertical lines denote the log fold change cut-off (−1.0 fold on the left and +1.0 fold on the right), distinguishing between minor and major alterations. Cold-colored circles represent proteins with significant decreases (dark blue for log_2_FC < −1.0; light blue for −1.0 ≤ log_2_FC > 0), whereas warm-colored circles indicate proteins with significant increases (orange for 0 < log_2_FC ≤ 1.0; red for log_2_FC > 1.0). Proteins without significant differences are shown in gray. The top five most increased and decreased proteins are also displayed. In part (**C**), a horizontal bar chart shows the proteins with the most notable abundance changes, along with their corresponding fold changes (log_2_) and p-values, which are indicated within the bars.

The names of all the differentially abundant proteins (DAPs) are shown in Table S2. Among these proteins, 158 were downregulated and 81 upregulated (Figure 2B). The proteins that exhibited the highest increases were the tumor-suppressor protein involved in DNA repair ABRAXAS1, the calcium sensor for endo- and exocytosis of synaptic vesicles SYT1; the mitochondrial RNA polymerase POLRMT; the eukaryotic translation initiation factor EIF5A2 and TTC7B, an isoform of a subunit of the PI4KIIIα complex, responsible for the first step in plasma membrane phosphoinositide synthesis. Among the proteins that were found to decrease in abundance, the ones that exhibited substantial downregulation were the serine/threonine kinase BRAF, the transcriptional corepressor TLE3, the transcriptional coactivator TRIP12 (a.k.a. MED1), the linker Histone H1.4 (HIST1H1E) and the α-tubulin isoform TUBA3E (Figure 2B and 2C).

### LSD regulates proteins involved in proteostasis, energetic metabolism, and neuronal plasticity

LSD-induced DAPs were subjected to enrichment analysis in Metascape (using KEGG and Reactome databases). The top 10 statistically enriched terms are shown in Figure 3A. The result exhibits a predominance of terms associated with cellular proteostasis (selective autophagy [-log_10_P = 7.77]; apoptosis [-log_10_P = 7.62]; and cellular responses to stress [-log_10_P = 6.46]), energetic metabolism (glycolysis [-log_10_P = 5.81]) and neuronal plasticity (signaling by Rho GTPases, Miro GTPases and RHOBTB3 [-log_10_P = 16.97]; membrane trafficking [-log_10_P = 12.63]; axon guidance [-log_10_P = 8.41]; RHOQ GTPase cycle [-log_10_P = 5.13]; RHOBTB GTPase cycle [-log_10_P = 5.10]; and transmission across chemical synapses [-log_10_P = 4.94]). Table S3 shows the complete results derived from the pathway and process enrichment analysis.

**Figure 3.**
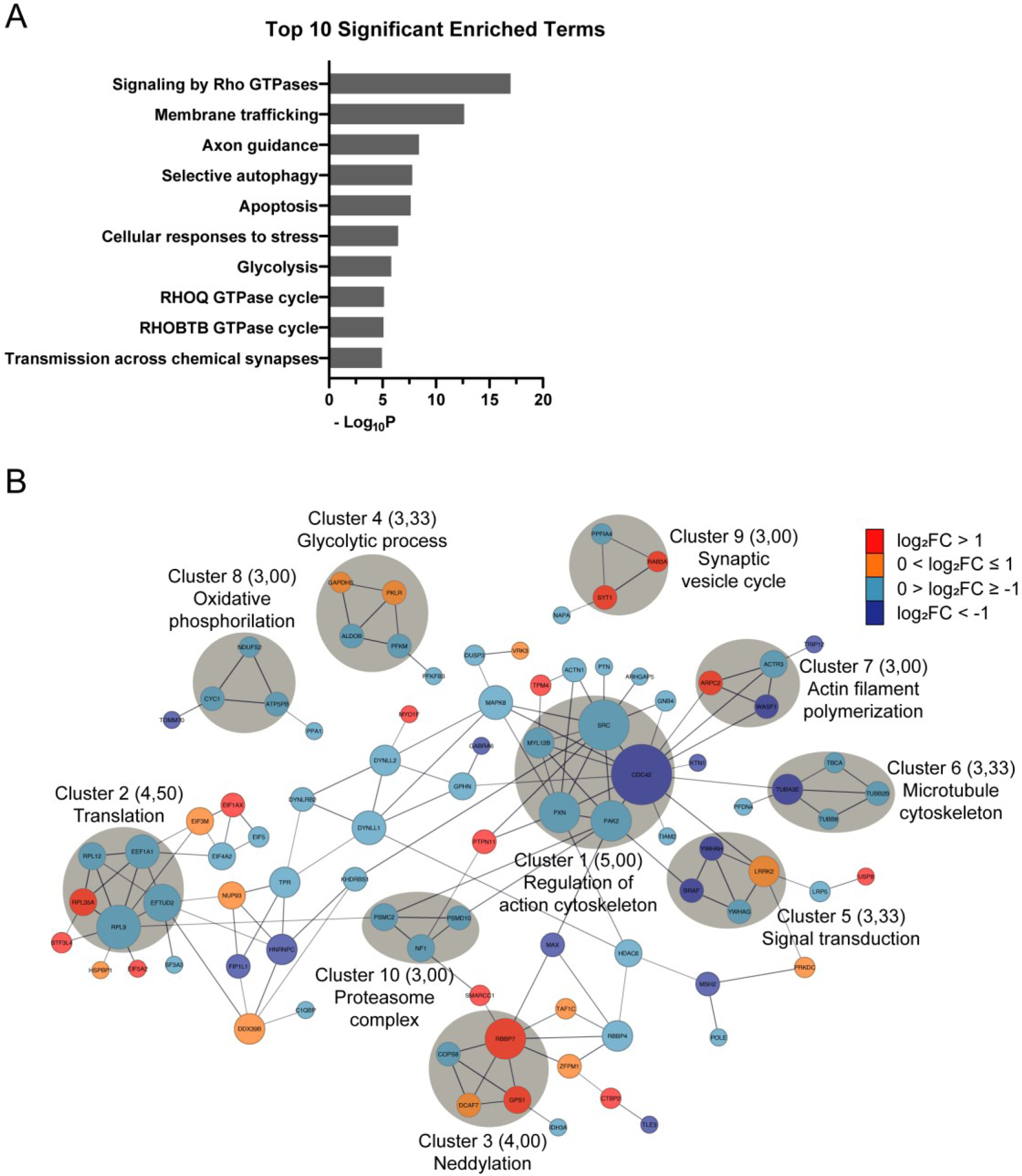
LSD influences proteins associated with proteostasis, cellular energy metabolism, and neuronal plasticity, as evidenced by: (**A**) The functional distribution of DAPs, which are categorized based on KEGG Pathway and Reactome Gene Sets; (**B**) A PPI network of DAPs induced by 100 nM LSD. For this network, the interaction score was set to a high confidence level (0.7). The node sizes represent the degrees (number of connections), with significant clusters highlighted in gray. The node colors indicate the extent of fold change: dark blue for more than two-fold decrease, light blue for less than two-fold decrease, orange for less than two-fold increase, and red for more than two-fold increase.

We also constructed a protein-protein interaction (PPI) network with LSD-induced DAPs (Figure 3B) using the STRING database. The resulting network comprised 237 nodes (proteins) and 159 edges (interactions). Notably, CDC42 displayed the highest degree of connection (14 degrees), followed by SRC (11 degrees), RPL9 (9 degrees), PAK2 (8 degrees), RBBP7 (8 degrees), and PXN (8 degrees). Among these highly connected nodes, RPL9 and RBBP7 were associated with proteostasis. At the same time, CDC42, SRC, PAK2, and PXN were linked to neuroplasticity and are part of a significant cluster related to this process. Altogether, we identified ten significant clusters using MCODE analysis. Three clusters (2, 3, and 10) were related to cellular proteostasis, two clusters (4 and 8) to energy metabolism, and three clusters (1, 7, and 9) to neuronal plasticity. Specifically, clusters 2, 3, and 10 exhibited enrichments in translation (p = 6.0E-6), neddylation (p = 1.1E-5), and the proteasome complex (p = 5.5E-3), respectively. Clusters 4 and 8 significantly correlated with the glycolytic process (p = 1.1E-8) and oxidative phosphorylation (p = 2.6E-4), respectively. Clusters 1, 7, and 9 were associated with the regulation of actin cytoskeleton (p = 5.9E-7), microtubule cytoskeleton (p = 8.6E-7), and synaptic vesicle cycle (p = 9.5E-3), in that order.

Regarding proteostasis, selective autophagy is crucial for neuronal homeostasis (32) and plays key roles in guidance signaling, dendritic spine architecture, spine pruning, vesicular release, and synaptic plasticity in neurons (33). Neddylation is a post-translational modification, wherein a particular family of E3 ubiquitin ligases are the best characterized substrates. This modification within these E3 ligases facilitates the ubiquitination of their respective targets (34,35). Furthermore, the enrichment of ’cellular responses to stress’ and ’apoptosis’ may stem from the abundant number of proteins involved in the proteostasis network, as ribosomal and proteasomal proteins. Various components of this network respond to proteotoxic stress, significantly influencing cellular decisions between apoptosis and survival (36–38). Thus, fundamentally, the regulation of the proteostasis network profoundly affects both the composition and functionality of the cellular proteome.

In reference to cellular energy metabolism, it encompasses the metabolic pathways responsible for ATP synthesis through NADH turnover. The two primary pathways involved in these processes are glycolysis/fermentation and oxidative phosphorylation (39). Notably, when investigating the functional enrichment and network analyses of DAPs induced by LSD, glycolysis prominently emerges as a modulated pathway in both analyses, while oxidative phosphorylation surfaces in the last one. Thus, it appears that LSD induces changes in proteins associated with cellular energy metabolism.

Neuroplasticity can be classified into structural and functional plasticity. Structural plasticity refers to changes in neuronal morphology (40,41), while functional plasticity encompasses modifications in synaptic transmission strength. Alterations in presynaptic neurotransmitter release and postsynaptic responses mediated by specific receptors can influence such changes (42). As part of structural plasticity, signaling by Rho GTPases and axon guidance emerged as potential pathways elicited by LSD in the functional enrichment and network analyses (Figures 3A and 3B). The precise control of neuronal cytoskeletal dynamics plays a pivotal role in orchestrating structural plasticity (40), whereby the Rho family of small GTPases emerge as critical modulators (43,44). Many signaling cascades implicated in neuronal structural modifications converge upon actin and microtubule cytoskeletal networks as shared terminal effectors (40). Moreover, Rho GTPases and their regulatory factors serve as critical downstream constituents of guidance signaling pathways (45), thereby establishing a connection between axon guidance molecules, pivotal for nervous system development, and their significance in governing synaptic plasticity in the mature brain (46,47). In terms of functional plasticity, our analysis uncovered enriched terms related to neurotransmitter release, membrane trafficking, and synaptic transmission, besides a cluster associated with the synaptic vesicle cycle in the interaction analysis. Modulation of the structural and functional aspects of neuroplasticity appears to be highly relevant when assessing the effects of LSD on these organoids.

### The proteostasis network dynamically adapts the proteome, enabling cellular responses to LSD stimuli

Given that our enrichment and network analysis revealed terms, hubs, and clusters associated with the molecular machinery responsible for protein quantity and quality control, we focused on the modulation of the proteostasis network pathway. In Figure 4, LSD-modulated proteins associated with each step of the network are identified and color-coded based on their regulation. We noted in this pathway that LSD-induced DAPs, which are predominantly reduced, potentially impact protein synthesis, folding, maturation, transport, targeting, and degradation via the ubiquitin-proteasome system and autophagy. A reduction in degradative pathways may extend the lifespan of key synaptic proteins, as their metabolic turnover is dependent on synthesis and degradation rates (48). Additionally, autophagy is known to be inhibited by mTOR pathway activation (49), a pathway significantly implicated in LSD’s neuroplastic effects (14). However, further research is necessary to confirm whether LSD specifically influences any specific aspect of the proteostasis network, or if these observations are merely homeostatic responses to the stimulus. Neuroplasticity itself poses a challenge to the proteostasis network (50), as signals inducing plasticity necessitate the adaptation of protein content to elicit specific responses (51).

**Figure 4.**
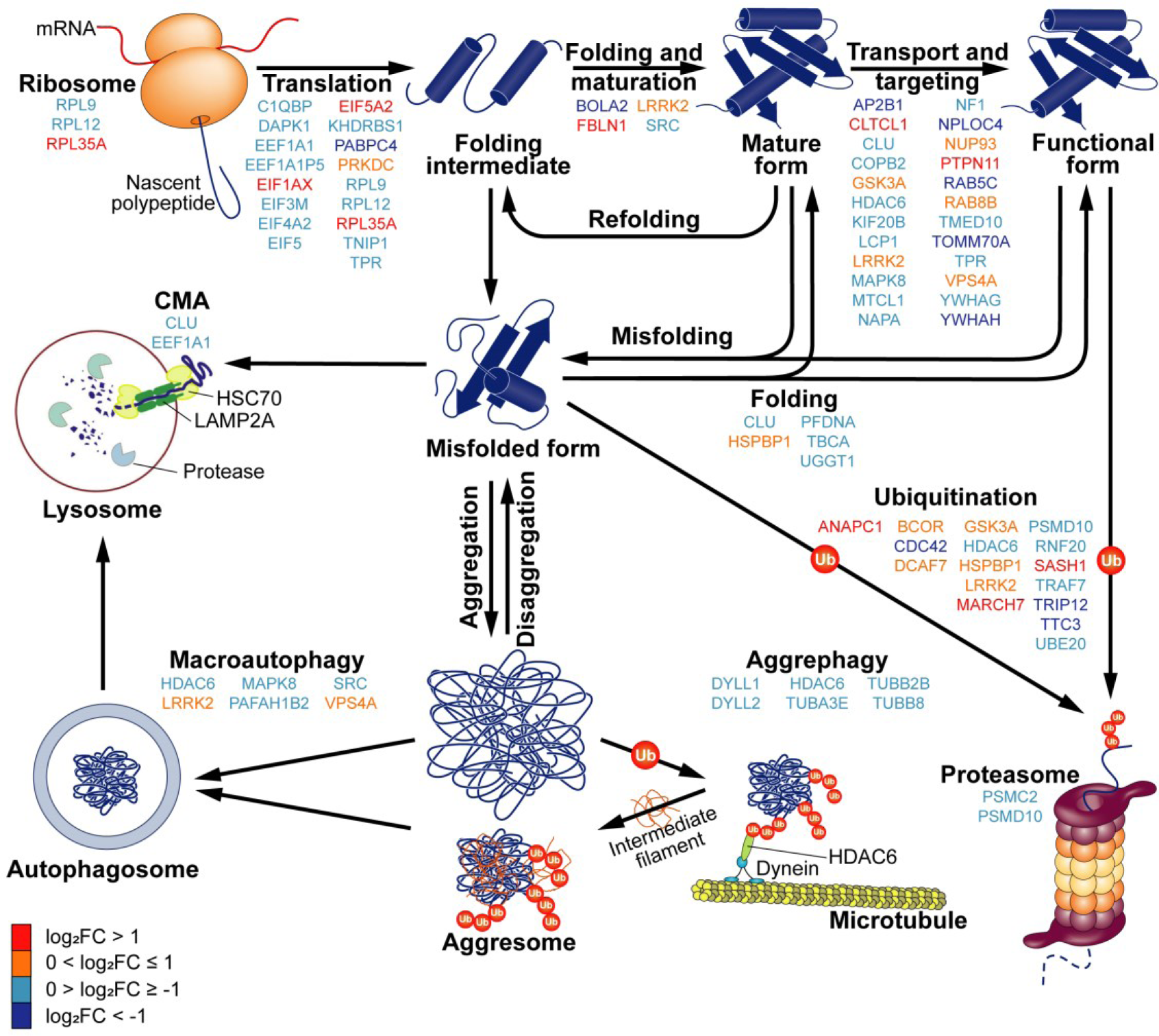
LSD modulates proteins within the proteostasis network. The schematic diagram illustrates the components of this network, comprising the synthesis, folding, maturation, targeting, and degradation of proteins. In this diagram, the DAPs affected by LSD are color-coded, reflecting the extent of their modulation: dark blue for more than two-fold decrease, light blue for less than two-fold decrease, orange for less than two-fold increase, and red for more than two-fold increase.

### Changes in neural energy metabolism to support neuroplasticity

Our enrichment and network analyses suggest that LSD modulates key pathways in energetic metabolism, including glycolysis, the tricarboxylic acid (TCA) cycle, and oxidative phosphorylation, as depicted in Figure 5. We observed that glycolysis is a pathway with four out of ten regulated steps. Modulated targets include phosphofructokinase PFKM and pyruvate kinase PKLR, representing two out of the three key steps responsible for the irreversible reactions of glycolysis. Furthermore, several proteins critical to the TCA cycle and oxidative phosphorylation were found to be downregulated. Upregulation of PKLR (log_2_FC = 0.37), coupled with the downregulation of proteins in the subsequent steps of mitochondrial energy metabolism, may favor lactate formation. In the brain, astrocytes emerge as the primary producers of this metabolite (52). Considering that periods of heightened neuronal activity and plasticity demand increased energy (53) and recognizing astrocyte-derived lactate as an alternative energy source for neurons (54), these findings may reflect an enhancement in the astrocyte-neuron lactate shuttle, aiming to support the high energetic demand of neuroplasticity.

**Figure 5.**
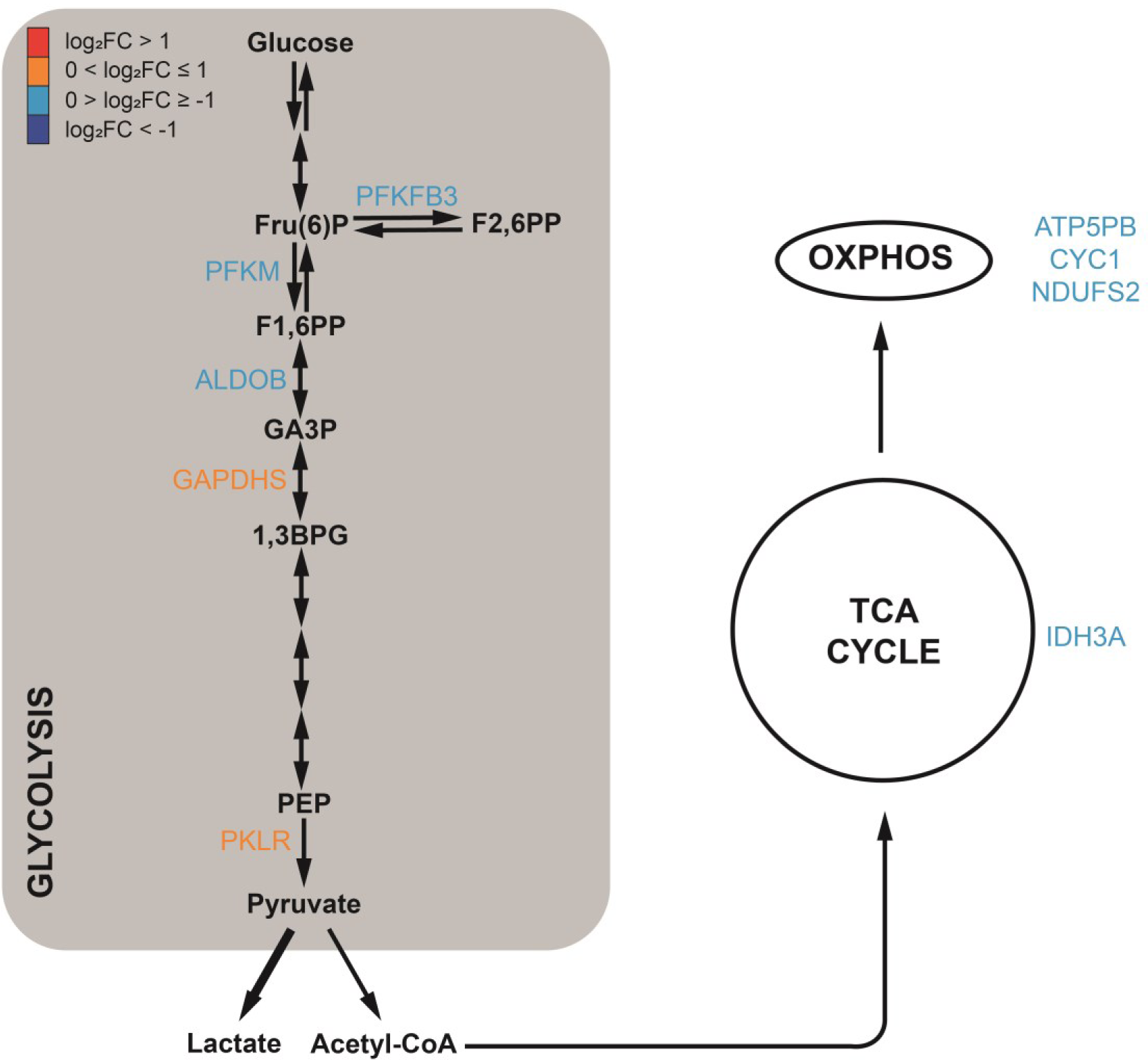
LSD exerts regulatory effects on proteins involved in key cellular energy metabolism pathways. The schematic representation highlights the most critical pathways: glycolysis, the TCA cycle, and oxidative phosphorylation. Within this schematic, the DAPs influenced by LSD in each pathway are displayed. These are color-coded to indicate the extent of their change: dark blue for more than two-fold decrease, light blue for less than two-fold decrease, orange for less than two-fold increase, and red for more than two-fold increase. 1,3BPG, 1,3-bisphosphoglycerate; F1,6PP, Fructose 1,6-bisphosphate; F2,6PP, Fructose 2,6-bisphosphate; Fru(6)P, Fructose 6-phosphate; GA3P, Glyceraldehyde 3-phosphate; OXPHOS, oxidative phosphorylation; PEP, Phosphoenolpyruvate.

### LSD modulates proteins involved in cytoskeleton regulation and release of synaptic vesicles pathways

LSD exerts modulation on several crucial biological processes involved in neuroplasticity, with two noteworthy examples being the actin cytoskeleton pathway (KEGG ID: hsa04810; p= 8.6E-3) and the synaptic vesicle cycle pathway (KEGG ID: hsa04721; p= 3.2E-2). We present a simplified representation of their regulation in response to DAPs induced by LSD (Figure 6A and 6B), as determined through KEGG pathway analysis. These findings underpin the previously described neuroplastic effects of LSD (14). Table 1 displays the representation names of the DAPs within pathways alongside their corresponding encoding genes.

**Figure 6.**
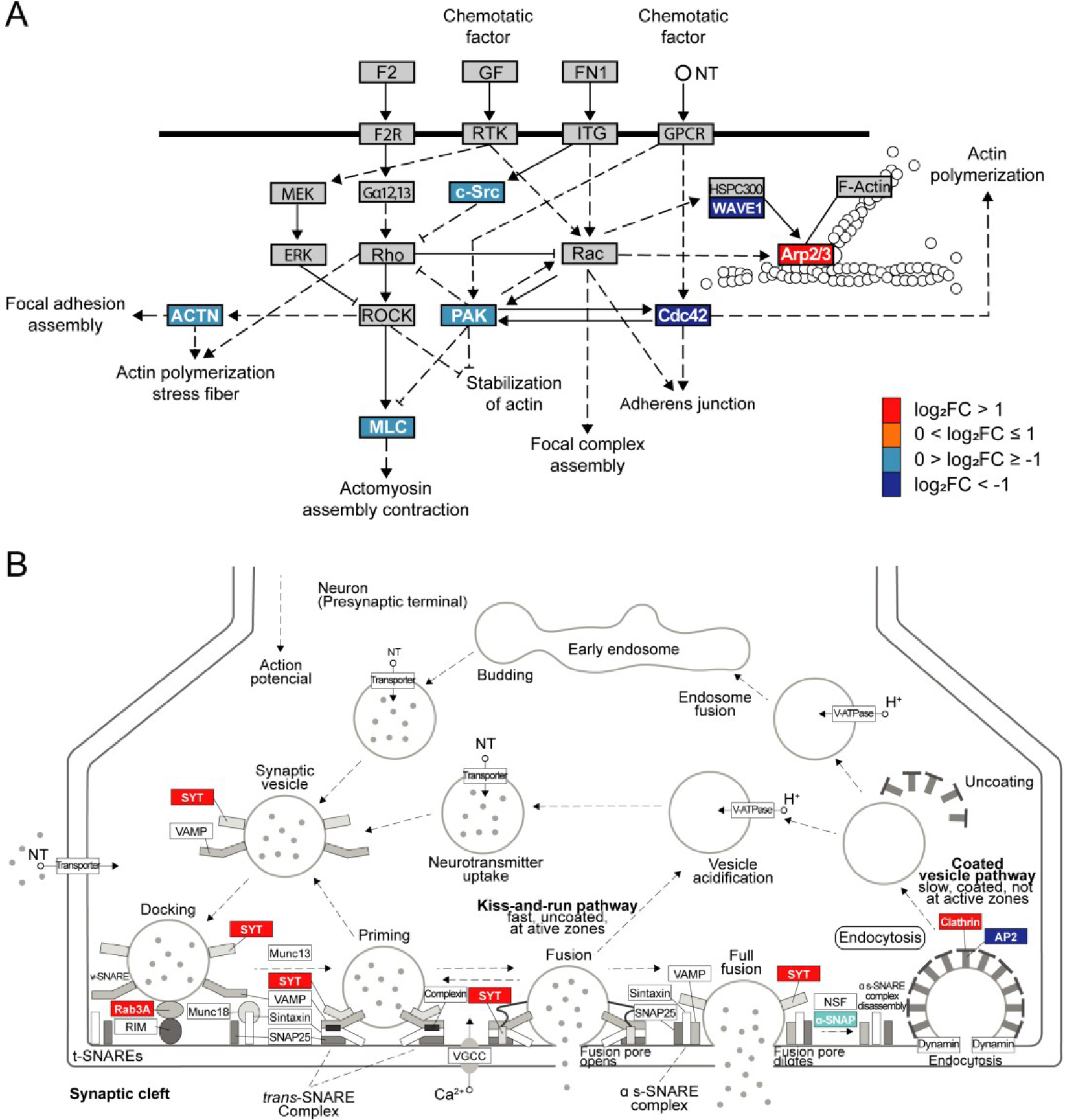
Signaling pathway diagrams exhibit LSD-modulated proteins involved in cytoskeleton regulation and neurotransmitter release. The LSD-induced DAPs involved in each step are depicted with distinct colors: dark blue (higher than two-fold decreased), light blue (less than two-fold decreased), orange (less than two-fold increased), or red (higher than two-fold increased): (**A**) LSD-induced modulation of proteins involved in the regulation of the actin cytoskeleton pathway. Figure adapted from KEGG (KEGG ID: hsa04810; p= 8.6E-3). ACTN, actinin; ERM, ezrin, radixin or moesin; F2, coagulation factor II; F2R, coagulation factor II receptor; FN1, fibronectin 1; GF, growth factor; GPCR, G protein-coupled receptors; Gα12,13, G protein subunit alpha 12, 13; ITG, integrin; MLC, myosin light chain; NT, neurotransmitter; ROCK, Rho kinase; (**B**) LSD-induced changes in proteins involved in synaptic vesicle cycle. Figure modified from KEGG (KEGG ID: hsa04721; p= 3.2E-2). AP2, adaptor protein 2 complex; NSF, N-ethylmaleimide-sensitive fusion protein; NT, neurotransmitter; RIM, Rab3 interacting molecule; SNAP, soluble NSF attachment proteins; SNARE, soluble NSF adaptor protein receptor; SYT, synaptotagmin; t-SNARE, target SNARE; VAMP, vesicle-associated membrane protein; VGCC, voltage-gated calcium channel; V-ATPase, vacuolar ATPase; v-SNARE, vesicle SNARE.

**Table 1.** DAP-encoding gene symbols and protein names in Figures 6A and 6B and their respective representations in the pathways. The upper section corresponds to Figure 6A, while the lower section refers to Figure 6B.

| Representation | Gene symbol | Protein name | Log<sub>2</sub>FC |
| --- | --- | --- | --- |
| ACTN | ACTN1 | Actinin alpha 1 | -0.543033 |
| Arp2/3 | ACTR3 | Actin related protein 3 | -0.395243 |
|  | ARPC2 | Actin related protein 2/3 complex subunit 2 | 1.40449 |
| Cdc42 | CDC42 | Cell division cycle 42 | -1.27146 |
| c-Src | SRC | SRC proto-oncogene, non-receptor tyrosine kinase | -0.296145 |
| MLC | MYL12B | Myosin light chain 12B | -0.675952 |
| PAK | PAK2 | p21 (RAC1) activated kinase 2 | -0.559334 |
| WAVE1 | WASF1 | WASP family member 1 | -1.06246 |
| α-SNAP | NAPA | NSF attachment protein alpha | -0.39629 |
| AP2 | AP2B1 | Adaptor related protein complex 2 subunit beta 1 | -1.20892 |
| Clathrin | CLTCL1 | Clathrin heavy chain like 1 | 1.23632 |
| Rab3A | RAB3A | RAB3A, member RAS oncogene family | 1.42592 |
| SYT | SYT1 | Synaptotagmin 1 | 4.44077 |

According to the regulation of actin cytoskeleton map (Figure 6A), signaling to the cytoskeleton can be mediated through G protein-coupled receptors (GPCRs), integrins, and receptor tyrosine kinases (RTKs), leading to a wide range of effects, including alterations in cell shape. Indeed, it is well known that LSD can activate various G protein-coupled serotonin receptors, also increasing the levels of BDNF, which acts through the tyrosine kinase receptor TrkB (3,24). Additionally, it was recently shown that LSD can directly bind TrkB receptors, thereby facilitating the action of BDNF (6) and regulates the expression of extracellular matrix proteins, including fibronectin (55).

Intracellularly, cellular responses are regulated through numerous signaling cascades, which include the Rho family of small GTPases and their downstream protein kinase effectors. Arp2/3 complex is a key factor in actin filament branching and polymerization, essential for dendritic spine structural plasticity and stability (56) which was upregulated in LSD exposed organoids. LSD also modulated Rho GTPase CDC42, the protein kinase PAK1, and WASF1, a member of the actin regulatory WAVE complex. Therefore, our findings suggest modulation of actin cytoskeleton, including focal adhesions, adherens junctions, actomyosin, and stress fibers (Figure 6A).

In the synaptic vesicle cycle pathway, we observed upregulation of proteins involved in the fusion of synaptic vesicles to the plasma membrane (Figure 6B). Rab proteins located on the vesicle membrane can form complexes with effector proteins, such as rabphilin and rab-interacting molecule (RIM), to facilitate the docking of synaptic vesicles (57). SYT1, one of the most upregulated proteins in the analysis, functions as the primary calcium sensor for neurotransmitter release. Upon Ca^2+^ binding, SYT1 triggers the complete assembly of the soluble NSF adaptor protein receptor (SNARE) complex, which leads to rapid and synchronized membrane fusion (58,59).

Additionally, proteins associated with the recovery and recycling of synaptic vesicles, particularly in clathrin-mediated endocytosis (CME), exhibited significant alterations in abundance following exposure to LSD (Figure 6B). SYT1, up-regulated, not only functions as the Ca^2+^ sensor for neurotransmitter release but also acts in endocytosis (59). The soluble N-ethylmaleimide-sensitive factor (NSF) attachment protein α (α-SNAP, encoded by NAPA) is downregulated. α-SNAP, along with NSF, disassembles the SNARE complex after fusion (60). One isoform of the clathrin heavy chain was upregulated, whereas one subunit of the clathrin adaptor protein 2 (AP2) complex (AP2B1) was downregulated (Figure 6B).

### Two distinct LSD concentrations share modulation of proteins involved in regulating cell morphology and synaptic processes

We compared our dataset of human cerebral organoids exposed to 100 nM LSD with our prior data obtained from exposure to 10 nM LSD (23). The comparison showed that distinct LSD concentrations resulted in a similar number of significantly modulated proteins, being 234 when exposed to 10 nM LSD and 239 to 100 nM LSD. Upon exploring the datasets, we identified 26 proteins that were modulated in both concentrations, as shown in figure 7A (purple lines). These persistently modulated proteins are potential candidates for a molecular pattern of LSD action in human brain. These 26 proteins are listed in Table S4.

**Figure 7.**
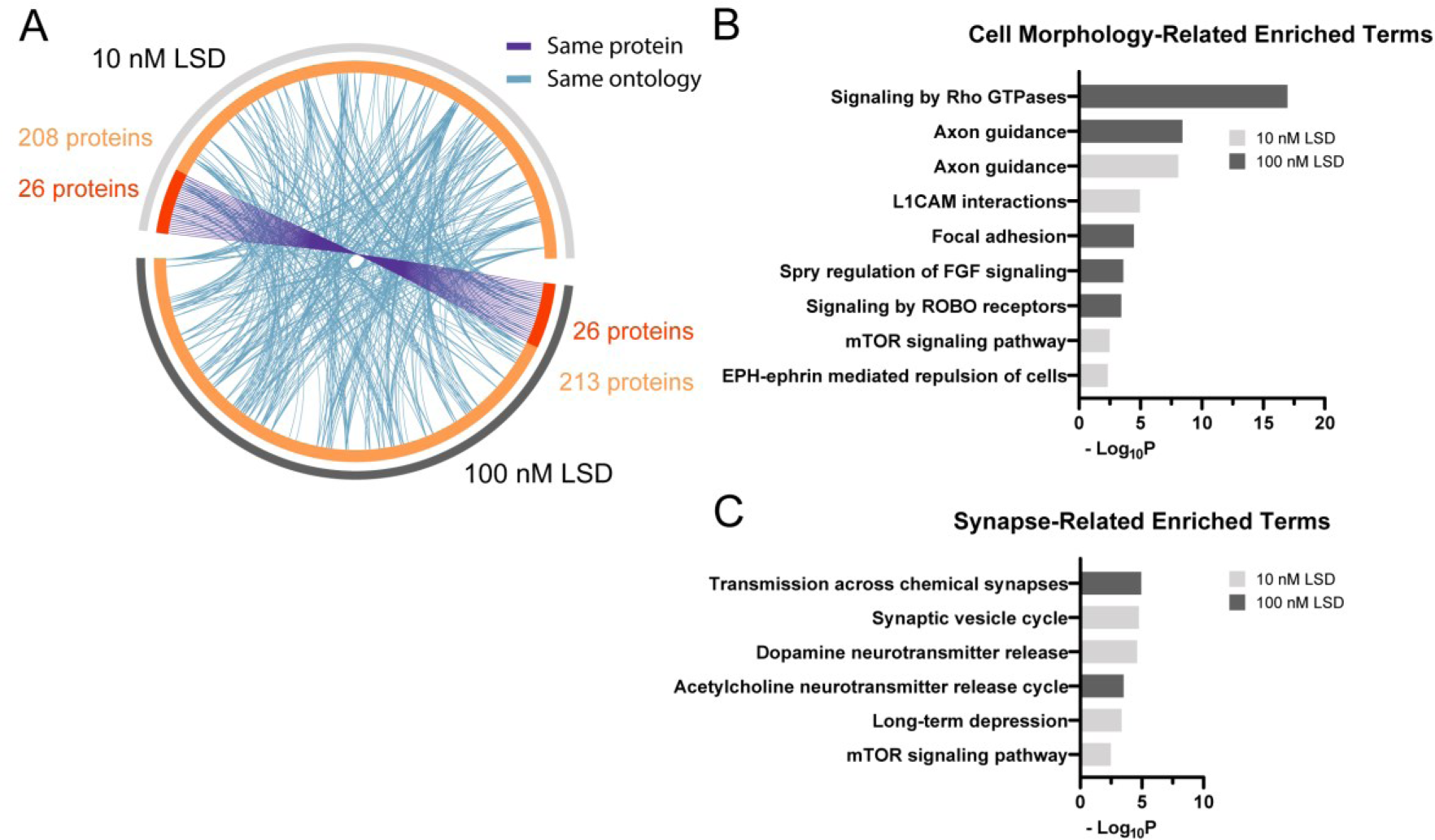
Different LSD concentrations share modulation of several biological processes: (**A**) Overlap between genes encoding DAPs induced by 10 and 100 nM LSD. On the outside, the light gray arch corresponds to the 10 nM LSD gene list, and the dark gray to 100 nM LSD. On the inside, light orange represents genes uniquely modulated by one condition, and in dark orange those regulated in both conditions. The purple lines link the same genes common in both 10 nM and 100 nM LSD conditions. Blue lines link different genes with the same ontology term; (**B**) and (**C**) Significant terms associated with alterations in cell morphology and synaptic processes, respectively. The light gray bars correspond to data from the 10 nM LSD concentration, while the dark gray bars represent data from the 10 nM LSD concentration.

Among the consistently modulated proteins, the ɑ6 subunit of GABA_A_ receptors (GABRA6) and the protein O-GlcNAcase (OGA) emerge as commonly downregulated proteins. Besides the well-established roles of GABA_A_ receptors, emerging evidence has identified this very specific subunit as a promising therapeutic target for addressing a wide spectrum of neurological and neuropsychiatric disorders (61,62). Interestingly, functional magnetic resonance imaging (fMRI) studies suggest that LSD perturbs the excitation/inhibition balance of the brain (63). On the other hand, OGA is an enzyme that mediates the removal of O-GlcNAc from target proteins (64). This modification was described to occur in many biological processes such as regulation of gene expression, signal transduction, cellular stress response, metabolism (65), and, more recently, in synaptic function (66). Reduction of OGA levels can decrease dendritic spine density in primary neurons (66). Although the exact roles of O-GlcNAcylation in synaptic function remain unclear, research has extensively investigated its role in aging-associated neurodegenerative diseases, such as Alzheimer’s and Parkinson’s disease. Elevated O-GlcNAcylation has been shown to prevent protein aggregation and slow neurodegeneration, making it a promising therapeutic target for these conditions (64). Another noteworthy modulation is the LSD-induced upregulation of copine-1 (CPNE1) in both concentrations. Studies have demonstrated that CPNE1 activates the AKT-mTOR signaling pathway in neural stem cells (NSCs) (67). Given the established association between mTOR and psychedelic-induced neural plasticity (14,23), CPNE1 and other copines may be potential upstream regulators of LSD-induced mTOR-mediated plasticity in other cell types as well.

While only a few hits were common at both concentrations, numerous proteins modulated in one group shared a common ontology with those from the other group (Figure 7A – blue lines). This functional overlap underscores the remarkable similarity in the elicited biological processes, despite the modulation of several distinct proteins. By comparing the top 20 enriched terms from both analyses, we were able to identify processes commonly modulated. The presence of multiple terms associated with the regulation of cell morphology (Figure 7B) and synaptic-related processes (Figure 7C), which are crucial for structural and functional plasticity, respectively, suggests a characteristic of LSD action at both concentrations.

### LSD induces neurite outgrowth in human brain cells

To specifically investigate the impact of LSD on structural plasticity, we conducted a neurite outgrowth assay, and the results are presented in Figure 8. In this assay, brain spheroids were initially plated on poly-ornithine/laminin-coated plates for 24 hours. They were then exposed to LSD at concentrations of 10 nM and 100 nM for an equivalent duration. Representative images are displayed in Figure 8A. In our research, we employed the Sholl analysis to quantify changes on arbor complexity induced by LSD. This method focuses on the evaluation of neurite intersections against a sequence of concentric circles of gradually increased radius, drawn around the core. A visual representation of the Sholl analysis conducted on these images is available in Figure S1.

**Figure 8.**
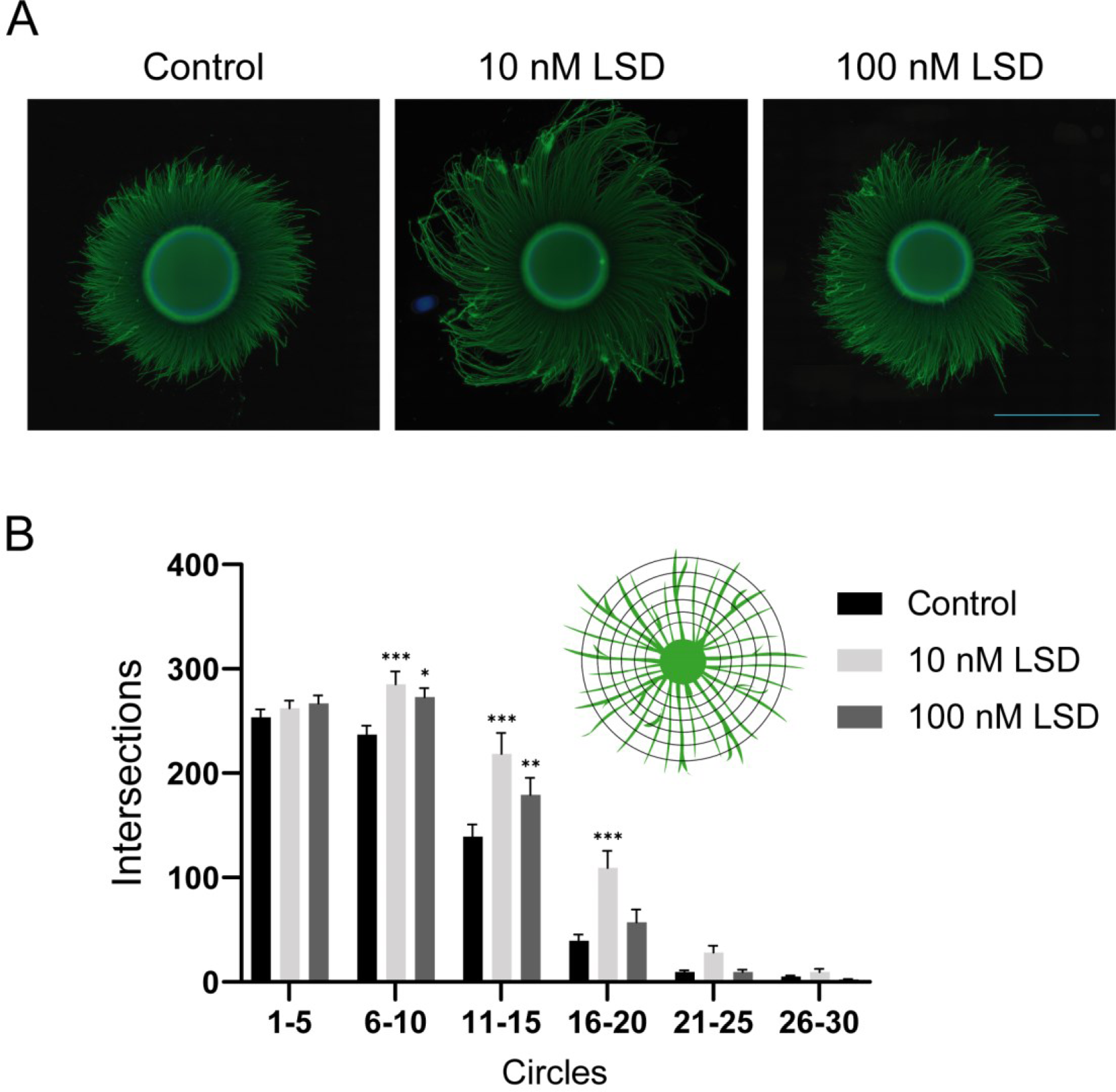
Neurite Outgrowth in Response to LSD: Human brain spheroids were initially plated for 24 hours and then subjected to exposure to either 10 nM or 100 nM LSD for an additional 24 hours. For this analysis, the control group included 18 spheroids, while the groups exposed to 10 nM and 100 nM LSD consisted of 17 and 15 spheroids, respectively. (**A**) The samples were analyzed using immunofluorescence staining for β-tubulin III (TUJ1; depicted in green) and DAPI staining to highlight nuclei (shown in blue). (**B**) Sholl analysis was employed to assess neurite outgrowth. An inset in this section provides a schematic of the Sholl analysis method. The analysis was conducted on the mean numbers of crossings for each group of five circles. The quantification was carried out over five independent experiments. Statistical significance is indicated as follows: *p<0.05, **p<0.01, ***p<0.001, in comparison to the control group. A scale bar indicating 1000 µm is included for reference.

Neurite intersections of circles were analyzed by mean crossings in groups of five successive circles. Both concentrations, 10 and 100 nM LSD, led to an increase in the average number of neurite crossings in circles 6 to 10 (control: 236.77 ± 8.65; 10 nM LSD: 285.33 ± 12.04, p<0.01; 100 nM LSD: 272.84 ± 8.65, p<0.05), and 11 to 15 (control: 139.13 ± 11.75; 10 nM LSD: 218.46 ± 20.00, p<0.001; 100 nM LSD: 179.01 ± 16.62, p<0.05). The concentration of 10 nM LSD still increased the number of crossings in circles 16 to 20 (control: 39.63 ± 5.92; 10 nM LSD: 109.16 ± 16.44, p<0.001; 100 nM LSD: 57.07 ± 12.44). These findings suggest that both concentrations led to increased arbor complexity (branching and/or elongation). Interestingly, different LSD concentrations can impact neurite outgrowth dynamics in varying ways. Therefore, further studies are essential to comprehend these dose-dependent variations and temporally distinct responses (Figure 8B).

The lack of significant changes in the number of crossings in circles 1 to 5 (control: 253.63 ± 7.66; 10 nM LSD: 262.09 ± 7.28; 100 nM LSD: 266.85 ± 7.70) suggests that the observed effects weren’t influenced by alterations in the number of primary neurites (Figure 8B). One the other hand, the absence of significant changes in intersections of circles 21 to 25 (control: 9.49 ± 1.78; 10 nM LSD: 28.04 ± 6.83; 100 nM LSD: 9.39 ± 2.30) and 26 to 30 (control: 5.19 ± 1.14; 10 nM LSD: 9.42 ± 3.18; 100 nM LSD: 2.61 ± 0.38) indicates that the effect was more pronounced in the proximal segments of the neurites (Figure 8B).

These findings are consistent with the results of Ly et al. (2018) in primary cortical rat cultures, where LSD was also found to promote neurite outgrowth. Similar to our observations, Ly et al. reported that LSD had a limited impact on the number of primary neurites and did not cause changes in the length of the longest dendrite (14). This parallel in outcomes across different experimental models underscores the potential of LSD in modulating neural structure, hinting at a consistent biological response irrespective of the model organism.

## CONCLUSIONS

Our study reveals that LSD exposure leads to a significant alteration in the abundance of numerous proteins in human cerebral organoids, marking a shift in the proteomic profile of human neural cells. The enrichment analysis of these DAPS indicates that LSD affects processes such as proteostasis, energy metabolism, and neuroplasticity.

LSD modulates proteins involved in various aspects of the proteostasis network, including protein synthesis, folding, maturation, transport, autophagy, and proteasomal degradation. A notable observation is the reduction in most proteostasis proteins, potentially extending the lifespan of synaptic proteins by decelerating turnover rates reliant on a balance between synthesis and degradation (48). Additionally, LSD appears to inhibit autophagy, possibly through activation of the mTOR pathway (49), a known mechanism of LSD-induced neuroplasticity (14). However, it remains to be investigated whether LSD’s regulation of proteostasis is a direct effect or an indirect homeostatic response. The adaptation in proteostasis is crucial for proteome remodeling and cellular plasticity (50,51).

LSD impacts the abundance of proteins involved in glycolysis, the TCA cycle, and oxidative phosphorylation. This suggests that psychedelics could induce metabolic changes to accommodate the high demands during neural excitation and plasticity (53). Our data points to an increase in the lactate production, a primary energy source from astrocytes supporting neuronal plasticity (52,54).

Our analysis also implicates LSD in pathways essential for structural and functional neuroplasticity, including cytoskeletal regulation and neurotransmitter release. The remodeling of dendrites requires precise control over actin and microtubule dynamics, typically mediated by Rho GTPases (40,43). Additionally, LSD seems to enhance synaptic vesicle fusion proteins while reducing components of clathrin-mediated endocytosis, hinting at increased neurotransmitter release, though its implications for reuptake warrant further investigation.

Lastly, the comparison of proteins modulated in human cerebral organoids exposed to 100 nM LSD and those exposed to 10 nM LSD (23) shows a significant overlap in ontology among the modulated proteins at both concentrations. Interestingly, this overlap is particularly pronounced in terms associated with regulation of cell morphology, and synaptic-related processes. The presence of these terms points towards events encompassing structural and functional plasticity, respectively. These biological processes, consistently regulated at both concentrations, appear to be important hallmarks of LSD action in the human brain. Furthermore, our research revealed that LSD stimulates neurite outgrowth in iPSC-derived brain spheroids. We observed this effect at both concentrations, 10 nM and 100 nM, where LSD was found to enhance the complexity of the neurites. This finding suggests a broader spectrum of LSD biological activity on neuronal plasticity.

In conclusion, our proteomic analysis uncovers potential mechanisms behind the LSD-induced plasticity previously reported (14). Neuroplasticity induced by LSD was demonstrated in both proteomics and neurite outgrowth assay. Overall, these findings confirm neuroplastic effects induced by LSD in human cellular models and underscores the potential of psychedelics in treating conditions associated with impaired plasticity. Our study also highlights the value of human cerebral organoids as a tool for characterizing metabolic responses to psychedelics and deciphering aspects of neuroplasticity.

## METHODS

### hiPSC-derived human cerebral organoids

GM23279A, an hiPSC line from a healthy female donor obtained from the NIGMS Repository of the Coriell Institute, was used in this study. Cells were cultured on Matrigel (Corning, USA) coated plates with mTeSR1 (STEMCELL Technologies, Canada) at 37°C and in a 5% CO_2_ atmosphere. Colonies were manually passed when 80% confluence was reached. hiPSCs were differentiated into cerebral organoids as described by Goto-Silva et al. (68). Shortly afterward, hiPSCs were detached and dissociated with Accutase (MP Biomedicals, USA) for 5 min at 37°C, generating single cells in suspension. Then, 9,000 cells were added in each well of an ultra-low binding 96-well plate (Corning, USA) in human embryonic stem cells (hESC) media (Dulbecco’s modified eagle medium/Ham’s F12 [DMEM/F12; Gibco, USA], 20% KnockOut Serum Replacement [KSR; Gibco, USA], 3% fetal bovine serum, certified, United States [Gibco, USA], 1% Glutamax [Gibco, USA], 1% minimum essential media-nonessential amino acids [MEM-NEAA; Gibco, USA], 0.7% β-mercaptoethanol [Gibco, USA] and 1% Penicillin-Streptomycin [Pen-Strep; Gibco, USA]) with 4 ng/ml basic fibroblast growth factor (bFGF; Invitrogen, USA) and 50 µM Rho-associated protein kinase inhibitor (ROCKi; Calbiochem, USA). This media was changed every other day for 6 days, and the embryoid bodies transferred to low-adhesion 24-well plates (Corning, USA) in neural induction media (DMEM/F12, 1% N2 supplement [Gibco, USA], 1% Glutamax, 1% MEM-NEAA, 1 µg/ml heparin [Sigma-Aldrich, USA]) and 1% Pen-Strep. The neural induction media was changed every other day for 4 days. On day 10, tissues were immersed, for 1h at 37°C, in a solution of Matrigel diluted in DMEM/F-12, according to the dilution factor given by the manufacturer. After this step, coated organoids were returned to the 24-well ultra-low-attachment plates with differentiation media without vitamin A [1:1 mixture of DMEM/F12 and Neurobasal (Gibco, USA), 0.5% N2 supplement, 1% B27 supplement without vitamin A (Gibco, USA), 3.5 ml/l 2-mercaptoethanol, 1:4,000 insulin (Sigma-Aldrich, USA), 1% Glutamax, 0.5% MEM-NEAA and 1% Pen-Strep] for 4 days, changing the medium every 48h. After the stationary growth, the organoids were transferred to 6 well plates under agitation (90 RPM), containing differentiation media with vitamin A [same composition of differentiation media above, except B27 supplement with vitamin A (Gibco, USA)]. These media were replaced every 4 days until day 45.

### LSD exposure

LSD (Lipomed, USA) was dissolved in ultrapure water (18.2 MΩ.cm) at room temperature, protected from light exposure. 45-day-old human cerebral organoids were exposed to 100 nM LSD for 24 hours, and the control group received culture medium.

### Immunofluorescence in organoid sections

Five cerebral organoids were collected per condition (control and LSD) from each of the three independent experiments. Organoids were fixed overnight in 4% paraformaldehyde (PFA), rinsed with phosphate-buffered saline (PBS), dehydrated, cryoprotected in a 30% sucrose solution, and kept at 4°C for 48 – 72h. They were transferred into Tissue-Tek O.C.T. Compound (Sakura Finetek Japan, Japan), snap-frozen on dry ice, and stored at -80°C. Organoids were sectioned with 20 µM thickness in a cryostat (Leica Biosystems, Germany), and the slides were maintained at -80°C until staining with specific markers.

For immunofluorescence, slides were thawed for 30 min at 37°C, washed three times with PBS, and permeabilized with 0.3% Triton X-100 (Sigma-Aldrich, USA) diluted in 1X PBS for 15 min. For specific stainings (5-HT2A and PAX6), antigen retrieval procedure was performed before the permeabilization step. Sections were incubated in 10mM citrate buffer, 0.05% Tween 20, pH=6 for 10 min at 98°C. Following this step, the sections were blocked with 1% BSA + 10% normal goat serum (NGS) in PSB for 2 hours. All antibodies were diluted in this blocking solution. Cryosections were incubated overnight with primary antibody at 4°C, washed three times for 5 min with PBS, and incubated with secondary antibody for 2h. The antibodies used in this study are indicated in Table S5. After three more washes with PBS for 5 min, the sections were incubated with 300 nM DAPI (Invitrogen, USA) for 5 min, for nuclear staining. Then, a last round of three washes with PBS was performed, and the slides were cover-slipped with Aqua-Poly/Mount (Polysciences, USA).

Images of organoid slices were acquired in a Leica TCS SP8 confocal microscope. For the immunofluorescence images, a 20x oil-immersion objective lens was used.

### Liquid chromatography-mass spectrometry

Liquid chromatography-mass spectrometry. Ten 45-day-old human cerebral organoids were collected from each of three independent experimental batches. Five of these organoids were exposed to 100 nM LSD for 24 hours, and the other five received only medium (control condition). After exposure, organoids were pelleted and frozen at −80 °C until sample processing for mass spectrometry-based label-free shotgun proteomics. Organoids were lysed in a buffer containing 7 M urea, 2 M thiourea, 1% CHAPS, 70 mM DTT, and Complete Protease Inhibitor Cocktail (Roche, Switzerland). Total protein content was measured using the bicinchoninic acid (BCA) method. The protein extracts (50 µg) were shortly loaded to an 10% SDS-PAGE gel by electrophoresis, and slices of each sample (containing all proteins) were subjected to in gel reduction (2 mM DTT), alkylation (10 mM iodoacetamide), and then digested with trypsin at 37°C overnight (1:100, w/w, Sigma-Aldrich, USA). Peptides were extracted from gels with acetonitrile/formic acid (50%/5%) solution and 100% acetonitrile solution. Peptides were then dried with SpeedVac (Thermo Fisher Scientific) and stored at -80°C until use. Prior to analyses peptides were diluted in 0.1% formic acid, and applied to a reverse-phase liquid chromatographer [Acquity UPLC M-Class System (Waters Corporation, USA)], coupled to a Synapt G2-Si mass spectrometer (Waters Corporation, USA). Data-independent acquisition (DIA) strategy was used with ion mobility separation (high-definition data-independent mass spectrometry; HDMSE). Peptides were loaded onto a first-dimension chromatography on an M-Class BEH C18 Column (130 Å, 5 µm, 300 µm X 50 mm, Waters Corporation, USA) and eluted by discontinuous fractionation steps (13%, 18%, and 50% acetonitrile). After each step, the peptide loads were directed to a second-dimension chromatography on a nanoACQUITY UPLC HSS T3 Column (1.8 µm, 75 µm X 150 mm; Waters Corporation, USA) and eluted with acetonitrile [7 to 40% gradient (v/v), for 54 min, at a flow rate of 0.4 µL/min] into a Synapt G2-Si. MS/MS analysis was performed using nano-electrospray ionization in positive ion mode [nanoESI (+)] and a NanoLock Spray ionization source (Waters Corporation, UK). The lock mass channel was sampled every 30 seconds. Mass spectrometer calibration was performed with a [Glu1]-fibrinopeptide B human (Glu-Fib) (Sigma-Aldrich, USA) solution with an MS/MS spectrum reference from the NanoLock Spray source. Samples were run in technical duplicates of biological triplicates.

### Database search and quantification

HDMSE raw data was imported to Progenesis QI for Proteomics software, version 4.0 (Waters Corporation, USA), where protein identification and quantification was processed using dedicated algorithms (including Apex3D, peptide eD, and ion accounting informatics). Using default parameters for ion accounting and quantitation, peptide identification was carried out using UniProt’s human proteomic database (version 2018/09, reviewed and non-reviewed). During the database queries, the databases used were reversed “on the fly” and appended to the original database to assess the false positive identification rate. Parameters for peptide identification: 1) in trypsin digestion; 2) variable modifications by oxidation (M) and fixed modification by carbamidomethyl (C); and 3) FDR less than 1%. Identifications not satisfying these criteria were not included in the analysis. Relative quantitation of proteins was done with Hi-N method (N=3). The quantitative analysis was carried out on the log_2_ values of the measured intensities. The mass spectrometry proteomics data have been deposited to the ProteomeXchange Consortium via the PRIDE (69) partner repository with the dataset identifiers PXD027369 and PXD037814.

### In silico analysis

The fold change in the abundance of each protein was determined as a ratio of the intensity values from LSD-exposed and control organoids in each batch. Statistical analysis, aiming for the identification of the DAPs between control and LSD-exposed organoids, was carried out in Perseus software (version 2.0.7.0, Max-Planck-Gesellschaft, München) (70,71). Fold change values were log_2_ transformed, and a one sample t-test was performed to evaluate whether the abundance of each protein was significantly changed. Significant hits (p < 0.05) were considered DAPs induced by LSD and subjected to the bioinformatic analysis.

For biological process and pathway enrichment analysis, given DAPs were launched on the Metascape platform (http://metascape.org/) (72). The analysis considered the following ontology databases: Reactome Gene Sets and KEGG Pathway. Terms with a minimum overlap of 3.0, P value cutoff 0.01, and minimum enrichment of 1.5 were collected and groupe into clusters. For PPI analysis, a network with given DAPs was constructed using the STRING 11.5 database (https://string-db.org/) (73) with a confidence score > 0.7 and visualized in Cytoscape 3.9.1 (74). Molecular Complex Detection (MCODE; Network scoring: loops not included, degree cutoff = 2; Cluster finding: haircut parameter, node score cutoff = 0.2, k-core = 2, and max. depth = 100) and CytoHubba (degree ranking method) (75) plugins of Cytoscape software were applied to obtain clusters and hub proteins, respectively. The functional annotation of the clusters was obtained in the version 2021 of the Database for Annotation, Visualization, and Integrated Discovery (DAVID 2021) (https://david.ncifcrf.gov/). The thresholds were count = 2 and EASE = 0.1. For an in-depth study of possibly affected pathways, schemes were constructed, showing where the given DAPs would act in each pathway. The pathways were chosen based on previous analysis. Some schemes were constructed based on the literature, and for others, the Kyoto Encyclopedia of genes and genomes (KEGG) pathway enrichment in DAVID 2021 was employed.

In the comparative analyses, we contrasted the data obtained from the analysis of this paper (100 nM LSD) with the data from Ornelas et al.’s study (23) (10 nM LSD). To assess the similarity level in the ontology, we employed Metascape to perform an overlap correlation. Furthermore, we compared the top 20 most enriched terms in both analyses, aiming to identify the commonly modulated processes.

### hiPSC-derived human brain spheroids

hiPSCs were differentiated into NSCs using PSC Neural Induction Medium (Thermo Fisher Scientific, USA), following the manufacturer’s guidelines. Media was changed every other day until day 7. At this stage, NSCs were split and expanded in neural expansion medium, composed of a 1:1 ratio of Advanced DMEM/F12 and Neurobasal medium supplemented with neural induction supplement (Thermo Fisher Scientific, USA). To form spheroids (76), 8.0 × 10^4^ NSCs in 150 µL were seeded into each well of round-bottom ultra-low attachment 96-well plates (Corning, USA) and centrifuged at 300 g for 3 minutes to facilitate settling. After 72 hours, the medium was switched to a differentiation medium (1:1 mixture of Neurobasal medium and DMEM, enriched with B27 and N2 supplements), with subsequent media replacements every other day. By day 10 of aggregation, spheroids were ready for the neurite outgrowth assay.

### Neurite outgrowth assay

Each well in 96-well plates (Perkin Elmer, USA) was seeded with a single spheroid, coated with 100 μg/ml poly-ornithine and 20 μg/ml laminin. After settling for 24 hours, spheroids were exposed to 10 and 100 nM LSD for an additional 24 hours. Subsequently, each one of them was submitted to both immunofluorescence and Sholl analysis.

### Immunofluorescence in plated brain spheroids

Following a 15-minute fixation in a 4% paraformaldehyde solution, brain spheroids were washed with PBS and then permeabilized using 0.3% Triton X-100 in PBS for 15 minutes. Subsequently, a blocking step was carried out by incubating in a solution containing 3% fetal calf serum in PBS for 1 hour. Following blocking, they were subjected to overnight incubation at 4°C with the primary antibody TUJ1 (Neuromics, USA) at a dilution of 1:2000 in the blocking solution. After incubation, spheroids underwent three washes with PBS and were re-blocked using 3% fetal calf serum for 20 minutes. Next, they were incubated with the secondary antibody, specifically goat anti-mouse Alexa Fluor 594 (Thermo Fisher Scientific, USA) at a dilution of 1:400, for 60 minutes. To visualize nuclei, a counterstain using 0.5 μg/mL DAPI was performed for 5 minutes. Finally, a solution composed of a 1:1 mixture of glycerol and PBS was added to the plates, and images were captured using a 20x objective on an Agilent BioTek Cytation 1 imaging system.

### Sholl analysis in plated brain spheroids

This study used Sholl analysis to assess neurite outgrowth in plated spheroids. The Neuroanatomy plugin within Fiji ImageJ software (version 1.54f) was employed for this purpose (77). The analysis involved setting up 30 circles originated from the spheroid body, each spaced at 31.5 µm intervals. Neurite outgrowth was quantified by analyzing the mean crossings in groups of five circles. Statistical analysis was performed using two-way ANOVA, complemented by Dunnets’s post-hoc test for multiple comparisons. A visual representation of the Sholl analysis can be found in Figure S1.

## Author Contributions

S.K.R., and M.N.C. conceived and designed the study; M. N. C. performed all cell cultures; J.M.N. carried out the proteomics and assisted in the interpretation; D.M.S. supervised proteomic experiments and data interpretation; M.N.C., L.G.S., and P.T. analyzed the majority of the proteomic data; L. G. S. and I. D. undertook conducted the experiments related to neurite outgrowth; M.N.C. drafted the manuscript and figures; K. K. undertook the characterization of cerebral organoids; S.K.R coordinated all the study; A.F. contributed to the conceptual framework of the proposal and provided critical review and feedback on the manuscript; All authors reviewed and approved the final manuscript prior to submission.

## Notes

The authors declare no competing interest.

## ACKNOWLEDGEMENTS

We gratefully acknowledge the Brazilian National Council for Scientific and Technological Development (CNPq), the Pioneering Science Initiative, and the D’Or Institute for Research and Education (IDOR) for the research fellowships provided during the development of this work. We also acknowledge the technical support provided by Beatriz Luzia de Mello Lima Guimarães, Gabriela Lopes Vitória, Ismael Carlos da Silva Gomes, and Jhonata de Sousa do Nascimento. Our appreciation extends to Joana Cardoso for her contribution to the immunofluorescence experiments and the acquisition of corresponding images. We also thank Michele do Nascimento Costa for her assistance in the editing of images. Lastly, we would like to underscore the use of the GPT-3.5 language model, developed by OpenAI, as a valuable tool for spelling correction. Importantly, its application in this specific context had no significant impact on the content or substance of the text.

## ABBREVIATIONS

α-SNAP: soluble NSF attachment protein α
5-HT: 5-hydroxytryptamine
ACTN: actinin
AP2: adaptor protein 2 complex
ATP: adenosine triphosphate
BDNF: brain-derived neurotrophic factor
BSA: bicinchoninic acid
BSA: bovine serum albumin
cDNA: Complementary DNA
CME: clathrin-mediated endocytosis
CNS: central nervous system
CPNE1: copine-1
DAP: differentially abundant proteins
DAPI: 4’,6-Diamidino-2-Phenylindole
DAVID: Database for Annotation Visualization and Integrated Discovery
DIA: data-independent acquisition
DMEM/F12: Dulbecco’s Modified Eagle Medium/Nutrient Mixture F-12
DNA: deoxyribonucleic acid
DTT: dithiothreitol
ERM: ezrin, radixin or moesin
F2: coagulation factor II
F2R: coagulation factor II receptor
FC: fold change
FDR: false discovery rate
fMRI: functional magnetic resonance imaging
FN1: fibronectin 1
Gα12,13: G protein subunit alpha 12, 13
GABA: *gamma*-aminobutyric acid
GF: growth factor
GPCR: G protein-coupled receptor
GTPase: guanosine triphosphatases
Glu-Fib: [Glu1]-fibrinopeptide B human
HDMSE: high-definition data-independent mass spectrometry
hESC: human embryonic stem cell
HGNC: HUGO Gene Nomenclature Committee
hiPSC: human induced pluripotent stem cell
HUGO: Human Genome Organization
ITG: integrin
KEGG: Kyoto Encyclopedia of Genes and Genomes
KSR: KnockOut Serum Replacement
LC-MS/MS: liquid chromatography tandem mass spectrometry
LSD: lysergic acid diethylamide
M-MLV: moloney murine leukemia virus
MAP2: microtubule-associated protein 2
MCODE: Molecular Complex Detection
MEM-NEAA: minimum essential media-nonessential amino acids
MLC: myosin light chain
mRNA: messenger ribonucleic acid
mTOR: mammalian target of rapamycin
mTeSR1: modified TeSR1 medium
NADH: nicotinamide adenine dinucleotide
NSF: N-ethylmaleimide-sensitive fusion protein
NSC: neural stem cell
NT: neurotransmitter
OGA: O-GlcNAcase
PBS: phosphate-buffered saline
PCR: polymerase chain reaction
PFA: paraformaldehyde
Pen-Strep: Penicillin-Streptomycin
PPI: protein-protein interaction
Rho: Rat sarcoma virus homolog
ROCK: Rho kinase
ROCKi: Rho-associated protein kinase inhibitor
RIM: Rab-interacting molecule
RNA: ribonucleic acid
SNAP: soluble NSF attachment proteins
SNARE: soluble NSF adaptor protein receptor
RPM: rotation per minute
RTK: Receptor Tyrosine Kinase
STRING: Search Tool for the Retrieval of Interacting Genes
t-SNARE: target SNARE
TCA: tricarboxylic acid
Taq: *Thermus aquaticus*
TeSR1: serum-free medium for human pluripotent stem cells
TrkB: tropomyosin receptor kinase B
VAMP: vesicle-associated membrane protein
VGCC: voltage-gated calcium channel
V-ATPase: vacuolar ATPase
v-SNARE: vesicle SNARE.

