## Supplemental Information for "LSD Modulates Proteins Involved in Cell Proteostasis, Energy Metabolism and Neuroplasticity in Human Cerebral Organoids"

**Table S1.** Protein analyses from cerebral organoids exposed to 100 nM LSD (L100) and control samples (CTRL) in distinct experimental batches (n=3). The hits are sorted based on the t-test results, and those with p-values less than 0.05 have their gene symbols highlighted in bold. Standard deviation (SD) was used as a measure of dispersion.

| L100/ CTRL1 | L100/ CTRL2 | L100/ CTRL3 | Mean | SD | T-test p-value | Accession | Gene Symbol |
|---|---|---|---|---|---|---|---|
| -0.7798 | -0.7820 | *-0.7593* | -0.7737 | 0.0125 | 0.0001 | Q9Y4B5;J3QLE1 | **MTCL1** |
| -1.4369 | -1.4033 | *-1.4616* | -1.4339 | 0.0293 | 0.0001 | Q9UPP1;H0Y3N9;H0Y589;B0QZE1;B0QZZ2;B0QZZ3;B0QZZ4;Q5JPR8 | **PHF8** |
| 1.3685 | 1.3355 | *1.3909* | 1.3650 | 0.0279 | 0.0001 | Q86YV5 | **PRAG1** |
| -1.8061 | -1.7799 | *-1.7263* | -1.7708 | 0.0407 | 0.0002 | Q9Y3P9;B5MCD9;C9JGR5 | **RABGAP1** |
| -0.2818 | -0.2669 | *-0.2769* | -0.2752 | 0.0076 | 0.0003 | P29966 | **MARCKS** |
| 2.6823 | 2.6477 | *2.4125* | 2.5809 | 0.1468 | 0.0011 | A0A0C4DGK5;H0YJS2;H0YK02;Q86TV6 | **TTC7B** |
| -1.2436 | -1.4225 | *-1.3661* | -1.3441 | 0.0914 | 0.0015 | Q86YS3;K7EL58 | **RAB11FIP4** |
| -0.3267 | -0.3724 | *-0.3770* | -0.3587 | 0.0278 | 0.0020 | Q8TD57;Q5T7N2 | **DNAH3** |
| 0.9624 | 0.8812 | *1.0343* | 0.9593 | 0.0766 | 0.0021 | Q7Z7G8 | **VPS13B** |
| 0.5395 | 0.6117 | *0.6303* | 0.5938 | 0.0480 | 0.0022 | Q96HN2;H0Y8B3;C9K0S0 | **AHCYL2** |
| 2.4216 | 2.8761 | *2.6512* | 2.6496 | 0.2273 | 0.0024 | Q9GZV4;C9J7B5;F8WCJ1 | **EIF5A2** |
| 1.8716 | 2.2188 | *2.1693* | 2.0866 | 0.1878 | 0.0027 | Q5JQF8 | **PABPC1L2A** |
| -0.5160 | -0.5385 | *-0.4505* | -0.5017 | 0.0457 | 0.0028 | P49755;G3V2K7 | **TMED10** |
| -0.3778 | -0.3283 | *-0.3156* | -0.3406 | 0.0329 | 0.0031 | Q3ZCM7;A0A075B736;Q5SQY0 | **TUBB8** |
| 0.5358 | 0.5642 | *0.4656* | 0.5219 | 0.0507 | 0.0031 | Q8IX07;A0A087WWQ0;A0A087WZP1 | **ZFPM1** |
| -0.5846 | -0.5062 | *-0.4833* | -0.5247 | 0.0531 | 0.0034 | Q9HAV0;C9JD14;H7C5J5 | **GNB4** |
| -0.6978 | -0.5774 | *-0.6898* | -0.6550 | 0.0673 | 0.0035 | P16949;B5BU83;A2A2D0 | **STMN1** |
| -0.4954 | -0.4278 | *-0.5271* | -0.4835 | 0.0507 | 0.0037 | P52209;K7ELN9;K7EM49;K7EMN2 | **PGD** |
| -0.9111 | -0.8334 | *-1.0374* | -0.9273 | 0.1029 | 0.0041 | Q14194;E9PD68 | **CRMP1** |
| -0.7469 | -0.6098 | *-0.7435* | -0.7001 | 0.0782 | 0.0041 | Q93045;E5RGX5;Q6ZRC1 | **STMN2** |
| 0.9643 | 0.7679 | *0.8767* | 0.8696 | 0.0984 | 0.0042 | M0QYA8;Q8IV63;M0R073;M0QXD7;M0QYG0;M0QZ79;M0R025 | **VRK3** |
| -1.2854 | -1.2847 | *-1.5664* | -1.3788 | 0.1624 | 0.0046 | Q12802;H0YMW2;A0A087WTD7 | **AKAP13** |
| -1.1457 | -1.3664 | *-1.4477* | -1.3200 | 0.1563 | 0.0046 | O75410;R4GMT7;E7ET87;E7EVI4; | **TACC1** |

| L100/ CTRL1 | L100/ CTRL2 | L100/ CTRL3 | Mean | SD | T-test p-value | Accession | Gene Symbol |
|---|---|---|---|---|---|---|---|
| | | | | | | H0YAY0;E5RFM9;E5RJG6 | |
| -0.8805 | -0.9217 | *-1.1045* | -0.9689 | 0.1193 | 0.0050 | Q8NEZ3;D6R9P6;D6RE75;D6RAI4 | **WDR19** |
| -1.2439 | -1.0776 | *-1.3955* | -1.2390 | 0.1590 | 0.0054 | P98164 | **LRP2** |
| -0.4805 | -0.5801 | *-0.6213* | -0.5607 | 0.0724 | 0.0055 | O15061;A0A075B7B1;C9JIE4 | **SYNM** |
| -0.2245 | -0.2913 | *-0.2704* | -0.2621 | 0.0342 | 0.0056 | Q8WXA3;H0YD93 | **RUFY2** |
| -0.2799 | -0.2137 | *-0.2511* | -0.2483 | 0.0332 | 0.0059 | Q9H0Q0;C9IYV6;C9JPE5 | **FAM49A** |
| 2.2756 | 1.7846 | *2.2917* | 2.1173 | 0.2882 | 0.0061 | P47813;O14602;X6RAC9;A6NJH9 | **EIF1AX** |
| -0.1577 | -0.1503 | *-0.1199* | -0.1426 | 0.0200 | 0.0065 | Q9BZF9;F5H2B9;H0YNH8 | **UACA** |
| -0.3006 | -0.2269 | *-0.2507* | -0.2594 | 0.0376 | 0.0069 | A0A0B4J207;P21108 | **PRPS1L1** |
| -1.5736 | -1.8667 | *-1.3962* | -1.6122 | 0.2376 | 0.0072 | A0A0A0MTT5 | **CTAG1A** |
| -0.6085 | -0.6209 | *-0.7910* | -0.6734 | 0.1020 | 0.0076 | O95989 | **NUDT3** |
| -0.7770 | -0.7051 | *-0.5718* | -0.6847 | 0.1041 | 0.0076 | Q15029;K7EP67;K7EJ74;K7EIT3 | **EFTUD2** |
| -1.2941 | -0.9482 | *-1.1143* | -1.1189 | 0.1730 | 0.0079 | Q9NQW7;Q5T6H7;Q5T6H2 | **XPNPEP1** |
| 0.8105 | 0.9075 | *1.1093* | 0.9424 | 0.1524 | 0.0086 | Q13838;F6WLT2;A0A0A0MT12;F6TRA5;F6S4E6;A0A140T9X9;A0A140TA18;F6UJC5;H0Y400;H0YCC6;A0A0G2JJL7;F6QYI9;F6R6M7;A0A0G2JHN7;F6S2B7;F6U6E2;A0A140T9N3;F6SXL5 | **DDX39B** |
| -0.4441 | -0.4177 | *-0.3213* | -0.3943 | 0.0646 | 0.0088 | Q01581;D6RIW1 | **HMGCS1** |
| -0.5476 | -0.3991 | *-0.5283* | -0.4917 | 0.0808 | 0.0089 | P21359;J3KSB5;H0Y465 | **NF1** |
| -0.4551 | -0.3266 | *-0.3860* | -0.3893 | 0.0643 | 0.0090 | P05062;A0A087WXX2 | **ALDOB** |
| 0.9338 | 0.8608 | *0.6703* | 0.8216 | 0.1361 | 0.0090 | Q15572 | **TAF1C** |
| 1.2416 | 1.1363 | *0.8858* | 1.0879 | 0.1828 | 0.0093 | C9K025;P18077;F8WB72;F8WBS5 | **RPL35A** |
| 0.4567 | 0.3248 | *0.4128* | 0.3981 | 0.0672 | 0.0094 | Q562R1 | **ACTBL2** |
| -0.7637 | -0.6317 | *-0.5426* | -0.6460 | 0.1112 | 0.0097 | Q07666 | **KHDRBS1** |
| 1.9997 | 1.5224 | *1.4784* | 1.6669 | 0.2891 | 0.0099 | Q9H0S4;F5H1N9 | **DDX47** |
| -0.3336 | -0.2788 | *-0.2348* | -0.2824 | 0.0495 | 0.0101 | Q5VTE0;P68104;A0A087WV01;Q5JR01;A6PW80;A0A0A0MST8;C9JRP1;E9PFN4;H0YBD9;H0YC27;H0YC67;H7C3C4;P12757;Q9Y6M7 | **EEF1A1P5** |
| 0.7298 | 0.5776 | *0.8248* | 0.7107 | 0.1247 | 0.0101 | Q9H0W5 | **CCDC8** |

| L100/ CTRL1 | L100/ CTRL2 | L100/ CTRL3 | Mean | SD | T-test p-value | Accession | Gene Symbol |
|---|---|---|---|---|---|---|---|
| -0.5944 | -0.8347 | *-0.6663* | -0.6985 | 0.1233 | 0.0102 | B1AJZ9;H0Y3C6; H0YEU3;Q5JYW1 | **FHAD1** |
| 0.4566 | 0.3704 | *0.3223* | 0.3831 | 0.0681 | 0.0104 | Q6NZY4 | **ZCCHC8** |
| -0.7414 | -0.6456 | *-0.9146* | -0.7672 | 0.1364 | 0.0104 | O60502;H7C3X0 | **OGA** |
| -2.1961 | -1.5234 | *-1.9479* | -1.8891 | 0.3402 | 0.0106 | A0A075B6H4;Q9H8E8 | **KAT14** |
| 0.4192 | 0.3318 | *0.4792* | 0.4101 | 0.0741 | 0.0107 | A0A087WWI6;P61962 | **DCAF7** |
| 1.4409 | 1.6954 | *1.1715* | 1.4360 | 0.2620 | 0.0109 | P23142;B1AHM7;B1AHM9;H7C1M6;B1AHM6;B1AHM8;B1AHN3 | **FBLN1** |
| -0.7429 | -0.7821 | *-1.0301* | -0.8517 | 0.1557 | 0.0110 | A0A0A0MSK6;Q13017;G3V444;G3V5I7 | **ARHGAP5** |
| -0.4576 | -0.5244 | *-0.3592* | -0.4471 | 0.0831 | 0.0113 | O75197;E9PHY1 | **LRP5** |
| 2.0529 | 2.0616 | *2.8224* | 2.3123 | 0.4418 | 0.0119 | E7EVR1;Q9NR11 | **ZNF302** |
| -0.4569 | -0.4190 | *-0.3098* | -0.3952 | 0.0764 | 0.0122 | P61158;B4DXW1;F8WDR7;F8WE84;F8WEW2 | **ACTR3** |
| -1.8096 | -1.3378 | *-1.2855* | -1.4776 | 0.2887 | 0.0125 | Q86UP2;G3V4Y7;B7Z6P3;G3V5G2;H0YJV5;G3V5P0 | **KTN1** |
| 0.8956 | 0.8692 | *1.2211* | 0.9953 | 0.1960 | 0.0127 | E5RG59 | **ZNF395** |
| 3.8070 | 5.4424 | *4.0729* | 4.4408 | 0.8776 | 0.0128 | P21579;J3KQA0; C9JX50;F8VXH0; F8VYH8;F8VZY3;F8W1U9 | **SYT1** |
| -0.4026 | -0.4285 | *-0.2894* | -0.3735 | 0.0740 | 0.0128 | P35998;A0A1W2PQS1;C9JLS9 | **PSMC2** |
| 1.0342 | 1.5550 | *1.4159* | 1.3351 | 0.2696 | 0.0133 | Q96K17;E9PL10 | **BTF3L4** |
| 0.8333 | 1.2381 | *1.1822* | 1.0845 | 0.2193 | 0.0134 | Q5T1J5;Q9Y6H1 | **CHCHD2P9** |
| -0.2857 | -0.3451 | *-0.4288* | -0.3532 | 0.0719 | 0.0135 | Q99627;E9PGT6; H7C3S9 | **COPS8** |
| -1.1987 | -1.3824 | *-0.9082* | -1.1631 | 0.2391 | 0.0138 | P43246;V9H019; A0A2R8Y7S8;V9H0B2;V9H015;C9J809;A0A2R8Y713 | **MSH2** |
| -2.7149 | -1.7863 | *-2.2149* | -2.2387 | 0.4647 | 0.0141 | P61244;G3V5L1 | **MAX** |
| 0.1838 | 0.1191 | *0.1565* | 0.1531 | 0.0325 | 0.0146 | A0A1C7CYX1;Q6PJG2;A0A0A0MSU2 | **ELMSAN1** |
| -1.7467 | -2.7124 | *-2.2809* | -2.2467 | 0.4838 | 0.0151 | Q13310;B1ANR0;H0Y5F5;B1ANR1;H0YCC8;H0YEQ8;H0YEU6 | **PABPC4** |
| 0.5617 | 0.5183 | *0.3638* | 0.4813 | 0.1040 | 0.0152 | Q8N1F7;H3BVG0;H3BV15;H3BM93;H3BP95;H3BRI8;H3BMX0;H3BNG7;H3BNN5;H3BPA9 | **NUP93** |

| L100/ CTRL1 | L100/ CTRL2 | L100/ CTRL3 | Mean | SD | T-test p-value | Accession | Gene Symbol |
|---|---|---|---|---|---|---|---|
| -1.0733 | -1.0077 | *-1.4814* | -1.1874 | 0.2567 | 0.0152 | A8MT87;P11086 | **PNMT** |
| -2.2950 | -2.1244 | *-3.1480* | -2.5224 | 0.5484 | 0.0154 | Q6UN15;H0Y8P7 | **FIP1L1** |
| 0.4703 | 0.3425 | *0.5374* | 0.4501 | 0.0991 | 0.0158 | K7EP73;O14556 | **GAPDHS** |
| -0.5482 | -0.3869 | *-0.6056* | -0.5136 | 0.1134 | 0.0159 | Q15126 | **PMVK** |
| 0.6650 | 1.0247 | *0.7863* | 0.8253 | 0.1830 | 0.0160 | O95359;E7EMZ9;E9PBC6;D6RAA5;Q4VXL4;Q4VXL8;H0Y9Y7 | **TACC2** |
| 1.1650 | 1.7937 | *1.7343* | 1.5643 | 0.3471 | 0.0160 | P67936;A0A2R8YGX3;A0A2R8YE05;K7EP68;A0A2R8YHD2;K7EMU5;K7EPV9 | **TPM4** |
| -0.7073 | -0.4703 | *-0.5173* | -0.5649 | 0.1255 | 0.0161 | A0A0A0MRZ4;Q15025 | **TNIP1** |
| -1.2733 | -2.0218 | *-1.8193* | -1.7048 | 0.3872 | 0.0168 | A0A0B4J239;F5GXX5;F5H895;P61803 | **DAD1** |
| 2.1406 | 1.3854 | *1.6030* | 1.7097 | 0.3887 | 0.0168 | O94885 | **SASH1** |
| -0.8917 | -0.6344 | *-0.5947* | -0.7069 | 0.1612 | 0.0169 | Q9C0C9;K7ES11;K7EQ12 | **UBE2O** |
| -1.8567 | -2.4343 | *-1.5576* | -1.9495 | 0.4457 | 0.0170 | A0A0C4DGQ1;Q9BUG6 | **ZSCAN5A** |
| 1.2747 | 0.8003 | *1.1522* | 1.0757 | 0.2462 | 0.0170 | Q86UE4;E5RJU9;H0YBE0 | **MTDH** |
| 0.7354 | 0.6954 | *0.4662* | 0.6323 | 0.1453 | 0.0171 | Q9BTV4 | **TMEM43** |
| -0.6901 | -0.6022 | *-0.9328* | -0.7417 | 0.1712 | 0.0173 | Q5W041 | **ARMC3** |
| -0.3943 | -0.6158 | *-0.6204* | -0.5435 | 0.1292 | 0.0183 | P02792 | **FTL** |
| 0.7263 | 0.4486 | *0.5736* | 0.5828 | 0.1391 | 0.0185 | A0A087WZE9;Q15651 | **HMGN3** |
| 1.2318 | 1.2692 | *1.8578* | 1.4529 | 0.3511 | 0.0189 | P52737;C9J2Q9;C9JJK8 | **ZNF136** |
| -0.1260 | -0.1279 | *-0.1896* | -0.1478 | 0.0361 | 0.0194 | Q12874 | **SF3A3** |
| 0.3334 | 0.3034 | *0.2023* | 0.2797 | 0.0687 | 0.0195 | Q1X8D7;J3QSG3;H3BNV0;H3BQI9;H3BRB2;H3BRP7;H3BSQ6 | **LRRC36** |
| -1.2240 | -0.7448 | *-0.9601* | -0.9763 | 0.2400 | 0.0196 | P08574 | **CYC1** |
| 0.3013 | 0.2641 | *0.1813* | 0.2489 | 0.0614 | 0.0197 | Q6W2J9;H7BYY2 | **BCOR** |
| -0.3220 | -0.2527 | *-0.1953* | -0.2567 | 0.0635 | 0.0198 | Q9UBN7;A0A2R8YDE6;A0A2R8YG4G2 | **HDAC6** |
| -1.0474 | -0.7126 | *-1.1893* | -0.9831 | 0.2447 | 0.0200 | Q8IY63 | **AMOTL1** |
| -2.4077 | -2.1844 | *-3.4357* | -2.6759 | 0.6674 | 0.0201 | Q9NRX4 | **PHPT1** |
| -0.7735 | -1.0970 | *-1.3010* | -1.0572 | 0.2660 | 0.0205 | O94826 | **TOMM70** |
| -0.7808 | -1.0513 | *-1.3077* | -1.0466 | 0.2635 | 0.0205 | Q58EX2;H7C2P2 | **SDK2** |
| -0.7384 | -1.0036 | *-0.6135* | -0.7851 | 0.1992 | 0.0208 | Q15181;Q5SQT6 | **PPA1** |
| -2.1608 | -3.4305 | *-3.5673* | -3.0528 | 0.7756 | 0.0208 | Q14669;C9JLD7;C9JLJ5;C9JSX9;F8W9P3;G5E9G6 | **TRIP12** |

| L100/ CTRL1 | L100/ CTRL2 | L100/ CTRL3 | Mean | SD | T-test p-value | Accession | Gene Symbol |
|---|---|---|---|---|---|---|---|
| -1.2341 | -0.9063 | *-0.7543* | -0.9649 | 0.2452 | 0.0209 | F5H039;Q9NQX3;G3V582;H0YJR5;H0YJ30 | **GPHN** |
| -0.6420 | -0.4614 | *-0.3967* | -0.5000 | 0.1271 | 0.0209 | O15540 | **FABP7** |
| 0.8866 | 0.9969 | *1.4222* | 1.1019 | 0.2828 | 0.0213 | A0A0A0MSK5;J3KN66;Q5JTV8;H0Y4R4 | **TOR1AIP1** |
| -1.2211 | -1.0375 | *-0.7167* | -0.9918 | 0.2553 | 0.0214 | A0A0A0MSU4;O94911 | **ABCA8** |
| -0.7398 | -0.4745 | *-0.4947* | -0.5697 | 0.1477 | 0.0217 | P53355;F8WCQ3 | **DAPK1** |
| -0.2427 | -0.1417 | *-0.2098* | -0.1981 | 0.0515 | 0.0218 | A1L4K2;B5BUB8;P45983;A6NF29;C9J762 | **MAPK8** |
| 1.0598 | 1.3333 | *0.7805* | 1.0579 | 0.2764 | 0.0220 | Q8IXT1;E9PMA7;E9PN94 | **DDIAS** |
| -0.9871 | -1.6846 | *-1.5373* | -1.4030 | 0.3676 | 0.0221 | Q7Z7L1;K7EIM3;C9J902;C9JDG6;C9JUT2;K7ER38;K7EKT7;K7ES87 | **SLFN11** |
| 1.8033 | 2.7146 | *3.0990* | 2.5390 | 0.6655 | 0.0221 | Q9P0J1;E5RI96;E5RIE5;E5RIV4 | **PDP1** |
| -0.7686 | -0.4579 | *-0.7500* | -0.6588 | 0.1743 | 0.0225 | P32969;A0A2R8Y5Y7;D6RAN4;E7ESE0;H0Y9V9;H0Y9R4 | **RPL9** |
| -0.9934 | -1.4106 | *-1.7300* | -1.3780 | 0.3694 | 0.0231 | M0R2K1;Q9H0K4 | **RSPH6A** |
| 0.4820 | 0.5023 | *0.7625* | 0.5823 | 0.1564 | 0.0232 | Q7L2H7;J3KNJ2 | **EIF3M** |
| -0.9814 | -1.2353 | *-0.7087* | -0.9751 | 0.2634 | 0.0235 | Q6Q0C0;H3BR17 | **TRAF7** |
| 0.5053 | 0.7922 | *0.8850* | 0.7275 | 0.1980 | 0.0238 | Q9UL01;A0A2R8Y6J1;A0A2U3TZJ0 | **DSE** |
| -0.7060 | -0.4099 | *-0.5268* | -0.5476 | 0.1491 | 0.0238 | Q8N157 | **AHI1** |
| -0.4038 | -0.7022 | *-0.6560* | -0.5873 | 0.1606 | 0.0240 | P29762;B5MCB5 | **CRABP1** |
| 0.8945 | 0.8236 | *1.3388* | 1.0190 | 0.2792 | 0.0241 | O94804 | **STK10** |
| 1.2120 | 1.4274 | *0.8049* | 1.1481 | 0.3162 | 0.0244 | Q13098;A0A096LP07;A0A096LPJ3;A8K070;C9JFE4;J3KRJ4;J3KSA5;J3KTB0;J3QLT0;J3QS88;J3KRE8;J3QL53;J3QLE8;J3QQX0;J3QS84 | **GPS1** |
| -0.4048 | -0.2299 | *-0.3742* | -0.3363 | 0.0934 | 0.0248 | Q9NYU2;H7BZG0 | **UGGT1** |
| -0.4875 | -0.6732 | *-0.3893* | -0.5167 | 0.1442 | 0.0250 | P68402;J3KNE3 | **PAFAH1B2** |
| -0.7319 | -0.5610 | *-0.9823* | -0.7584 | 0.2119 | 0.0251 | Q96D15;M0QZH0 | **RCN3** |
| -0.3956 | -0.5173 | *-0.6937* | -0.5355 | 0.1499 | 0.0251 | Q5VTR2;C9J0A5;C9JXC9 | **RNF20** |
| -2.9407 | -3.4723 | *-1.9377* | -2.7836 | 0.7793 | 0.0251 | Q6PEY2 | **TUBA3E** |
| -0.7622 | -0.6146 | *-1.0635* | -0.8134 | 0.2288 | 0.0254 | O75347;E5RHG6;E5RJD8;E5RIW3;E5RIX8 | **TBCA** |

| L100/ CTRL1 | L100/ CTRL2 | L100/ CTRL3 | Mean | SD | T-test p-value | Accession | Gene Symbol |
|---|---|---|---|---|---|---|---|
| -0.3655 | -0.3928 | *-0.2216* | -0.3267 | 0.0920 | 0.0254 | Q07864;F5H1D6; F5H7E4 | **POLE** |
| 0.4657 | 0.3330 | *0.2674* | 0.3554 | 0.1010 | 0.0259 | Q5S007;E9PC85 | **LRRK2** |
| 1.0700 | 1.3749 | *0.7659* | 1.0703 | 0.3045 | 0.0259 | F5H5K1;J3KTP0; J3QSU1;Q96QE4 | **LRRC37B** |
| -1.0424 | -0.9570 | *-0.5821* | -0.8605 | 0.2449 | 0.0260 | B5BUI8;P51452; K7ES89 | **DUSP3** |
| -0.4104 | -0.6551 | *-0.7454* | -0.6036 | 0.1733 | 0.0264 | Q5T5U3;A0A1B0 GV73;E7ESW5;F 8W9U9 | **ARHGAP21** |
| -1.6465 | -1.1428 | *-2.0795* | -1.6230 | 0.4688 | 0.0267 | Q6UWP8;K7ESC 4 | **SBSN** |
| -0.3780 | -0.6271 | *-0.4064* | -0.4705 | 0.1364 | 0.0269 | P08237;A0A2R8 Y891;F8VNX2;F8 VZQ1;F8VP00;F8 VX13;F8VSL1;F8 VW30 | **PFKM** |
| -1.5819 | -1.8298 | *-2.7084* | -2.0400 | 0.5919 | 0.0269 | Q04917 | **YWHAH** |
| -0.5982 | -1.0557 | *-0.7520* | -0.8020 | 0.2328 | 0.0270 | Q96Q89;A0A0A0 MSJ5 | **KIF20B** |
| -0.5069 | -0.2768 | *-0.4051* | -0.3963 | 0.1153 | 0.0271 | P54920;M0R0Y2; M0R2M1;M0R02 7;M0R0I4;M0R21 3;M0R058 | **NAPA** |
| -0.4464 | -0.7013 | *-0.4273* | -0.5250 | 0.1530 | 0.0272 | P61981 | **YWHAG** |
| -0.8594 | -0.7008 | *-0.4676* | -0.6760 | 0.1971 | 0.0272 | O14950;J3QRS3; P19105 | **MYL12B** |
| 0.8149 | 1.4998 | *1.2210* | 1.1785 | 0.3444 | 0.0273 | P56545;Q5SQP8 | **CTBP2** |
| -0.6057 | -0.3912 | *-0.3612* | -0.4527 | 0.1333 | 0.0277 | P82909 | **MRPS36** |
| 1.0427 | 0.8589 | *0.5609* | 0.8208 | 0.2432 | 0.0280 | Q9NVP4 | **DZANK1** |
| -1.1181 | -0.5983 | *-0.9239* | -0.8801 | 0.2627 | 0.0284 | P63167;F8VRV5; F8VXI7;F8VXL2 | **DYNLL1** |
| -0.3830 | -0.2048 | *-0.3007* | -0.2961 | 0.0891 | 0.0289 | P12931;J3QRU1; P07947 | **SRC** |
| -0.6502 | -0.3976 | *-0.3976* | -0.4818 | 0.1458 | 0.0292 | Q9P0X4 | **CACNA1I** |
| 0.7060 | 0.7653 | *0.4101* | 0.6271 | 0.1903 | 0.0293 | P35913;H7C4P9 | **PDE6B** |
| -0.7156 | -0.5740 | *-0.3783* | -0.5560 | 0.1694 | 0.0296 | Q09028;H0YF10; H0YCT5;C9JPP3 | **RBBP4** |
| 2.2207 | 3.5459 | *4.2115* | 3.3261 | 1.0134 | 0.0296 | O00411;K7EMH3 | **POLRMT** |
| -0.9572 | -0.5399 | *-0.9985* | -0.8319 | 0.2537 | 0.0296 | Q96QB1;A0A0J9 YWS8;R4GMP5 | **DLC1** |
| -0.9826 | -0.6487 | *-0.5585* | -0.7299 | 0.2234 | 0.0298 | Q96FJ2 | **DYNLL2** |
| -0.3910 | -0.7129 | *-0.7120* | -0.6053 | 0.1856 | 0.0299 | Q8N9K5 | **ZNF565** |
| -2.7468 | -1.5495 | *-2.8847* | -2.3936 | 0.7343 | 0.0300 | P51148;F8VVK3; K7ERI8;K7ERQ8; K7ENY4;F8VVZ0 ;K7EIP6 | **RAB5C** |
| 0.2893 | 0.5468 | *0.4212* | 0.4191 | 0.1288 | 0.0301 | A0A0B4J224;E9 PLJ4;E9PQL7;O1 5535;U3KQB2;U 3KQV4 | **ZSCAN9** |
| 2.1378 | 1.2021 | *1.4277* | 1.5892 | 0.4884 | 0.0301 | Q9H992 | **MARCH7** |
| -0.3923 | -0.5985 | *-0.7556* | -0.5821 | 0.1822 | 0.0311 | P30050 | **RPL12** |

| L100/ CTRL1 | L100/ CTRL2 | L100/ CTRL3 | Mean | SD | T-test p-value | Accession | Gene Symbol |
|---|---|---|---|---|---|---|---|
| -0.5190 | -1.0069 | *-0.8726* | -0.7995 | 0.2520 | 0.0316 | A0A1B0GTU4;F5GZ78;P49023;A0A1B0GU60;A0A1B0GWE7;A0A1B0GV30 | **PXN** |
| 0.9957 | 1.3694 | *1.8901* | 1.4184 | 0.4492 | 0.0318 | P35499 | **SCN4A** |
| -4.7155 | -2.4095 | *-3.9620* | -3.6956 | 1.1758 | 0.0321 | A0A2R8Y8E0;A0A2U3TZI2;H7C560;P15056;A0A2R8Y467;A0A2R8YDP5 | **BRAF** |
| -1.6431 | -1.5174 | *-2.6542* | -1.9383 | 0.6232 | 0.0328 | A0A0D9SF60;Q99569;E7EST6;E7EMY7;E7EP40;E9PHJ1 | **PKP4** |
| 1.4192 | 0.8482 | *1.6866* | 1.3180 | 0.4282 | 0.0334 | Q5T282;Q92966 | **SNAPC3** |
| -0.3782 | -0.2251 | *-0.4483* | -0.3505 | 0.1141 | 0.0336 | Q5TZA2;B1AKD8;A0A087WW81;A0A087WU09;Q86T23;B0QYQ9;B0QYR0;B0QYR1;H0YKS0;Q8IVE0;Q9Y2F9 | **CROCC** |
| 0.3099 | 0.2912 | *0.5091* | 0.3700 | 0.1208 | 0.0337 | P30613 | **PKLR** |
| -0.4247 | -0.5606 | *-0.2842* | -0.4232 | 0.1382 | 0.0338 | Q86WZ0 | **HEATR4** |
| -0.3708 | -0.5708 | *-0.7364* | -0.5593 | 0.1831 | 0.0339 | Q13177;H7C1X3 | **PAK2** |
| -0.7254 | -0.3647 | *-0.6650* | -0.5850 | 0.1932 | 0.0345 | Q8TC05 | **MDM1** |
| -0.5995 | -0.6101 | *-0.3142* | -0.5079 | 0.1679 | 0.0345 | Q13510;A0A1B0GTM3;A0A1B0GUA4;A0A1B0GUH5;A0A1B0GW68;E7EMM4;A0A1B0GTP7;A0A1B0GTZ5;A0A1B0GUG1;A0A1B0GUE3;A0A1B0GUW4;A0A1B0GV06;A0A1B0GUB3;A0A1B0GU62;A0A1B0GV88;A0A1B0GVC9;A0A1B0GVG2;A0A1B0GVJ1;A0A1B0GW48;A0A1B0GTD4;A0A1B0GTQ7;A0A1B0GU06;A0A1B0GV95;A0A1B0GVE7;Q86WS4 | **ASAH1** |
| -1.3771 | -0.6831 | *-1.1272* | -1.0625 | 0.3515 | 0.0346 | Q5SZK4;Q5SZK5;Q92558;Q5SZK3;Q9UPY6 | **WASF1** |
| -0.3909 | -0.5832 | *-0.7774* | -0.5838 | 0.1932 | 0.0346 | P24539;Q5QNZ2 | **ATP5PB** |
| 0.2971 | 0.5048 | *0.6015* | 0.4678 | 0.1555 | 0.0349 | Q9Y520;E7EPN9;A0A0A0MS30 | **PRRC2C** |
| 5.0170 | 5.8906 | *2.9086* | 4.6054 | 1.5330 | 0.0350 | A0A087WXK1 | **ABRAXAS1** |

| L100/ CTRL1 | L100/ CTRL2 | L100/ CTRL3 | Mean | SD | T-test p-value | Accession | Gene Symbol |
|---|---|---|---|---|---|---|---|
| -0.7171 | -0.9389 | *-1.3835* | -1.0132 | 0.3393 | 0.0354 | Q9H3K6;A0A087WZT3;A0A0B4J295;H3BTW0;H3BV85 | **BOLA2** |
| -1.4647 | -1.0114 | *-0.7519* | -1.0760 | 0.3607 | 0.0355 | P07910;B2R5W2;B4DY08;G3V4C1;G3V4W0;G3V2Q1;G3V576;G3V555;G3V575;G3V251;B4DSU6;G3V3K6;G3V5X6;G3V2D6;A0A0G2JPF8;B7ZW38;G3V4M8;O60812;P0DMR1;G3V2H6;G3V5V7 | **HNRNPC** |
| -0.3964 | -0.2268 | *-0.4621* | -0.3618 | 0.1214 | 0.0356 | P48449;A0A0G2JQD0;C9J315;A0A0G2JS81 | **LSS** |
| 0.2026 | 0.2607 | *0.1282* | 0.1972 | 0.0664 | 0.0358 | P78527;H0YG84 | **PRKDC** |
| 0.1818 | 0.1047 | *0.2147* | 0.1671 | 0.0564 | 0.0360 | Q8N9W4;F8WBV9 | **GOLGA6L2** |
| -0.2930 | -0.2296 | *-0.1434* | -0.2220 | 0.0751 | 0.0361 | P10909;H0YC35;H0YAS8;E5RJZ5;E7ERK6;E7ETB4;H0YLK8 | **CLU** |
| -0.3300 | -0.6443 | *-0.6508* | -0.5417 | 0.1834 | 0.0361 | Q16875;A0A1W2PR17;Q5VX20;Q5W015;F2Z2I2;A0A1W2PNV9;H0Y483;I1Z9G3;Q16877;Q4VBA9;Q66S35 | **PFKFB3** |
| -0.7783 | -0.9398 | *-1.4674* | -1.0618 | 0.3604 | 0.0363 | K7EP59;Q15831;K7EMR0;K7EQN8 | **STK11** |
| -2.3489 | -1.1444 | *-1.8535* | -1.7823 | 0.6054 | 0.0364 | Q96JF6;I3L508 | **ZNF594** |
| 1.7880 | 1.0329 | *2.1284* | 1.6498 | 0.5606 | 0.0364 | Q5FVE4;K7EKE4;K7EL11;K7ERT0;K7ESC8;K7ESF1 | **ACSBG2** |
| -0.4012 | -0.2150 | *-0.4398* | -0.3520 | 0.1202 | 0.0367 | A0A0A0MR51;O60427 | **FADS1** |
| 1.3703 | 0.6613 | *1.1321* | 1.0545 | 0.3608 | 0.0369 | Q9BQ70;H3BMJ8;H3BSP8;H3BTU3;Q9H7D3 | **TCF25** |
| -0.7256 | -0.3661 | *-0.7269* | -0.6062 | 0.2080 | 0.0371 | B1AJY5;B1AJY7;O75832 | **PSMD10** |
| -0.5952 | -0.5933 | *-0.2981* | -0.4955 | 0.1710 | 0.0375 | Q9BWM7;A0A0A0MS41;A0A1P0AYU5;S4R3N9 | **SFXN3** |
| 0.6697 | 0.7196 | *1.2138* | 0.8677 | 0.3008 | 0.0378 | Q4G0X9;I3L477 | **CCDC40** |
| -0.7871 | -1.2495 | *-0.6559* | -0.8975 | 0.3118 | 0.0380 | Q16658;H7C459;A0A0A0MSB2;C9JFC0;C9JPH9 | **FSCN1** |
| -1.0070 | -1.9664 | *-1.2823* | -1.4186 | 0.4940 | 0.0381 | Q8TAP6 | **CEP76** |

| L100/ CTRL1 | L100/ CTRL2 | L100/ CTRL3 | Mean | SD | T-test p-value | Accession | Gene Symbol |
|---|---|---|---|---|---|---|---|
| 1.0380 | 0.5954 | *1.2531* | 0.9622 | 0.3353 | 0.0382 | Q9UPM6;H0YMY8 | **LHX6** |
| 0.9314 | 0.8115 | *1.5199* | 1.0876 | 0.3791 | 0.0382 | Q92922 | **SMARCC1** |
| -2.1287 | -1.0750 | *-2.1949* | -1.7995 | 0.6283 | 0.0383 | F5H1Y4;Q9HD26;A0A0J9YVX5 | **GOPC** |
| -1.9425 | -1.1303 | *-1.0803* | -1.3844 | 0.4840 | 0.0384 | Q96HA7 | **TONSL** |
| -0.2394 | -0.4435 | *-0.5030* | -0.3953 | 0.1382 | 0.0384 | A0A087WVQ9 | **EEF1A1** |
| 0.2042 | 0.4299 | *0.3421* | 0.3254 | 0.1138 | 0.0384 | Q92930;H0YNE9 | **RAB8B** |
| -1.7320 | -0.8436 | *-1.2388* | -1.2715 | 0.4451 | 0.0385 | P60953;Q5JYX0 | **CDC42** |
| -0.7222 | -0.6164 | *-0.3420* | -0.5602 | 0.1962 | 0.0385 | Q9UP95;I3L1N8 | **SLC12A4** |
| 1.4391 | 0.7945 | *1.6808* | 1.3048 | 0.4582 | 0.0387 | Q16576;E9PC52;Q5JP02;Q5JNZ6;Q5JP01 | **RBBP7** |
| 0.2760 | 0.3654 | *0.5495* | 0.3970 | 0.1395 | 0.0388 | P49840;A8MT37;M0QYV0 | **GSK3A** |
| -0.9143 | -1.0563 | *-1.7257* | -1.2321 | 0.4333 | 0.0388 | Q16445 | **GABRA6** |
| 0.3974 | 0.5505 | *0.8046* | 0.5842 | 0.2057 | 0.0389 | Q9NZL4 | **HSPBP1** |
| 1.7389 | 1.6924 | *0.8464* | 1.4259 | 0.5024 | 0.0390 | P20336;S4R3Q3 | **RAB3A** |
| -0.7540 | -1.2169 | *-1.5885* | -1.1865 | 0.4181 | 0.0390 | Q8TAT6 | **NPLOC4** |
| -0.8591 | -0.4757 | *-0.4971* | -0.6107 | 0.2155 | 0.0391 | Q8TF09;H3BQI1;Q9NP97;B1AKR6;H3BNG9;H3BPA0;Q7Z4M1 | **DYNLRB2** |
| 0.7486 | 1.3648 | *1.5955* | 1.2363 | 0.4378 | 0.0394 | P53675;A0A087WX41;F5H5N6;A0A087WXH4;H0YGJ9 | **CLTCL1** |
| -1.3452 | -0.6448 | *-0.9671* | -0.9857 | 0.3506 | 0.0397 | P50213;H0YL72;H0YKD0;H0YLI6;H0YMU3;H0YM64;H0YNF5;H0YNF8;H0YM46 | **IDH3A** |
| -0.2514 | -0.4515 | *-0.5387* | -0.4138 | 0.1473 | 0.0397 | Q96JM4;H0YCJ9 | **LRRIQ1** |
| 1.7198 | 1.7069 | *3.0553* | 2.1607 | 0.7748 | 0.0403 | G3XAE0;Q5HY98 | **ZNF766** |
| -0.7437 | -0.6060 | *-1.1870* | -0.8456 | 0.3036 | 0.0404 | Q8IVF5;E9PMZ8;F5H6W6;E9PKT1;F5H6R0 | **TIAM2** |
| 0.8567 | 1.5640 | *1.8510* | 1.4239 | 0.5117 | 0.0405 | Q92820 | **GGH** |
| -0.7257 | -0.7894 | *-1.3494* | -0.9549 | 0.3432 | 0.0405 | E9PQY2;Q9NQP4 | **PFDN4** |
| 0.8358 | 1.0282 | *1.6410* | 1.1683 | 0.4205 | 0.0406 | P23786;A0A1B0GTB8;A0A1B0GV75;A0A1B0GVF3;A0A1B0GWC0 | **CPT2** |
| 0.3414 | 0.7231 | *0.6907* | 0.5851 | 0.2116 | 0.0410 | Q6ZNA1;A0A0A0MR57;A0A075B7G2;A0A075B7G3;A0A087WUU8;A0A087WV98;A0A087WWI3;A0A087X254;A0A087X2A5;A0A087X2B0;A0A0C4DGP9;A0 | **ZNF836** |

| L100/ CTRL1 | L100/ CTRL2 | L100/ CTRL3 | Mean | SD | T-test p-value | Accession | Gene Symbol |
|---|---|---|---|---|---|---|---|
| | | | | | | A0U1RQK1;A0A1W2PNY2;A0A1W2PQL4;A0A1W2PRC0;A2RRD8;A6NP11;A8MTY0;A8MUV8;B1APK8;B4DU55;B4DX44;B9EG95;C9K0H3;E7EWC5;F5H032;F5H290;F8W6Y9;F8W889;H0Y892;H3BS42;H9KV89;K7EK80;K7EP55;M0QX96;M0QXU9;M0QY24;M0QYS4;M0R0F3;M0R1M8;O14709;O43309;O43345;O43361;O95600;O95780;P0CB33;P0DPD5;P10073;P17019;P17022;P17035;P17097;P51522;P51815;P52744;Q03924;Q09FC8;Q15929;Q15937;Q2M3W8;Q2M3X9;Q2VY69;Q3ZCX4;Q5JNZ3;Q5SXM1;Q5VIY5;Q6AZW8;Q6P9G9;Q6PDB4;Q6ZMW2;Q6ZN06;Q6ZN19;Q6ZN57;Q76KX8;Q7Z3V5;Q86TJ5;Q86UE3;Q86V71;Q86XN6;Q86Y25;Q8IW36;Q8N8C0;Q8N8J6;Q8N972;Q8N988;Q8N9F8;Q8N9M3;Q8NDQ6;Q8NEM1;Q8NHY6;Q8TB69;Q8TBZ5;Q8TF39;Q8WXB4;Q96CX3;Q96IR2;Q96LX8;Q96MR9;Q96PE6;Q99676;Q9H5H4;Q9H7R5;Q9H963;Q9HBT7;Q9HCL3;Q9NQX6;Q9P0L1;Q9Y473 | |
| -2.7188 | -1.8069 | *-3.8352* | -2.7870 | 1.0158 | 0.0415 | P10412 | **HIST1H1E** |
| -0.3177 | -0.1587 | *-0.2008* | -0.2257 | 0.0824 | 0.0417 | P12270 | **TPR** |

| L100/ CTRL1 | L100/ CTRL2 | L100/ CTRL3 | Mean | SD | T-test p-value | Accession | Gene Symbol |
|---|---|---|---|---|---|---|---|
| -0.5747 | -0.4100 | *-0.8531* | -0.6126 | 0.2239 | 0.0418 | P55010;H0YLZ1;H0YN40;H0YM54;H0YMJ8 | **EIF5** |
| -0.9074 | -1.9659 | *-1.8207* | -1.5647 | 0.5738 | 0.0420 | E9PCE7;E9PMP8;P53804 | **TTC3** |
| 1.9818 | 1.2444 | *0.9872* | 1.4045 | 0.5163 | 0.0422 | O15144;H7C3F9;G5E9J0 | **ARPC2** |
| -0.4772 | -0.3567 | *-0.2190* | -0.3510 | 0.1292 | 0.0423 | Q9BVA1;K7EK43;G3V4U2;M0R2T4;I3L0U9;I3L0V9;I3L4T6;I3L4U4;K7EJ64;K7EJZ4;K7EN98;K7EPE5;K7EQT3;K7ERA8;K7ESQ3;Q8TBP0 | **TUBB2B** |
| -0.6637 | -1.2745 | *-1.4718* | -1.1367 | 0.4213 | 0.0429 | O60664;K7ERZ3;K7EL96;K7ER39 | **PLIN3** |
| -0.2244 | -0.3525 | *-0.4876* | -0.3548 | 0.1316 | 0.0429 | Q99829;B0QZ18;A6PVH9;F2Z2V0;E7ENH5;Q5JX45;Q5JX44;Q5JX56;Q5JX58;Q5JX59;Q5JX60;H0Y524;Q5JX52;Q5JX61;Q5JX55;E7EV27;Q5JX53;Q5JX57;Q5JX54 | **CPNE1** |
| -2.7997 | -2.6731 | *-4.9776* | -3.4835 | 1.2955 | 0.0431 | H0YKN8;H0YKT5;H0YL70;H0YNT2;Q04726;A0A0D9SES8;F5H7D6;H0YNI7;Q04724 | **TLE3** |
| 0.7500 | 1.5641 | *1.6347* | 1.3163 | 0.4917 | 0.0435 | Q9H1A4;H0Y564;A0A2R8YF63 | **ANAPC1** |
| -0.3011 | -0.5869 | *-0.6782* | -0.5220 | 0.1967 | 0.0442 | Q07021;I3L3B0;I3L3Q7 | **C1QBP** |
| -0.6845 | -1.5347 | *-1.3745* | -1.1979 | 0.4517 | 0.0443 | Q6GPH4;C9J7Z8;I3L3D9;I3L509;I3L2B3;I3L3B3;I3L3Q2;I3L534 | **XAF1** |
| 1.8655 | 0.8321 | *1.6831* | 1.4603 | 0.5516 | 0.0444 | Q06124;A0A1W2PPU4;A0A0U1RRI0 | **PTPN11** |
| -0.1444 | -0.2873 | *-0.3258* | -0.2525 | 0.0956 | 0.0446 | Q14240;E7EQG2;J3KSN7;E7EMV8;E9PBH4;F8WE11;J3KS93;J3KT04;I3L3H2;J3QQP0 | **EIF4A2** |
| -0.5328 | -1.1847 | *-1.1118* | -0.9431 | 0.3572 | 0.0446 | Q9BY12;H3BPM0;H3BS25;H3BU24;H3BR40;H3BT27;H3BTL8 | **SCAPER** |
| -0.4250 | -0.5836 | *-0.8985* | -0.6357 | 0.2410 | 0.0447 | P21246;C9JR52 | **PTN** |
| -0.5641 | -0.4082 | *-0.2540* | -0.4088 | 0.1551 | 0.0448 | O75306 | **NDUFS2** |
| -0.6974 | -1.5131 | *-1.5252* | -1.2452 | 0.4745 | 0.0451 | P37059 | **HSD17B2** |

| L100/ CTRL1 | L100/ CTRL2 | L100/ CTRL3 | Mean | SD | T-test p-value | Accession | Gene Symbol |
|---|---|---|---|---|---|---|---|
| 0.4798 | 0.8703 | *1.0929* | 0.8144 | 0.3103 | 0.0452 | Q7Z7A4;W5RWE6 | **PXK** |
| -0.7796 | -1.6963 | *-1.1509* | -1.2089 | 0.4611 | 0.0452 | P63010;A0A087X253;A0A087WU93;A0A087WZQ6;A0A087WYD1;K7ERB2;K7EN71;A0A087WXS3;K7EJX1;K7EKZ5 | **AP2B1** |
| 0.4681 | 0.9159 | *1.0710* | 0.8183 | 0.3131 | 0.0455 | Q8TE73 | **DNAH5** |
| -0.7609 | -0.3401 | *-0.7268* | -0.6093 | 0.2337 | 0.0457 | Q86VS3;H3BU17 | **IQCH** |
| -0.3789 | -0.4636 | *-0.2015* | -0.3480 | 0.1338 | 0.0459 | Q9NXD2 | **MTMR10** |
| 0.1859 | 0.3977 | *0.4212* | 0.3350 | 0.1296 | 0.0465 | Q9UN37;I3L4J1;O75351 | **VPS4A** |
| -0.5233 | -0.3140 | *-0.2546* | -0.3640 | 0.1411 | 0.0466 | K7EJR9;K7EPW9;Q9NPI9;K7EKJ4;K7ELL5 | **KCNJ16** |
| -0.2116 | -0.3166 | *-0.4686* | -0.3323 | 0.1292 | 0.0469 | Q05707;J3QT83;Q4G0W3 | **COL14A1** |
| -0.4647 | -0.4170 | *-0.2009* | -0.3609 | 0.1406 | 0.0471 | P35606;D6R997;D6RBT6;D6RBG7;D6RBZ7;D6RCL6;H0YAC7 | **COPB2** |
| 0.9512 | 2.0834 | *1.3647* | 1.4664 | 0.5729 | 0.0473 | A0A0G2JLQ8;A0A0G2JMG8;A0A0G2JNC8;A0A0G2JPQ2;J3KN39;Q9NX02;A0A0G2JPB6;A0A0G2JLX3;A0A0G2JP37;K7EMK2 | **NLRP2** |
| -0.2871 | -0.1307 | *-0.2966* | -0.2381 | 0.0932 | 0.0474 | P13796;Q5TBN3 | **LCP1** |
| 1.5949 | 1.7449 | *0.7521* | 1.3639 | 0.5352 | 0.0477 | P40818 | **USP8** |
| -0.2407 | -0.5628 | *-0.5056* | -0.4364 | 0.1719 | 0.0480 | O75335;B1APN9;M0QZB5 | **PPFIA4** |
| -0.5840 | -0.4300 | *-0.9290* | -0.6477 | 0.2555 | 0.0482 | Q15075 | **EEA1** |
| 1.6014 | 2.3361 | *3.5751* | 2.5042 | 0.9975 | 0.0490 | O00160 | **MYO1F** |
| -0.5437 | -0.7594 | *-0.3260* | -0.5430 | 0.2167 | 0.0492 | P12814;H9KV75;G3V2W4;G3V2N5;H7C5W8;H0YJW3;G3V2X9;H0YJ11;G3V5M4;G3V2E8 | **ACTN1** |
| -0.4250 | -0.9646 | *-0.6394* | -0.6763 | 0.2716 | 0.0498 | Q86XU0;M0R297 | **ZNF677** |
| -0.2621 | -0.5282 | *-0.6356* | -0.4753 | 0.1923 | 0.0505 | Q8N9H8 | EXD3 |
| -0.2824 | -0.4626 | *-0.6633* | -0.4694 | 0.1906 | 0.0508 | P26378;B1APY8;A0A0R4J2E6;B1APY9;B1AM48 | ELAVL4 |
| 0.4661 | 0.4502 | *0.8806* | 0.5990 | 0.2440 | 0.0511 | O95678 | KRT75 |
| 0.1865 | 0.0936 | *0.2291* | 0.1697 | 0.0693 | 0.0513 | Q96GM5;F8VRQ4;F8VUB0;F8VZ70 | SMARCD1 |
| 0.2983 | 0.5673 | *0.7303* | 0.5320 | 0.2182 | 0.0518 | Q9Y2X3;H7BZ72;F8WED0 | NOP58 |

| L100/ CTRL1 | L100/ CTRL2 | L100/ CTRL3 | Mean | SD | T-test p-value | Accession | Gene Symbol |
|---|---|---|---|---|---|---|---|
| 1.2410 | 0.8204 | *0.5353* | 0.8656 | 0.3550 | 0.0518 | O75534;E9PLT0;E9PKN4;E9PLD4;E9PNG3 | CSDE1 |
| -0.7892 | -1.4312 | *-0.6967* | -0.9724 | 0.4000 | 0.0520 | P08865;C9J9K3;A0A0C4DG17;F8WD59 | RPSA |
| -2.5678 | -3.4538 | *-1.4167* | -2.4794 | 1.0214 | 0.0522 | C9JFW8;Q9UQQ1;E9PII9;E9PKG8;E9PKW7;E9PLR8 | NAALADL1 |
| 0.4307 | 1.0654 | *0.8543* | 0.7834 | 0.3232 | 0.0523 | Q6AW86;M0R1X9;M0R3B5 | ZNF324B |
| -0.5130 | -0.8074 | *-0.3533* | -0.5579 | 0.2303 | 0.0524 | Q8N111 | CEND1 |
| 0.9830 | 1.2832 | *0.5216* | 0.9292 | 0.3836 | 0.0524 | Q9P0V3;C9JED2 | SH3BP4 |
| 0.1622 | 0.1826 | *0.3338* | 0.2262 | 0.0937 | 0.0527 | Q99460;A0A087WW66;H7C378;C9J9M4;H7BZR6;F8WCE3 | PSMD1 |
| 1.2580 | 1.6558 | *2.7812* | 1.8984 | 0.7900 | 0.0532 | P35659;B4DFG0;D6RDA2;H0Y8X0;D6R9L5 | DEK |
| 2.7251 | 3.4624 | *5.9772* | 4.0549 | 1.7051 | 0.0542 | Q5VZM2 | RRAGB |
| -1.3056 | -2.0462 | *-3.1109* | -2.1542 | 0.9075 | 0.0544 | Q9H6S3 | EPS8L2 |
| 0.5200 | 0.9030 | *1.2798* | 0.9010 | 0.3799 | 0.0545 | Q13367;A0A2R8Y2A8 | AP3B2 |
| -0.7032 | -0.7363 | *-1.4143* | -0.9513 | 0.4013 | 0.0545 | Q13123;A0A0C4DGW5;D6REL4;E7EQZ7;Q9H4P6;D6RAY9 | IK |
| -1.0979 | -0.6550 | *-1.6002* | -1.1177 | 0.4729 | 0.0548 | Q96AH8 | RAB7B |
| -0.7595 | -0.4587 | *-1.1167* | -0.7783 | 0.3294 | 0.0549 | E7ESY4;H0Y4T7;Q13330;F8W9Y9 | MTA1 |
| -1.2184 | -1.5020 | *-2.6494* | -1.7899 | 0.7577 | 0.0549 | P22105;A0A140T8Y3;A0A140T902;A0A140T9C0;A0A140TA33;A0A140TA41;A0A140TA52;A0A140T8Z8;A0A087X0I0;A0A140T9L7;Q16473 | TNXB |
| -0.4275 | -0.2221 | *-0.2116* | -0.2871 | 0.1218 | 0.0551 | O43143 | DHX15 |
| -3.4722 | -1.8129 | *-1.7061* | -2.3304 | 0.9903 | 0.0553 | F8VWP7;F8W927 | CAPS2 |
| -0.8735 | -0.3472 | *-0.8155* | -0.6787 | 0.2886 | 0.0553 | O76094;D6RDY6;R4GNC1 | SRP72 |
| -0.5297 | -0.2581 | *-0.6610* | -0.4829 | 0.2055 | 0.0554 | M0R344 | SPHK2 |
| -1.0159 | -1.0101 | *-0.4131* | -0.8130 | 0.3464 | 0.0555 | P60660;F8W1R7;G3V1V0;J3KND3;B7Z6Z4;G8JLA2;G3V1Y7;F8VPF3;F8VZU9;F8W180;H0YI43;F8VXL3 | MYL6 |

| L100/ CTRL1 | L100/ CTRL2 | L100/ CTRL3 | Mean | SD | T-test p-value | Accession | Gene Symbol |
|---|---|---|---|---|---|---|---|
| 2.1969 | 1.0438 | *1.1813* | 1.4740 | 0.6299 | 0.0558 | F8VU39;J3KPG5; Q9UIF9;A0A0C4 DGI9 | BAZ2A |
| -1.2052 | -0.4665 | *-0.9810* | -0.8842 | 0.3788 | 0.0561 | A0A0A0MTI5;B8 ZWD1;P07108 | DBI |
| 2.3732 | 0.9960 | *2.5272* | 1.9655 | 0.8431 | 0.0562 | Q14161;F8VXI9; F8WAK2;R4GNG 3;B5BU58;F8W8 22;Q6FI58 | GIT2 |
| 0.3734 | 0.9594 | *0.8662* | 0.7330 | 0.3149 | 0.0564 | Q9Y2F5 | ICE1 |
| -1.2888 | -1.5814 | *-2.8251* | -1.8984 | 0.8158 | 0.0564 | O00408;F5H130 | PDE2A |
| -0.6896 | -1.0068 | *-1.6261* | -1.1075 | 0.4763 | 0.0565 | P49006 | MARCKSL1 |
| 0.5591 | 1.3580 | *1.4128* | 1.1100 | 0.4778 | 0.0566 | Q6ZSZ6;H0YN23 ;H0YKA1 | TSHZ1 |
| -0.5601 | -0.7117 | *-1.2506* | -0.8408 | 0.3629 | 0.0569 | Q8NFW8;F5GYM 0 | CMAS |
| -0.6761 | -0.9367 | *-1.5692* | -1.0607 | 0.4593 | 0.0572 | Q9BUT1;D6R9P2 | BDH2 |
| 2.4810 | 1.1802 | *1.3044* | 1.6552 | 0.7178 | 0.0574 | Q8N4Y2;E9PPF3 ;E9PHZ8;E9PK0 4;E9PRE5 | CRACR2B |
| 0.4661 | 0.1875 | *0.3116* | 0.3217 | 0.1396 | 0.0574 | O15195;E9PFV5; H7BZ43 | VILL |
| 0.2385 | 0.6208 | *0.5687* | 0.4760 | 0.2073 | 0.0578 | A0A1B0GV05;Q6 84P5 | RAP1GAP2 |
| -2.3866 | -0.9616 | *-2.4517* | -1.9333 | 0.8421 | 0.0578 | P42785;E9PIG4; E9PKN6;E9PL85; E9PQB5;E9PQN 3;E9PLY4;E9PNJ 1;E9PNF7;E9PL4 9 | PRCP |
| -1.5793 | -1.5308 | *-0.6176* | -1.2426 | 0.5418 | 0.0579 | B5MD58;P36956; S4R3B4;X6RBE4 | SREBF1 |
| 0.9858 | 1.0448 | *2.0395* | 1.3567 | 0.5921 | 0.0580 | P37235 | HPCAL1 |
| -1.2285 | -3.2132 | *-2.9742* | -2.4720 | 1.0835 | 0.0585 | Q9UHC1 | MLH3 |
| -0.3618 | -0.7410 | *-0.9560* | -0.6863 | 0.3008 | 0.0585 | O94822;H7BYG8 | LTN1 |
| -0.8656 | -1.1570 | *-1.9974* | -1.3400 | 0.5877 | 0.0585 | Q9H2G2 | SLK |
| -0.9535 | -2.5131 | *-2.2760* | -1.9142 | 0.8404 | 0.0587 | P51648;J3QRD1; J3KTD9 | ALDH3A2 |
| -0.4543 | -1.1584 | *-1.1612* | -0.9246 | 0.4073 | 0.0590 | Q9UM47 | NOTCH3 |
| -2.3241 | -0.8705 | *-2.0331* | -1.7426 | 0.7691 | 0.0592 | Q9UKN8 | GTF3C4 |
| -0.1103 | -0.2949 | *-0.2606* | -0.2220 | 0.0982 | 0.0595 | P07205 | PGK2 |
| -0.8062 | -1.1724 | *-0.4526* | -0.8104 | 0.3599 | 0.0599 | P53999;D6R970; D6RC37 | SUB1 |
| -0.4627 | -1.0399 | *-0.5702* | -0.6909 | 0.3069 | 0.0599 | P07951;A7XZE4; Q5TCU3;Q5TCU 8;U3KQK2 | TPM2 |
| -1.4465 | -1.0601 | *-0.5447* | -1.0171 | 0.4524 | 0.0601 | P63104;E7ESK7; B0AZS6;B7Z2E6; H0YB80;E7EVZ2 ;E9PD24;E5RIR4 ;E5RGE1 | YWHAZ |
| -0.8729 | -0.3399 | *-0.5864* | -0.5997 | 0.2668 | 0.0601 | Q5JTJ3 | COA6 |
| -2.0783 | -0.7683 | *-1.6887* | -1.5118 | 0.6727 | 0.0601 | P61020;F8VUA5 | RAB5B |

| L100/ CTRL1 | L100/ CTRL2 | L100/ CTRL3 | Mean | SD | T-test p-value | Accession | Gene Symbol |
|---|---|---|---|---|---|---|---|
| 0.3930 | 0.3739 | *0.7810* | 0.5160 | 0.2297 | 0.0602 | P50416 | CPT1A |
| 0.9174 | 1.5215 | *0.6292* | 1.0227 | 0.4554 | 0.0602 | Q9UJ14 | GGT7 |
| -0.9132 | -1.8328 | *-2.4474* | -1.7311 | 0.7721 | 0.0604 | F8VNV8;P54284;F8VUW8;F8VV14 | CACNB3 |
| 0.9581 | 0.6843 | *1.6357* | 1.0927 | 0.4897 | 0.0609 | A0A087WUD3;Q9NRP0 | OSTC |
| -0.9479 | -0.9892 | *-0.3717* | -0.7696 | 0.3452 | 0.0610 | Q15907;H3BMH2;H3BSC1;P62491;B4DQU5 | RAB11B |
| 0.5176 | 0.8511 | *1.3193* | 0.8960 | 0.4028 | 0.0612 | A0A0A0MS50;Q96RE9 | ZNF300 |
| -0.3485 | -0.6394 | *-0.2814* | -0.4231 | 0.1903 | 0.0613 | O95747;C9JIG9 | OXSR1 |
| 0.7825 | 0.8678 | *1.6924* | 1.1142 | 0.5025 | 0.0616 | F5H0R1;Q9H6E5;H3BRB1;F8WA97;J3KN81 | TUT1 |
| -5.0216 | -1.8967 | *-3.4659* | -3.4614 | 1.5625 | 0.0617 | B7ZM68;Q6ZNL6 | FGD5 |
| -0.2774 | -0.1706 | *-0.4361* | -0.2947 | 0.1336 | 0.0622 | Q9BXS5;K7EJL1;E7ENJ6;K7EQX3 | AP1M1 |
| -0.2863 | -0.6469 | *-0.7969* | -0.5767 | 0.2625 | 0.0626 | P36957;Q86SW4;G3V3F0;G3V5M3;H0YJF9;B7ZAZ8 | DLST |
| -0.5859 | -0.7428 | *-0.2669* | -0.5319 | 0.2425 | 0.0628 | O43765;K7EMD6 | SGTA |
| -0.2640 | -0.4652 | *-0.6976* | -0.4756 | 0.2170 | 0.0629 | P16144;J3QQL2;J3QRK0 | ITGB4 |
| 0.7793 | 1.9729 | *1.2135* | 1.3219 | 0.6041 | 0.0631 | O43854 | EDIL3 |
| -1.0059 | -0.3711 | *-0.9803* | -0.7857 | 0.3593 | 0.0632 | P23258;Q9NRH3;K7EIS0;K7EKE5 | TUBG1 |
| -0.3721 | -0.4913 | *-0.1771* | -0.3468 | 0.1586 | 0.0632 | P37108;H0YLA2;H0YLW0 | SRP14 |
| -0.8787 | -2.1787 | *-2.4587* | -1.8387 | 0.8431 | 0.0635 | E5RFH5;F5H8F7;Q9UBL3 | ASH2L |
| -1.0722 | -0.3979 | *-0.7271* | -0.7324 | 0.3372 | 0.0639 | P10599 | TXN |
| -0.4409 | -0.5506 | *-0.1950* | -0.3955 | 0.1821 | 0.0639 | Q5JZY3;J3KQG3 | EPHA10 |
| -1.1638 | -0.7074 | *-0.4588* | -0.7767 | 0.3576 | 0.0640 | B7Z645;H0YAA0;O60506;F6UXX1 | SYNCRIP |
| -0.1917 | -0.3890 | *-0.5319* | -0.3709 | 0.1708 | 0.0640 | P14618;B4DNK4;H3BQ34;H3BTJ2;H3BUW1;H3BT25;H3BU13;H3BN34;H3BQZ3;H3BTN5 | PKM |
| 1.8305 | 2.4336 | *0.8689* | 1.7110 | 0.7892 | 0.0642 | R4GNB9 | TRIM11 |
| 0.7241 | 0.3646 | *1.0095* | 0.6994 | 0.3232 | 0.0644 | O15085 | ARHGEF11 |
| 0.9630 | 2.1132 | *2.7224* | 1.9329 | 0.8934 | 0.0644 | Q9Y2G4;E5RJ05 | ANKRD6 |
| 0.6229 | 1.4566 | *0.7854* | 0.9550 | 0.4419 | 0.0646 | A8MUS3;H7BY10;K7EJV9;K7ERT8;P62750;K7EMA7 | RPL23A |
| 2.1617 | 2.1265 | *0.7873* | 1.6919 | 0.7836 | 0.0646 | O60296;Q53RS6 | TRAK2 |
| 0.8404 | 2.0195 | *1.1189* | 1.3263 | 0.6163 | 0.0650 | E9PKC0;Q6IQ23;A0A1B0GTN9;A0A1B0GUN0;H0YDE2 | PLEKHA7 |

| L100/ CTRL1 | L100/ CTRL2 | L100/ CTRL3 | Mean | SD | T-test p-value | Accession | Gene Symbol |
|---|---|---|---|---|---|---|---|
| -2.4738 | -1.4641 | *-0.9806* | -1.6395 | 0.7619 | 0.0650 | Q9H0U4;E9PLD0 | RAB1B |
| -0.4025 | -0.3790 | *-0.8251* | -0.5356 | 0.2511 | 0.0661 | O43684;J3QT28; J3QSX4 | BUB3 |
| -0.2861 | -0.5413 | *-0.2309* | -0.3528 | 0.1656 | 0.0662 | Q8WXX0 | DNAH7 |
| -0.3899 | -0.1393 | *-0.2704* | -0.2665 | 0.1253 | 0.0665 | Q12906;K7EKJ9; K7EQR9;K7EKY 0;K7ER69;K7EJ0 9;K7EM82;K7EN K6;K7EQ75;K7E RM6 | ILF3 |
| -1.1221 | -0.9248 | *-0.3855* | -0.8108 | 0.3813 | 0.0665 | A0A0A0MR39;Q9 P2K5;A0A087WU T0;A0A0A0MQW 0;A0A087WWC8; A0A0C4DGV1;H 0YKS1;H0YN19 | MYEF2 |
| -1.0465 | -0.4199 | *-1.2303* | -0.8989 | 0.4249 | 0.0671 | E9PR30;P62861 | FAU |
| -2.3396 | -5.1125 | *-2.4701* | -3.3074 | 1.5646 | 0.0672 | Q3BBV0;S4R2X0 ;S4R2Z6 | NBPF1 |
| -0.7543 | -0.2578 | *-0.5881* | -0.5334 | 0.2527 | 0.0674 | P62280;M0QZC5 ;M0R1H6;M0R1H 5 | RPS11 |
| -0.2031 | -0.4941 | *-0.5980* | -0.4317 | 0.2047 | 0.0675 | E9PM92;O00193; E9PQA1;E9PRZ9 | C11orf58 |
| -0.2113 | -0.1097 | *-0.3090* | -0.2100 | 0.0996 | 0.0675 | Q8IZS8;B4DSF6; C9JAV5 | CACNA2D3 |
| -0.4082 | -0.4457 | *-0.9096* | -0.5878 | 0.2793 | 0.0677 | P35232;C9JW96; E7ESE2;E9PCW 0;C9JZ20;D6RBK 0 | PHB |
| -0.4966 | -0.4573 | *-0.1703* | -0.3747 | 0.1781 | 0.0678 | P80723 | BASP1 |
| -0.5383 | -0.6227 | *-0.2108* | -0.4572 | 0.2176 | 0.0679 | P49189 | ALDH9A1 |
| 1.2271 | 1.1360 | *0.4186* | 0.9272 | 0.4428 | 0.0683 | P38117;M0QY67 | ETFB |
| -0.7292 | -1.4784 | *-0.6531* | -0.9536 | 0.4561 | 0.0685 | P51587;H0YE37; H0YD86 | BRCA2 |
| -0.4849 | -0.1668 | *-0.4650* | -0.3722 | 0.1782 | 0.0686 | P60174;U3KPZ0; U3KQF3;U3KPS 5 | TPI1 |
| 0.6694 | 0.9662 | *1.7061* | 1.1139 | 0.5339 | 0.0688 | A6NNF4;A0A0A0 MR41;P0DKX0 | ZNF726 |
| -0.1730 | -0.4832 | *-0.3124* | -0.3229 | 0.1553 | 0.0692 | P28482;K7EK24; Q9NS82 | MAPK1 |
| 1.1832 | 3.2817 | *2.0912* | 2.1853 | 1.0524 | 0.0694 | A0A0A0MRN9;Q 9GZV1;Q5T457 | ANKRD2 |
| 0.5730 | 0.8128 | *1.4553* | 0.9471 | 0.4562 | 0.0694 | Q5VYS8;Q5VYS 9 | TUT7 |
| -0.4526 | -0.7110 | *-0.2564* | -0.4733 | 0.2280 | 0.0694 | P27824;D6RGY2 ;D6RAU8;D6RB8 5;D6RDP7;H0Y9 Q7;D6RFL1;H0Y 9H1;D6RAQ8;D6 RD16;D6RHJ3 | CANX |
| -1.0724 | -0.9964 | *-2.2391* | -1.4360 | 0.6965 | 0.0703 | V9GYU8 | EBF4 |
| 1.2041 | 0.4232 | *0.7837* | 0.8036 | 0.3908 | 0.0706 | Q86VS8 | HOOK3 |
| -4.2274 | -2.0089 | *-5.9820* | -4.0728 | 1.9911 | 0.0713 | E7EQA9;Q92844 | TANK |

| L100/ CTRL1 | L100/ CTRL2 | L100/ CTRL3 | Mean | SD | T-test p-value | Accession | Gene Symbol |
|---|---|---|---|---|---|---|---|
| -0.3107 | -0.1143 | *-0.3512* | -0.2587 | 0.1267 | 0.0715 | Q8NCB2;E7ETR1;C9J9E2;B4DM24;B4DSW8;F8WDJ4 | CAMKV |
| -0.7698 | -0.2615 | *-0.5222* | -0.5178 | 0.2542 | 0.0718 | Q8NEV1 | CSNK2A3 |
| -0.5809 | -1.1690 | *-1.7195* | -1.1565 | 0.5694 | 0.0722 | Q9H5L6;D6RCT5;D6REM3;F2Z371 | THAP9 |
| -0.5755 | -1.1157 | *-0.4538* | -0.7150 | 0.3523 | 0.0723 | Q9P2J8;C9J5H1;J3QKY7 | ZNF624 |
| 0.6065 | 0.4308 | *1.1113* | 0.7162 | 0.3533 | 0.0724 | P0DPH8;P0DPH7;F8W0F6;C9K0S6;A0A087WVM1;A6XGL0;F8VXZ7;H0YFQ2;H0YMX6;P78382;Q5W1L7 | TUBA3D |
| -0.8112 | -0.2762 | *-0.5360* | -0.5411 | 0.2676 | 0.0727 | Q9HC77;F6VUX8 | CENPJ |
| -0.5832 | -0.1982 | *-0.6054* | -0.4623 | 0.2290 | 0.0729 | Q9NSD9 | FARSB |
| 0.4612 | 0.7756 | *1.2917* | 0.8428 | 0.4193 | 0.0735 | Q8IZH2;H7C5E4;C9JCZ8 | XRN1 |
| -0.7687 | -0.4188 | *-0.2941* | -0.4939 | 0.2460 | 0.0737 | P30086;O15522;O95096 | PEBP1 |
| -0.2587 | -0.6279 | *-0.3127* | -0.3998 | 0.1994 | 0.0738 | O95394;A0A087WT27;J3KN95;D6RCQ8;D6RIS6;D6RCD1;H0Y8I3;H0Y987 | PGM3 |
| -1.6481 | -0.9375 | *-2.6807* | -1.7554 | 0.8765 | 0.0740 | Q08379;A0A2Q2TH77;B7ZC06;A0A087WYC0;R4GND7 | GOLGA2 |
| 0.9588 | 0.9234 | *2.0874* | 1.3232 | 0.6620 | 0.0743 | Q969N2;A0A1W2PP57;A0A1W2PPC3;A0A1W2PPS0;A0A1W2PP53;A0A1W2PPQ7;A0A1W2PRZ8;A0A1W2PP13;A0A1W2PNP0;A0A1W2PQ52;F6W983;A0A1W2PPR6 | PIGT |
| -0.5481 | -0.8353 | *-1.4810* | -0.9548 | 0.4778 | 0.0743 | Q9UPN4;I3L2J8;I3L2X7;I3L4M5;I3L316 | CEP131 |
| -0.5445 | -0.1707 | *-0.4451* | -0.3868 | 0.1936 | 0.0743 | J3KT51;Q9UK76;J3KSH8 | JPT1 |
| -0.9000 | -0.3014 | *-0.9432* | -0.7149 | 0.3588 | 0.0747 | E7EM64;Q7L5N1;H7C3T0 | COPS6 |
| -1.5008 | -0.4740 | *-1.1347* | -1.0365 | 0.5204 | 0.0747 | P78352;C9JWP9;C9JYG3;K7EKP9;K7EKU8;O14909 | DLG4 |
| -0.1208 | -0.1822 | *-0.0605* | -0.1212 | 0.0609 | 0.0747 | O94759;E9PGK7;C9JZQ8 | TRPM2 |
| -0.5059 | -0.1707 | *-0.5422* | -0.4063 | 0.2048 | 0.0753 | P63000;A4D2P1 | RAC1 |
| 0.3952 | 0.9660 | *0.4758* | 0.6123 | 0.3089 | 0.0754 | F8W6K2 | CHN1 |

| L100/ CTRL1 | L100/ CTRL2 | L100/ CTRL3 | Mean | SD | T-test p-value | Accession | Gene Symbol |
|---|---|---|---|---|---|---|---|
| -0.2932 | -0.7485 | *-0.9484* | -0.6634 | 0.3358 | 0.0758 | P25788;G3V4X5;G3V3W4;G3V5N4 | PSMA3 |
| -0.5854 | -1.4259 | *-0.6938* | -0.9017 | 0.4572 | 0.0761 | Q969E4;Q6IPX3 | TCEAL3 |
| 0.4696 | 1.3691 | *1.5220* | 1.1202 | 0.5686 | 0.0762 | A8K0Z3 | WASHC1 |
| 3.9267 | 2.1899 | *1.4331* | 2.5166 | 1.2785 | 0.0763 | Q13111;K7EJF1 | CHAF1A |
| 0.7817 | 2.4847 | *2.4030* | 1.8898 | 0.9605 | 0.0764 | O00410;E7EQT5;E7ETV3;E7EV12;C9JMV5;C9JZ53;C9J875;C9JQT6;C9JXE0;C9JZD8;E7ESA1;E7ESZ1;E7ETV8;E7EWK4;E7EX05 | IPO5 |
| -0.4646 | -0.6174 | *-0.1911* | -0.4244 | 0.2160 | 0.0766 | Q07065 | CKAP4 |
| -1.1316 | -0.4871 | *-1.5586* | -1.0591 | 0.5394 | 0.0767 | F8W8I6;P31483;H7BY49 | TIA1 |
| -0.8457 | -1.0586 | *-0.3235* | -0.7426 | 0.3782 | 0.0767 | Q14203;E7EX90;Q6AWB1;E7EWF7;E9PCY0;C9JJD0;C9JKG6;C9J1B7;C9JJN7;C9JTE5;C9JZA4 | DCTN1 |
| -0.1369 | -0.0830 | *-0.2356* | -0.1518 | 0.0774 | 0.0767 | Q99426;K7EK42;K7EP07;K7EL99;K7EQH0;K7ER04 | TBCB |
| 0.5710 | 1.4027 | *1.8525* | 1.2754 | 0.6502 | 0.0768 | Q8N806;G3V253;G3V2G3;G3V336;H0YJA0 | UBR7 |
| 1.2673 | 2.2426 | *0.8053* | 1.4384 | 0.7338 | 0.0769 | A2RUR9;C9JT67;A6NG92;A0A087WSY3;Q3MJ40;A6NJB5;C9JAY6 | CCDC144A |
| -0.4273 | -0.2961 | *-0.7940* | -0.5058 | 0.2580 | 0.0769 | A0A0A0MR49;Q9HCS2 | CYP4F12 |
| -0.7010 | -0.5857 | *-0.2118* | -0.4995 | 0.2557 | 0.0774 | Q6IAA8;F5GX19;F5H3Y3;F5H479;H0YFI1 | LAMTOR1 |
| -0.5078 | -0.6393 | *-1.2857* | -0.8109 | 0.4164 | 0.0778 | Q9NX78;G3V320 | TMEM260 |
| 0.1589 | 0.2800 | *0.4684* | 0.3024 | 0.1559 | 0.0783 | P11532;A0A075B6G3;E9PDN5;H0Y304;H0Y864;Q4G0X0;A0A0B4J1W6;Q14172 | DMD |
| 2.4187 | 2.4680 | *0.7613* | 1.8827 | 0.9714 | 0.0784 | F8VXU5;Q9UBQ0 | VPS29 |
| -0.6998 | -2.3025 | *-2.1908* | -1.7310 | 0.8948 | 0.0787 | A0A024RCR6;A0A0G2JK23;P46379;A0A0G2JL47;H0Y710;A0A0G2JJM1;A0A0G2JJR8;A0A1B0GX79;F6S6P2;F6TC96;F6U1F2;F6U341;F6UR09;F6VEM6;F6WML8;F6X9 | BAG6 |

| L100/ CTRL1 | L100/ CTRL2 | L100/ CTRL3 | Mean | SD | T-test p-value | Accession | Gene Symbol |
|---|---|---|---|---|---|---|---|
| | | | | | | W3;F6XTU0;X6REW1 | |
| 0.4295 | 1.4441 | *1.2454* | 1.0397 | 0.5377 | 0.0788 | Q9GZV7;Q5T3J1 | HAPLN2 |
| 1.0027 | 3.3602 | *2.6026* | 2.3218 | 1.2036 | 0.0791 | P11274;A9UF01 | BCR |
| -0.4104 | -0.6844 | *-0.2294* | -0.4414 | 0.2290 | 0.0792 | Q15424;K7ES42;H7C3F4 | SAFB |
| -0.4766 | -0.1865 | *-0.2351* | -0.2994 | 0.1554 | 0.0792 | P27448;A0A0A0MSZ1;J3KNR0;H0YIY6 | MARK3 |
| -0.3780 | -0.1791 | *-0.5705* | -0.3759 | 0.1957 | 0.0797 | Q02386;K7EPV5 | ZNF45 |
| 0.9271 | 2.3794 | *1.1689* | 1.4918 | 0.7781 | 0.0800 | Q8IY21;D6R944;H0Y9B2 | DDX60 |
| -0.3633 | -0.7146 | *-0.2695* | -0.4491 | 0.2346 | 0.0802 | O60292 | SIPA1L3 |
| -0.5777 | -0.3406 | *-1.0001* | -0.6395 | 0.3341 | 0.0802 | P53007 | SLC25A1 |
| -1.2522 | -2.2008 | *-0.7571* | -1.4034 | 0.7336 | 0.0803 | Q8IWV7;A0A087WTJ9;H3BUC4;H3BUZ4 | UBR1 |
| 2.2333 | 1.1003 | *0.8646* | 1.3994 | 0.7317 | 0.0803 | Q9BU68 | PRR15L |
| 0.1983 | 0.0806 | *0.2706* | 0.1832 | 0.0959 | 0.0805 | Q96DG6 | CMBL |
| -0.7771 | -0.6203 | *-1.5966* | -0.9980 | 0.5243 | 0.0810 | Q9P258 | RCC2 |
| -0.4356 | -0.9699 | *-1.4296* | -0.9450 | 0.4974 | 0.0813 | Q8TF46 | DIS3L |
| -0.8530 | -0.2644 | *-0.5545* | -0.5573 | 0.2943 | 0.0817 | O94762;J3KTQ2;J3QLU0 | RECQL5 |
| 0.7227 | 1.9977 | *2.5266* | 1.7490 | 0.9273 | 0.0823 | Q8NDV7 | TNRC6A |
| -0.6130 | -1.7898 | *-0.9929* | -1.1319 | 0.6006 | 0.0824 | L8E9D3 | FANCA |
| -0.5739 | -0.8425 | *-0.2513* | -0.5559 | 0.2960 | 0.0829 | A5YKK6;H3BMH0;H3BT18;H3BU44 | CNOT1 |
| -1.2425 | -2.1804 | *-0.7225* | -1.3818 | 0.7389 | 0.0835 | Q7Z3U7;A0A286YFF8;F8VZV1 | MON2 |
| -0.2256 | -0.1449 | *-0.0688* | -0.1464 | 0.0784 | 0.0837 | Q8NBS9 | TXNDC5 |
| -0.7768 | -0.2289 | *-0.8007* | -0.6021 | 0.3234 | 0.0842 | M0R117;M0R1A7;Q02543;M0R0P7;M0R3D6 | RPL18A |
| -0.4784 | -1.3844 | *-0.7393* | -0.8674 | 0.4664 | 0.0844 | P04844;Q5JYR7;Q5JYR4;F2Z3K5;Q5JYR3 | RPN2 |
| 1.5150 | 2.1855 | *4.2514* | 2.6506 | 1.4263 | 0.0845 | Q9NQA5;A0A0A6YY98;H7C2J6 | TRPV5 |
| -0.1283 | -0.4607 | *-0.4166* | -0.3352 | 0.1805 | 0.0846 | E7EV01;O15484;E9PS73;K7EP62 | CAPN5 |
| -0.7323 | -0.6431 | *-1.6146* | -0.9966 | 0.5370 | 0.0847 | Q9Y2L6;E9PGA7 | FRMD4B |
| -0.6867 | -0.2581 | *-0.3294* | -0.4247 | 0.2297 | 0.0852 | Q9BPW8;H7C2U6;C9JDV8;F8WCR5 | NIPSNAP1 |
| -0.5870 | -0.4795 | *-0.1614* | -0.4093 | 0.2213 | 0.0852 | P48681 | NES |
| -0.6720 | -0.5128 | *-1.3826* | -0.8558 | 0.4631 | 0.0853 | Q92736;H7BY35 | RYR2 |
| 1.2968 | 0.4680 | *1.6885* | 1.1511 | 0.6232 | 0.0854 | Q96H55;A0A087WU55;A0A087WW10;A0A087WY49;A0A087WYT7;A0A087WZQ9 | MYO19 |

| L100/ CTRL1 | L100/ CTRL2 | L100/ CTRL3 | Mean | SD | T-test p-value | Accession | Gene Symbol |
|---|---|---|---|---|---|---|---|
| 0.4240 | 0.1167 | *0.3358* | 0.2922 | 0.1582 | 0.0854 | P62753;A2A3R5; A2A3R7 | RPS6 |
| -0.1955 | -0.1318 | *-0.3774* | -0.2349 | 0.1274 | 0.0857 | Q96Q15;J3KRA9;I3L0W2;I3L400; E9PNP6;A0A087WXM7;A0A087WW29;C9JIV0;C9JLP9;C9JVA0;H3BPS6;H3BR09;Q6P435;Q6ZU64;Q96GD3 | SMG1 |
| -0.9005 | -2.1970 | *-3.1554* | -2.0843 | 1.1317 | 0.0858 | Q8NDA2 | HMCN2 |
| -0.4944 | -0.2814 | *-0.8714* | -0.5490 | 0.2988 | 0.0861 | Q15349;F2Z2J1; B7Z3B5 | RPS6KA2 |
| -0.1767 | -0.3253 | *-0.5593* | -0.3538 | 0.1929 | 0.0864 | Q9P2D0;E7EPI0; E9PDR5 | IBTK |
| -0.2626 | -0.7635 | *-0.3984* | -0.4748 | 0.2591 | 0.0865 | O14618;J3KNF4; E9PP76 | CCS |
| 0.4679 | 1.0889 | *0.4475* | 0.6681 | 0.3646 | 0.0866 | Q7Z6Z7;A0A1B0GXC7;Q5H963;H0Y659;A0A087X146;A0A087X1S3; B1AJU2;H0YF09 | HUWE1 |
| -2.7996 | -6.0339 | *-2.2942* | -3.7093 | 2.0290 | 0.0869 | Q12965;H0YNQ8;H0YLE5;H0YLJ4 | MYO1E |
| -0.2209 | -0.3692 | *-0.6732* | -0.4211 | 0.2306 | 0.0871 | P11586;V9GYY3; F5H2F4 | MTHFD1 |
| -0.5323 | -0.1427 | *-0.4432* | -0.3727 | 0.2041 | 0.0871 | Q9P0L0;J3QKM9 | VAPA |
| -0.5127 | -1.8959 | *-1.7466* | -1.3851 | 0.7592 | 0.0872 | O75122;J3KR49; E3W994;A0A0U1RQI6;D6RBU8;H7C4M5;C9J668;H7C4I5;E7ERI8;E7EW49 | CLASP2 |
| -0.4210 | -0.7454 | *-0.2385* | -0.4683 | 0.2567 | 0.0873 | Q16186;A0A087WX59 | ADRM1 |
| 1.6587 | 0.4406 | *1.4210* | 1.1734 | 0.6457 | 0.0878 | Q9Y250 | LZTS1 |
| -0.7262 | -0.8305 | *-0.2208* | -0.5925 | 0.3261 | 0.0879 | Q8NI27;A0A0C4DG98;H0Y7U4;H7C477 | THOC2 |
| -0.2607 | -0.9150 | *-0.9711* | -0.7156 | 0.3949 | 0.0883 | Q9Y228;E2QRE5 | TRAF3IP3 |
| -1.0336 | -0.6405 | *-1.9366* | -1.2036 | 0.6645 | 0.0884 | Q96A35 | MRPL24 |
| -0.5959 | -0.2926 | *-0.9811* | -0.6232 | 0.3451 | 0.0888 | Q9NX14 | NDUFB11 |
| 0.9004 | 3.0904 | *1.9298* | 1.9735 | 1.0957 | 0.0892 | O75164 | KDM4A |
| -0.7511 | -0.7789 | *-0.2080* | -0.5793 | 0.3219 | 0.0893 | P08559;Q5JPT9; Q5JPU0;Q5JPU1;Q5JPU2;Q5JPU3 | PDHA1 |
| -0.1525 | -0.4541 | *-0.2350* | -0.2805 | 0.1559 | 0.0893 | P12081;B3KWE1;B4DDD8;B4E1C5;E7ETE2;A0A2R8YCI2;A0A2R8YFR1;D6RF05;A0A2R8Y4X6;B4DQ67;D6RB22 | HARS |

| L100/ CTRL1 | L100/ CTRL2 | L100/ CTRL3 | Mean | SD | T-test p-value | Accession | Gene Symbol |
|---|---|---|---|---|---|---|---|
| -0.3295 | -0.9263 | *-1.2400* | -0.8320 | 0.4625 | 0.0894 | P10606 | COX5B |
| 2.1736 | 0.7309 | *2.7991* | 1.9012 | 1.0607 | 0.0900 | Q9Y230;M0R0Y3;X6R2L4;M0QXI6;M0R0Z0;A0A1W2PS48 | RUVBL2 |
| -0.4895 | -0.7629 | *-0.2160* | -0.4895 | 0.2734 | 0.0902 | A0A0A0MS54;B2RB89;P22694;A0A087WVC4;B1APF9;B1APG3;B1APF8;B1APG2;B1APG0;B1APG1;B1APF7 | PRKACB |
| -1.4175 | -1.4572 | *-0.3859* | -1.0869 | 0.6074 | 0.0902 | Q01546 | KRT76 |
| -1.3352 | -0.9959 | *-2.7939* | -1.7083 | 0.9553 | 0.0903 | O60925;E5RGS4 | PFDN1 |
| -0.7760 | -0.7684 | *-1.8776* | -1.1407 | 0.6383 | 0.0904 | Q12955;A0A087WTF3;A0A087WZ26;B1AQT1;H0Y3A4;A0A087WTE8;A0A087WVC2;A0A087X0B4;D6RFK6;A0A087WV39;D6RBY7;D6RHY3;H0YA66;K7EQY5;P42679 | ANK3 |
| -3.1088 | -1.3276 | *-1.2289* | -1.8884 | 1.0580 | 0.0906 | Q96GE4 | CEP95 |
| -1.0964 | -0.7204 | *-2.1504* | -1.3224 | 0.7413 | 0.0907 | Q8TBY8;F5H5J2 | PMFBP1 |
| -1.0736 | -0.2693 | *-0.9473* | -0.7634 | 0.4326 | 0.0924 | Q96CS3 | FAF2 |
| -0.3786 | -0.2673 | *-0.7805* | -0.4755 | 0.2700 | 0.0927 | P42166;P42167;G5E972;H0YJH7 | TMPO |
| -0.8250 | -0.3792 | *-1.3424* | -0.8488 | 0.4820 | 0.0928 | Q96AC1;H0YJ34;A0A0U1RRM8;G3V281;G3V5R2 | FERMT2 |
| -0.3220 | -1.2503 | *-0.9011* | -0.8245 | 0.4689 | 0.0930 | A1L0T0;E9PJS0;M0R026;E9PL44 | ILVBL |
| -0.6157 | -1.0501 | *-0.3053* | -0.6570 | 0.3741 | 0.0932 | O75445 | USH2A |
| 0.3658 | 0.5146 | *1.0697* | 0.6500 | 0.3710 | 0.0936 | P60983;G3V4P8;G3V3X4;M0QYG8;M0QYJ8;M0R0C1;M0R1D2;O60234 | GMFB |
| -0.7799 | -3.1052 | *-2.9595* | -2.2816 | 1.3025 | 0.0936 | P17858 | PFKL |
| -0.4029 | -0.5622 | *-0.1439* | -0.3697 | 0.2111 | 0.0937 | P17066 | HSPA6 |
| -1.5106 | -0.9210 | *-2.8977* | -1.7764 | 1.0148 | 0.0937 | Q13164;C9JUK9;J3KT61;S4R3I1 | MAPK7 |
| 0.3259 | 0.2759 | *0.7424* | 0.4481 | 0.2562 | 0.0939 | H0Y6H0;Q8NB78;Q08EI0 | KDM1B |
| 1.0877 | 1.0803 | *2.6871* | 1.6184 | 0.9255 | 0.0939 | Q9Y5G4 | PCDHGA9 |
| -0.8759 | -1.0230 | *-2.3451* | -1.4147 | 0.8091 | 0.0939 | Q96C19 | EFHD2 |
| -0.3976 | -0.2720 | *-0.8118* | -0.4938 | 0.2825 | 0.0939 | Q6IQ55;A0A0B4J292;H3BTY5;Q8IWY7 | TTBK2 |
| 2.2686 | 1.3212 | *0.6564* | 1.4154 | 0.8102 | 0.0941 | Q15928 | ZNF141 |
| -0.4677 | -0.4549 | *-1.1455* | -0.6894 | 0.3950 | 0.0942 | E5RGN3;E5RIM7;O00244 | ATOX1 |

| L100/ CTRL1 | L100/ CTRL2 | L100/ CTRL3 | Mean | SD | T-test p-value | Accession | Gene Symbol |
|---|---|---|---|---|---|---|---|
| 0.4216 | 1.2823 | *0.6374* | 0.7804 | 0.4478 | 0.0945 | Q9Y4E6;A2RRE0;K7EMB8 | WDR7 |
| -1.1012 | -4.1653 | *-4.4656* | -3.2440 | 1.8618 | 0.0945 | Q8IWZ3;E9PDP5;H0Y4P6;H3BLS9;H7C2F5 | ANKHD1 |
| 1.2500 | 0.3133 | *0.9083* | 0.8239 | 0.4740 | 0.0949 | O75145;R4GN36;R4GNF1 | PPFIA3 |
| -0.2746 | -0.5021 | *-0.9155* | -0.5641 | 0.3249 | 0.0951 | Q9BZV1;K7EP32;K7ELN1 | UBXN6 |
| 0.0939 | 0.3364 | *0.3925* | 0.2743 | 0.1587 | 0.0959 | A0A2R8Y7Y4;Q8N8A2;H7C209;H7C4A0 | ANKRD44 |
| -0.2608 | -0.5336 | *-0.9185* | -0.5710 | 0.3305 | 0.0959 | C9JRR5;F8WCG5;Q3SYG4 | BBS9 |
| -1.0941 | -0.4547 | *-1.7185* | -1.0891 | 0.6319 | 0.0963 | Q9Y285;K7ER00;K7ER16;K7EK06 | FARSA |
| -1.0543 | -1.6775 | *-3.3513* | -2.0277 | 1.1879 | 0.0979 | E9PNW5;A0A1W2PRI9 | C4orf50 |
| 1.5207 | 1.6101 | *3.9687* | 2.3665 | 1.3883 | 0.0981 | P54764;E9PG71;C9JFX5;C9JIX8;F5GZZ5;F8WBU0 | EPHA4 |
| -1.4968 | -0.3825 | *-1.6395* | -1.1729 | 0.6882 | 0.0982 | Q96SN8;A0A0A0MRG9;B1AMJ5;F8WCI3 | CDK5RAP2 |
| 0.5220 | 1.9099 | *2.2712* | 1.5677 | 0.9234 | 0.0988 | Q8WX94;A0A0G2JM25;A0A0G2JMB6;A0A0G2JNK1;A0A0G2JNK3;A0A0G2JPH3;K7ERG0 | NLRP7 |
| -0.6451 | -0.2187 | *-0.2944* | -0.3861 | 0.2275 | 0.0989 | Q99497;K7ELW0;K7EN27 | PARK7 |
| -0.6491 | -0.2847 | *-0.2279* | -0.3872 | 0.2286 | 0.0992 | Q92522 | H1FX |
| -0.9244 | -1.3102 | *-0.3142* | -0.8496 | 0.5022 | 0.0994 | Q6ZW49 | PAXIP1 |
| -1.7711 | -0.8220 | *-0.5865* | -1.0598 | 0.6271 | 0.0996 | Q9BZE9;J3QR50;J3QRW3;C9JAL9;J3KRG1 | ASPSCR1 |
| 0.1702 | 0.4255 | *0.6698* | 0.4219 | 0.2498 | 0.0997 | P46940;H0YLE8;A0A0J9YXZ5 | IQGAP1 |
| -2.2048 | -0.5009 | *-1.8258* | -1.5105 | 0.8946 | 0.0997 | O94933;C9K0R4 | SLITRK3 |
| 0.2050 | 0.4630 | *0.7744* | 0.4808 | 0.2851 | 0.0999 | P47756;B1AK85;B1AK87;B1AK88 | CAPZB |
| 0.8757 | 3.3296 | *1.9848* | 2.0634 | 1.2288 | 0.1007 | Q6PGP7 | TTC37 |
| -0.1820 | -0.7236 | *-0.8143* | -0.5733 | 0.3419 | 0.1009 | P35221;G3XAM7;E5RIB1;E5RG03;E5RGY6;E5RHV7;E5RIE0;E5RGD2;E5RJL0;E5RJ41;E5RFM3;E5RFM5;E5RJP7;A0A087WZL6;E5RFG3;E5RFK9;E5RGU3;E5RHJ5;E5RGG4;E5RGS1;E5RIT8;E5RJ43;E5RJZ2 | CTNNA1 |

| L100/ CTRL1 | L100/ CTRL2 | L100/ CTRL3 | Mean | SD | T-test p-value | Accession | Gene Symbol |
|---|---|---|---|---|---|---|---|
| -0.3140 | -0.0700 | *-0.2682* | -0.2174 | 0.1297 | 0.1010 | A0A2Q3DQE3;Q13555;Q5SWX3;Q8WU40;H0Y6G2 | CAMK2G |
| -1.0735 | -1.6893 | *-3.4654* | -2.0761 | 1.2420 | 0.1015 | E9PR03 | IRF7 |
| 0.1346 | 0.3590 | *0.5508* | 0.3481 | 0.2083 | 0.1015 | Q96JN2;C9IYI5;C9J884;C9JAD8;C9JE17;C9JU31 | CCDC136 |
| -0.5779 | -0.5887 | *-0.1339* | -0.4335 | 0.2595 | 0.1016 | Q9NP80 | PNPLA8 |
| 2.1549 | 3.9659 | *1.1079* | 2.4096 | 1.4459 | 0.1020 | J3KSM2;Q96A59 | MARVELD3 |
| -1.1357 | -0.2721 | *-1.2297* | -0.8791 | 0.5278 | 0.1021 | M0QZR4;Q92888;M0QYC1;M0R2C7 | ARHGEF1 |
| -0.4211 | -0.8540 | *-1.5389* | -0.9380 | 0.5636 | 0.1022 | P11234;C9J6B1 | RALB |
| -0.4515 | -0.6681 | *-1.4210* | -0.8469 | 0.5089 | 0.1022 | B7ZAP0;F5H8L0;A0A0U1RQV7;E9PS63 | RABGAP1L |
| -1.7372 | -0.3837 | *-1.3577* | -1.1595 | 0.6982 | 0.1026 | Q9H9B4;D6RFI0;S4R2X2;D6RDG7;D6RAE9;H0Y9J5 | SFXN1 |
| 0.9051 | 2.5053 | *1.0250* | 1.4785 | 0.8913 | 0.1028 | A0RZB6;D6REA1;Q9H173 | SIL1 |
| -3.0611 | -0.9822 | *-1.4124* | -1.8186 | 1.0974 | 0.1030 | Q71DI3 | HIST2H3A |
| -1.5809 | -0.9714 | *-3.2221* | -1.9248 | 1.1641 | 0.1034 | A0A0G2JQ62;Q8NF37;A0A0G2JRI7 | LPCAT1 |
| -0.4426 | -1.3929 | *-1.9555* | -1.2637 | 0.7647 | 0.1035 | Q6VY07;B4DF77;H0YCU5;E9PNG7;E9PPK2 | PACS1 |
| -0.2122 | -0.6000 | *-0.9049* | -0.5724 | 0.3472 | 0.1039 | Q86TI0;H0Y8P0;C9JIE2;H0YA01;H7C0Z7;H7C380 | TBC1D1 |
| -0.5304 | -0.1401 | *-0.2996* | -0.3233 | 0.1962 | 0.1040 | Q9UNM6;A0A087WUL9;J3KNQ3;E9PL38;H0YD73;E9PQG3;E9PPD2 | PSMD13 |
| 0.5019 | 0.6435 | *0.1394* | 0.4283 | 0.2600 | 0.1040 | Q9P0L2;A0A087X0I6;B4DIB3 | MARK1 |
| -0.1180 | -0.0330 | *-0.1526* | -0.1012 | 0.0615 | 0.1042 | E9PKF4;J3KNW4 | FHL2 |
| 2.7997 | 1.6121 | *0.7238* | 1.7118 | 1.0416 | 0.1044 | Q9H3S7 | PTPN23 |
| 1.8416 | 2.0061 | *0.4260* | 1.4245 | 0.8687 | 0.1048 | Q9P2G1 | ANKIB1 |
| 1.2512 | 0.2703 | *0.9415* | 0.8210 | 0.5014 | 0.1051 | O76041;Q5JUU7 | NEBL |
| 0.8852 | 2.6004 | *1.1000* | 1.5285 | 0.9344 | 0.1053 | Q01850 | CDR2 |
| -0.3200 | -0.4234 | *-0.9719* | -0.5718 | 0.3503 | 0.1057 | Q8IV33 | KIAA0825 |
| -0.3047 | -0.7694 | *-1.2579* | -0.7773 | 0.4766 | 0.1058 | Q4LDE5;A0A0A0MSD0 | SVEP1 |
| 0.1924 | 0.3569 | *0.0959* | 0.2151 | 0.1320 | 0.1060 | P13804;H0YLU7;H0YK49;H0YKF0;H0YNX6;H0YL12;H0YL83 | ETFA |

| L100/ CTRL1 | L100/ CTRL2 | L100/ CTRL3 | Mean | SD | T-test p-value | Accession | Gene Symbol |
|---|---|---|---|---|---|---|---|
| 0.3257 | 0.4928 | *1.0596* | 0.6260 | 0.3846 | 0.1062 | O75150;H3BP71;A0A087WTK2;H3BS50 | RNF40 |
| -0.3252 | -0.0679 | *-0.3038* | -0.2323 | 0.1428 | 0.1062 | Q86VP6;A0A0C4DGH5;H0YH27 | CAND1 |
| -0.2346 | -1.0804 | *-0.7744* | -0.6965 | 0.4282 | 0.1063 | P08603 | CFH |
| 0.2957 | 0.3256 | *0.8233* | 0.4815 | 0.2964 | 0.1065 | E7EQ29;P16278;C9J4G9 | GLB1 |
| -0.1681 | -0.6841 | *-0.4017* | -0.4179 | 0.2584 | 0.1073 | Q9NYB0 | TERF2IP |
| -0.5170 | -0.1461 | *-0.2591* | -0.3074 | 0.1901 | 0.1074 | P51610;A6NEM2;H7C1C4 | HCFC1 |
| 0.5785 | 0.1972 | *0.8798* | 0.5518 | 0.3421 | 0.1078 | A8MTJ3 | GNAT3 |
| -0.3021 | -0.0674 | *-0.1982* | -0.1892 | 0.1176 | 0.1082 | Q15435;H7C003;C9JD73;B5MBZ8;C9J177;C9JRC4;H7C3Q5 | PPP1R7 |
| -0.4052 | -0.1868 | *-0.7334* | -0.4418 | 0.2751 | 0.1086 | O43852;H0Y875 | CALU |
| -0.5413 | -1.0887 | *-0.3050* | -0.6450 | 0.4020 | 0.1088 | Q99962 | SH3GL2 |
| -0.1019 | -0.5140 | *-0.4481* | -0.3547 | 0.2214 | 0.1091 | Q9UEU0;H0YJL5;H0YJJ5 | VTI1B |
| -0.6053 | -2.6748 | *-1.6913* | -1.6571 | 1.0352 | 0.1092 | Q9BQI6 | SLF1 |
| 0.9858 | 4.0457 | *2.3320* | 2.4545 | 1.5336 | 0.1092 | Q12770;C9JFY0;C9JQ35;D6RA39;F8W921;F8W9W7;F8WDP3 | SCAP |
| -1.8270 | -0.4314 | *-2.1946* | -1.4843 | 0.9302 | 0.1098 | J3KTF0 | FASN |
| -1.0114 | -0.2391 | *-0.5990* | -0.6165 | 0.3865 | 0.1098 | Q99536;K7ERT7;K7EJM4;K7ESA3;K7EM19;K7ENX2 | VAT1 |
| 0.3167 | 0.3218 | *0.8637* | 0.5007 | 0.3143 | 0.1101 | Q9NSK0;H7C4M1;C9JZE5;C9JQU1;C9JXT5;C9K0D5 | KLC4 |
| 0.1356 | 0.0282 | *0.1430* | 0.1022 | 0.0642 | 0.1103 | O15068;A2A3H1;F8WBK7 | MCF2L |
| 1.2634 | 2.3277 | *0.5902* | 1.3938 | 0.8760 | 0.1103 | Q8N103 | TAGAP |
| -0.2367 | -1.0481 | *-0.6491* | -0.6446 | 0.4057 | 0.1106 | P37837 | TALDO1 |
| 0.1422 | 0.7316 | *0.6028* | 0.4922 | 0.3099 | 0.1106 | A0A0C4DH07;Q8N2S1 | LTBP4 |
| -0.2026 | -1.0359 | *-0.8148* | -0.6844 | 0.4317 | 0.1110 | Q9NYD6 | HOXC10 |
| -1.0067 | -0.1931 | *-0.8099* | -0.6699 | 0.4245 | 0.1118 | Q14566 | MCM6 |
| -0.3292 | -0.2025 | *-0.7085* | -0.4134 | 0.2633 | 0.1128 | Q5VYK3;J3KN16;R4GMY1 | ECPAS |
| -0.7962 | -1.1685 | *-0.2374* | -0.7340 | 0.4686 | 0.1132 | O60566 | BUB1B |
| 0.0983 | 0.5324 | *0.4661* | 0.3656 | 0.2339 | 0.1136 | Q9UPV0;E9PI05;E9PLS8 | CEP164 |
| 0.2884 | 1.5680 | *1.3805* | 1.0790 | 0.6910 | 0.1138 | Q5T6L9;K7EPX8;K7ER62;K7ERF7 | ERMARD |
| 0.2240 | 0.0979 | *0.4077* | 0.2432 | 0.1558 | 0.1139 | A0A0G2JL69;B4DQI1;E9PDZ0;P06681;A0A0G2JK28;H0Y868 | C2 |

| L100/ CTRL1 | L100/ CTRL2 | L100/ CTRL3 | Mean | SD | T-test p-value | Accession | Gene Symbol |
|---|---|---|---|---|---|---|---|
| -1.1669 | -0.7146 | *-2.5211* | -1.4676 | 0.9400 | 0.1139 | Q08174;D6RAX3;D6RBG2 | PCDH1 |
| -0.2472 | -0.6619 | *-0.2320* | -0.3804 | 0.2440 | 0.1141 | P17612;Q15136;K7ERP6;P22612 | PRKACA |
| 0.3036 | 0.1710 | *0.0709* | 0.1818 | 0.1167 | 0.1143 | P38159;H0Y6E7;H3BT71;H3BUY5;Q96E39;H3BR27;A0A1B0GUK8;H3BNC1 | RBMX |
| -0.6597 | -0.9324 | *-0.1823* | -0.5915 | 0.3797 | 0.1143 | Q96HP0;K7ESB7;K7EKT0;K7EP20 | DOCK6 |
| -1.3726 | -1.0879 | *-3.3827* | -1.9478 | 1.2508 | 0.1144 | P11217 | PYGM |
| -0.4093 | -2.1352 | *-2.2173* | -1.5872 | 1.0210 | 0.1147 | A0A024RCV8;A0A0G2JK42;A0A0G2JK88;A0A140T982;A0A140T9J3;H7C2J4;O43196;A2ABE9;A0A0G2JHB4;A0A0G2JIC1;A0A0G2JJ70;A0A140T927;A0A140T9I9;A0A140T9P6;A2ABF0 | MSH5-SAPCD1 |
| 0.5517 | 0.3831 | *0.1082* | 0.3477 | 0.2239 | 0.1149 | Q14152 | EIF3A |
| 1.2587 | 2.1990 | *4.6249* | 2.6942 | 1.7369 | 0.1151 | O14939 | PLD2 |
| -0.2156 | -0.9128 | *-0.5012* | -0.5432 | 0.3505 | 0.1153 | O95613;H7C2A3 | PCNT |
| 0.9944 | 1.1230 | *0.2007* | 0.7727 | 0.4995 | 0.1156 | Q9NR99;G3V423;G3V456;G3V5U1 | MXRA5 |
| -1.2199 | -1.4580 | *-0.2612* | -0.9797 | 0.6335 | 0.1157 | O15126;A0A087WXB0;A0A087WZA6;A0A087WTX8;A0A087WU14 | SCAMP1 |
| 0.2808 | 1.0313 | *1.4631* | 0.9251 | 0.5982 | 0.1157 | Q96M96;B7Z493;E9PJX4;F8W1R0;E9PNX0;E9PQT1;F8VVF1;J3KSS3 | FGD4 |
| 0.2372 | 0.9466 | *0.4851* | 0.5563 | 0.3600 | 0.1158 | P52943;H0YFA4;H0YHD8 | CRIP2 |
| 2.3407 | 1.5013 | *0.4792* | 1.4404 | 0.9323 | 0.1159 | Q99698 | LYST |
| -0.7008 | -2.7378 | *-3.7458* | -2.3948 | 1.5512 | 0.1160 | F5GX29;H0YKE7;H0YNG9;Q99653;H0YLX1 | CHP1 |
| -2.9097 | -0.5157 | *-2.6622* | -2.0292 | 1.3166 | 0.1163 | Q13574;E9PNL8 | DGKZ |
| -0.1415 | -0.5031 | *-0.2280* | -0.2909 | 0.1888 | 0.1164 | Q9H799 | CPLANE1 |
| -0.2007 | -1.0749 | *-1.1224* | -0.7993 | 0.5190 | 0.1165 | O75582 | RPS6KA5 |
| -0.8818 | -1.4876 | *-0.3225* | -0.8973 | 0.5827 | 0.1165 | O95716;H0YMN7;M0R257 | RAB3D |
| -0.7120 | -3.6278 | *-2.4223* | -2.2540 | 1.4652 | 0.1167 | P00338;F5GXY2;F5GYU2;F5GXH2;F5H5J4;F5H6W8;F5GZQ4 | LDHA |

| L100/ CTRL1 | L100/ CTRL2 | L100/ CTRL3 | Mean | SD | T-test p-value | Accession | Gene Symbol |
|---|---|---|---|---|---|---|---|
| -0.5807 | -0.2030 | *-0.9594* | -0.5810 | 0.3782 | 0.1170 | A0A0A0MTN0;Q13617;Q5T2B5;Q5T2B7 | CUL2 |
| -0.3269 | -1.1689 | *-0.5291* | -0.6750 | 0.4396 | 0.1170 | A0A1B0GTJ8;A0A1B0GVH0 | ARID1B |
| 1.3904 | 1.2226 | *3.6637* | 2.0923 | 1.3635 | 0.1172 | Q7Z7M1;A0A087WWT7;A0A1W2PRP3 | ADGRD2 |
| -1.8125 | -0.3650 | *-2.1005* | -1.4260 | 0.9301 | 0.1174 | Q96HE7;G3V3E6;G3V5B3;G3V503 | ERO1A |
| 0.7608 | 1.9392 | *0.6210* | 1.1070 | 0.7241 | 0.1179 | Q9P0I2;S4R3U9 | EMC3 |
| -0.5453 | -0.1249 | *-0.2955* | -0.3219 | 0.2114 | 0.1187 | P54577;A0A0C4DGZ5 | YARS |
| -1.9397 | -1.6970 | *-0.3286* | -1.3218 | 0.8686 | 0.1188 | Q9ULU8;F1T0E5;H7C4T6;H7C4P2 | CADPS |
| -0.6812 | -0.4613 | *-1.5934* | -0.9120 | 0.6003 | 0.1192 | Q14155;A0A2R8YG42;B1ALK7;E9PDQ5;E7EUY6;E7ENL8 | ARHGEF7 |
| -0.7067 | -0.2536 | *-0.2442* | -0.4015 | 0.2644 | 0.1192 | Q92734;Q05BK6;C9JJP5;C9JUE0;C9JTY3 | TFG |
| -2.8397 | -1.5272 | *-0.6496* | -1.6722 | 1.1022 | 0.1194 | P16615;H7C5W9;J3QSY6 | ATP2A2 |
| 1.2710 | 0.3421 | *0.5789* | 0.7307 | 0.4827 | 0.1199 | Q8TC12;G3V2G6;H0YIZ8;G3V234;H0YJ10;H0YJ46 | RDH11 |
| -0.7051 | -0.1298 | *-0.4818* | -0.4389 | 0.2901 | 0.1199 | E9PDF6;O43795;E7EQD9;C9JYW1;C9JUP5 | MYO1B |
| -0.3982 | -0.2334 | *-0.8717* | -0.5011 | 0.3313 | 0.1201 | P40926;G3XAL0 | MDH2 |
| 0.4254 | 1.1824 | *2.0290* | 1.2123 | 0.8022 | 0.1202 | P53597 | SUCLG1 |
| -0.2979 | -0.9231 | *-1.5022* | -0.9077 | 0.6023 | 0.1207 | O43896 | KIF1C |
| 0.2662 | 1.5581 | *1.5960* | 1.1401 | 0.7570 | 0.1209 | Q6NSJ2 | PHLDB3 |
| -0.4302 | -2.6326 | *-2.4178* | -1.8269 | 1.2143 | 0.1211 | Q86XL3;F5H6J0 | ANKLE2 |
| -0.2025 | -0.9209 | *-1.2023* | -0.7752 | 0.5156 | 0.1212 | P26639 | TARS |
| -1.7283 | -1.0842 | *-0.3333* | -1.0486 | 0.6982 | 0.1214 | Q6ZRP7;H0Y430 | QSOX2 |
| -0.3842 | -0.0626 | *-0.3203* | -0.2557 | 0.1703 | 0.1214 | Q9P1U1;A0A0A0MTI9;Q9C0K3;H7C4J1;C9IZN3 | ACTR3B |
| -1.3541 | -3.4165 | *-1.0395* | -1.9367 | 1.2912 | 0.1217 | E9PNC0 | POLD3 |
| 1.6518 | 0.4047 | *2.3257* | 1.4608 | 0.9747 | 0.1219 | Q16760;H7C0L1;C9JY42 | DGKD |
| 0.0888 | 0.5177 | *0.3754* | 0.3273 | 0.2185 | 0.1219 | Q9P265 | DIP2B |
| -1.0709 | -0.2515 | *-1.4791* | -0.9338 | 0.6252 | 0.1225 | P21796;C9JI87 | VDAC1 |
| 4.1633 | 1.0749 | *6.0881* | 3.7754 | 2.5290 | 0.1227 | Q16795 | NDUFA9 |
| -2.2071 | -5.9670 | *-1.9279* | -3.3673 | 2.2557 | 0.1227 | Q9H8Y8 | GORASP2 |
| -0.8800 | -0.2974 | *-1.4944* | -0.8906 | 0.5986 | 0.1233 | P05023;Q5TC01;Q5TC02 | ATP1A1 |
| 1.8535 | 2.0293 | *0.3176* | 1.4001 | 0.9416 | 0.1234 | J3KNK3;Q17RQ9 | NKPD1 |
| -0.9907 | -0.1971 | *-0.5811* | -0.5896 | 0.3969 | 0.1236 | P11177;F8WF02 | PDHB |

| L100/ CTRL1 | L100/ CTRL2 | L100/ CTRL3 | Mean | SD | T-test p-value | Accession | Gene Symbol |
|---|---|---|---|---|---|---|---|
| 0.8028 | 0.1284 | *0.6273* | 0.5195 | 0.3499 | 0.1237 | O60673 | REV3L |
| -0.0825 | -0.4861 | *-0.3368* | -0.3018 | 0.2040 | 0.1245 | E9PK91;Q9NYF8;A0A1W2PQ43;E9PK09;E9PKI6;E9PQN2;E9PJA7 | BCLAF1 |
| 0.6057 | 2.1365 | *3.3585* | 2.0336 | 1.3793 | 0.1252 | Q14774 | HLX |
| 0.6826 | 1.1843 | *2.6475* | 1.5048 | 1.0209 | 0.1252 | E7ESX4;P10523 | SAG |
| -0.8436 | -0.1316 | *-0.8635* | -0.6129 | 0.4170 | 0.1258 | O00186 | STXBP3 |
| -0.0535 | -0.3290 | *-0.2343* | -0.2056 | 0.1399 | 0.1259 | A1L4H1 | SSC5D |
| 0.7745 | 3.2934 | *4.7109* | 2.9263 | 1.9937 | 0.1261 | Q9ULA0;A0A024R442;E7ETB3;E7EMB6;C9JBE1;B9ZVU2;E5RJ35;E7EPX3;F8WAN0 | DNPEP |
| -0.3261 | -0.4388 | *-1.1159* | -0.6269 | 0.4272 | 0.1261 | P02794;G3V192;G3V1D1;E9PPQ4 | FTH1 |
| 0.7930 | 2.0687 | *3.8168* | 2.2262 | 1.5181 | 0.1263 | K7EK91;Q9UFH2 | DNAH17 |
| -0.5499 | -0.1639 | *-0.9022* | -0.5387 | 0.3693 | 0.1274 | Q9NZK5;B4E3Q4;C9IZA8;F5H7J3 | ADA2 |
| -0.4238 | -0.2912 | *-0.0683* | -0.2611 | 0.1797 | 0.1282 | Q9BPU6;E7EWB4;E7ESV0;E9PHT0;A0A1C7CYY2;Q9H2K0 | DPYSL5 |
| -0.0889 | -0.6220 | *-0.5993* | -0.4368 | 0.3015 | 0.1288 | J3KPH8;Q8WUI4;C9J102;C9JNI4;F8VWY3;C9JAH2;C9JBC2;C9JEB6;C9JF80;C9JVZ1 | HDAC7 |
| 0.3323 | 0.5003 | *0.0815* | 0.3047 | 0.2107 | 0.1292 | Q5T124;X6R8K8;X6R8M6;Q5T130;X6RDY0;X6RIY5 | UBXN11 |
| -0.2460 | -0.1538 | *-0.5866* | -0.3288 | 0.2279 | 0.1297 | Q13200;C9JPC0;H7C1H2;H7C2Q3;F8WBS8 | PSMD2 |
| -0.1300 | -0.8919 | *-0.9501* | -0.6573 | 0.4576 | 0.1306 | P83881;H0Y5B4;H7BZ11;J3KQN4;H0Y3V9;H7BY91 | RPL36A |
| -0.9783 | -2.0898 | *-4.3633* | -2.4771 | 1.7254 | 0.1307 | P19086 | GNAZ |
| 0.2734 | 0.3870 | *0.9864* | 0.5489 | 0.3831 | 0.1312 | Q9NZ71;F6WH68;D6RA96;A0A2R8YD56;X6R5I7 | RTEL1 |
| -0.6451 | -0.8519 | *-0.1206* | -0.5392 | 0.3769 | 0.1315 | M0QZL7 | TUBB4A |
| -0.4565 | -1.4572 | *-0.5057* | -0.8064 | 0.5641 | 0.1316 | O15327;E7EQN9;D6RJC3;E9PCZ3;E9PG59;E9PHC0;H0YA10 | INPP4B |
| -1.6287 | -0.5365 | *-2.8954* | -1.6869 | 1.1805 | 0.1318 | Q7RTT5 | SSX7 |
| 0.0664 | 0.2668 | *0.4135* | 0.2489 | 0.1742 | 0.1318 | Q01844;A0A0D9SFL3;B0QYK0;C9JGE3;H7BY36 | EWSR1 |
| -0.5550 | -0.0896 | *-0.6696* | -0.4381 | 0.3072 | 0.1322 | O60268;H3BMU3;H3BNA7 | KIAA0513 |

| L100/ CTRL1 | L100/ CTRL2 | L100/ CTRL3 | Mean | SD | T-test p-value | Accession | Gene Symbol |
|---|---|---|---|---|---|---|---|
| -0.3132 | -0.2703 | *-0.0413* | -0.2082 | 0.1462 | 0.1324 | P27797;K7EJB9; K7EL50 | CALR |
| 0.6091 | 0.1281 | *0.3027* | 0.3466 | 0.2435 | 0.1325 | Q9NQ48;H7C488 | LZTFL1 |
| 0.3236 | 1.3033 | *2.0304* | 1.2191 | 0.8565 | 0.1326 | F8VWY2;Q9UBM8 | MGAT4C |
| -0.2687 | -1.4323 | *-1.9199* | -1.2070 | 0.8483 | 0.1327 | Q9NS86 | LANCL2 |
| 0.8856 | 0.3205 | *0.2626* | 0.4896 | 0.3442 | 0.1327 | Q14980;A0A087WY61;H0YFY6;F5H4J1;F5H6Y5;K4DIE0;F5H763;F5H0Z7;F5GZW1; F5H1L0;F5H2F3; F5H3L6;F8W6T3 | NUMA1 |
| 0.3370 | 1.8352 | *1.0270* | 1.0664 | 0.7499 | 0.1328 | Q63HM2;B5MC47;B6ZDM2;H0YIY3 | PCNX4 |
| -1.4214 | -0.3793 | *-2.3154* | -1.3721 | 0.9690 | 0.1337 | Q9UL62 | TRPC5 |
| 0.4988 | 0.0639 | *0.4652* | 0.3426 | 0.2420 | 0.1337 | A0A087WWH3;Q9NZM4 | BICRA |
| 0.3997 | 1.5370 | *0.6227* | 0.8531 | 0.6027 | 0.1338 | P58304 | VSX2 |
| -1.4157 | -0.3295 | *-2.1661* | -1.3037 | 0.9234 | 0.1343 | J3KQJ9;Q8N2E2; E5RG96 | VWDE |
| -0.5124 | -0.4718 | *-1.5150* | -0.8331 | 0.5909 | 0.1347 | P18124;A8MUD9;C9JIJ5;C9JZ88 | RPL7 |
| 0.2223 | 0.2205 | *0.0283* | 0.1570 | 0.1115 | 0.1348 | P55795 | HNRNPH2 |
| -0.4553 | -0.3374 | *-1.2128* | -0.6685 | 0.4750 | 0.1350 | P18084;V9GZ57 | ITGB5 |
| -0.6383 | -1.3951 | *-2.9584* | -1.6639 | 1.1832 | 0.1352 | Q9BZF1;F8VQX7;F8VUA7 | OSBPL8 |
| -0.6183 | -1.4258 | *-0.3380* | -0.7940 | 0.5648 | 0.1353 | Q9UI12;G3V126; H0YB41 | ATP6V1H |
| 0.7565 | 1.3478 | *0.2364* | 0.7802 | 0.5561 | 0.1357 | Q9NYF5;D6RAT6;D6RCA0;D6RDL7 | FAM13B |
| -1.5737 | -0.3067 | *-0.8057* | -0.8954 | 0.6382 | 0.1357 | P42345;B1AKP8 | MTOR |
| -0.3319 | -0.6646 | *-1.4737* | -0.8234 | 0.5872 | 0.1358 | Q8NEV4;A0A2R8Y4D5;F5H0U9 | MYO3A |
| -0.2178 | -0.3525 | *-0.8649* | -0.4784 | 0.3415 | 0.1360 | P10827;J3KTF3; J3QRA9;J3QRW5 | THRA |
| -0.8523 | -0.8489 | *-0.1059* | -0.6024 | 0.4300 | 0.1360 | Q5T5P2 | KIAA1217 |
| -0.7717 | -1.4050 | *-0.2496* | -0.8088 | 0.5786 | 0.1365 | B0YIW6;P48444; Q6P1Q5 | ARCN1 |
| 0.2249 | 0.0974 | *0.4689* | 0.2637 | 0.1888 | 0.1366 | Q96L34;Q6IPE9; K7EK17;K7EKG8;K7EN95 | MARK4 |
| -0.9631 | -0.1209 | *-0.9990* | -0.6943 | 0.4970 | 0.1366 | P52701;A0A087WWJ1;C9J7Y7;C9J8Y8;C9JH55 | MSH6 |
| -0.1938 | -0.9225 | *-0.4358* | -0.5174 | 0.3712 | 0.1371 | O43865 | AHCYL1 |
| 0.6854 | 0.9203 | *2.4863* | 1.3640 | 0.9790 | 0.1372 | P31350;A0A286YFD6;C9JXC1 | RRM2 |
| -0.3851 | -0.3156 | *-1.0910* | -0.5973 | 0.4290 | 0.1374 | P46777;A0A2R8Y6J3;Q5T7N0;A0A2R8Y4A2 | RPL5 |

| L100/ CTRL1 | L100/ CTRL2 | L100/ CTRL3 | Mean | SD | T-test p-value | Accession | Gene Symbol |
|---|---|---|---|---|---|---|---|
| -0.1259 | -0.0950 | *-0.3429* | -0.1879 | 0.1351 | 0.1376 | H7C3G9;Q9UJ70;C9JEV6;H0YEB7;H7C1L7;H0YE82;H0YF44 | NAGK |
| 0.3200 | 0.1542 | *0.0652* | 0.1798 | 0.1293 | 0.1377 | Q96T58;F6WRY4;H0Y5U7 | SPEN |
| -2.4275 | -0.3397 | *-2.8923* | -1.8865 | 1.3595 | 0.1381 | Q5T123;Q9H299 | SH3BGRL3 |
| -0.2907 | -0.0961 | *-0.5405* | -0.3091 | 0.2228 | 0.1381 | Q5JPF3;E9PJI0;A0A087WX87;A0A087WYV7;H0YDI7 | ANKRD36C |
| -1.4031 | -2.9063 | *-0.5873* | -1.6322 | 1.1764 | 0.1381 | A0A2R8Y3T0;G3V5G6;O43405 | COCH |
| -0.0991 | -0.6655 | *-0.8284* | -0.5310 | 0.3828 | 0.1382 | A0A1B0GTS2 | OBSCN |
| -0.7362 | -1.0888 | *-0.1487* | -0.6579 | 0.4749 | 0.1385 | P20339 | RAB5A |
| -0.3127 | -0.7061 | *-1.5044* | -0.8411 | 0.6072 | 0.1385 | Q96EI5;A2RQR6;D6RD44;D6RCE9;D6RIH3;D6RHZ1;J3KQY4 | TCEAL4 |
| -0.6651 | -0.5068 | *-1.8294* | -1.0004 | 0.7223 | 0.1385 | O95271;E7EQ52;A0A0C4DGE3 | TNKS |
| -1.8934 | -0.2724 | *-2.3250* | -1.4970 | 1.0822 | 0.1388 | K7ES00;K7EK07;P84243 | H3F3B |
| -2.3717 | -2.1532 | *-0.2605* | -1.5951 | 1.1609 | 0.1403 | D3YTB1;F8W727;P62910;D3YTI8 | RPL32 |
| 0.8997 | 3.6123 | *1.4209* | 1.9776 | 1.4395 | 0.1404 | J3KNV4;G3V2C6 | ITGA7 |
| 0.0643 | 0.4945 | *0.5815* | 0.3801 | 0.2769 | 0.1406 | Q8IZQ1;D6RJE4 | WDFY3 |
| -0.0978 | -0.5103 | *-0.7513* | -0.4531 | 0.3305 | 0.1408 | Q6T4P5 | PLPPR3 |
| -0.1167 | -0.9176 | *-1.0677* | -0.7006 | 0.5113 | 0.1409 | Q8TAT2 | FGFBP3 |
| 0.4714 | 0.1164 | *0.7830* | 0.4569 | 0.3336 | 0.1410 | E7EUN2;A0A087X1U1;F5GXM9;Q9UPQ3;A0A087X044;C9J8Z2;C9J975;E7ESL9;H7C4F1 | AGAP1 |
| -0.0809 | -0.5112 | *-0.2910* | -0.2944 | 0.2152 | 0.1413 | P40939;H0YFD6;A0A2R8Y4F5;A0A2R8YG21;A0A2R8Y688 | HADHA |
| 1.0237 | 0.8022 | *0.1156* | 0.6472 | 0.4735 | 0.1415 | Q9Y3R5 | DOP1B |
| 0.2297 | 0.3383 | *0.8978* | 0.4886 | 0.3585 | 0.1421 | Q13619;A0A0A0MR50 | CUL4A |
| 1.0810 | 0.2015 | *0.5321* | 0.6049 | 0.4442 | 0.1424 | Q96L73;D6RA90;D6RBP3 | NSD1 |
| 0.3225 | 0.9087 | *0.2472* | 0.4928 | 0.3622 | 0.1425 | Q5JSZ5;Q5JSZ9 | PRRC2B |
| -0.4457 | -0.3064 | *-1.1926* | -0.6482 | 0.4766 | 0.1426 | Q13263;M0R0K9;M0R3C0;M0QZE6 | TRIM28 |
| -0.7052 | -0.7628 | *-0.0790* | -0.5156 | 0.3793 | 0.1427 | P14314;K7ELL7;A0A0C4DGP4;K7EPW7;K7EJ70;K7EKX1 | PRKCSH |
| 3.1092 | 2.9915 | *0.3234* | 2.1414 | 1.5755 | 0.1428 | P49796 | RGS3 |
| -0.0674 | -0.1964 | *-0.3833* | -0.2157 | 0.1588 | 0.1429 | P19022;C9J8J8;C9J126;A0A087 | CDH2 |

| L100/ CTRL1 | L100/ CTRL2 | L100/ CTRL3 | Mean | SD | T-test p-value | Accession | Gene Symbol |
|---|---|---|---|---|---|---|---|
| | | | | | | WX99;A0A0G2JRA4;P55283 | |
| -0.4345 | -0.7473 | *-0.1115* | -0.4311 | 0.3179 | 0.1433 | Q16555;C9J1F2;A0A1C7CYX9 | DPYSL2 |
| -0.0365 | -0.3541 | *-0.3507* | -0.2471 | 0.1824 | 0.1436 | P98172 | EFNB1 |
| -0.3104 | -2.1816 | *-1.3129* | -1.2683 | 0.9364 | 0.1436 | A0A087WSV8;P80303;E9PKG6;Q2L696;A0A2R8Y6G7;E9PLE9;E9PLR0;H0YEG8;E9PJP3;E9PM22;H0YD18 | NUCB2 |
| -2.4072 | -3.8463 | *-0.5186* | -2.2574 | 1.6689 | 0.1439 | Q96J02 | ITCH |
| 0.9486 | 0.2761 | *1.7326* | 0.9858 | 0.7289 | 0.1439 | J3KNE0;A6NKT7 | RGPD3 |
| -0.2900 | -2.3030 | *-2.8014* | -1.7981 | 1.3297 | 0.1439 | J3KSW6;Q8IX18;J3KSX9;J3KTK0 | DHX40 |
| -0.6183 | -2.2685 | *-0.7893* | -1.2254 | 0.9074 | 0.1443 | O96019;H7C5S0 | ACTL6A |
| -1.2768 | -0.3238 | *-0.4727* | -0.6911 | 0.5126 | 0.1446 | P25685;M0R080 | DNAJB1 |
| -3.0526 | -1.3050 | *-6.6415* | -3.6664 | 2.7206 | 0.1447 | Q96F07;A0A087WWZ1;A0A087WTQ3;A0A087WVE1 | CYFIP2 |
| -1.2851 | -0.8220 | *-3.3687* | -1.8253 | 1.3565 | 0.1451 | P98174 | FGD1 |
| -1.1019 | -0.1070 | *-1.0145* | -0.7411 | 0.5509 | 0.1451 | Q9Y266;A0A0A0MSS4;A0A0A0MSU9 | NUDC |
| 0.1702 | 0.5696 | *1.0612* | 0.6003 | 0.4463 | 0.1451 | O15254 | ACOX3 |
| -0.0379 | -0.3642 | *-0.2731* | -0.2251 | 0.1684 | 0.1466 | Q9UNH7;A0A0A0MRI2;G3V5X9;G3V4Z5;H0YJF8 | SNX6 |
| -0.1334 | -1.3831 | *-1.4208* | -0.9791 | 0.7326 | 0.1467 | P61160;F5H6T1 | ACTR2 |
| 0.5855 | 2.1252 | *0.7133* | 1.1413 | 0.8544 | 0.1468 | O00763;F8W8T8;H0YGH5;A0A087WUA1 | ACACB |
| 2.8312 | 1.6022 | *0.4082* | 1.6139 | 1.2116 | 0.1474 | O95104 | SCAF4 |
| -0.1921 | -0.4690 | *-0.1023* | -0.2545 | 0.1911 | 0.1475 | P00367 | GLUD1 |
| -0.2517 | -0.1283 | *-0.6024* | -0.3275 | 0.2460 | 0.1475 | Q9H2U1;E7EWK3 | DHX36 |
| -1.1648 | -0.1114 | *-0.9254* | -0.7339 | 0.5522 | 0.1480 | A0A0U1RRB6;Q9Y2D4;J3QT38 | EXOC6B |
| -1.3275 | -0.8333 | *-3.5039* | -1.8882 | 1.4209 | 0.1480 | B1AHC2;B1AHF8;F8WCV8;P85298;F8W6F4 | ARHGAP8 |
| -1.0414 | -0.2422 | *-1.7778* | -1.0205 | 0.7680 | 0.1480 | P61758;B4DWR3 | VBP1 |
| 1.3620 | 0.6008 | *0.2668* | 0.7432 | 0.5613 | 0.1488 | Q9UKI9 | POU2F3 |
| 0.0634 | 0.4737 | *0.2779* | 0.2717 | 0.2052 | 0.1489 | Q9Y2K5;A0A0U1RRA6;B5MCG9;B5MCU0;H0YIX9;V9GYY9 | R3HDM2 |
| -1.2354 | -0.4090 | *-0.3374* | -0.6606 | 0.4991 | 0.1489 | A0A087WUM2;Q6ZMR3 | LDHAL6A |
| -0.2368 | -0.6099 | *-0.1390* | -0.3286 | 0.2485 | 0.1492 | F5GXQ8;H0Y325;H0Y326 | SYNE1 |

| L100/ CTRL1 | L100/ CTRL2 | L100/ CTRL3 | Mean | SD | T-test p-value | Accession | Gene Symbol |
|---|---|---|---|---|---|---|---|
| -0.3271 | -0.3160 | *-1.0718* | -0.5716 | 0.4332 | 0.1496 | Q9C093;A0A1B0GWC1;H0Y989;H0YAC0;A0A1B0GWD8 | SPEF2 |
| 0.1724 | 2.0436 | *1.8360* | 1.3507 | 1.0257 | 0.1501 | P68036;A0A1B0GUS4 | UBE2L3 |
| 0.3836 | 0.7523 | *0.1194* | 0.4184 | 0.3179 | 0.1502 | Q05513;E9PBE1 | PRKCZ |
| -0.3440 | -3.0351 | *-3.8245* | -2.4012 | 1.8248 | 0.1503 | Q9BX84 | TRPM6 |
| -0.3734 | -1.0984 | *-0.2866* | -0.5861 | 0.4457 | 0.1505 | P09543;K7ERC4;C9K0L8;K7EN66;K7ERZ0 | CNP |
| -0.0695 | -0.2977 | *-0.1114* | -0.1595 | 0.1215 | 0.1508 | Q9UQM7 | CAMK2A |
| 0.2028 | 1.0249 | *1.6735* | 0.9671 | 0.7370 | 0.1510 | Q9NXG6 | P4HTM |
| -0.7308 | -7.7774 | *-8.8552* | -5.7878 | 4.4125 | 0.1510 | Q70CQ4;A0A087WXV9 | USP31 |
| -0.2185 | -0.6540 | *-0.1727* | -0.3484 | 0.2656 | 0.1510 | P51957;E7EX48;F8WAX1 | NEK4 |
| 0.5104 | 0.2286 | *1.1771* | 0.6387 | 0.4871 | 0.1511 | A0A075B6F3;Q96JB1;H0Y7V4 | DNAH8 |
| 1.0114 | 0.3474 | *2.0819* | 1.1469 | 0.8751 | 0.1512 | Q9UN86;D6RAC7;D6RB17;D6RBW8;D6RE13;D6RGJ4;D6RBM9;D6RBR0;D6REX8 | G3BP2 |
| -0.1037 | -0.8620 | *-0.5273* | -0.4977 | 0.3800 | 0.1514 | Q01484;I6L894;E9PHW9;D6RHE1;A0A0U1RQN6;B7Z651;E9PCH6;H0Y931;H0Y8Y2;D6R9U4;H0YAG3;D6RIY9 | ANK2 |
| 0.3500 | 0.2897 | *1.0736* | 0.5711 | 0.4362 | 0.1515 | Q9UGU0;I3L1M7 | TCF20 |
| -0.3048 | -0.0590 | *-0.1313* | -0.1650 | 0.1263 | 0.1520 | C9J4R3;F8WEB4;Q96RG2 | PASK |
| -0.2927 | -0.3565 | *-0.0297* | -0.2263 | 0.1732 | 0.1520 | P18206;A0A096LPE1 | VCL |
| -3.3724 | -4.8586 | *-0.4849* | -2.9053 | 2.2239 | 0.1520 | Q96RL7;H0Y7P8 | VPS13A |
| -0.4486 | -0.4553 | *-1.5270* | -0.8103 | 0.6207 | 0.1522 | P49448 | GLUD2 |
| -0.5685 | -0.3966 | *-0.0558* | -0.3403 | 0.2609 | 0.1524 | Q9Y4A5;F2Z2U4;C9K0N1 | TRRAP |
| 1.7272 | 0.2991 | *0.8053* | 0.9439 | 0.7241 | 0.1525 | O14514;A0A2R8Y5M7 | ADGRB1 |
| 1.2339 | 0.2085 | *1.9151* | 1.1192 | 0.8591 | 0.1527 | Q15370 | ELOB |
| -0.3721 | -0.4590 | *-1.4078* | -0.7463 | 0.5745 | 0.1534 | Q9UL54 | TAOK2 |
| -0.2390 | -0.9148 | *-0.3010* | -0.4849 | 0.3736 | 0.1535 | P30101;H7BZJ3 | PDIA3 |
| 1.5618 | 0.2180 | *2.2581* | 1.3460 | 1.0370 | 0.1536 | Q86V15 | CASZ1 |
| 0.7018 | 0.2860 | *1.5790* | 0.8556 | 0.6600 | 0.1539 | Q5T670;Q7L590;C9J600 | MCM10 |
| 0.6794 | 1.6060 | *0.3102* | 0.8652 | 0.6676 | 0.1539 | Q9BZL4 | PPP1R12C |
| -0.2319 | -0.7147 | *-0.1878* | -0.3781 | 0.2923 | 0.1544 | Q13428;J3KQ96;E7ETY2;H0Y8Y7;A0A2R8Y857;H0YAB7 | TCOF1 |
| -0.2798 | -0.5146 | *-0.0693* | -0.2879 | 0.2228 | 0.1546 | P12956;B1AHC9 | XRCC6 |

| L100/ CTRL1 | L100/ CTRL2 | L100/ CTRL3 | Mean | SD | T-test p-value | Accession | Gene Symbol |
|---|---|---|---|---|---|---|---|
| -0.1420 | -0.5272 | *-0.1667* | -0.2786 | 0.2156 | 0.1546 | P05091;S4R3S4; F8VP50 | ALDH2 |
| 0.0753 | 0.9225 | *1.0420* | 0.6799 | 0.5270 | 0.1550 | Q8IXB1;A0A087 WXH7;E7EP04;Q 71S60 | DNAJC10 |
| -0.6760 | -0.3710 | *-0.0888* | -0.3786 | 0.2937 | 0.1552 | B7Z2U2;Q6ZVM7 | TOM1L2 |
| -7.8248 | -1.0832 | *-4.1357* | -4.3479 | 3.3758 | 0.1554 | B1AHB1;P33992 | MCM5 |
| -0.8874 | -0.7359 | *-2.7914* | -1.4716 | 1.1455 | 0.1560 | Q13620;K4DI93 | CUL4B |
| -0.8927 | -0.7091 | *-2.7517* | -1.4512 | 1.1301 | 0.1561 | A0A2R8Y595;A0 A2R8YFL0;P497 11 | CTCF |
| -2.0698 | -0.1492 | *-2.1852* | -1.4681 | 1.1436 | 0.1562 | Q14CN2 | CLCA4 |
| -0.0380 | -0.5596 | *-0.5549* | -0.3842 | 0.2998 | 0.1567 | P17661 | DES |
| 0.0630 | 0.6445 | *0.4191* | 0.3755 | 0.2932 | 0.1568 | Q9Y2I7;E9PDH4; C9JL08;F8WEZ0 | PIKFYVE |
| 0.2723 | 0.7180 | *0.1532* | 0.3812 | 0.2977 | 0.1569 | P16520;E9PCP0; F5H0S8;F5H100; F5H8J8 | GNB3 |
| 0.2966 | 0.4578 | *1.2880* | 0.6808 | 0.5320 | 0.1570 | Q9NUL3;A0A0A0 MTD1;E7EPX0;E 7EVJ4;E9PH62;F 8VPI7;A0A0A0M TC6;E7EVI1;A0A 0A0MTC4;E5RJN 7;G5EA18;E5RJ6 7 | STAU2 |
| -0.2812 | -0.7384 | *-0.1564* | -0.3920 | 0.3064 | 0.1570 | P10768;X6RA14; H7BZT7;U3KQT1 | ESD |
| 0.1237 | 1.6176 | *1.2377* | 0.9930 | 0.7765 | 0.1571 | Q9H269 | VPS16 |
| -0.4096 | -0.2162 | *-0.0550* | -0.2269 | 0.1775 | 0.1573 | P04179;F5GYZ5; F5H3C5;F5H4R2 ;G8JLJ2;F5GXZ9 ;G5E9P6 | SOD2 |
| -1.8252 | -0.9931 | *-4.7530* | -2.5238 | 1.9749 | 0.1573 | Q7Z3V4;S4R3H8 | UBE3B |
| -0.7269 | -0.3061 | *-0.1346* | -0.3892 | 0.3048 | 0.1575 | P28066 | PSMA5 |
| 0.3006 | 0.6785 | *0.1171* | 0.3654 | 0.2863 | 0.1576 | P08729;A0A1W2 PRP1 | KRT7 |
| 0.7563 | 0.1187 | *1.1931* | 0.6894 | 0.5403 | 0.1577 | Q9UBU9;E9PIN3 ;E9PLA7;E9PMV 7 | NXF1 |
| 0.2016 | 0.3258 | *0.9046* | 0.4773 | 0.3752 | 0.1584 | Q16568 | CARTPT |
| -0.4956 | -0.1620 | *-1.0476* | -0.5684 | 0.4472 | 0.1587 | P26196;Q8IV96 | DDX6 |
| 0.3272 | 1.3469 | *2.5311* | 1.4017 | 1.1030 | 0.1587 | O75069;A0A0C4 DFR1;A0A1B0G UZ3;G5E963 | TMCC2 |
| - 11.042 5 | -6.5573 | *-1.1915* | -6.2637 | 4.9320 | 0.1588 | A6NHK2;P62304 | SNRPE |
| 0.3209 | 0.7476 | *1.7711* | 0.9465 | 0.7453 | 0.1588 | Q92621 | NUP205 |
| -0.9777 | -0.0672 | *-1.0839* | -0.7096 | 0.5588 | 0.1589 | Q6ZNG0 | ZNF620 |
| -0.7700 | -0.4149 | *-2.0149* | -1.0666 | 0.8402 | 0.1590 | Q14697;F5H6X6; E9PKU7;E9PNH 1 | GANAB |

| L100/ CTRL1 | L100/ CTRL2 | L100/ CTRL3 | Mean | SD | T-test p-value | Accession | Gene Symbol |
|---|---|---|---|---|---|---|---|
| -0.1001 | -1.0338 | *-1.3669* | -0.8336 | 0.6567 | 0.1590 | A0A0A0MRJ0;Q5VT25;A0A0A0MRJ1 | CDC42BPA |
| -0.6077 | -0.0499 | *-0.4249* | -0.3608 | 0.2844 | 0.1590 | P19623;K7EL89;K7EQ47 | SRM |
| 0.3001 | 1.8958 | *3.0500* | 1.7486 | 1.3808 | 0.1595 | A0A2R8YH03;Q5VWN6 | FAM208B |
| 0.1223 | 1.0277 | *1.4985* | 0.8828 | 0.6994 | 0.1604 | O75146 | HIP1R |
| -0.0527 | -0.5564 | *-0.7483* | -0.4525 | 0.3593 | 0.1609 | Q969Q0 | RPL36AL |
| -0.3857 | -0.0388 | *-0.5321* | -0.3189 | 0.2534 | 0.1611 | Q16204 | CCDC6 |
| 0.2171 | 0.0856 | *0.5041* | 0.2689 | 0.2140 | 0.1614 | P05787;F8VUG2;F8W1U3 | KRT8 |
| 0.8634 | 0.9092 | *3.1468* | 1.6398 | 1.3053 | 0.1615 | Q9NVE7;A0A0G2JR38;E9PHT6;H0YA26;H0Y9E4 | PANK4 |
| -0.6630 | -2.8561 | *-5.3845* | -2.9679 | 2.3628 | 0.1616 | Q07960 | ARHGAP1 |
| 0.3268 | 0.4350 | *1.3446* | 0.7021 | 0.5590 | 0.1616 | P15907;C9J6X5;C9K0R8 | ST6GAL1 |
| -1.2611 | -0.7453 | *-3.4913* | -1.8326 | 1.4595 | 0.1617 | O95758 | PTBP3 |
| -0.6008 | -1.4245 | *-0.2483* | -0.7578 | 0.6036 | 0.1617 | Q9HCK8;A0A2R8Y840;H0YJG4;A0A2R8Y4P3;A0A2R8Y808 | CHD8 |
| -0.2719 | -1.3085 | *-2.3711* | -1.3172 | 1.0497 | 0.1618 | O60733 | PLA2G6 |
| 1.4301 | 0.5283 | *0.2953* | 0.7512 | 0.5993 | 0.1621 | O60285 | NUAK1 |
| -1.4562 | -1.8740 | *-5.9075* | -3.0792 | 2.4582 | 0.1623 | O75419 | CDC45 |
| 0.1129 | 0.1848 | *0.5226* | 0.2734 | 0.2188 | 0.1628 | P99999;C9JFR7 | CYCS |
| 0.2206 | 0.1081 | *0.5713* | 0.3000 | 0.2416 | 0.1645 | Q8TDW7;E9PQ73 | FAT3 |
| -3.2831 | -0.1553 | *-3.3764* | -2.2716 | 1.8333 | 0.1650 | Q9BZ23;V9GYZ0 | PANK2 |
| -0.0324 | -0.6669 | *-0.5704* | -0.4232 | 0.3419 | 0.1652 | Q99755;A6PW58 | PIP5K1A |
| -0.3560 | -1.0987 | *-0.2553* | -0.5700 | 0.4606 | 0.1653 | A0A0G2JRI9;A0A0G2JRM4;C9JJE7;A0A0G2JRT2 | MUC20 |
| -0.0398 | -0.7549 | *-0.6056* | -0.4667 | 0.3772 | 0.1653 | O60568;H7C2S8;H7C2V1;H7C0B8 | PLOD3 |
| 0.3687 | 0.9259 | *2.2104* | 1.1683 | 0.9445 | 0.1654 | P49792 | RANBP2 |
| -2.2587 | -0.1012 | *-2.2793* | -1.5464 | 1.2516 | 0.1657 | H0Y6T8 | RAB18 |
| 1.2433 | 2.2243 | *0.2279* | 1.2318 | 0.9983 | 0.1660 | Q9Y5E6;Q9Y5E4 | PCDHB3 |
| -0.4569 | -2.0523 | *-0.6828* | -1.0640 | 0.8633 | 0.1663 | Q9Y295;H0YI06 | DRG1 |
| -2.2681 | -1.4094 | *-6.5920* | -3.4232 | 2.7777 | 0.1664 | O75165 | DNAJC13 |
| 0.1958 | 1.2287 | *0.5240* | 0.6495 | 0.5278 | 0.1667 | P58876;P62807;Q99877;U3KQK0;O60814;Q99879;Q99880;P57053;Q96A08;A0A2R8Y619 | HIST1H2BD |
| 3.4718 | 2.3708 | *0.2310* | 2.0246 | 1.6479 | 0.1672 | Q15257;A6PVN5;F6WIT2;B7ZBQ0;Q68CR8;A6PVN9 | PTPA |
| 0.7055 | 1.3200 | *0.1430* | 0.7228 | 0.5887 | 0.1673 | Q7L014;A0A0C4DG89;D6RJA6 | DDX46 |

| L100/ CTRL1 | L100/ CTRL2 | L100/ CTRL3 | Mean | SD | T-test p-value | Accession | Gene Symbol |
|---|---|---|---|---|---|---|---|
| 0.0410 | 0.8264 | *0.6467* | 0.5047 | 0.4115 | 0.1676 | Q6ZS17;A0A0A0MTL6;H3BSX9;B5MDQ0;H3BMG9;H3BQI5;H3BSV5;H3BU12;Q2NKX8 | RIPOR1 |
| -0.0111 | -0.1599 | *-0.2110* | -0.1273 | 0.1038 | 0.1676 | Q969G3;B4DGM3;A0A2R8Y855;A0A2R8Y4T4;A0A2R8Y765;A0A2R8Y7I9;A0A2R8Y7U4;A0A2R8YES3;A0A2U3TZQ7;J3QKS7;J3QR61;H7C048;K7EMQ8;A0A2R8YD78;J3KT85;A0A2R8YEB8;J3QKX6 | SMARCE1 |
| 4.0103 | 3.1400 | *0.1922* | 2.4475 | 2.0010 | 0.1683 | E2QRD4;Q6ZRQ5;H0Y8J1 | MMS22L |
| 0.4913 | 4.4577 | *2.3256* | 2.4249 | 1.9851 | 0.1686 | O95398 | RAPGEF3 |
| -0.7616 | -0.1149 | *-1.3048* | -0.7271 | 0.5957 | 0.1688 | Q15477;H7C5N0 | SKIV2L |
| -0.0586 | -0.8059 | *-1.0994* | -0.6546 | 0.5366 | 0.1690 | P62906 | RPL10A |
| 1.2447 | 0.0450 | *1.1694* | 0.8197 | 0.6720 | 0.1690 | G3V2S6;Q9Y5K8;H0YJH8;G3V2V6;G3V559 | ATP6V1D |
| -0.1443 | -0.8751 | *-0.3554* | -0.4583 | 0.3761 | 0.1693 | Q86WI3 | NLRC5 |
| 0.9914 | 1.4546 | *4.4878* | 2.3113 | 1.8991 | 0.1696 | Q9GZU2;M0QXI3;M0QZD4;M0R155 | PEG3 |
| -0.1748 | -0.1745 | *-0.6483* | -0.3325 | 0.2734 | 0.1698 | Q08209;E7ETC2;E9PK68;E9PPC8 | PPP3CA |
| -4.9176 | -2.3857 | *-0.6019* | -2.6351 | 2.1686 | 0.1700 | A0A087WXL3;O75417 | POLQ |
| -0.2339 | -0.0399 | *-0.0921* | -0.1220 | 0.1004 | 0.1700 | M0QXL7 | ZNF208 |
| 0.6028 | 0.5294 | *2.1021* | 1.0781 | 0.8875 | 0.1701 | Q93050;B7Z2A9;B7Z641;K7EPG4;K7EM24;K7ELZ6;K7EN36 | ATP6V0A1 |
| -1.1361 | -2.0593 | *-5.8028* | -2.9994 | 2.4713 | 0.1703 | Q69YN4 | VIRMA |
| -0.0267 | -0.4101 | *-0.2673* | -0.2347 | 0.1938 | 0.1708 | Q9NUQ9;E5RI16;A0A087X178;E5RGI7;E5RIR8;E5RJE1;E5RJL8;E5RFS4;E5RHU5;E5RK61 | FAM49B |
| -0.6803 | -0.4729 | *-0.0377* | -0.3970 | 0.3279 | 0.1710 | Q7L7V1 | DHX32 |
| -4.5684 | -0.7155 | *-8.0794* | -4.4544 | 3.6833 | 0.1712 | P05198 | EIF2S1 |
| 0.3544 | 0.0737 | *0.6890* | 0.3724 | 0.3080 | 0.1713 | A0A067XG54;Q8NB49 | ATP11C |
| -0.0898 | -0.2389 | *-0.5783* | -0.3023 | 0.2503 | 0.1716 | P10809;E7EXB4;E7ESH4;C9JCQ4;C9JL19;C9J0S9;C9JF90;C9K0L6;G3V3D4;H0Y4N1 | HSPD1 |

| L100/ CTRL1 | L100/ CTRL2 | L100/ CTRL3 | Mean | SD | T-test p-value | Accession | Gene Symbol |
|---|---|---|---|---|---|---|---|
| | | | | | | ;K7ENX5;Q9BPX7;Q9HAC7;Q9Y2R4 | |
| -0.7993 | -0.5662 | *-0.0404* | -0.4686 | 0.3888 | 0.1721 | P13637;A0A0A0MT26;A0A2R8YEY8;M0R116;M0QXF2 | ATP1A3 |
| 0.0766 | 0.1412 | *0.3991* | 0.2057 | 0.1707 | 0.1721 | Q14593 | ZNF273 |
| -0.0059 | -0.2166 | *-0.2110* | -0.1445 | 0.1200 | 0.1725 | Q63HN8;A0A0A0MTR7 | RNF213 |
| 0.8462 | 2.7145 | *0.6058* | 1.3889 | 1.1544 | 0.1726 | Q14353;A0A1W2PR36 | GAMT |
| -0.6424 | -0.2275 | *-0.1220* | -0.3306 | 0.2751 | 0.1728 | P24666;G5E9R5;F2Z2Q9 | ACP1 |
| -0.1632 | -0.2282 | *-0.7328* | -0.3747 | 0.3118 | 0.1729 | Q9Y6G9;E9PHI6;C9JGM7;C9JLW1 | DYNC1LI1 |
| 1.1120 | 1.1977 | *0.0315* | 0.7804 | 0.6500 | 0.1731 | Q02880;E9PCY5 | TOP2B |
| -0.3762 | -1.7059 | *-0.5342* | -0.8721 | 0.7264 | 0.1731 | Q02818;C9JKZ2;H7BZI1;C9J3C1;C9JBD3 | NUCB1 |
| -0.4153 | -0.4031 | *-0.0104* | -0.2762 | 0.2303 | 0.1734 | O00534;B4DHS6 | VWA5A |
| -0.6119 | -5.0241 | *-2.3418* | -2.6593 | 2.2232 | 0.1741 | Q8N1I0;H0Y599;H0Y7H7;C9J637;C9J7D9 | DOCK4 |
| 0.3293 | 0.2241 | *0.0168* | 0.1901 | 0.1590 | 0.1742 | Q9UMR2;H3BQK0;F6QDS0;H3BN59;H3BMQ5 | DDX19B |
| -0.5478 | -2.3190 | *-4.8079* | -2.5582 | 2.1401 | 0.1742 | Q7Z460;F8WA11;H0Y5T1;C9J151;C9JP76 | CLASP1 |
| -0.8652 | -0.0889 | *-1.3860* | -0.7800 | 0.6527 | 0.1743 | P23378;A0A1W2PP74;A0A1W2PPB1 | GLDC |
| -0.6008 | -0.2878 | *-0.0690* | -0.3192 | 0.2673 | 0.1745 | P08195;F5GZS6;J3KPF3;F5GZI0;H0YFS2;H0YFX4;F5H0E2 | SLC3A2 |
| -0.9764 | -0.4012 | *-2.4973* | -1.2917 | 1.0830 | 0.1748 | Q9UIG0 | BAZ1B |
| -1.0808 | -1.4453 | *-4.7967* | -2.4409 | 2.0483 | 0.1751 | Q9NP79;A0A087WY55;Q5TGM0 | VTA1 |
| 2.5225 | 0.2490 | *1.2959* | 1.3558 | 1.1379 | 0.1751 | P62273;A0A2R8Y851;A0A087WTT6;A0A2R8Y6P7 | RPS29 |
| -0.1135 | -0.1758 | *-0.5479* | -0.2791 | 0.2349 | 0.1758 | P24534;C9JZW3;F2Z2G2;F8WF65 | EEF1B2 |
| -0.3338 | -0.2926 | *-0.0070* | -0.2112 | 0.1780 | 0.1762 | P53396;K7ESG8 | ACLY |
| 0.5631 | 1.8761 | *0.4176* | 0.9522 | 0.8034 | 0.1765 | Q9NZV7 | ZIM2 |
| -0.2067 | -2.9455 | *-1.7253* | -1.6258 | 1.3721 | 0.1766 | A0A2R8Y5N2;A0A2R8Y5S3;A0A2R8YFV7;Q92574;A0A2R8Y6S1;A0A2R8YGX7;A0A2R8Y6W1;A0A2R8Y7E9;A0A2R8YD74;Q59IT9;Q86WV8;A0A2R8Y5 | TSC1 |

| L100/ CTRL1 | L100/ CTRL2 | L100/ CTRL3 | Mean | SD | T-test p-value | Accession | Gene Symbol |
|---|---|---|---|---|---|---|---|
| | | | | | | F8;A0A2R8Y5J1;A0A2R8Y5Q4;A0A2R8Y6N1;A0A2R8Y6S8;A0A2R8Y756;A0A2R8YGL0 | |
| 3.5468 | 1.8684 | *10.1666* | 5.1939 | 4.3875 | 0.1768 | O95248 | SBF1 |
| -1.3322 | -0.5227 | *-3.3950* | -1.7500 | 1.4810 | 0.1773 | F2Z2X0;O00746;Q4TT34;A0A087WVT9;A2IDC9;H0Y6J0 | NME4 |
| -0.2460 | -1.1660 | *-0.3610* | -0.5910 | 0.5013 | 0.1779 | P20020;E7ERY9;F8W1V5;A0A0U1RQU3;H0YHH6 | ATP2B1 |
| 0.7602 | 1.1422 | *0.0526* | 0.6516 | 0.5528 | 0.1780 | Q9Y6R4;F5H4R1;F5H538;J3KNB8 | MAP3K4 |
| 0.6488 | 0.9260 | *0.0359* | 0.5369 | 0.4555 | 0.1780 | P35579;Q5BKV1;B1AH99 | MYH9 |
| -0.5945 | -0.7100 | *-0.0140* | -0.4395 | 0.3730 | 0.1781 | A0A0D9SFE4;A0A0U1RQP1;Q05193;A0A0D9SFB1;A0A1B0GU67;A0A1B0GUX5;K7ENE7 | DNM1 |
| 0.5034 | 0.5739 | *0.0091* | 0.3621 | 0.3077 | 0.1784 | Q96RP9;C9IZ01;F8WAU4 | GFM1 |
| 0.0234 | 1.6217 | *1.4651* | 1.0367 | 0.8811 | 0.1784 | O60239 | SH3BP5 |
| -0.2781 | -0.5496 | *-0.0519* | -0.2932 | 0.2492 | 0.1784 | P39019;M0QXK4;M0R140;M0QYF7;M0R2L9;A0A075B6E2 | RPS19 |
| -0.1864 | -1.3595 | *-2.4507* | -1.3322 | 1.1324 | 0.1785 | P23368;A0A1W2PPH1;A0A1W2PQT3;A0A1W2PQY8;A0A1W2PR68;A0A1W2PQH3;A0A1W2PQF8;A0A1W2PQ37 | ME2 |
| -4.0860 | -0.4295 | *-1.9562* | -2.1572 | 1.8365 | 0.1789 | Q9H3H9 | TCEAL2 |
| 1.3842 | 0.1996 | *0.5565* | 0.7134 | 0.6077 | 0.1790 | Q96NY7 | CLIC6 |
| -0.4617 | -0.7050 | *-2.2557* | -1.1408 | 0.9732 | 0.1794 | Q13439;H0Y6I0;E7EVX2;C9JHJ5;A0A087WTW2;M0R0N0;O95182;Q86W71 | GOLGA4 |
| -0.0337 | -1.3484 | *-1.6918* | -1.0247 | 0.8752 | 0.1798 | O15417;H9KVB4 | TNRC18 |
| -0.5669 | -0.0505 | *-0.2884* | -0.3020 | 0.2585 | 0.1803 | P31150;G5E9U5 | GDI1 |
| -0.0553 | -0.3377 | *-0.6505* | -0.3478 | 0.2977 | 0.1804 | Q15555;K7EL66;K7ERD8;K7ENB3;M0QX52 | MAPRE2 |
| 0.1971 | 2.7545 | *1.5283* | 1.4933 | 1.2791 | 0.1805 | Q15700;B7Z264;B7Z2T4;E9PN83;E9PIW2;B5MCC5;F8W750;A8MVA8;E9PPV7;E9PQT9;E9PRL2;C9 | DLG2 |

| L100/ CTRL1 | L100/ CTRL2 | L100/ CTRL3 | Mean | SD | T-test p-value | Accession | Gene Symbol |
|---|---|---|---|---|---|---|---|
| | | | | | | JFF9;H7C325;Q5JUW8;Q92796 | |
| -0.0391 | -0.7827 | *-1.1342* | -0.6520 | 0.5591 | 0.1808 | Q08J23 | NSUN2 |
| -0.0096 | -0.2284 | *-0.3216* | -0.1865 | 0.1602 | 0.1812 | Q6QNY1;A0A087X1N6;J3QRU7 | BLOC1S2 |
| 0.1645 | 1.4262 | *0.6355* | 0.7421 | 0.6376 | 0.1814 | A0A2R8Y4H4;Q6ZRR7;H3BUS4 | LRRC9 |
| -2.3730 | -0.2222 | *-1.1684* | -1.2545 | 1.0780 | 0.1814 | Q8IV08;M0R3G9 | PLD3 |
| 0.3296 | 0.0576 | *0.6528* | 0.3467 | 0.2979 | 0.1814 | E7ET52;Q8WW38;E5RJX0 | ZFPM2 |
| -3.2205 | -3.3684 | *-13.0486* | -6.5459 | 5.6321 | 0.1817 | P10074;Q5SY20;Q5SY21 | ZBTB48 |
| 0.8748 | 0.2543 | *2.0517* | 1.0603 | 0.9129 | 0.1819 | Q15418;E9PGT3;E9PMM7;Q5SVM7 | RPS6KA1 |
| -0.8282 | -0.7763 | *-3.1926* | -1.5990 | 1.3803 | 0.1826 | Q5VT06;H0Y6Q4;H0Y7F7;E9PIK0;Q30KQ4;Q86VL8;Q9Y2L8 | CEP350 |
| 3.7426 | 0.8679 | *1.0110* | 1.8738 | 1.6200 | 0.1830 | O15015;C9J3L0 | ZNF646 |
| -1.3812 | -0.3255 | *-3.0443* | -1.5836 | 1.3706 | 0.1833 | Q9UHJ3 | SFMBT1 |
| -0.2036 | -1.5049 | *-0.6001* | -0.7695 | 0.6670 | 0.1837 | Q86YA3;G3XAL8;D6REN9;D6RB47;D6REQ7 | ZGRF1 |
| -0.5102 | -0.5122 | *0.0004* | -0.3406 | 0.2954 | 0.1838 | Q58FF8 | HSP90AB2P |
| 0.1864 | 0.6643 | *1.5594* | 0.8034 | 0.6970 | 0.1840 | Q96K21;H3BRM1 | ZFYVE19 |
| -0.4163 | -2.1366 | *-0.6630* | -1.0720 | 0.9302 | 0.1841 | Q86UQ4;A0A0A0MT16;F5H7B7;H7C0U5 | ABCA13 |
| -0.8185 | -0.0305 | *-0.5283* | -0.4591 | 0.3985 | 0.1842 | Q9BRX8 | PRXL2A |
| -0.1829 | -3.1929 | *-1.8319* | -1.7359 | 1.5073 | 0.1842 | P28340;M0R2B7;M0QZR8;A0A2R8Y7K6;A0A2R8Y705;M0QZB4 | POLD1 |
| 1.1811 | 0.5925 | *3.4618* | 1.7451 | 1.5156 | 0.1843 | O00423;F8W717;G3V3N9;G3V500 | EML1 |
| 0.4937 | 0.0190 | *0.3133* | 0.2754 | 0.2396 | 0.1848 | A0A2R8Y7K4;F5GZB4;Q8IVV2;J3KRE7;J3QKX9 | LOXHD1 |
| -1.6378 | -0.1427 | *-0.8021* | -0.8609 | 0.7493 | 0.1849 | H3BMM9;H3BTC0;Q15287;H3BMS0;H3BV80;H3BPG5 | RNPS1 |
| -2.6292 | -0.0017 | *-2.2787* | -1.6366 | 1.4266 | 0.1853 | H3BQ65;Q3V5L5;H3BR20 | MGAT5B |
| -0.4134 | -0.1544 | *-0.0606* | -0.2095 | 0.1827 | 0.1854 | P49756 | RBM25 |
| -0.3331 | -0.6938 | *-2.0170* | -1.0146 | 0.8866 | 0.1860 | Q4VXU2 | PABPC1L |
| -0.0629 | -0.2211 | *-0.0468* | -0.1103 | 0.0963 | 0.1860 | Q13557;D6R938;E9PF82;E9PBG7;H0Y9C2;E7EQE4;E9PBE8;E7ETC9;D6RHX9;H0Y9J2 | CAMK2D |

| L100/ CTRL1 | L100/ CTRL2 | L100/ CTRL3 | Mean | SD | T-test p-value | Accession | Gene Symbol |
|---|---|---|---|---|---|---|---|
| 0.3829 | 1.6539 | *0.4328* | 0.8232 | 0.7198 | 0.1861 | P38646;D6RJI2;D6RA73;H0YBG6;H0Y8S0 | HSPA9 |
| 0.1125 | 2.1865 | *3.3936* | 1.8975 | 1.6595 | 0.1862 | E7ESC5;F5GZA4;Q96AQ1;Q96LY2 | CCDC74B |
| -2.6056 | -0.1151 | *-1.5609* | -1.4272 | 1.2506 | 0.1867 | H0YKD8;P46779 | RPL28 |
| -0.3982 | -0.3126 | *-1.4420* | -0.7176 | 0.6288 | 0.1867 | P25325;B1AH49 | MPST |
| 0.2799 | 2.8808 | *1.3207* | 1.4938 | 1.3091 | 0.1868 | Q8NFA0;K7EQL6;K7EL53;K7EKZ1;K7EN13 | USP32 |
| -0.2734 | -0.4689 | *-1.4862* | -0.7428 | 0.6512 | 0.1868 | P22090;C9JEH7 | RPS4Y1 |
| 0.0834 | 1.0249 | *1.7508* | 0.9530 | 0.8360 | 0.1870 | P49454;A0A087X1R9;E9PL74 | CENPF |
| 1.1691 | 0.0525 | *1.8010* | 1.0075 | 0.8854 | 0.1875 | Q9NR45;Q5TBR1;Q5TBR0 | NANS |
| -0.0524 | -1.0074 | *-0.5721* | -0.5439 | 0.4781 | 0.1876 | P08621;M0QYR1 | SNRNP70 |
| -0.2815 | -0.5371 | *-1.6384* | -0.8190 | 0.7210 | 0.1880 | F5GX23;F5H5V4;J3KN29;O00233;F5H169;F5H7X1 | PSMD9 |
| -0.4002 | -1.1063 | *-0.1633* | -0.5566 | 0.4906 | 0.1883 | Q9HAV4;H0Y3W3;H0Y3Q8 | XPO5 |
| -0.8094 | -3.5599 | *-0.9249* | -1.7647 | 1.5557 | 0.1884 | Q05086;A0A1B0GVL3;A0A0D9SG77;A0A0D9SG63;S4R306;A0A1B0GTB3 | UBE3A |
| -1.0429 | -0.5523 | *-0.0662* | -0.5538 | 0.4883 | 0.1885 | P16157 | ANK1 |
| -0.5742 | -0.0186 | *-0.8564* | -0.4831 | 0.4263 | 0.1887 | Q92973;S4R398 | TNPO1 |
| 0.7229 | 1.0105 | *3.5530* | 1.7622 | 1.5576 | 0.1891 | Q86WR0;B7Z2L8;G3V121 | CCDC25 |
| -0.0503 | -0.6048 | *-1.0594* | -0.5715 | 0.5054 | 0.1892 | B5BUE6;J3KTA4;P17844;J3KRZ1;J3QSF1;J3QRQ7;J3QLG9;J3QR62;J3QRN5 | DDX5 |
| -0.5221 | -0.3215 | *-1.7189* | -0.8542 | 0.7556 | 0.1893 | O15212 | PFDN6 |
| -0.4754 | -0.3297 | *-0.0073* | -0.2708 | 0.2396 | 0.1893 | O94856;X6RKN2;H7BY57;D6RBU5;D6RHX4;H7C0L6;A0A0C4DG92;H7C073;H7C5S4 | NFASC |
| 0.6852 | 1.1231 | *0.0397* | 0.6160 | 0.5450 | 0.1894 | P39687;H0YN26;H7BZ09;O43423 | ANP32A |
| -0.1588 | -1.9554 | *-3.4243* | -1.8462 | 1.6355 | 0.1897 | Q8WYL5;F8VS18 | SSH1 |
| 0.9525 | 0.6379 | *0.0184* | 0.5363 | 0.4752 | 0.1898 | Q5JRA6;A0A0A0MRH6 | MIA3 |
| -0.5754 | -0.9810 | *-0.0394* | -0.5319 | 0.4723 | 0.1904 | Q99798;A2A274 | ACO2 |
| -0.1162 | -0.0068 | *-0.0617* | -0.0616 | 0.0547 | 0.1905 | Q14679;E7EX20;E9PH58;H7BZY4;H7C2S3;H7C421 | TTLL4 |

| L100/ CTRL1 | L100/ CTRL2 | L100/ CTRL3 | Mean | SD | T-test p-value | Accession | Gene Symbol |
|---|---|---|---|---|---|---|---|
| -1.2938 | -0.0043 | *-1.7283* | -1.0088 | 0.8967 | 0.1907 | O95865;A0A140T971;Q5SRR8;Q5SSV3;H0Y7N1 | DDAH2 |
| -0.2294 | -5.5474 | *-3.2169* | -2.9979 | 2.6658 | 0.1908 | Q96F86;H3BPW9;H3BQ37;H3BMB8;H3BNJ7;H3BQA1;H3BQP5;H3BTF8;H3BTH0;H3BU87 | EDC3 |
| -0.0090 | 0.9375 | *1.1397* | 0.6894 | 0.6132 | 0.1909 | A0A2R8YFA9;O94851 | MICAL2 |
| -1.2819 | -1.1307 | *-5.0401* | -2.4842 | 2.2148 | 0.1915 | A0A087WU27;A0A087WUS4;A6NKB8;C9JMZ3;Q9H4A4 | RNPEP |
| -8.2270 | -1.4106 | *-2.6084* | -4.0820 | 3.6393 | 0.1915 | Q8TAE8 | GADD45GIP1 |
| 1.1760 | 0.1078 | *0.5271* | 0.6036 | 0.5382 | 0.1915 | Q92824 | PCSK5 |
| 1.5221 | 1.3569 | *-0.0236* | 0.9518 | 0.8487 | 0.1916 | Q9H1K0 | RBSN |
| -0.1679 | -0.8474 | *-0.2401* | -0.4185 | 0.3732 | 0.1916 | Q13618;H7C399 | CUL3 |
| -0.0308 | -0.1228 | *-0.2923* | -0.1486 | 0.1326 | 0.1918 | P15311;E7EQR4;E9PNP4;H0YHN3 | EZR |
| -0.1542 | -1.0900 | *-2.2065* | -1.1502 | 1.0275 | 0.1921 | P29597;E9PPF2;E9PQL2 | TYK2 |
| -0.6243 | -3.7183 | *-1.1892* | -1.8439 | 1.6477 | 0.1922 | M0QXH3 | CPAMD8 |
| -0.1689 | -1.4856 | *-2.8577* | -1.5041 | 1.3445 | 0.1923 | P37268;A0A1W2PQ47;E9PNM1;E9PJG4;E9PS69 | FDFT1 |
| 0.0792 | 2.5336 | *1.5224* | 1.3784 | 1.2335 | 0.1926 | Q14315 | FLNC |
| -0.3666 | -0.1207 | *-0.9615* | -0.4830 | 0.4323 | 0.1926 | Q96CN7;D6RGE2 | ISOC1 |
| -0.7033 | -0.0763 | *-1.3442* | -0.7079 | 0.6339 | 0.1928 | Q9ULI0;H7BYF1;C9JG15 | ATAD2B |
| 0.4331 | 3.3008 | *1.2262* | 1.6534 | 1.4808 | 0.1928 | O75891;C9IZ36;C9JY00;C9JYZ6;D6RFJ7;F2Z324;F8WC34 | ALDH1L1 |
| -1.7991 | -0.1438 | *-0.8389* | -0.9273 | 0.8311 | 0.1930 | Q9P2D7 | DNAH1 |
| 0.0427 | -1.8888 | *-1.8181* | -1.2214 | 1.0953 | 0.1932 | X6RHV1 | SETDB1 |
| 0.2409 | 0.5447 | *1.6016* | 0.7957 | 0.7142 | 0.1934 | O14828 | SCAMP3 |
| 0.1695 | 0.4500 | *0.0568* | 0.2254 | 0.2025 | 0.1936 | P12268;H0Y4R1;E7ETK5 | IMPDH2 |
| -0.5112 | -0.2817 | *-1.6479* | -0.8136 | 0.7316 | 0.1939 | P13645;C4AM86;F8VZY9;K7EPJ9;O76014;O76015;Q14532;Q92764 | KRT10 |
| 0.0282 | -2.1487 | *-1.7280* | -1.2829 | 1.1547 | 0.1942 | P78332;E9PGM9 | RBM6 |
| 0.8116 | 1.0542 | *-0.0098* | 0.6187 | 0.5576 | 0.1946 | A0AUZ9 | KANSL1L |
| 0.2780 | 0.6517 | *1.9043* | 0.9447 | 0.8518 | 0.1947 | P13611;E9PF17;D6RGZ6;Q86W61 | VCAN |
| 0.3450 | 0.8979 | *2.5229* | 1.2553 | 1.1321 | 0.1948 | Q9C040;A0A0J9YX34;C9J084 | TRIM2 |

| L100/ CTRL1 | L100/ CTRL2 | L100/ CTRL3 | Mean | SD | T-test p-value | Accession | Gene Symbol |
|---|---|---|---|---|---|---|---|
| -0.7588 | -2.2422 | *-0.3329* | -1.1113 | 1.0023 | 0.1948 | Q5T1H1 | EYS |
| -3.4446 | -0.0446 | *-5.1461* | -2.8784 | 2.5975 | 0.1949 | P42892;B4DKB2 | ECE1 |
| -0.2528 | -0.8370 | *-2.1618* | -1.0839 | 0.9781 | 0.1949 | Q8N5M1;C9J2Q2;K7ELC1 | ATPAF2 |
| 1.0676 | 0.0782 | *0.5028* | 0.5495 | 0.4963 | 0.1952 | Q9NV72;M0R085 | ZNF701 |
| 0.3326 | 0.5659 | *0.0166* | 0.3051 | 0.2757 | 0.1954 | P53602;H3BP35; H3BQ47 | MVD |
| 0.0855 | 0.3165 | *0.7903* | 0.3974 | 0.3593 | 0.1954 | Q6ZVL6;H0YDE5 | KIAA1549L |
| -0.6697 | -0.5938 | *0.0161* | -0.4158 | 0.3759 | 0.1955 | O75947;F5H608 | ATP5PD |
| 4.9405 | 1.7580 | *0.6658* | 2.4548 | 2.2209 | 0.1957 | Q13029 | PRDM2 |
| 0.0251 | 0.9171 | *0.5423* | 0.4948 | 0.4479 | 0.1958 | A0A0C4DGH6;A0JNW5;F8VW65; F8W1I2;F8W665 | UHRF1BP1L |
| 11.7687 | 13.9685 | *-0.2938* | 8.4811 | 7.6785 | 0.1959 | P23284 | PPIB |
| -0.0582 | -0.7908 | *-0.3684* | -0.4058 | 0.3677 | 0.1961 | P00492 | HPRT1 |
| 0.2717 | -0.0001 | *0.3902* | 0.2206 | 0.2001 | 0.1964 | Q99447;I3L1R7;I3L3V9;I3L1L9;I3L2Q1;I3L1C4 | PCYT2 |
| -1.3363 | -4.0850 | *-0.6226* | -2.0146 | 1.8282 | 0.1965 | Q6P2E9 | EDC4 |
| -0.2455 | -0.4183 | *-1.4062* | -0.6900 | 0.6262 | 0.1966 | O60462 | NRP2 |
| 0.6470 | 0.1455 | *1.5381* | 0.7769 | 0.7053 | 0.1967 | Q8TBB0 | THAP6 |
| 0.4319 | -0.0093 | *0.3569* | 0.2598 | 0.2361 | 0.1968 | P22392;Q32Q12; J3KPD9;O60361; E7ERL0;F6XY72; C9K028;E5RHP0 | NME2 |
| 0.9261 | 0.5441 | *0.0241* | 0.4981 | 0.4528 | 0.1970 | O14525 | ASTN1 |
| 0.0028 | 0.9173 | *0.6187* | 0.5129 | 0.4663 | 0.1971 | Q8N8E3;F5GYE8;J3QSA5;J3KSN4 | CEP112 |
| 1.1172 | 0.0980 | *2.1246* | 1.1133 | 1.0133 | 0.1974 | P50440;H0YKW9 | GATM |
| -0.3503 | -0.4643 | *-1.7593* | -0.8579 | 0.7826 | 0.1980 | Q9UFE4 | CCDC39 |
| -0.1135 | -1.0947 | *-0.4352* | -0.5478 | 0.5002 | 0.1983 | Q92545;F6S8H2 | TMEM131 |
| -0.0138 | 0.8932 | *1.2216* | 0.7003 | 0.6398 | 0.1985 | Q9H999;E5RHA5 | PANK3 |
| -0.2175 | -0.0006 | *-0.3265* | -0.1816 | 0.1659 | 0.1985 | P49915 | GMPS |
| -0.3085 | -0.5321 | *-0.0132* | -0.2846 | 0.2602 | 0.1987 | P0CG39 | POTEJ |
| -2.0906 | -0.9125 | *-0.1712* | -1.0581 | 0.9679 | 0.1988 | P06753;Q5HYB6; Q5VU61;D6RFM2 | TPM3 |
| 0.4496 | 1.7089 | *0.3375* | 0.8320 | 0.7615 | 0.1990 | O75376;A0A088AWL3;E7EVU5;C9JAP0;E7EVK1;E7EW50;J3KS51 | NCOR1 |
| -0.3983 | -0.1263 | *-0.0610* | -0.1952 | 0.1789 | 0.1993 | P18615;A0A0A0MT02;A0A0A0MSN9;E9PD43;A0A0G2JI50;H7C1J7 | NELFE |
| -0.4154 | 0.0139 | *-0.5065* | -0.3027 | 0.2779 | 0.1999 | O00231;J3QRY4; J3QS13 | PSMD11 |
| -0.3213 | -0.2076 | *-0.0012* | -0.1767 | 0.1622 | 0.1999 | P04899 | GNAI2 |
| 2.2937 | 3.3181 | *-0.0249* | 1.8623 | 1.7127 | 0.2004 | P53779;A0A1P0B7D2;A0A286YE | MAPK10 |

| L100/ CTRL1 | L100/ CTRL2 | L100/ CTRL3 | Mean | SD | T-test p-value | Accession | Gene Symbol |
|---|---|---|---|---|---|---|---|
| | | | | | | W9;A0A286YEX7;A0A286YF62;A0A286YF85;A0A286YF97;A0A286YFC0;A0A286YFJ6;A8MWW6;A0A286YEN5;A0A286YEQ0;A0A286YEQ7;A0A286YES9;A0A286YEV3;A0A286YEY0;A0A286YF19;A0A286YF35;A0A286YFA6;A0A286YFB6;A0A286YFD7;A0A286YFI3;A0A286YFJ4;A0A286YFM6;A0A286YFN2;D6RJF9;A0A286YES0;A0A286YEV8;A0A286YF02;Q499Y8;A0A286YF83;D6RCB1;D6RDG1;A0A286YFE2;A8MPV1;D6RAJ0;A0A1W2PPW1;A0A1W2PQ03;A0A1W2PRZ2;A0A286YES8;A0A286YEV5;A0A286YF53;A0A286YF80;A0A286YF95;A0A286YFA3;A0A286YFA7;A0A286YFB7;A0A286YFC3;A0A286YFK3;D6R9C1;D6RAU3;D6RBH2;D6RFX8;D6RFZ7 | |
| -0.1580 | -0.3353 | *-1.0535* | -0.5156 | 0.4742 | 0.2004 | P07737;K7EJ44;I3L3D5 | PFN1 |
| -0.6343 | -0.9134 | *-3.3788* | -1.6422 | 1.5104 | 0.2004 | O75038;B9DI82 | PLCH2 |
| -0.0392 | -1.0740 | *-0.5748* | -0.5626 | 0.5175 | 0.2004 | O94906 | PRPF6 |
| -0.0991 | -1.3401 | *-2.5379* | -1.3257 | 1.2195 | 0.2004 | Q00526 | CDK3 |
| -0.2579 | -0.6432 | *-0.0625* | -0.3212 | 0.2955 | 0.2004 | O75373;M0R1D1 | ZNF737 |
| -0.0455 | -0.3650 | *-0.7673* | -0.3926 | 0.3617 | 0.2009 | Q99700;A0A2R8Y5A6;F8VQP2;F8WB06;V9GY86;A0A2R8Y7P6;A0A2R8YDM9;H0YH87;F8VRK6 | ATXN2 |
| -0.0426 | -0.7437 | *-1.3651* | -0.7171 | 0.6616 | 0.2013 | P35251 | RFC1 |
| -0.0544 | 1.3473 | *1.5844* | 0.9591 | 0.8857 | 0.2015 | Q15858;A0A2R8YDP4 | SCN9A |
| 0.0148 | -0.5487 | *-0.4323* | -0.3221 | 0.2975 | 0.2016 | C9J813 | CALD1 |

| L100/ CTRL1 | L100/ CTRL2 | L100/ CTRL3 | Mean | SD | T-test p-value | Accession | Gene Symbol |
|---|---|---|---|---|---|---|---|
| -0.0298 | -0.6126 | *-1.1154* | -0.5859 | 0.5433 | 0.2027 | P18074;E7EVE9; A8MX75;K7EKF3 | ERCC2 |
| 0.2207 | 0.4247 | *1.4076* | 0.6843 | 0.6346 | 0.2028 | P49721;A0A087 WVV1 | PSMB2 |
| 0.1302 | 0.2683 | *0.8659* | 0.4215 | 0.3911 | 0.2029 | O14531;Q5T0Q6 | DPYSL4 |
| -0.7403 | -1.0324 | *0.0187* | -0.5847 | 0.5425 | 0.2029 | Q9NQ29;A8MYV 2;B8ZZ10;B8ZZ0 9 | LUC7L |
| -0.9832 | -0.3367 | *-2.7897* | -1.3699 | 1.2714 | 0.2030 | Q9Y4J8;M0QZ28 ;M0R0C4 | DTNA |
| -0.9561 | -0.9000 | *-4.1531* | -2.0031 | 1.8622 | 0.2035 | M0R366;Q9BTV5 | FSD1 |
| -0.0457 | 1.3201 | *1.0753* | 0.7832 | 0.7282 | 0.2035 | O15078;J3KNF5; A0A0A0MS86;F8 VS29;S4R322;F8 W097;F8W0V9;A 0A087WUX4 | CEP290 |
| -0.2682 | -0.4029 | *-1.4918* | -0.7210 | 0.6710 | 0.2038 | P14927;B7Z2R2; E5RHG9 | UQCRB |
| -0.3688 | -0.1243 | *-1.0464* | -0.5132 | 0.4777 | 0.2039 | O14715;J3KQ37; F8W705;C9J1W9 ;C9J6W1;C9JF75 ;C9J1P2 | RGPD8 |
| 3.2022 | 0.2372 | *1.3798* | 1.6064 | 1.4954 | 0.2039 | F5H829;Q96EP1 | CHFR |
| -0.0126 | 0.2548 | *0.2662* | 0.1695 | 0.1578 | 0.2039 | P11498;E9PS68 | PC |
| -0.4435 | -0.2812 | *-1.6103* | -0.7783 | 0.7250 | 0.2041 | P49591;Q5T5C7 | SARS |
| -0.0448 | 1.0593 | *1.3336* | 0.7827 | 0.7296 | 0.2043 | O14578;H0YGG8 | CIT |
| -1.8690 | -0.2688 | *-0.5873* | -0.9083 | 0.8470 | 0.2044 | Q14019 | COTL1 |
| -0.0069 | 0.8941 | *0.5854* | 0.4909 | 0.4579 | 0.2044 | Q07812;K4JQN1 | BAX |
| -0.7005 | -0.6803 | *0.0347* | -0.4487 | 0.4188 | 0.2046 | Q9Y3I0 | RTCB |
| 0.0648 | -1.4782 | *-1.2928* | -0.9021 | 0.8424 | 0.2048 | H3BPZ4 | SH2B1 |
| -0.6254 | -0.8269 | *0.0243* | -0.4760 | 0.4448 | 0.2050 | P35080;C9J712; C9J2N0;C9JQ45; G5E9Q6;C9J0J7 | PFN2 |
| -0.6923 | -0.3753 | *-2.3682* | -1.1453 | 1.0709 | 0.2052 | Q9UH77;D6RH2 1;D6R9K4 | KLHL3 |
| -1.5485 | -0.7161 | *-4.9766* | -2.4138 | 2.2582 | 0.2053 | P23468;F5GWR7 ;Q3KPI9;C9J8S8; C9J6E4 | PTPRD |
| 0.0499 | 0.3642 | *0.8093* | 0.4078 | 0.3816 | 0.2054 | Q15393;I3L4G7; H3BMB0;J3QL37 ;J3QRB2 | SF3B3 |
| 1.7444 | 0.2394 | *0.5577* | 0.8472 | 0.7932 | 0.2056 | Q96EY1;I3L1I6;I3 L1T6;I3L1T9;I3L3 D3 | DNAJA3 |
| 0.2611 | 0.1683 | *-0.0021* | 0.1425 | 0.1335 | 0.2058 | Q8N3C0;E5RFZ0 | ASCC3 |
| -2.3377 | -0.3887 | *-5.4582* | -2.7282 | 2.5572 | 0.2059 | Q96I24 | FUBP3 |
| 1.5840 | 1.3457 | *-0.0708* | 0.9530 | 0.8946 | 0.2063 | H0YI59 | WDR90 |
| 2.5156 | 0.4824 | *0.6251* | 1.2077 | 1.1349 | 0.2066 | P19174;A0A0D9 SEK2;V9GY71;V 9GYH5 | PLCG1 |
| -0.3236 | -0.4355 | *0.0133* | -0.2486 | 0.2336 | 0.2067 | P31930 | UQCRC1 |
| 1.7235 | -0.0821 | *2.2378* | 1.2930 | 1.2184 | 0.2074 | Q14676;A2AB05; A2AB07 | MDC1 |

| L100/ CTRL1 | L100/ CTRL2 | L100/ CTRL3 | Mean | SD | T-test p-value | Accession | Gene Symbol |
|---|---|---|---|---|---|---|---|
| -0.4804 | -0.1413 | *-1.3394* | -0.6537 | 0.6176 | 0.2082 | Q12765;C9K052; C9J7U9;B8ZZP4 | SCRN1 |
| -0.7126 | -0.0156 | *-0.3773* | -0.3685 | 0.3486 | 0.2086 | Q15652 | JMJD1C |
| 0.1277 | -2.0712 | *-2.1749* | -1.3728 | 1.3005 | 0.2090 | P61201;B4DIH5 | COPS2 |
| -1.1954 | -1.2867 | *0.0749* | -0.8024 | 0.7611 | 0.2094 | P61289;K9J957; K7ENH2;K7ESG 5;B3KQ25;A0A08 7WTV2;K7EKR3; K7EPX6 | PSME3 |
| 2.9618 | 2.9920 | *-0.1868* | 1.9223 | 1.8266 | 0.2099 | Q86UE8;J3KRK0 ;J3QLK5;J3KST4 ;J3QS73 | TLK2 |
| 0.1200 | 1.0209 | *0.3390* | 0.4933 | 0.4698 | 0.2106 | Q05397;E7ESA6; H0YBP1;H0YB16 ;H0YBZ1;E9PEI4 ;B4DWJ1;E5RHD 8;E5RI03;E5RG8 0;E5RJQ2 | PTK2 |
| 0.0128 | -0.3501 | *-0.2550* | -0.1974 | 0.1881 | 0.2107 | Q8N1W1 | ARHGEF28 |
| -0.3612 | -1.2858 | *-0.1985* | -0.6152 | 0.5865 | 0.2109 | P54802 | NAGLU |
| 2.0589 | 0.3393 | *0.5400* | 0.9794 | 0.9402 | 0.2130 | O00584;A0A087 WZM2;D6RHI9;A 0A087WWI1;J3Q Q64 | RNASET2 |
| 0.2576 | 2.4405 | *0.8349* | 1.1777 | 1.1311 | 0.2131 | Q8NFF5 | FLAD1 |
| 0.0431 | -0.9359 | *-0.6991* | -0.5306 | 0.5107 | 0.2138 | O15054;I3L0Z0 | KDM6B |
| 0.0846 | 0.5829 | *1.3976* | 0.6884 | 0.6628 | 0.2139 | A0A0G2JRX7;A0 A0G2JRX8;B5M CY4;B5MD04;E7 EMH2;E7EW99; H7C2P3;P78395 | PRAME |
| 0.2216 | 0.1616 | *0.9096* | 0.4310 | 0.4156 | 0.2144 | Q9NTZ6 | RBM12 |
| 0.0595 | 1.5206 | *3.0125* | 1.5309 | 1.4765 | 0.2144 | O95372;Q5QPQ0 ;Q5QPN9;Q5QP Q1;Q5QPQ2;Q5 QPQ3;Q5QPN5 | LYPLA2 |
| 0.0557 | -0.7445 | *-0.8646* | -0.5178 | 0.5003 | 0.2149 | Q29RF7;H0Y9X6 | PDS5A |
| -0.0024 | 0.7574 | *1.3506* | 0.7018 | 0.6782 | 0.2149 | P43003;A0A087X 0U3;A0A087WT8 7 | SLC1A3 |
| -0.1662 | 2.3059 | *2.1962* | 1.4453 | 1.3966 | 0.2149 | Q58FF3 | HSP90B2P |
| -0.2948 | -0.0803 | *-0.8392* | -0.4048 | 0.3912 | 0.2150 | P55209;F5H4R6; F8VY35;F8W0J6; H0YIV4;F8VUX1; F8VV59;F8W020; B7Z9C2;F8VXI6; F8W118;F8W543 ;H0YH88;H0YHC 3;F8VRJ2;B7Z4K 9;F8VVB5 | NAP1L1 |
| 1.0198 | 1.2008 | *-0.0775* | 0.7144 | 0.6918 | 0.2156 | A6NEC2 | NPEPPSL1 |
| 0.0985 | -1.2988 | *-1.5489* | -0.9164 | 0.8878 | 0.2157 | Q9BRJ6;C9JQV0 | C7orf50 |
| -0.0299 | -5.1839 | *-2.7649* | -2.6596 | 2.5786 | 0.2159 | P11047;R4GNC7 | LAMC1 |
| 0.0840 | 1.0727 | *2.3430* | 1.1666 | 1.1325 | 0.2163 | D6RDN9;Q15119 | PDK2 |

| L100/ CTRL1 | L100/ CTRL2 | L100/ CTRL3 | Mean | SD | T-test p-value | Accession | Gene Symbol |
|---|---|---|---|---|---|---|---|
| 0.3874 | 1.1449 | *3.5253* | 1.6859 | 1.6374 | 0.2165 | O75175;B7Z6J7; H7C148;A0A0G2 JNJ7;A0A2R8Y4 W0;A0A2R8Y691 ;A0A2R8Y7Z8;H0 Y5X7 | CNOT3 |
| -2.9210 | -0.0043 | *-1.5836* | -1.5030 | 1.4601 | 0.2165 | A0A075B7G4;Q8 IYB9 | ZNF595 |
| -0.1115 | -0.0571 | *-0.3998* | -0.1895 | 0.1841 | 0.2167 | P23526 | AHCY |
| -0.0504 | 0.6243 | *0.6487* | 0.4075 | 0.3968 | 0.2172 | A0A1W2PPA4;A0 A1X7SBU6;A0A1 W2PP32;A0A1W 2PRR5;A0A1W2 PRC7;A0A1W2P NS5;A0A1W2PR 84 | ADGRV1 |
| -0.2630 | 0.0174 | *-0.2192* | -0.1549 | 0.1509 | 0.2173 | Q92598;A0A0A0 MSM0;R4GN69; Q5TBM3 | HSPH1 |
| -0.2082 | -0.4023 | *-0.0036* | -0.2047 | 0.1994 | 0.2173 | O60701;E7ER83; E7ETF4;E7EV97; D6RHF4;E7ER95 ;E9PBD2 | UGDH |
| -0.6901 | -0.8940 | *0.0517* | -0.5108 | 0.4977 | 0.2174 | A4UGR9;A0A087 WVH1;Q9H4E7 | XIRP2 |
| 1.3904 | 3.1787 | *0.1347* | 1.5679 | 1.5297 | 0.2178 | P27482 | CALML3 |
| 1.4690 | -0.1211 | *1.7023* | 1.0167 | 0.9923 | 0.2179 | Q05932 | FPGS |
| 0.7307 | 0.0837 | *1.7253* | 0.8466 | 0.8269 | 0.2182 | Q2M2I5 | KRT24 |
| 0.2876 | 1.1269 | *0.1787* | 0.5311 | 0.5189 | 0.2183 | Q9H0W8;M0QZC 7;M0R2N0;M0R0 U0 | SMG9 |
| 2.8213 | 4.3139 | *-0.1591* | 2.3254 | 2.2774 | 0.2190 | Q5JRD6 | PDSS2 |
| 0.1635 | 0.7533 | *2.0658* | 0.9942 | 0.9737 | 0.2190 | O75054 | IGSF3 |
| -0.5843 | -0.8339 | *-3.4608* | -1.6263 | 1.5936 | 0.2192 | O43719;Q5H918; Q5H919 | HTATSF1 |
| -2.8062 | 0.2395 | *-2.9062* | -1.8243 | 1.7880 | 0.2192 | P55286;J3KRI5;X 6R3Y6;J3QKW5; J3QLE6;J3KTG8 | CDH8 |
| -1.9844 | -1.1785 | *-7.6778* | -3.6136 | 3.5427 | 0.2193 | Q14289;C9JHV9; E5RJ77;E5RHL2; E5RK84 | PTK2B |
| -0.4436 | -0.1479 | *-1.3852* | -0.6589 | 0.6461 | 0.2194 | Q9P2B2 | PTGFRN |
| -0.3830 | -0.4514 | *0.0332* | -0.2671 | 0.2622 | 0.2198 | A6NFX8;A6NJU6 ;Q9UKK9;A6NCQ 0;H0YEY4;C9JY Y9 | NUDT5 |
| 0.0144 | -0.2429 | *-0.3726* | -0.2004 | 0.1970 | 0.2201 | Q9C0B1;A0A1B0 GTC3;A0A1B0G TC5;A0A1B0GU2 6;A0A1B0GUC3; A0A1B0GV98;A0 A1B0GVH5;A0A1 B0GTY1;A0A1B0 GTZ8;A0A1B0G UY7;X6R3I0;A0A 1B0GTI3 | FTO |

| L100/ CTRL1 | L100/ CTRL2 | L100/ CTRL3 | Mean | SD | T-test p-value | Accession | Gene Symbol |
|---|---|---|---|---|---|---|---|
| 0.3406 | -4.1310 | *-5.3342* | -3.0416 | 2.9901 | 0.2202 | P17213;A0A0D9SFX6;A0A2R8YDF1;H0Y738 | BPI |
| 1.3840 | 0.5568 | *0.0764* | 0.6724 | 0.6614 | 0.2203 | Q08722;A0A2R8Y484;H7BYS8 | CD47 |
| 0.0317 | -0.4327 | *-0.6128* | -0.3379 | 0.3325 | 0.2204 | Q6IQ26;H0YDE6 | DENND5A |
| -0.3211 | -0.1508 | *-0.0071* | -0.1597 | 0.1572 | 0.2205 | P07195;A8MW50;C9J7H8;F5H793 | LDHB |
| 0.0437 | -0.4792 | *-0.5386* | -0.3247 | 0.3204 | 0.2213 | Q2LD37 | KIAA1109 |
| 0.8586 | 1.1931 | *-0.0677* | 0.6613 | 0.6531 | 0.2216 | A0A0A0MRW5;E7ERH8;Q8NEE6 | FBXL13 |
| -1.2113 | -0.2446 | *-3.3247* | -1.5936 | 1.5752 | 0.2218 | Q9Y6M1;F8W930 | IGF2BP2 |
| 0.5774 | 0.7562 | *-0.0496* | 0.4280 | 0.4232 | 0.2219 | D6RAS9;O14827 | RASGRF2 |
| 0.6300 | 1.6593 | *0.1063* | 0.7985 | 0.7901 | 0.2221 | Q5T3Q7;Q9H583;Q6P664 | HEATR1 |
| -0.6653 | -0.0890 | *-1.6691* | -0.8078 | 0.7996 | 0.2223 | Q12797;E5RG29;E5RG56;E5RHJ2;E5RHK2;G3XAN5 | ASPH |
| -0.2161 | -1.7294 | *-4.2825* | -2.0760 | 2.0553 | 0.2223 | Q6PL18 | ATAD2 |
| 2.5160 | 0.4244 | *0.5855* | 1.1753 | 1.1639 | 0.2224 | P15428 | HPGD |
| -0.3163 | -0.7607 | *-2.6177* | -1.2316 | 1.2208 | 0.2227 | A2A288;H0Y4Y6 | ZC3H12D |
| -1.4914 | -0.3850 | *-0.2177* | -0.6980 | 0.6921 | 0.2228 | P48382;F8W689;A0A0A0MSM9;A0A0A0MSQ2;A0A0A0MT34;F6R6G4;F6S3S0;F6UE82;F6X9D6 | RFX5 |
| -0.7806 | -1.3024 | *-5.1747* | -2.4192 | 2.4005 | 0.2230 | J3KQ21;Q9P2S6;C9JZ56;H7C254;Q5CZB7;J3KPY5;Q6GPI0 | ANKMY1 |
| 1.2169 | -0.1087 | *1.1332* | 0.7471 | 0.7423 | 0.2234 | P51532;Q9HBD4;A0A0A0MT49;A0A2R8Y4P4;A0A2R8Y7S2;A0A2R8YGG3;A0A2R8Y440;A0A2R8YG32;A0A2R8Y7F3;A0A2R8Y6N0;A0A2R8Y526;A0A2R8YF58;A0A2R8Y7Y7;A0A2R8Y6V2;A0A2R8YF80;A0A2R8Y523;A0A2R8Y5K3;K7EQF0;A0A2R8YFK5;A0A2R8YGP5;K7EP28;A0A0U1RRF5;A0A2R8YCY3;A0A2R8Y4R6;A0A2R8YF38;A0A2R8YFV8 | SMARCA4 |
| 0.0517 | -0.5712 | *-0.5364* | -0.3520 | 0.3500 | 0.2237 | Q6ZS30 | NBEAL1 |

| L100/ CTRL1 | L100/ CTRL2 | L100/ CTRL3 | Mean | SD | T-test p-value | Accession | Gene Symbol |
|---|---|---|---|---|---|---|---|
| -0.9759 | -0.3405 | *-0.0768* | -0.4644 | 0.4622 | 0.2239 | P62424;Q5T8U2; Q5T8U3 | RPL7A |
| 3.8444 | 1.7583 | *0.0788* | 1.8938 | 1.8865 | 0.2242 | E7EVM7;Q9H5I5; A0A2R8Y3N5 | PIEZO2 |
| 1.3710 | 1.1044 | *-0.1065* | 0.7896 | 0.7874 | 0.2246 | Q6NXT6 | TAPT1 |
| 1.4226 | 1.2857 | *-0.1274* | 0.8603 | 0.8581 | 0.2246 | P18505;D6REM0 | GABRB1 |
| 1.8960 | 2.8823 | *-0.1417* | 1.5455 | 1.5422 | 0.2247 | A0A096LNH7;Q9Y5E3;Q9UN67;Q9Y5F1 | PCDHB6 |
| -0.2388 | 0.0238 | *-0.2776* | -0.1642 | 0.1639 | 0.2249 | P20648 | ATP4A |
| 0.2428 | 0.3671 | *1.5401* | 0.7167 | 0.7158 | 0.2251 | E9PCX7;D6RCR6;Q13423;D6RAI5;D6RHU2 | NNT |
| 0.3481 | 0.5043 | *-0.0292* | 0.2744 | 0.2743 | 0.2252 | Q9UJU6;B4DDD6;F8WBG8;F2Z2V3;F2Z3E3;F8WB73;F8WBB2;F8WC20;F8WCK3; F8WFE1 | DBNL |
| -1.4116 | -0.3570 | *-4.2148* | -1.9945 | 1.9939 | 0.2253 | A0AVI2;A0A286YFD1;A0A286YFJ1;A0A286YEQ6 | FER1L5 |
| 0.4957 | -0.0491 | *0.6179* | 0.3549 | 0.3551 | 0.2256 | Q9H2M9 | RAB3GAP2 |
| -0.6610 | -0.7812 | *0.0673* | -0.4583 | 0.4591 | 0.2259 | Q02878;F8VZ45; U3KQR5;F8VR69 | RPL6 |
| -0.0786 | -0.6595 | *-0.1849* | -0.3077 | 0.3093 | 0.2270 | P62277;J3KMX5; E9PS50 | RPS13 |
| 0.7362 | 1.9081 | *0.0945* | 0.9129 | 0.9196 | 0.2277 | Q9Y239;G3XAL1;A0A1B0GX71 | NOD1 |
| -0.0638 | 0.8670 | *1.3910* | 0.7314 | 0.7368 | 0.2277 | O00159 | MYO1C |
| -0.1767 | -0.7773 | *-0.1251* | -0.3597 | 0.3626 | 0.2279 | Q9H773 | DCTPP1 |
| 0.6365 | 1.6339 | *0.0741* | 0.7815 | 0.7899 | 0.2287 | E7ENI6;B9ZVM7; E9PDL4;C9IZC7; C9J3Y4;C9J8Y3; F8WET5;Q6LCR0;Q6LCT9;Q05084 | ICA1 |
| -0.0932 | -1.6189 | *-0.5948* | -0.7690 | 0.7776 | 0.2289 | P57678;I3L2C7;E7EN12;I3L399;I3L4A4;I3L4M4 | GEMIN4 |
| -0.0631 | 0.9437 | *0.6583* | 0.5130 | 0.5189 | 0.2290 | P14868;C9J7S3; C9JLC1;H7BZ35; C9JQM9;H7C278 | DARS |
| -0.0282 | 0.9648 | *0.5412* | 0.4926 | 0.4983 | 0.2290 | Q8NE71;H0YGW7;Q5STZ8;F5GYK6 | ABCF1 |
| 0.1652 | 0.3528 | *1.3239* | 0.6140 | 0.6219 | 0.2294 | Q6IA86 | ELP2 |
| -0.2946 | -0.6507 | *-0.0078* | -0.3177 | 0.3221 | 0.2297 | P60891;B1ALA9; A0A2R8Y7H4;B7ZB02;B1ALA7;Q15244 | PRPS1 |
| -0.5700 | -2.4304 | *-0.3646* | -1.1217 | 1.1381 | 0.2299 | Q16696 | CYP2A13 |
| 0.2086 | 0.9014 | *0.1371* | 0.4157 | 0.4222 | 0.2302 | A0A087X080;A0A0U1RRH1;A0A0 | RYR3 |

| L100/ CTRL1 | L100/ CTRL2 | L100/ CTRL3 | Mean | SD | T-test p-value | Accession | Gene Symbol |
|---|---|---|---|---|---|---|---|
| | | | | | | X1KG73;Q15413; A0A1B0GTF2 | |
| 0.0193 | -0.5898 | *-0.3324* | -0.3009 | 0.3057 | 0.2303 | P07864;F5H245; G3XAP5;F5H155 ;F5H5G7 | LDHC |
| -0.3630 | -0.1636 | *-1.3513* | -0.6260 | 0.6360 | 0.2304 | Q9UMY4;A0A087 X0R6 | SNX12 |
| -0.0242 | -0.2228 | *-0.5724* | -0.2731 | 0.2776 | 0.2304 | A0A0A0MRC4;A 0A0A0MSK4;Q86 YR5;A0A087WV F5 | GPSM1 |
| -0.0713 | 1.5551 | *0.9379* | 0.8073 | 0.8210 | 0.2307 | Q8WXA9 | SREK1 |
| -0.4154 | -2.1642 | *-0.4051* | -0.9949 | 1.0127 | 0.2309 | Q86SQ0;A0A1W 2PRT1;E9PFQ4; E9PGF6 | PHLDB2 |
| 0.1382 | -1.5089 | *-1.2187* | -0.8631 | 0.8792 | 0.2312 | G3V0E5;P02786; H7C3V5 | TFRC |
| -0.2369 | -1.0379 | *-3.1396* | -1.4715 | 1.4991 | 0.2312 | P81274 | GPSM2 |
| 0.1784 | 2.9748 | *1.0548* | 1.4027 | 1.4303 | 0.2315 | Q9P2R6;H7BYW 9;K7EJQ1 | RERE |
| 0.0767 | 0.2188 | *0.7525* | 0.3494 | 0.3563 | 0.2316 | Q9UQ35;I3L4D8; A0A087X1W1;I3L 182;I3L1C0;I3L1I 8;I3L3Q8 | SRRM2 |
| 0.0796 | -0.7095 | *-0.7108* | -0.4469 | 0.4559 | 0.2316 | Q8IV36 | HID1 |
| 0.8655 | -0.1004 | *1.0829* | 0.6160 | 0.6299 | 0.2324 | Q86SX6 | GLRX5 |
| -0.4848 | 0.0558 | *-0.4885* | -0.3058 | 0.3132 | 0.2328 | Q13153;B3KNX7; E9PM17;H0YCG 5;E9PKH9;E9PQ W5;E9PRP6;H0Y CM0;E9PJF8;E9 PMP2 | PAK1 |
| 0.1562 | 1.9063 | *0.6008* | 0.8878 | 0.9096 | 0.2330 | Q6ZNJ1;H0Y764; H7C354;H7C3Y7 | NBEAL2 |
| -1.0530 | 0.1232 | *-1.0831* | -0.6710 | 0.6880 | 0.2332 | P62993 | GRB2 |
| -0.1837 | -0.0778 | *-0.6815* | -0.3143 | 0.3223 | 0.2333 | P26038 | MSN |
| -0.3083 | -0.8561 | *-0.0451* | -0.4031 | 0.4137 | 0.2335 | Q8IVG5 | SAMD9L |
| -0.6073 | 0.0720 | *-0.6242* | -0.3865 | 0.3972 | 0.2339 | P61586;C9JNR4; C9JX21;C9JRM1 | RHOA |
| -1.5228 | 0.1504 | *-1.2443* | -0.8722 | 0.8965 | 0.2340 | Q8NBJ4;C9JYM4 | GOLM1 |
| -0.8013 | -0.7449 | *0.0900* | -0.4854 | 0.4991 | 0.2341 | Q13492;B5BU72; E9PJT1;E9PK13; H0YEF7;H0YEH1 | PICALM |
| -0.0597 | -0.4428 | *-1.2236* | -0.5754 | 0.5932 | 0.2349 | Q9Y2T7 | YBX2 |
| 0.0460 | 0.0080 | *0.1342* | 0.0627 | 0.0648 | 0.2355 | P31327;Q5R211 | CPS1 |
| 0.2425 | -2.0631 | *-1.9641* | -1.2616 | 1.3035 | 0.2357 | Q9UKA9 | PTBP2 |
| -0.0770 | -0.0652 | *-0.3866* | -0.1763 | 0.1823 | 0.2359 | Q9UBB6;H7C2R 2;C9J5H8 | NCDN |
| -0.6463 | -0.4518 | *0.0528* | -0.3484 | 0.3608 | 0.2364 | Q01469;I6L8B7;A 8MUU1 | FABP5 |
| -0.3839 | -0.1883 | *-1.5422* | -0.7048 | 0.7318 | 0.2372 | O60486;F5H3A2 | PLXNC1 |
| 0.0650 | -0.5605 | *-0.8312* | -0.4423 | 0.4596 | 0.2375 | Q86VH2;F2Z355 | KIF27 |
| -9.0390 | 1.1610 | *-9.6579* | -5.8453 | 6.0755 | 0.2376 | P02549 | SPTA1 |

| L100/ CTRL1 | L100/ CTRL2 | L100/ CTRL3 | Mean | SD | T-test p-value | Accession | Gene Symbol |
|---|---|---|---|---|---|---|---|
| -0.9722 | -0.2296 | *-0.1269* | -0.4429 | 0.4613 | 0.2382 | P45880;A0A0A0MR02;Q5JSD1;Q5JSD2;A2A3S1 | VDAC2 |
| 0.2150 | 0.3619 | *1.5798* | 0.7189 | 0.7492 | 0.2384 | Q16836;A0A0A0MSE2;A0A1W2PNM1;A0A1W2PQV5;E9PF18;A0A1W2PQ78;A0A1W2PQC2;A0A1W2PRT2;A0A1W2PP40;A0A1W2PQ55;A0A0D9SFP2 | HADH |
| 0.0295 | -0.2446 | *-0.2246* | -0.1466 | 0.1528 | 0.2385 | Q96JG9;H3BS19 | ZNF469 |
| -0.4323 | -0.9964 | *-0.0024* | -0.4770 | 0.4985 | 0.2393 | Q9NTK5;J3KQ32;C9JTK6;C9JCJ9 | OLA1 |
| 0.0407 | -1.3163 | *-0.6720* | -0.6492 | 0.6788 | 0.2394 | P49589;B4DKY1 | CARS |
| -1.1128 | 0.1491 | *-1.1919* | -0.7185 | 0.7525 | 0.2400 | O95347;Q5T821 | SMC2 |
| -0.1325 | -0.7963 | *-2.3855* | -1.1047 | 1.1577 | 0.2402 | P19087;A0A087WZE5 | GNAT2 |
| 0.0087 | -0.8060 | *-0.3699* | -0.3890 | 0.4077 | 0.2402 | Q96CW5 | TUBGCP3 |
| -2.7430 | -0.3039 | *-7.6283* | -3.5584 | 3.7297 | 0.2402 | Q13342;U3KPV9 | SP140 |
| -0.0469 | 0.3820 | *0.5763* | 0.3038 | 0.3189 | 0.2407 | O14686;H0YEF2 | KMT2D |
| -0.0702 | -0.1294 | *0.0058* | -0.0646 | 0.0678 | 0.2408 | O43707;F5GXS2;H7C144;K7EP19 | ACTN4 |
| 0.1132 | -0.9380 | *-0.8114* | -0.5454 | 0.5739 | 0.2415 | P41743 | PRKCI |
| -0.4258 | -0.3766 | *0.0524* | -0.2500 | 0.2631 | 0.2415 | P25398 | RPS12 |
| -0.2465 | -0.2500 | *0.0332* | -0.1544 | 0.1625 | 0.2416 | P41091 | EIF2S3 |
| -0.0124 | 0.1819 | *0.1104* | 0.0933 | 0.0983 | 0.2418 | O75396;A0A087X1A9 | SEC22B |
| -0.1753 | 0.0109 | *-0.3512* | -0.1719 | 0.1811 | 0.2419 | P35241;A0A2R8Y5S7;A0A2R8Y7M3;A0A2R8Y5P0;A0A2R8Y4H6;E9PKN5 | RDX |
| -1.1510 | 0.0222 | *-0.5416* | -0.5568 | 0.5867 | 0.2420 | A0MZ66 | SHTN1 |
| -6.5423 | 0.3345 | *-3.5936* | -3.2671 | 3.4500 | 0.2426 | Q8TBA6;H0YK03 | GOLGA5 |
| 0.5584 | 0.1297 | *0.0693* | 0.2525 | 0.2666 | 0.2427 | Q9UEG4 | ZNF629 |
| -0.2160 | 1.7875 | *2.8525* | 1.4747 | 1.5580 | 0.2428 | Q9P2G3 | KLHL14 |
| -0.0025 | -0.8021 | *-1.9008* | -0.9018 | 0.9531 | 0.2429 | A0A0J9YXM6;Q562E7;E9PDG3 | WDR81 |
| -0.0529 | 1.3458 | *0.6921* | 0.6617 | 0.6998 | 0.2432 | F5H101;F8WE42;Q76FK4 | NOL8 |
| 0.1225 | 1.0716 | *0.2604* | 0.4848 | 0.5128 | 0.2432 | P29401;A0A0B4J1R6;E9PFF2;F8W888;F8WAX4 | TKT |
| 2.0777 | 2.4381 | *-0.2994* | 1.4055 | 1.4874 | 0.2434 | Q9ULE3;F8WAT8 | DENND2A |
| 0.4431 | -0.0630 | *0.6014* | 0.3272 | 0.3470 | 0.2441 | Q92623 | TTC9 |
| 0.3303 | 0.9217 | *0.0304* | 0.4275 | 0.4536 | 0.2442 | Q9C0H2 | TTYH3 |
| -0.4669 | -0.9640 | *0.0286* | -0.4674 | 0.4963 | 0.2444 | Q9H5N1;B4DHR0;H3BNR2;H3BNR8;H3BT64;H3BU67 | RABEP2 |

| L100/ CTRL1 | L100/ CTRL2 | L100/ CTRL3 | Mean | SD | T-test p-value | Accession | Gene Symbol |
|---|---|---|---|---|---|---|---|
| -1.6260 | 0.2345 | *-2.1874* | -1.1930 | 1.2677 | 0.2447 | E7ENS9;H7BXU4;M0QYX2;Q9H2X3 | CLEC4M |
| -0.9743 | -4.4140 | *-0.5715* | -1.9866 | 2.1118 | 0.2448 | Q8N6Q8 | METTL25 |
| -0.4551 | -0.0773 | *-1.4171* | -0.6498 | 0.6908 | 0.2448 | P78347;A0A087X277 | GTF2I |
| 0.0854 | -0.6677 | *-0.5785* | -0.3869 | 0.4115 | 0.2450 | P01116;G3V4K2;G3V5T7 | KRAS |
| -0.1971 | -0.4222 | *0.0100* | -0.2031 | 0.2162 | 0.2453 | P34932;A0A087WYC1;A0A087WTS8 | HSPA4 |
| -0.0960 | -0.5131 | *-1.6379* | -0.7490 | 0.7975 | 0.2453 | Q8WYA0;H0YHE2;F8W1J4 | IFT81 |
| -0.6564 | -0.2968 | *-2.7069* | -1.2200 | 1.3002 | 0.2456 | P62857 | RPS28 |
| 0.0877 | 0.4414 | *1.4358* | 0.6550 | 0.6989 | 0.2460 | Q5T655 | CFAP58 |
| 0.0282 | -0.3256 | *-0.6345* | -0.3106 | 0.3316 | 0.2462 | Q96SI9 | STRBP |
| -0.3526 | -0.2084 | *0.0252* | -0.1786 | 0.1907 | 0.2462 | P50991 | CCT4 |
| 0.0546 | 0.2307 | *0.7843* | 0.3565 | 0.3808 | 0.2463 | Q9UBS4;H7C2Y5 | DNAJB11 |
| -0.0760 | -0.3112 | *-0.0332* | -0.1401 | 0.1497 | 0.2464 | Q9Y262;B0QY89;B0QY90;C9JHP4;C9K0Q7 | EIF3L |
| 0.8245 | 0.0609 | *0.2370* | 0.3741 | 0.3999 | 0.2465 | Q5JNZ5 | RPS26P11 |
| 1.5912 | 2.4598 | *-0.2195* | 1.2772 | 1.3670 | 0.2470 | Q7Z3T8 | ZFYVE16 |
| 0.6954 | 0.7662 | *-0.1053* | 0.4521 | 0.4840 | 0.2471 | P31939;H7C1S2;F8WEF0 | ATIC |
| 0.8592 | 0.1143 | *2.6066* | 1.1934 | 1.2794 | 0.2475 | Q7Z4G4;K7ENP1 | TRMT11 |
| 0.2065 | -1.3368 | *-1.6022* | -0.9108 | 0.9767 | 0.2476 | P27338 | MAOB |
| -0.0664 | 1.3684 | *3.0282* | 1.4434 | 1.5487 | 0.2478 | O60281;J3KNV1;E5RFE6;H0YAU0 | ZNF292 |
| 0.0422 | -1.0616 | *-0.5233* | -0.5142 | 0.5520 | 0.2479 | A0FGR9 | ESYT3 |
| -0.0453 | -0.0421 | *-0.2580* | -0.1151 | 0.1238 | 0.2484 | P36405 | ARL3 |
| -0.0576 | -0.4415 | *-0.0931* | -0.1974 | 0.2122 | 0.2484 | O75955;A0A140T9R1;A0A140T9X0;A2AB09;A2AB10;A0A140T959;A2AB12;A0A140T9C3;A0A0G2JJQ6;A2AB11;A0A140T910;A0A140T957;A2AB13;A0A140T9B1;A0A140T9W4;A0A140T907 | FLOT1 |
| 0.5721 | -0.0763 | *0.4835* | 0.3264 | 0.3516 | 0.2491 | P48735;H0YL11;H0YLL5 | IDH2 |
| 0.0929 | 0.5991 | *1.8928* | 0.8616 | 0.9282 | 0.2491 | Q8IYM0;A0A0C4DGG0;F8VRJ5;E9PMM1 | FAM186B |
| -0.0126 | 0.3976 | *0.1862* | 0.1904 | 0.2051 | 0.2491 | O15145;C9JZD1;F8VR50;K7ESH3 | ARPC3 |
| 0.5112 | 4.5321 | *1.0370* | 2.0268 | 2.1855 | 0.2495 | B7Z524;H0YDX7;J3QT46;Q9Y450;E9PHZ9;E9PS53 | HBS1L |

| L100/ CTRL1 | L100/ CTRL2 | L100/ CTRL3 | Mean | SD | T-test p-value | Accession | Gene Symbol |
|---|---|---|---|---|---|---|---|
| 2.0594 | 3.1387 | *-0.3022* | 1.6320 | 1.7598 | 0.2495 | P12111;C9JNG9;I3L392 | COL6A3 |
| -0.1739 | -0.5989 | *-0.0393* | -0.2707 | 0.2921 | 0.2497 | Q8IVL1;A0A0A0MTE8;A0A0A0MTL4;E9PNV5;E9PLU3 | NAV2 |
| 0.6370 | 0.3095 | *-0.0257* | 0.3069 | 0.3314 | 0.2498 | Q7Z4S6;A0A1B0GV47;H0YHT2;H0YI78 | KIF21A |
| -0.6962 | -0.0571 | *-2.0030* | -0.9187 | 0.9919 | 0.2498 | P61266 | STX1B |
| 0.1341 | 0.1690 | *0.9031* | 0.4021 | 0.4343 | 0.2500 | Q14693 | LPIN1 |
| -0.7126 | -1.7927 | *-0.0006* | -0.8353 | 0.9023 | 0.2500 | Q8TBZ2;H0Y7R1;C9JXR6 | MYCBPAP |
| 0.4014 | 2.5808 | *0.4615* | 1.1479 | 1.2413 | 0.2504 | I3L4C2;Q9UQB8;I3L0M4;I3L113;I3L125;I3L1C8;I3L2M4;I3L327;I3L526;I3L2J6;I3L3J7 | BAIAP2 |
| -0.0171 | 0.2139 | *0.4471* | 0.2146 | 0.2321 | 0.2504 | P18859;A8MUH2 | ATP5PF |
| 0.8210 | 1.5856 | *-0.0869* | 0.7733 | 0.8373 | 0.2508 | Q49MG5;A2VCS9;E7ETZ8;A0A0C4DG83;C9JXH8 | MAP9 |
| 0.1437 | -1.1220 | *-0.8869* | -0.6217 | 0.6732 | 0.2508 | A0A0D9SFC8;A0A0R4J2E0;C9J9Y7 | KCNT1 |
| 0.0866 | 0.4413 | *1.4836* | 0.6705 | 0.7262 | 0.2509 | Q5VIR6;F6VX93;I3L184 | VPS53 |
| -0.0891 | -0.1919 | *0.0063* | -0.0915 | 0.0992 | 0.2509 | Q9NRD9;H0YK19 | DUOX1 |
| -2.7349 | 0.4432 | *-3.5092* | -1.9336 | 2.0945 | 0.2509 | A6NKZ9;E7ETJ9 | TIAL1 |
| -0.3175 | 0.0410 | *-0.5584* | -0.2783 | 0.3016 | 0.2511 | P49903;Q5T5U6 | SEPHS1 |
| -0.6129 | 0.0007 | *-1.5551* | -0.7224 | 0.7837 | 0.2514 | Q9GZP4;X6R8S9;X6RHB9 | PITHD1 |
| -0.2706 | 0.0441 | *-0.3110* | -0.1792 | 0.1944 | 0.2515 | Q9NZL3 | ZNF224 |
| -1.0654 | -0.0319 | *-2.8726* | -1.3233 | 1.4378 | 0.2519 | D6RDH9;E2QRM8;Q9Y5W7;D6RBA7;D6REK1 | SNX14 |
| -0.3310 | -0.5859 | *0.0432* | -0.2912 | 0.3164 | 0.2520 | H0YNL8;P48200;H0YLE0 | IREB2 |
| -0.8089 | -0.0041 | *-0.3084* | -0.3738 | 0.4064 | 0.2521 | P30085;Q5T0D2 | CMPK1 |
| -0.1853 | -0.9925 | *-0.1429* | -0.4402 | 0.4787 | 0.2522 | A0A0A0MRJ9;Q7Z6I6;A0A0A0MRJ8;E9PLT5 | ARHGAP30 |
| -0.1820 | -0.6093 | *-0.0327* | -0.2747 | 0.2992 | 0.2528 | Q04837;A0A0G2JLD8;C9K0U8;E7EUY5 | SSBP1 |
| -0.1097 | 0.9568 | *1.8379* | 0.8950 | 0.9753 | 0.2529 | P52294;C9JYI4 | KPNA1 |
| 0.0939 | 0.6855 | *0.1320* | 0.3038 | 0.3311 | 0.2529 | Q8TAA3;A0A087WYS6 | PSMA8 |
| -0.1873 | 1.1798 | *1.1754* | 0.7226 | 0.7880 | 0.2531 | P42704;A0A0C4DG06;B8ZZ38;C9JCA9 | LRPPRC |
| -0.0262 | 0.6833 | *1.6363* | 0.7644 | 0.8342 | 0.2534 | P17948;H9N1E7 | FLT1 |
| -0.0100 | 0.4866 | *0.2076* | 0.2281 | 0.2490 | 0.2534 | Q9ULM2 | ZNF490 |

| L100/ CTRL1 | L100/ CTRL2 | L100/ CTRL3 | Mean | SD | T-test p-value | Accession | Gene Symbol |
|---|---|---|---|---|---|---|---|
| -0.3696 | -0.1961 | *-1.7110* | -0.7589 | 0.8291 | 0.2538 | Q8TDY2 | RB1CC1 |
| 2.8338 | 2.4010 | *-0.4074* | 1.6092 | 1.7597 | 0.2541 | Q9H4G0;A0A1B0 GTW6;H0Y482 | EPB41L1 |
| 0.0477 | -0.5361 | *-0.3208* | -0.2697 | 0.2952 | 0.2544 | O95671 | ASMTL |
| 0.0645 | -0.3792 | *-0.4315* | -0.2487 | 0.2725 | 0.2547 | P07954 | FH |
| -0.3822 | -0.2174 | *0.0307* | -0.1896 | 0.2078 | 0.2548 | P00387;B1AHF3 | CYB5R3 |
| -0.0905 | 0.7113 | *0.5307* | 0.3838 | 0.4206 | 0.2548 | A0A087WTY0;B4 DXR9 | ZNF732 |
| -0.2062 | 1.5410 | *2.8218* | 1.3855 | 1.5200 | 0.2551 | P62829;J3KTJ3; B9ZVP7;C9JD32; J3KT29 | RPL23 |
| -0.6451 | -0.3120 | *-2.9218* | -1.2930 | 1.4204 | 0.2556 | Q13615;C9JLU3 | MTMR3 |
| 0.1296 | -0.0122 | *0.2774* | 0.1316 | 0.1448 | 0.2561 | Q9UMS4;F5GY5 6;F5H2I0 | PRPF19 |
| 0.4063 | 1.9477 | *0.2251* | 0.8597 | 0.9466 | 0.2563 | M0QXB4;O14579 ;M0R061 | COPE |
| 0.1587 | -1.0317 | *-1.7508* | -0.8746 | 0.9644 | 0.2568 | Q14CX7 | NAA25 |
| -8.2059 | -1.4766 | *-1.1473* | -3.6100 | 3.9836 | 0.2571 | C9JPR7;Q14DL9 ;Q9NY25 | CLEC5A |
| 0.6040 | 0.7125 | *-0.1069* | 0.4032 | 0.4451 | 0.2572 | O75475 | PSIP1 |
| 0.1943 | -1.2816 | *-1.0877* | -0.7250 | 0.8020 | 0.2579 | Q96HH9;D6R9P9 ;D6RFH3;E9PD0 9;D6REP5 | GRAMD2B |
| 1.4033 | 4.5104 | *0.1587* | 2.0241 | 2.2413 | 0.2582 | Q6PCB5;C9JM20 ;H7C2D3 | RSBN1L |
| 0.3133 | -1.7992 | *-1.8935* | -1.1265 | 1.2478 | 0.2583 | Q6ZQQ6;E7ESW 6 | WDR87 |
| -0.8948 | -0.0952 | *-0.1913* | -0.3938 | 0.4366 | 0.2586 | P30040;F8VY02 | ERP29 |
| 0.0144 | 0.3658 | *1.0615* | 0.4806 | 0.5329 | 0.2587 | Q05BV3;H0YJ79; H0YJX1;L8EBH5 | EML5 |
| -0.1494 | 0.8234 | *1.0823* | 0.5854 | 0.6494 | 0.2588 | P08133;E5RFF0; E5RK63;E5RJF5; E5RI05;H0YC77; E5RIU8;E5RJR0 | ANXA6 |
| -0.3087 | -3.7128 | *-0.8982* | -1.6399 | 1.8192 | 0.2588 | O95831;E9PMA0 | AIFM1 |
| -0.3917 | 0.0717 | *-0.4651* | -0.2617 | 0.2911 | 0.2597 | P35611;E7ENY0; E7EV99;A0A0A0 MSR2;H0Y9H2;D 6RF25;H0YFD8; D6RAH3;D6RJE2 | ADD1 |
| -0.0679 | 0.5929 | *1.2455* | 0.5902 | 0.6567 | 0.2599 | P35749 | MYH11 |
| -0.2900 | -0.4064 | *0.0535* | -0.2143 | 0.2391 | 0.2607 | O95571;M0QXB5 ;M0QY80 | ETHE1 |
| -0.0950 | 2.0288 | *0.9224* | 0.9521 | 1.0622 | 0.2607 | A0AVT1;H0Y8S8 | UBA6 |
| -2.6897 | 0.3526 | *-1.9115* | -1.4162 | 1.5805 | 0.2608 | Q9Y490 | TLN1 |
| 0.2127 | -0.0193 | *0.4870* | 0.2268 | 0.2535 | 0.2613 | Q92900 | UPF1 |
| -0.1204 | -0.5720 | *-0.0590* | -0.2505 | 0.2801 | 0.2615 | Q96HY6;A0A0A0 MRX2 | DDRGK1 |
| -0.6366 | -1.1417 | *0.1019* | -0.5588 | 0.6255 | 0.2618 | P60866;E5RJX2; E5RIP1 | RPS20 |
| -3.2901 | -0.1919 | *-0.8779* | -1.4533 | 1.6273 | 0.2620 | A0A0A0MRJ6;H7 BY58;P22061;F6 S8N6;C9J0F2;F8 | PCMT1 |

| L100/ CTRL1 | L100/ CTRL2 | L100/ CTRL3 | Mean | SD | T-test p-value | Accession | Gene Symbol |
|---|---|---|---|---|---|---|---|
| | | | | | | WAV5;F8WAX2;F8WDT3;H7C4X2 | |
| -0.8099 | -0.1878 | *-0.0685* | -0.3554 | 0.3981 | 0.2621 | E9PF63;O75116;Q14DU5 | ROCK2 |
| -0.8113 | -0.1711 | *-0.0824* | -0.3549 | 0.3977 | 0.2622 | F8W8Y7 | ARMCX4 |
| -0.0710 | -1.5446 | *-0.4383* | -0.6846 | 0.7671 | 0.2622 | A0A0U1RR07;A0A0U1RQP0;A0A0U1RQH1 | SYTL2 |
| -2.8360 | 0.0287 | *-7.8087* | -3.5387 | 3.9657 | 0.2622 | A0A1B0GTS4;A0A1B0GTX2;A0A1B0GU53;A0A1B0GUK4;A0A1B0GVU5;A0A1B0GXG0;E7EN11;H0Y476;O00305;A0A1B0GTA9;A0A1B0GTF6;A0A1B0GTN8;A0A1B0GTP5;A0A1C7CYX2 | CACNB4 |
| 0.1532 | -0.9646 | *-0.8092* | -0.5402 | 0.6055 | 0.2623 | P10644;K7EM13;K7EPB2;C9JSK5;H7BYW5;K7EKR1;P31321 | PRKAR1A |
| -0.0151 | 1.1561 | *3.1723* | 1.4378 | 1.6123 | 0.2624 | P26368;B5BU25;K7ENG2 | U2AF2 |
| 0.1320 | 0.4744 | *1.8966* | 0.8343 | 0.9358 | 0.2625 | Q6NUN0;I3L536 | ACSM5 |
| 0.0758 | 0.6501 | *2.1752* | 0.9670 | 1.0850 | 0.2626 | Q8N7U6 | EFHB |
| -0.1474 | 1.4466 | *0.8527* | 0.7173 | 0.8056 | 0.2630 | Q9UIJ7 | AK3 |
| -0.3954 | -0.6188 | *-3.2997* | -1.4380 | 1.6162 | 0.2632 | O15063;A0A0G2JQ76;A0A0G2JQ84;U3KPV0 | KIAA0355 |
| 0.7794 | 0.2827 | *-0.0042* | 0.3526 | 0.3964 | 0.2633 | F5H212;A0A0D9SFF3;C9J2N3;C9JAW2;C9JLG1 | HIVEP1 |
| 0.3173 | -0.0460 | *0.6265* | 0.2992 | 0.3366 | 0.2635 | Q15366;F8VZX2;H3BRU6;F8W0G4;F8VXH9;H3BSS4;F8VTZ0;H3BSP4;C9IZV9;C9J0A4;C9J5V4;C9J7A9;C9JSA6;C9JTY5;C9JZY3;C9K0A2;F8WC71;P57723 | PCBP2 |
| -0.0016 | -0.3115 | *-0.8932* | -0.4021 | 0.4526 | 0.2637 | Q68DA7;H0YM30;H0YL93 | FMN1 |
| -1.5313 | -1.0394 | *0.1973* | -0.7911 | 0.8906 | 0.2638 | O75382;E9PMK8;E9PMW5 | TRIM3 |
| 0.3051 | 0.6193 | *2.9999* | 1.3081 | 1.4736 | 0.2640 | Q01955;H7BXM4;A0A2R8Y2F0 | COL4A3 |
| -0.1875 | -1.4630 | *-0.2572* | -0.6359 | 0.7171 | 0.2644 | Q9NRC6 | SPTBN5 |
| 0.1502 | 0.2591 | *1.3394* | 0.5829 | 0.6574 | 0.2644 | F2Z2Z2;Q5THR1;Q9H5Z1 | DHX35 |
| 0.0375 | -0.1919 | *-0.2784* | -0.1443 | 0.1632 | 0.2654 | Q9UDT6;H7C4L6 | CLIP2 |
| 0.0907 | 0.1366 | *0.7555* | 0.3276 | 0.3713 | 0.2660 | Q15814 | TBCC |

| L100/ CTRL1 | L100/ CTRL2 | L100/ CTRL3 | Mean | SD | T-test p-value | Accession | Gene Symbol |
|---|---|---|---|---|---|---|---|
| 0.3567 | -1.9354 | *-1.8831* | -1.1539 | 1.3085 | 0.2662 | Q15052 | ARHGEF6 |
| 0.3445 | 0.0495 | *1.2281* | 0.5407 | 0.6133 | 0.2663 | Q9BR76;A0A087WW53;F5H390 | CORO1B |
| 2.8469 | 0.2602 | *0.6058* | 1.2376 | 1.4044 | 0.2664 | O75663 | TIPRL |
| 0.0712 | -2.8209 | *-1.0946* | -1.2814 | 1.4551 | 0.2667 | Q15067;I3L0T4;K7ELT1;I3L2U4 | ACOX1 |
| -0.3234 | -0.2673 | *-1.9822* | -0.8576 | 0.9743 | 0.2668 | P12532;F8WCN3;C9J6W7;C9J8F6;C9J995;C9JSQ1;C9JT96 | CKMT1A |
| -0.9325 | 0.1832 | *-1.4304* | -0.7266 | 0.8262 | 0.2672 | P62888;E5RI99;A0A0B4J213;A0A0C4DH44 | RPL30 |
| 0.3441 | 0.0222 | *1.1231* | 0.4965 | 0.5660 | 0.2680 | Q9Y4F9;F5GX51;A0A2R8YF77;A0A2R8YEE0;F5H029;A0A2R8Y4B9;A0A2R8Y7B3;B7Z6U4;H3BP45 | RIPOR2 |
| 2.3794 | -0.1675 | *1.1361* | 1.1160 | 1.2736 | 0.2684 | B4DDG0;B4E171;P41732 | TSPAN7 |
| -0.0776 | 1.0162 | *2.6016* | 1.1801 | 1.3471 | 0.2685 | Q8NEM7;B4E2D5;F6S7C4;R4GND2 | SUPT20H |
| -0.0343 | 0.1825 | *0.1740* | 0.1074 | 0.1228 | 0.2689 | P52888;K7EP46;K7EL02;K7EIK4;K7EL32 | THOP1 |
| -0.2720 | -0.4654 | *-2.4962* | -1.0779 | 1.2321 | 0.2689 | Q2TAC2 | CCDC57 |
| 0.1981 | -1.2031 | *-2.3739* | -1.1263 | 1.2877 | 0.2690 | A0A087X169;Q5T6S3;B0QZ72;X6RER8 | PHF19 |
| -0.7168 | -0.5960 | *0.1226* | -0.3968 | 0.4538 | 0.2692 | P28074;H0YJM8 | PSMB5 |
| -0.0441 | -0.3234 | *-1.1764* | -0.5146 | 0.5899 | 0.2699 | Q96JG6;H7BZP1 | VPS50 |
| -1.1747 | 0.0527 | *-3.2717* | -1.4646 | 1.6811 | 0.2703 | Q13885 | TUBB2A |
| 0.6746 | 0.1399 | *0.0594* | 0.2913 | 0.3344 | 0.2704 | Q66K74;M0QXQ9;M0QY41;M0QZ50 | MAP1S |
| -0.6127 | 0.0963 | *-0.4545* | -0.3236 | 0.3722 | 0.2710 | P22234;E9PBS1;D6RF62 | PAICS |
| -0.7297 | -0.2545 | *-3.3235* | -1.4359 | 1.6519 | 0.2711 | D6RBV2;D6RIU4;Q12907 | LMAN2 |
| -0.0146 | -0.2735 | *-0.0681* | -0.1187 | 0.1367 | 0.2714 | Q13554;H7BXS4;H7BZC6 | CAMK2B |
| 0.1304 | -0.8621 | *-0.6115* | -0.4477 | 0.5161 | 0.2718 | O00154;K7EKP8 | ACOT7 |
| -0.0813 | -0.8279 | *-0.1590* | -0.3561 | 0.4105 | 0.2718 | O94913;E9PNY7;E9PQ01 | PCF11 |
| 0.1675 | -0.8410 | *-0.8376* | -0.5037 | 0.5813 | 0.2722 | Q9ULM0 | PLEKHH1 |
| -0.2526 | -0.1319 | *-1.3318* | -0.5721 | 0.6607 | 0.2724 | Q9Y371;A0A087WW40 | SH3GLB1 |
| -0.1136 | 0.0240 | *-0.1753* | -0.0883 | 0.1020 | 0.2727 | A0A0J9YX86;A6NCC3;F8WBI6;H3BV12;H3BMP0;I6L899 | GOLGA8Q |
| -0.2707 | -1.2937 | *-0.1058* | -0.5567 | 0.6436 | 0.2728 | P42702 | LIFR |

| L100/ CTRL1 | L100/ CTRL2 | L100/ CTRL3 | Mean | SD | T-test p-value | Accession | Gene Symbol |
|---|---|---|---|---|---|---|---|
| 1.3630 | 1.1135 | *-0.2396* | 0.7456 | 0.8623 | 0.2729 | Q9NR48;F8VWK7 | ASH1L |
| 0.1821 | 0.6276 | *2.7541* | 1.1880 | 1.3745 | 0.2731 | P68133;A6NL76 | ACTA1 |
| -0.0166 | 0.3409 | *0.1404* | 0.1549 | 0.1792 | 0.2731 | P46459;I3L0N3;I3L2G1;K7EQD6;I3L0L3;I3L338;I3L4Q9 | NSF |
| 1.2639 | -0.0559 | *0.5069* | 0.5716 | 0.6623 | 0.2735 | P46734;Q6FI23 | MAP2K3 |
| -0.8974 | -1.1285 | *0.1956* | -0.6101 | 0.7072 | 0.2737 | Q9NSI6;H7BZR9 | BRWD1 |
| -1.7622 | -0.7548 | *0.1021* | -0.8050 | 0.9332 | 0.2737 | O14949 | UQCRQ |
| -0.4768 | -0.4904 | *0.0981* | -0.2897 | 0.3359 | 0.2738 | P62318;H3BT13 | SNRPD3 |
| 3.6179 | 3.1504 | *-0.6752* | 2.0310 | 2.3553 | 0.2739 | P20700;E9PBF6;A0A0D9SFE5;A0A0D9SFY5 | LMNB1 |
| 0.8274 | -5.2551 | *-3.7743* | -2.7340 | 3.1719 | 0.2740 | J3KPJ3;Q8N5S9 | CAMKK1 |
| 3.3498 | 2.0816 | *-0.4422* | 1.6631 | 1.9303 | 0.2742 | A4D1E1;A0A087WUA7 | ZNF804B |
| 0.0227 | 0.8031 | *0.2211* | 0.3490 | 0.4056 | 0.2746 | Q9UHV9 | PFDN2 |
| -0.1643 | 0.7424 | *0.9487* | 0.5089 | 0.5921 | 0.2750 | Q8N283;F6XZD3 | ANKRD35 |
| -0.2620 | -0.6038 | *0.0380* | -0.2759 | 0.3211 | 0.2750 | P46937;H0YCI3 | YAP1 |
| -0.6308 | 0.0631 | *-1.6454* | -0.7377 | 0.8593 | 0.2754 | Q92499;A0A087X2G1;F1T0B3 | DDX1 |
| -0.3691 | -0.6740 | *-3.7080* | -1.5837 | 1.8460 | 0.2756 | O15031;A6QRG9 | PLXNB2 |
| -0.0871 | -0.2609 | *-1.2178* | -0.5220 | 0.6089 | 0.2759 | Q8IWI9 | MGA |
| 1.2935 | 7.8009 | *0.8808* | 3.3251 | 3.8817 | 0.2761 | O00479 | HMGN4 |
| 0.0815 | -0.7225 | *-1.8446* | -0.8285 | 0.9674 | 0.2762 | Q8WX93;H0Y952;D6R9F5;D6R9Z5;D6RBB1;D6RBH5;F8WA26;H0YA05 | PALLD |
| -0.1655 | 0.7357 | *1.0718* | 0.5474 | 0.6398 | 0.2766 | O95155 | UBE4B |
| -0.3455 | 0.0698 | *-0.6436* | -0.3065 | 0.3583 | 0.2767 | P43243;A0A0R4J2E8;A8MXP9;D6REM6;B3KM87;H0Y8T4;D6R991;D6RBK5;D6R8Z5;Q68E03;A0A1B0GX04;D6R9F3;D6RB45;D6RBS2;D6RCM3;D6REK4 | MATR3 |
| 0.5509 | -0.1012 | *0.4491* | 0.2996 | 0.3508 | 0.2772 | Q9BX68 | HINT2 |
| -0.4127 | -2.7204 | *-0.3365* | -1.1565 | 1.3549 | 0.2774 | Q92620 | DHX38 |
| -0.8412 | -0.1244 | *-3.3173* | -1.4276 | 1.6753 | 0.2779 | Q9UNW9;F8VYI3;M0R1A0 | NOVA2 |
| -1.3039 | -0.1941 | *-5.1492* | -2.2158 | 2.6004 | 0.2780 | Q9H857;H7C519 | NT5DC2 |
| 0.4750 | 0.0224 | *0.1141* | 0.2038 | 0.2393 | 0.2780 | A0A0X1KG75;Q53SF7;H7C1G2 | COBLL1 |
| -0.7284 | -0.1984 | *-0.0175* | -0.3147 | 0.3695 | 0.2781 | O00267 | SUPT5H |
| -0.8006 | -0.7762 | *0.1670* | -0.4699 | 0.5518 | 0.2781 | P54727;Q5W0S5;H0Y579;K7ELW | RAD23B |

| L100/ CTRL1 | L100/ CTRL2 | L100/ CTRL3 | Mean | SD | T-test p-value | Accession | Gene Symbol |
|---|---|---|---|---|---|---|---|
| | | | | | | 1;K7ENJ0;P5472 5;Q5W0S4 | |
| -1.0737 | -0.5404 | *0.1060* | -0.5027 | 0.5908 | 0.2785 | P11277 | SPTB |
| 0.5947 | 4.9379 | *0.7485* | 2.0937 | 2.4644 | 0.2790 | Q07866;B5BU63; G3V2E7;F8W6L3 ;G5E9S8;H0YJT3 ;H0YJU9;H0YJL0 ;G3V2P7;H0YGB 8 | KLC1 |
| 1.5560 | 0.2642 | *0.1601* | 0.6601 | 0.7776 | 0.2793 | Q9P0K7 | RAI14 |
| -0.6768 | 0.1413 | *-0.6409* | -0.3921 | 0.4623 | 0.2796 | P51149;C9J8S3; C9J592;C9J4V0; C9IZZ0;C9J4S4; C9J7D1 | RAB7A |
| -0.4348 | -0.1801 | *0.0269* | -0.1960 | 0.2312 | 0.2798 | Q8WUD1;Q5HYI 5;E9PE37 | RAB2B |
| -0.0288 | -0.1524 | *-0.6389* | -0.2734 | 0.3226 | 0.2799 | Q6SZW1;J3KSG 7;J3QRE0 | SARM1 |
| 1.4652 | 0.0273 | *4.9687* | 2.1537 | 2.5416 | 0.2799 | Q6PCE3 | PGM2L1 |
| -0.9718 | -0.8978 | *0.2011* | -0.5561 | 0.6569 | 0.2802 | Q06203;D6RCC8 ;D6RE15 | PPAT |
| 0.2956 | 0.0636 | *1.2776* | 0.5456 | 0.6445 | 0.2802 | P54289 | CACNA2D1 |
| 0.5149 | -0.1212 | *0.7232* | 0.3723 | 0.4399 | 0.2804 | P20618 | PSMB1 |
| -0.0385 | -0.2656 | *-1.0662* | -0.4568 | 0.5399 | 0.2804 | Q9Y6F6;H0YI08; E9PJ61;H0YCL0 | MRVI1 |
| 0.2853 | -0.0205 | *0.8355* | 0.3668 | 0.4338 | 0.2807 | A1A4S6 | ARHGAP10 |
| -1.0208 | -3.1366 | *0.0491* | -1.3694 | 1.6212 | 0.2810 | Q9UKE5;C9JVV1 | TNIK |
| 0.2737 | 0.0475 | *1.1411* | 0.4874 | 0.5773 | 0.2811 | P30419;B7Z8J4; K7EN82 | NMT1 |
| 3.3135 | 0.7829 | *0.1467* | 1.4144 | 1.6752 | 0.2812 | Q8NBU5 | ATAD1 |
| -0.0907 | 0.4411 | *0.3935* | 0.2480 | 0.2942 | 0.2818 | E7EMS2;G3V3D 1;G3V3E8;H0YIZ 1;J3KMY5;P6191 6;G3V2V8 | NPC2 |
| 0.2269 | 1.5150 | *0.1759* | 0.6393 | 0.7589 | 0.2819 | Q3B7T1 | EDRF1 |
| 0.0264 | -0.4235 | *-1.2796* | -0.5589 | 0.6635 | 0.2820 | P15121;E9PCX2; E9PEF9 | AKR1B1 |
| -0.3718 | 0.0386 | *-1.0332* | -0.4555 | 0.5408 | 0.2820 | O14910;H0YIA8; H0YI92 | LIN7A |
| -2.6661 | -0.2154 | *-0.5021* | -1.1279 | 1.3399 | 0.2822 | A0A0A0MQW1;A 0A2R8Y4X0;A0A 2R8Y5H9;Q8IVT 5 | KSR1 |
| 0.4795 | 1.4800 | *-0.0251* | 0.6448 | 0.7660 | 0.2822 | Q0VGE8;A0A0X1 KG74;I3L0H5;M0 QY99;M0QZX7 | ZNF816 |
| 0.6935 | 3.1926 | *0.1792* | 1.3551 | 1.6120 | 0.2827 | P63208;E7ERH2; F8W8N3;E5RGM 3 | SKP1 |
| 1.2734 | -0.0834 | *0.5234* | 0.5712 | 0.6797 | 0.2828 | O95235 | KIF20A |
| -1.6343 | -0.2256 | *-0.2050* | -0.6883 | 0.8193 | 0.2829 | Q6UXN9 | WDR82 |
| -1.2114 | -2.3393 | *0.2624* | -1.0961 | 1.3047 | 0.2829 | Q8IVH8;F8WAZ1 ;V9GY95 | MAP4K3 |
| -0.9457 | 3.9058 | *5.5831* | 2.8477 | 3.3906 | 0.2830 | Q76LX8 | ADAMTS13 |

| L100/ CTRL1 | L100/ CTRL2 | L100/ CTRL3 | Mean | SD | T-test p-value | Accession | Gene Symbol |
|---|---|---|---|---|---|---|---|
| -0.0011 | 0.3383 | *1.1441* | 0.4937 | 0.5882 | 0.2832 | P23921;E9PL69; E9PP77;H0YCY7 | RRM1 |
| -0.2737 | -0.1767 | *-1.7066* | -0.7190 | 0.8566 | 0.2832 | Q9Y608;C9JSU1 | LRRFIP2 |
| -0.5296 | 0.0913 | *-1.2474* | -0.5619 | 0.6699 | 0.2834 | A0A1W2PPJ9;Q7RTN6;A0A1W2PNV7;A0A1W2PPG2;A0A1W2PQF1;A0A1W2PS04; J3QS66;A0A1W2PP78;A0A1W2PPM8;A0A1W2PQ00;A0A1W2PQE8;A0A1W2PR00; A0A1W2PR65;A0A1W2PRQ6;J3KSA2;J3QQS3;Q86YC8 | STRADA |
| 0.0198 | -0.3088 | *-0.9452* | -0.4114 | 0.4906 | 0.2835 | Q9ULH0;A0A1W2PPB7;E9PH70; H0Y8E4;A0A1W2PPY4 | KIDINS220 |
| 1.8545 | 1.6913 | *-0.3932* | 1.0509 | 1.2532 | 0.2835 | P11387 | TOP1 |
| -0.0833 | 2.7263 | *0.9292* | 1.1907 | 1.4230 | 0.2843 | P84098;J3KTE4; J3QR09 | RPL19 |
| -0.0698 | -0.1468 | *-0.8235* | -0.3467 | 0.4147 | 0.2846 | P00441;H7BYH4 | SOD1 |
| -1.1790 | -1.1166 | *0.2583* | -0.6791 | 0.8124 | 0.2846 | Q9Y3S1;F8W9F9;H0Y7T5;A6PVV2;H0Y493;H0Y7J9 | WNK2 |
| -2.2365 | 0.0097 | *-0.6625* | -0.9631 | 1.1529 | 0.2848 | Q92974;V9GYM8;Q5VY93;V9GYG5;V9GZ14;V9GYF5 | ARHGEF2 |
| -0.6979 | -0.3211 | *-3.8821* | -1.6337 | 1.9563 | 0.2850 | H3BNV7;J3KS95; Q9Y3D0 | CIAO2B |
| -0.0464 | -0.3782 | *-1.5454* | -0.6566 | 0.7873 | 0.2854 | Q15911 | ZFHX3 |
| 0.3710 | 0.0587 | *0.0371* | 0.1556 | 0.1869 | 0.2860 | Q5VUJ9;H7BY53;H0Y6F0;H0YHT5;H0Y588 | EFCAB2 |
| 0.1924 | -0.8526 | *-1.6763* | -0.7788 | 0.9365 | 0.2864 | O94874 | UFL1 |
| -0.9044 | 0.0877 | *-0.4208* | -0.4125 | 0.4961 | 0.2865 | Q15233;C9IZL7; H7C367;C9JYS8; C9J4X2 | NONO |
| 0.1484 | -0.6262 | *-0.6724* | -0.3834 | 0.4611 | 0.2865 | O95433;G3V438; H0YJG7;H0YJ63; H0YJU2;G3V3W9 | AHSA1 |
| 0.1608 | 0.0073 | *0.6047* | 0.2576 | 0.3102 | 0.2870 | Q99747;J3KTJ6; J3QKW4;J3QS28 | NAPG |
| -0.1548 | -0.2051 | *-1.4106* | -0.5902 | 0.7110 | 0.2871 | Q12905;B4DY09; X6R6Z1;A0A0A0MRL0 | ILF2 |
| 0.0201 | 2.0142 | *0.5391* | 0.8578 | 1.0346 | 0.2875 | Q92841;A0A1X7SBZ2;A0A1W2PQ51;A0A0U1RQJ0 | DDX17 |

| L100/ CTRL1 | L100/ CTRL2 | L100/ CTRL3 | Mean | SD | T-test p-value | Accession | Gene Symbol |
|---|---|---|---|---|---|---|---|
| -0.3390 | -1.6307 | *-7.4807* | -3.1501 | 3.8056 | 0.2881 | P33991;E5RG31; E5RFJ8 | MCM4 |
| 1.7816 | 2.3892 | *-0.4545* | 1.2388 | 1.4976 | 0.2883 | Q9Y2T4 | PPP2R2C |
| 0.6273 | 5.0112 | *0.6280* | 2.0888 | 2.5309 | 0.2891 | B0QY60 | SUN2 |
| 0.0395 | -0.3576 | *-1.0593* | -0.4591 | 0.5564 | 0.2892 | Q8NEY1;A0A0A0 MRJ3;H0Y6F6;H 7BZD9 | NAV1 |
| 5.4540 | 3.0972 | *-0.7712* | 2.5933 | 3.1431 | 0.2892 | Q15833;R4GMY7 | STXBP2 |
| 1.1693 | 1.2671 | *-0.2854* | 0.7170 | 0.8695 | 0.2894 | P01266;E7EVM0; H0YBJ2;H0YBY1 | TG |
| 0.7650 | 0.2254 | *3.8576* | 1.6160 | 1.9600 | 0.2894 | P40938 | RFC3 |
| -0.3275 | -0.5897 | *0.0813* | -0.2786 | 0.3381 | 0.2896 | P13489;H0YCR7; E9PIM9;E9PLZ3; E9PMJ3;E9PIK5; E9PMN0 | RNH1 |
| -0.1255 | -0.4387 | *-2.2066* | -0.9236 | 1.1221 | 0.2900 | Q8NA03;A0A2R8 YHB5 | FSIP1 |
| -0.0386 | 0.2750 | *0.7701* | 0.3355 | 0.4078 | 0.2902 | P08134;Q5JR08; E9PQH6;Q5JR07 ;E9PN11;Q5JR06 ;E9PLA2 | RHOC |
| 0.7916 | 0.4162 | *4.8258* | 2.0112 | 2.4447 | 0.2902 | Q9Y2B0 | CNPY2 |
| -0.3820 | 1.4788 | *2.4218* | 1.1729 | 1.4267 | 0.2905 | F8WJN3;Q16630 ;F8W084 | CPSF6 |
| -0.0177 | 2.9262 | *0.8290* | 1.2458 | 1.5156 | 0.2905 | Q9H361 | PABPC3 |
| -0.8530 | 0.2075 | *-1.6400* | -0.7618 | 0.9271 | 0.2906 | Q9Y5X3 | SNX5 |
| 0.3550 | -1.3775 | *-2.2980* | -1.1069 | 1.3470 | 0.2907 | Q14586;I3L4I6 | ZNF267 |
| 0.0710 | 2.5104 | *0.5870* | 1.0561 | 1.2856 | 0.2907 | Q9UL03 | INTS6 |
| -0.1812 | 0.0468 | *-0.3033* | -0.1459 | 0.1777 | 0.2910 | P26358;K7ENW7 | DNMT1 |
| 0.1811 | -0.8184 | *-1.7916* | -0.8096 | 0.9864 | 0.2910 | Q96J65 | ABCC12 |
| 0.0040 | -0.3915 | *-1.4158* | -0.6011 | 0.7327 | 0.2912 | P38935;F5GX64; F5H5K3;H3BRR1 | IGHMBP2 |
| 3.0959 | 0.5188 | *0.2510* | 1.2886 | 1.5709 | 0.2913 | Q6ZV73;F8VY01 | FGD6 |
| -0.0291 | -2.1315 | *-0.5381* | -0.8996 | 1.0968 | 0.2913 | E9PHG3;Q8NB1 2 | SMYD1 |
| 0.3158 | -0.0766 | *0.6268* | 0.2887 | 0.3525 | 0.2918 | Q9BWD1 | ACAT2 |
| -0.1626 | -0.1750 | *-1.3824* | -0.5733 | 0.7007 | 0.2921 | D6REX3;O94979 ;D6RHZ5;H7BXG 7;H0YAB3;H0Y8 V7;H0Y9K1 | SEC31A |
| 0.8656 | 1.3146 | *-0.2304* | 0.6499 | 0.7948 | 0.2923 | Q6ZT07 | TBC1D9 |
| -0.0651 | 0.3689 | *0.2438* | 0.1825 | 0.2234 | 0.2927 | O00533;A0A087 X0M8;C9J905 | CHL1 |
| -0.7165 | 0.0610 | *-2.3228* | -0.9927 | 1.2157 | 0.2928 | #N/D | AGO2 |
| -0.0428 | 0.1728 | *0.3412* | 0.1571 | 0.1925 | 0.2931 | Q9C0F0;A0A2R8 Y461 | ASXL3 |
| 0.2079 | 0.0980 | *1.2549* | 0.5203 | 0.6385 | 0.2936 | Q5JY65;Q9BZJ0; A0A0C4DGD7 | CRNKL1 |
| -0.1909 | -0.8173 | *-0.0187* | -0.3423 | 0.4203 | 0.2938 | P41229;F8W7H7; F8WDK1;F8WF5 6 | KDM5C |
| 3.6047 | -0.5955 | *2.2205* | 1.7432 | 2.1404 | 0.2938 | O43572;E7EMD6 | AKAP10 |

| L100/ CTRL1 | L100/ CTRL2 | L100/ CTRL3 | Mean | SD | T-test p-value | Accession | Gene Symbol |
|---|---|---|---|---|---|---|---|
| 0.0877 | -1.4642 | *-0.5310* | -0.6358 | 0.7813 | 0.2940 | Q8NDI1;B5MC86;C9IYU2;H7BZ98 | EHBP1 |
| -0.6632 | 0.1113 | *-1.8193* | -0.7904 | 0.9716 | 0.2942 | Q9UHD1;E9PHZ2 | CHORDC1 |
| -2.2322 | -0.5589 | *-0.0219* | -0.9376 | 1.1528 | 0.2943 | P27694;I3L4R8 | RPA1 |
| 1.3024 | -0.3544 | *2.0557* | 1.0012 | 1.2330 | 0.2948 | A8K3Y2;B5BU16;P52564;K7EIW3;A0A0A0MRF7;E9PRZ0;J3QR49;K7ELM6 | MAP2K6 |
| -0.5297 | -0.1124 | *-2.6008* | -1.0810 | 1.3327 | 0.2952 | P07858;E9PCB3;E9PHZ5;E9PJ67;E9PKQ7;E9PLY3;E9PNL5;E9PQM1;E9PR54;E9PSG5;E9PKX0;E9PL32;E9PS78;R4GMQ5 | CTSB |
| 0.2442 | 1.7917 | *0.1816* | 0.7391 | 0.9120 | 0.2955 | O76003 | GLRX3 |
| -1.5584 | -0.2508 | *-0.1205* | -0.6432 | 0.7952 | 0.2962 | Q96T76;Q5T454 | MMS19 |
| 0.1250 | -0.8919 | *-2.6837* | -1.1502 | 1.4220 | 0.2962 | Q02413 | DSG1 |
| 0.0425 | -0.5369 | *-0.2103* | -0.2349 | 0.2905 | 0.2963 | Q13409;E7EQL5;E7EV09;E7EMU4;E7ETL8;E7EQU2;E7ESD3;E7ET01;E7ERH4;E7ERR6;E7EUM4;E9PGG1;E7EU01 | DYNC1I2 |
| 0.5258 | 0.1458 | *-0.0066* | 0.2217 | 0.2742 | 0.2964 | D6RG30;Q2TB18 | ASTE1 |
| 0.1562 | -0.0246 | *0.4530* | 0.1949 | 0.2411 | 0.2965 | H0YEX6 | MAPK3 |
| 0.8411 | -3.1861 | *-3.6062* | -1.9837 | 2.4554 | 0.2966 | H3BMF4;Q9H2V7;H3BPQ9;H3BR82;H3BT44 | SPNS1 |
| -0.3886 | -1.4189 | *0.0157* | -0.5973 | 0.7397 | 0.2968 | Q14683;G8JLG1;H0Y7K8 | SMC1A |
| -0.3495 | -0.0700 | *-1.7242* | -0.7146 | 0.8855 | 0.2970 | P46109 | CRKL |
| -0.1722 | -4.9698 | *-1.0442* | -2.0621 | 2.5557 | 0.2971 | F6WFR7;Q9P121 | NTM |
| 0.8968 | 1.3236 | *-0.2520* | 0.6561 | 0.8149 | 0.2978 | P27708;F8VPD4;H7C2E4 | CAD |
| -0.8484 | -0.5673 | *0.1604* | -0.4184 | 0.5206 | 0.2985 | P15927;Q5TEJ0 | RPA2 |
| 0.8629 | 1.8525 | *-0.2184* | 0.8323 | 1.0358 | 0.2986 | Q9Y696 | CLIC4 |
| -0.6228 | -0.1338 | *-3.1845* | -1.3137 | 1.6385 | 0.2994 | Q8WYP5;H7C4S1 | AHCTF1 |
| -0.0504 | 1.1709 | *4.4305* | 1.8503 | 2.3165 | 0.3007 | G3V180;G3V1D3;Q9NY33;E9PQ14;E9PPK9;E9PKK8;E9PNX5 | DPP3 |
| -0.3544 | -0.5254 | *-3.8762* | -1.5853 | 1.9858 | 0.3009 | O75531 | BANF1 |
| -0.9728 | 0.2141 | *-2.5041* | -1.0876 | 1.3627 | 0.3010 | J3KR69;Q9H892;A8MTE9;H0YEF6 | TTC12 |
| 0.1014 | -0.6278 | *-1.9056* | -0.8107 | 1.0159 | 0.3010 | F8VSL2;F8VVA3;H0YIC9;P54619 | PRKAG1 |
| 0.0806 | 0.7048 | *0.0784* | 0.2879 | 0.3610 | 0.3012 | Q96A65 | EXOC4 |

| L100/ CTRL1 | L100/ CTRL2 | L100/ CTRL3 | Mean | SD | T-test p-value | Accession | Gene Symbol |
|---|---|---|---|---|---|---|---|
| -0.8858 | 0.2525 | *-1.5925* | -0.7419 | 0.9309 | 0.3015 | C9IZS8;Q03936; Q9Y2Q1 | ZNF92 |
| -0.2712 | -0.2199 | *-2.1828* | -0.8913 | 1.1188 | 0.3016 | P48995 | TRPC1 |
| 0.4316 | -0.0698 | *1.3291* | 0.5637 | 0.7088 | 0.3023 | P41219;H7C5W5 | PRPH |
| 0.3586 | 4.0541 | *0.5507* | 1.6545 | 2.0804 | 0.3023 | P24394 | IL4R |
| 0.0179 | 0.4226 | *0.0790* | 0.1732 | 0.2182 | 0.3030 | Q9Y620;E5RHN9 ;E5RI14 | RAD54B |
| -0.5274 | -0.8610 | *-6.2360* | -2.5415 | 3.2039 | 0.3032 | Q9Y6I3 | EPN1 |
| 0.5843 | -2.1404 | *-2.2904* | -1.2822 | 1.6181 | 0.3036 | H7BXF4;Q9NXE 4;H7C1Q6;C9J64 7;F2Z2I0;F2Z2W 5;F8WF03 | SMPD4 |
| -0.5022 | -0.1310 | *-2.7934* | -1.1422 | 1.4420 | 0.3037 | Q9P2N2;E9PMX 7;J3KT69;E9PL2 6;J3QRC2;J3KT C0;J3QLR3 | ARHGAP28 |
| -1.3136 | 0.2505 | *-0.8433* | -0.6354 | 0.8025 | 0.3038 | P62249;M0R210; M0R3H0;Q6IPX4 ;M0R1M5;M0QX 76 | RPS16 |
| -0.7542 | 0.2219 | *-1.3447* | -0.6256 | 0.7912 | 0.3043 | Q9P2M7;A6PVU 7 | CGN |
| 0.2350 | -0.8814 | *-0.8842* | -0.5102 | 0.6453 | 0.3044 | P62266;D6RD47; D6R9I7;D6RIX0; D6RDJ2 | RPS23 |
| -0.7348 | -4.2381 | *-0.2148* | -1.7292 | 2.1883 | 0.3045 | O60318 | MCM3AP |
| -0.5325 | -3.6536 | *-0.2726* | -1.4863 | 1.8815 | 0.3047 | O60443 | GSDME |
| -5.8584 | 0.3071 | *-1.8512* | -2.4675 | 3.1286 | 0.3052 | P29375 | KDM5A |
| 0.1486 | -1.0227 | *-0.5155* | -0.4632 | 0.5874 | 0.3053 | Q00341;H0Y394; C9J5E5;C9JIZ1; H7C0A4;C9JES8 ;C9JHS7;C9JHZ8 ;C9JK79;C9JT62; C9JZI8;C9J739;C 9JBS3;C9JEJ8;C 9JHN6;C9JHS9; C9JMQ6;C9JQ82 ;H7BZC3;H7C2D 1;A0A024R4E5 | HDLBP |
| 0.4825 | 1.4815 | *-0.0873* | 0.6256 | 0.7941 | 0.3057 | O75937 | DNAJC8 |
| 0.6083 | 1.7572 | *-0.1243* | 0.7471 | 0.9484 | 0.3057 | O75781;A0A087 WTK8;A0A087W WY4 | PALM |
| 0.3923 | 1.3596 | *-0.0494* | 0.5675 | 0.7206 | 0.3058 | Q92834 | RPGR |
| -0.5006 | -0.6044 | *0.1457* | -0.3198 | 0.4064 | 0.3061 | O43427;H0YCE7 ;E9PSD3 | FIBP |
| -0.6425 | -0.4408 | *0.1333* | -0.3167 | 0.4025 | 0.3061 | P18621;A0A087 WXM6;A0A0A6Y YL6;J3KRX5;J3Q QT2;A0A087WW H0;J3KRB3;J3Q S96;J3QLC8;A0A 087WY81;A0A0A 0MRF8;J3KSJ0 | RPL17 |
| -0.4140 | 0.1010 | *-0.3488* | -0.2206 | 0.2804 | 0.3061 | O95782 | AP2A1 |

| L100/ CTRL1 | L100/ CTRL2 | L100/ CTRL3 | Mean | SD | T-test p-value | Accession | Gene Symbol |
|---|---|---|---|---|---|---|---|
| -0.0784 | 0.2585 | *0.3923* | 0.1908 | 0.2426 | 0.3061 | O15371;B0QYA5; B0QYA6;B0QYA8 ;B0QYA4 | EIF3D |
| -0.0810 | 0.3745 | *1.0567* | 0.4501 | 0.5726 | 0.3065 | Q13045;J3KS54; J3QQQ2 | FLII |
| 0.8283 | 2.4062 | *-0.1714* | 1.0211 | 1.2996 | 0.3066 | P04626;J3QLU9; B4DTR1;J3KTI5 | ERBB2 |
| 0.6121 | -0.1275 | *1.7741* | 0.7529 | 0.9586 | 0.3067 | P68431 | HIST1H3A |
| 0.4432 | 0.4053 | *-0.1146* | 0.2446 | 0.3117 | 0.3070 | O94988;D6RCC1 ;D6RFM4;D6RG E4;Q6P521 | FAM13A |
| 0.1609 | -0.9481 | *-3.0408* | -1.2760 | 1.6258 | 0.3070 | P54098;A0A1B0 GTU7;A0A1B0G TQ6;A0A1B0GVT 8;A0A0D9SFM1; A0A1B0GW33;H 0YCD2;H0YCV2; H0YDF1;H0YE43 | POLG |
| 0.1881 | -0.6492 | *-0.7544* | -0.4052 | 0.5165 | 0.3072 | Q9HDC9;H0Y512 | APMAP |
| -1.0721 | -0.1947 | *-0.0417* | -0.4362 | 0.5560 | 0.3072 | Q96CN4 | EVI5L |
| -0.3186 | 1.2579 | *1.1004* | 0.6799 | 0.8683 | 0.3078 | Q96LX7;F2Z395 | CCDC17 |
| -0.2650 | -0.4988 | *0.0801* | -0.2279 | 0.2912 | 0.3081 | P62879;E7EP32; C9JIS1;C9JXA5; C9JZN1 | GNB2 |
| -4.4002 | -0.4764 | *-0.4521* | -1.7762 | 2.2725 | 0.3085 | F6U0I4;H9KV53; Q5U5Z8;E9PR59 ;E9PI49;E9PRJ1; E9PJH3;E9PS54 | AGBL2 |
| -0.5215 | -0.7846 | *0.1614* | -0.3816 | 0.4883 | 0.3086 | A0A1B0GVM0 | BEGAIN |
| -0.2603 | 0.0354 | *-0.9150* | -0.3800 | 0.4863 | 0.3086 | Q9P227;A0A087 WXU2;A0A087W ZZ2 | ARHGAP23 |
| -0.8151 | 0.2420 | *-0.9736* | -0.5155 | 0.6609 | 0.3092 | Q5TAX3;A0A0C4 DFM7;E9PKY2;E 9PKX1;E9PRG2; X6R5G7;H0YDJ1 ;H0YEE8;E9PQS 7;K7ERA4 | TUT4 |
| -0.1792 | -2.2151 | *-0.2849* | -0.8931 | 1.1461 | 0.3096 | P33316;H0YKC5; H0YNW5;A0A0C 4DGL3;H0YKI0;H 0YMM5;H0YNJ9 | DUT |
| -0.2601 | -0.8590 | *0.0447* | -0.3581 | 0.4597 | 0.3097 | P05387;H0YDD8 | RPLP2 |
| -0.9569 | 3.5536 | *3.3634* | 1.9867 | 2.5510 | 0.3098 | P48378 | RFX2 |
| 0.2094 | -1.2938 | *-4.3738* | -1.8194 | 2.3364 | 0.3098 | Q9H257;A0A286 YFD5 | CARD9 |
| 0.0882 | -0.5357 | *-0.2852* | -0.2442 | 0.3140 | 0.3103 | P18065;C9JMY1; C9JW52 | IGFBP2 |
| -0.3334 | 0.1050 | *-0.5409* | -0.2564 | 0.3297 | 0.3103 | P51991;H7C1J8 | HNRNPA3 |
| 0.3448 | -1.7269 | *-1.0917* | -0.8246 | 1.0614 | 0.3107 | Q8N3U4;B1AMT 4;B1AMS8;B1AM S9;B1AMT0;B1A MT1;B1AMT2;B1 AMT3;E7ERE6 | STAG2 |
| -0.3417 | -0.0503 | *-0.0208* | -0.1376 | 0.1773 | 0.3112 | P32119;A6NIW5 | PRDX2 |

| L100/ CTRL1 | L100/ CTRL2 | L100/ CTRL3 | Mean | SD | T-test p-value | Accession | Gene Symbol |
|---|---|---|---|---|---|---|---|
| -0.0424 | 0.1345 | *0.1913* | 0.0945 | 0.1219 | 0.3115 | Q5JU69 | TOR2A |
| -3.0225 | 0.9195 | *-3.6883* | -1.9304 | 2.4904 | 0.3115 | Q9H1I8 | ASCC2 |
| -0.4247 | -0.8179 | *0.1326* | -0.3700 | 0.4776 | 0.3117 | A0A0A0MRR1;Q5VV50;Q5VV52 | ZNF691 |
| 1.0962 | 0.2463 | *-0.0044* | 0.4461 | 0.5769 | 0.3124 | P09429;Q5T7C4;A0A0U1RRK2 | HMGB1 |
| -0.2949 | -0.0319 | *-1.5345* | -0.6204 | 0.8024 | 0.3124 | Q96PY6;H0Y8M6;D6RBG5 | NEK1 |
| -0.0772 | 0.3402 | *1.0138* | 0.4256 | 0.5505 | 0.3125 | A2IDC6;Q13084;Q4TT37 | MRPL28 |
| -0.5179 | -0.6671 | *0.1612* | -0.3413 | 0.4415 | 0.3125 | Q12768;E5RFU6;E7EQI7 | WASHC5 |
| 0.7537 | 3.5392 | *0.0129* | 1.4353 | 1.8593 | 0.3130 | Q5H9U9 | DDX60L |
| 0.3682 | -0.0583 | *1.3083* | 0.5394 | 0.6992 | 0.3132 | Q6P1X5;H0YB55 | TAF2 |
| -0.0429 | 0.1874 | *0.5615* | 0.2353 | 0.3050 | 0.3132 | P45974;F5H571 | USP5 |
| -0.0350 | 0.2057 | *0.7133* | 0.2947 | 0.3820 | 0.3133 | Q9H3N1;G3V448 | TMX1 |
| -0.1109 | 0.4328 | *0.3617* | 0.2279 | 0.2955 | 0.3134 | P23396;H0YEU2;F2Z2S8;H0YCJ7;H0YF32;E9PJH4;E9PK82;E9PQ96;E9PSF4;E9PQX2;H0YES8;E9PJN9 | RPS3 |
| -0.1556 | 0.5530 | *1.3787* | 0.5920 | 0.7679 | 0.3134 | O43491;E9PHY5;E9PK52;E9PII3;I6L9B1;A0A2R8Y5B3;E9PIG0;E9PJP4;E9PQD2;E9PQN0;E9PRG1;H0Y5B0 | EPB41L2 |
| 0.1389 | -0.4336 | *-0.8102* | -0.3683 | 0.4779 | 0.3136 | Q96M83;A0A1B0GWZ1;A0A1W2PNE7 | CCDC7 |
| -0.0678 | -0.3955 | *-0.0137* | -0.1590 | 0.2066 | 0.3142 | P22033 | MUT |
| -0.5362 | -0.1511 | *-3.3814* | -1.3562 | 1.7644 | 0.3145 | Q13435;E9PPJ0;E9PJ04;H0YCG1;E9PJT3;H0YEX5 | SF3B2 |
| -3.0146 | 0.3611 | *-1.2399* | -1.2978 | 1.6886 | 0.3146 | A6NMH8;E9PIF1;E9PJK1;E9PRJ8;H0YDJ9;H0YDL9;P60033;E9PM31;H0YEE2 | CD81 |
| 0.0405 | 0.4358 | *2.2861* | 0.9208 | 1.1988 | 0.3148 | P59901;A0A0G2JNG2;A0A075B7A5 | LILRA4 |
| 0.0458 | -0.1601 | *-0.4091* | -0.1745 | 0.2278 | 0.3158 | Q9BRP8 | PYM1 |
| 3.6764 | 0.9470 | *-0.1244* | 1.4997 | 1.9597 | 0.3162 | Q12879;F5GZ52 | GRIN2A |
| 2.0047 | 3.5321 | *-0.6644* | 1.6242 | 2.1240 | 0.3164 | P47755;F8W9N7;A0A0D9SET8;C9JUG7 | CAPZA2 |
| 0.5320 | 0.1800 | *3.5933* | 1.4351 | 1.8773 | 0.3165 | Q4G0M1 | ERFE |
| 1.0210 | 3.7171 | *-0.1714* | 1.5222 | 1.9921 | 0.3167 | P40692;H0Y818;H0Y5L7;H0Y5U4;H0Y793 | MLH1 |

| L100/ CTRL1 | L100/ CTRL2 | L100/ CTRL3 | Mean | SD | T-test p-value | Accession | Gene Symbol |
|---|---|---|---|---|---|---|---|
| 0.3931 | 1.9821 | *0.0164* | 0.7972 | 1.0433 | 0.3167 | Q7L099;D6RCQ1;D6REM9;H0Y8I0 | RUFY3 |
| -1.2397 | -0.0564 | *-0.1884* | -0.4948 | 0.6485 | 0.3172 | P02787;C9JVG0;F8WCI6;F8WEK9;C9JB55;F8WC57 | TF |
| -0.0691 | 0.2248 | *0.2520* | 0.1359 | 0.1781 | 0.3172 | P40925;B9A041;B8ZZ51;C9JRL4;C9JLV6 | MDH1 |
| -1.1742 | 0.3613 | *-1.3114* | -0.7081 | 0.9287 | 0.3175 | P13497 | BMP1 |
| -0.2622 | -0.2958 | *0.0811* | -0.1590 | 0.2086 | 0.3176 | O43747;H3BNR4;H3BR36;H3BUN9;B4DGE1;H3BN71;H3BN75;H3BRM7;H3BS13;H3BV30 | AP1G1 |
| 0.2229 | -0.6771 | *-0.9268* | -0.4603 | 0.6047 | 0.3181 | A0A087WVX5;A0A2U3TZM7;Q08289;A0A087WUH4;A0A087WWJ0;A0A2R8Y555;A6PVM6 | CACNB2 |
| -0.3704 | -4.4056 | *-0.4759* | -1.7507 | 2.2999 | 0.3181 | P38405;K7EPE2 | GNAL |
| -1.9422 | -1.9867 | *0.5786* | -1.1168 | 1.4684 | 0.3184 | P50148;B1AM21 | GNAQ |
| -0.0129 | -0.0191 | *-0.1675* | -0.0665 | 0.0875 | 0.3189 | A0A087WVA8;Q8IWB9 | TEX2 |
| -0.0356 | -0.2425 | *-1.4142* | -0.5641 | 0.7434 | 0.3193 | P21817 | RYR1 |
| -0.0932 | 0.5720 | *0.2797* | 0.2528 | 0.3334 | 0.3195 | Q9HC38;I3L3Q4;I3L1I0;I3L1F4;I3L277;I3L2C2;I3NI24;I3NI27 | GLOD4 |
| -0.7131 | -0.2834 | *-5.2057* | -2.0674 | 2.7263 | 0.3195 | Q9Y333 | LSM2 |
| -0.1933 | 0.0171 | *-0.8447* | -0.3403 | 0.4493 | 0.3199 | Q0VDD8;H7C3Z3;M9MMK7 | DNAH14 |
| -0.7358 | -0.5069 | *0.1716* | -0.3570 | 0.4719 | 0.3203 | Q96NB3;J3QQQ3 | ZNF830 |
| 5.0381 | 2.3401 | *-0.7756* | 2.2008 | 2.9093 | 0.3204 | Q86W11;A0A1W2PP20 | ZSCAN30 |
| -0.0507 | -0.4590 | *-0.0350* | -0.1816 | 0.2404 | 0.3209 | Q9H0C2 | SLC25A31 |
| -0.3619 | 1.1727 | *2.9851* | 1.2653 | 1.6754 | 0.3210 | P53004;C9J1E1 | BLVRA |
| 0.2312 | -0.0680 | *0.6301* | 0.2644 | 0.3502 | 0.3211 | Q9BQE3;F5H5D3;F8VS66;F8VRZ4;F8VWV9;F8VX09;F8VRK0;A6NHL2 | TUBA1C |
| -3.2388 | 0.5619 | *-1.6449* | -1.4406 | 1.9085 | 0.3212 | Q02779;E9PLB1;E9PRQ2;H0YCF5;M0R0A7 | MAP3K10 |
| 0.4460 | -2.5091 | *-1.3003* | -1.1211 | 1.4857 | 0.3213 | A0A075B780;O14787 | TNPO2 |
| -0.6001 | -1.4913 | *0.1887* | -0.6342 | 0.8405 | 0.3213 | Q8IWR0;I3L2K5 | ZC3H7A |
| -0.8055 | -0.0682 | *-4.5061* | -1.7933 | 2.3781 | 0.3215 | P55145;A8K878;H7C2D6 | MANF |
| -1.4007 | 0.0615 | *-0.3624* | -0.5672 | 0.7523 | 0.3216 | Q709C8;A0A0A0MSZ2;A0A0G2JI | VPS13C |

| L100/ CTRL1 | L100/ CTRL2 | L100/ CTRL3 | Mean | SD | T-test p-value | Accession | Gene Symbol |
|---|---|---|---|---|---|---|---|
| | | | | | | W2;A2AE48;A2AE50;A6NGY1;E5RJJ6;H0Y2Q8;H0YDD3;J3KT60;K7EN51;K7ENS6;K7EPV7;Q12899;Q5SPU2;Q64ET8;Q96QU4;Q9BVK2 | |
| -0.1302 | 0.3780 | *0.5606* | 0.2695 | 0.3579 | 0.3221 | A2A2F0;Q86X10;A0A0J9YW54 | RALGAPB |
| 0.1728 | -1.7096 | *-0.5989* | -0.7119 | 0.9463 | 0.3224 | Q58FG1 | HSP90AA4P |
| -3.5275 | 0.4016 | *-1.3160* | -1.4806 | 1.9697 | 0.3227 | Q8N3X6 | LCORL |
| -0.7692 | -0.9905 | *0.2586* | -0.5004 | 0.6666 | 0.3232 | P49773;D6RC06;D6RD60;D6RE99;D6REP8 | HINT1 |
| -0.1684 | -0.0158 | *-0.9779* | -0.3874 | 0.5171 | 0.3240 | P29992;K7EL62;A0A087WVZ3;O95837 | GNA11 |
| -0.4904 | 0.0003 | *-2.5644* | -1.0182 | 1.3614 | 0.3245 | Q9NY46;A0A1W2PRD1;E7EUE6 | SCN3A |
| -0.9654 | -0.4349 | *0.1497* | -0.4169 | 0.5577 | 0.3248 | Q9NPH3;C9J9W1;H7C3W4 | IL1RAP |
| 0.0134 | -0.1979 | *-0.9506* | -0.3784 | 0.5067 | 0.3251 | P51649;C9J8Q5 | ALDH5A1 |
| -0.9926 | -3.9428 | *0.1803* | -1.5850 | 2.1244 | 0.3254 | A0A087WVM4;B7ZM99;Q6UB35 | MTHFD1L |
| -0.1834 | 0.0313 | *-0.7454* | -0.2992 | 0.4011 | 0.3255 | O43526;A0A0D9SG49;A0A1B0GW14;Q4VXP6;A0A0D9SEV1;A0A0D9SF10;A0A0D9SGG3;A0A0G2JH35;A0A0G2JR54;A0A0G2JR98;A0A0G2JRN9;A0A0G2JRU6;A0A0G2JQC9;A0A0G2JQG6;A0A0G2JSD3;A0A0D9SGD4;A0A0G2JS89 | KCNQ2 |
| 0.3286 | -0.0998 | *0.3173* | 0.1821 | 0.2441 | 0.3256 | P09972;A8MVZ9;J3KSV6;J3QKP5;C9J8F3;J3QKK1;K7EKH5 | ALDOC |
| -0.0274 | -1.6207 | *-0.2712* | -0.6398 | 0.8582 | 0.3257 | O43166;G3V4Z3 | SIPA1L1 |
| 0.2870 | -0.8678 | *-1.0084* | -0.5298 | 0.7108 | 0.3258 | Q9BQ95 | ECSIT |
| -0.0318 | -0.0170 | *-0.2754* | -0.1081 | 0.1451 | 0.3260 | O14497;A0A1B0GTU5;H0Y488;A0A087WUV6;A0A1B0GVT5;E9PQW6 | ARID1A |
| -0.1732 | -0.1978 | *0.0570* | -0.1047 | 0.1406 | 0.3261 | Q9UNZ2;F2Z2K0;G3V4V8;R4GMY2;R4GNE6 | NSFL1C |
| 1.1051 | 0.0484 | *0.1489* | 0.4341 | 0.5832 | 0.3263 | Q8N137 | CNTROB |

| L100/ CTRL1 | L100/ CTRL2 | L100/ CTRL3 | Mean | SD | T-test p-value | Accession | Gene Symbol |
|---|---|---|---|---|---|---|---|
| -0.1515 | 0.4260 | *0.9024* | 0.3923 | 0.5278 | 0.3268 | P04792;F8WE04; C9J3N8 | HSPB1 |
| -0.3855 | -2.3507 | *-0.0407* | -0.9256 | 1.2461 | 0.3271 | P13798;C9JIF9;H7C393;C9JLK2;F8WEH5;H0YFE5 | APEH |
| -0.0472 | -0.1215 | *0.0158* | -0.0510 | 0.0687 | 0.3273 | A2VDJ0;H0Y2M0 | TMEM131L |
| 0.4360 | -2.3657 | *-1.2044* | -1.0447 | 1.4077 | 0.3274 | P52948;H0YEN4; H7C3P6;H0YES7;H0YCT1 | NUP98 |
| -4.4478 | -1.9451 | *0.6847* | -1.9027 | 2.5665 | 0.3278 | H3BPZ1;H3BRL8;H3BS72;Q9P035 | HACD3 |
| -0.9289 | -0.6404 | *0.2290* | -0.4468 | 0.6028 | 0.3279 | P14649;F8W1I5 | MYL6B |
| -0.6460 | 0.1968 | *-1.9253* | -0.7915 | 1.0685 | 0.3281 | B2R6F3;P84103; A0A087X2D0 | SFRS3 |
| -0.7075 | 2.0180 | *2.6761* | 1.3289 | 1.7940 | 0.3281 | C9JLV4;O14727 | APAF1 |
| 0.1805 | -0.7475 | *-2.7086* | -1.0919 | 1.4750 | 0.3283 | O43663 | PRC1 |
| -0.2489 | -0.6575 | *-5.2577* | -2.0547 | 2.7814 | 0.3291 | E9PFK9;Q9UJ41; A0A0A0MSJ3 | RABGEF1 |
| 0.3046 | -0.0845 | *0.2433* | 0.1545 | 0.2092 | 0.3293 | Q5TF21;E9PJP2; H3BRB8 | SOGA3 |
| -0.2279 | 0.0063 | *-1.2257* | -0.4824 | 0.6543 | 0.3298 | P09622;E9PEX6; A0A1W2PR83;F2Z2E3;F8WDM5 | DLD |
| 0.0421 | -0.6883 | *-3.5531* | -1.3998 | 1.9003 | 0.3301 | Q03164;E9PR05; H0YEU4;H7BYJ6;H7C5V8;H7C5W4 | KMT2A |
| -0.3278 | 0.9918 | *1.0725* | 0.5788 | 0.7862 | 0.3303 | Q16695;B4DEB1; K7EMV3;Q5TEC6;Q6NXT2;K7EP01 | HIST3H3 |
| -0.4362 | -2.6958 | *-0.0358* | -1.0559 | 1.4343 | 0.3304 | Q6DKJ4;A0A0G2JQK5;I3L4V6 | NXN |
| -0.5234 | 3.0773 | *1.4265* | 1.3268 | 1.8024 | 0.3304 | A0A0B4J1Y2 | TTC21A |
| -1.1569 | 0.3077 | *-0.8609* | -0.5700 | 0.7744 | 0.3304 | P09936;D6RE83; D6R956;D6R974 | UCHL1 |
| -1.4959 | -0.2642 | *-0.0001* | -0.5867 | 0.7984 | 0.3310 | P0C7V7 | SEC11B |
| -0.4409 | 0.1655 | *-0.8161* | -0.3638 | 0.4953 | 0.3312 | C9JJQ8 | TUBA4A |
| -2.7262 | -0.1362 | *-0.3186* | -1.0603 | 1.4456 | 0.3317 | Q15274;C9JCJ5 | QPRT |
| 0.0609 | 0.0156 | *0.4579* | 0.1781 | 0.2433 | 0.3325 | Q13023;G3V3H7; G3V3B5;G3V3H2 | AKAP6 |
| -0.3020 | -0.1479 | *-2.7373* | -1.0624 | 1.4526 | 0.3328 | H3BM40;H3BMR6;H3BN53;H3BN97;H3BNE7;H3BNK4;H3BNQ1;H3BNX9;H3BPH8;H3BQ86;H3BQJ4; H3BR92;H3BRT9;H3BU08;H3BVF3;Q9ULP0 | NDRG4 |
| -0.9558 | -0.2010 | *0.0276* | -0.3764 | 0.5147 | 0.3328 | P51858;H3BPQ6; M0R0J3 | HDGF |
| -0.5023 | 0.1910 | *-0.8964* | -0.4026 | 0.5505 | 0.3328 | Q07020;G3V203; J3QQ67;H0YHA7 | RPL18 |

| L100/ CTRL1 | L100/ CTRL2 | L100/ CTRL3 | Mean | SD | T-test p-value | Accession | Gene Symbol |
|---|---|---|---|---|---|---|---|
| | | | | | | ;F8VUA6;F8VYV2;A0A075B7A0 | |
| -0.6546 | -0.7101 | *0.2202* | -0.3815 | 0.5218 | 0.3329 | A0A0U1RR27 | DENND4A |
| -0.8104 | 0.3061 | *-1.7257* | -0.7433 | 1.0176 | 0.3332 | H7C3T2;Q2TAZ0 | ATG2A |
| -1.5863 | -0.1915 | *-0.0691* | -0.6156 | 0.8428 | 0.3332 | Q8WVV9;B7WPG3;H7BXH8;C9JJZ7;H7C464 | HNRNPLL |
| -0.1524 | 0.8043 | *3.5125* | 1.3881 | 1.9009 | 0.3334 | A0A0A0MSS2;A0A2R8YF04;Q3ZCN5;H7BXL6;H0YIL4 | OTOGL |
| 1.3569 | -0.4284 | *4.2891* | 1.7392 | 2.3819 | 0.3334 | Q6NVY1 | HIBCH |
| 0.8884 | -0.3281 | *2.1213* | 0.8939 | 1.2247 | 0.3336 | Q8NB25;A0A087X2A7;H7BY63;E7EQ67;H0YBC0;B9DI78 | FAM184A |
| 1.8367 | 4.6818 | *-0.6649* | 1.9512 | 2.6752 | 0.3338 | Q8ND24 | RNF214 |
| 0.2179 | -0.7287 | *-2.4608* | -0.9906 | 1.3584 | 0.3339 | A6NJL1 | ZSCAN5B |
| -0.5506 | 0.2116 | *-1.0683* | -0.4691 | 0.6438 | 0.3342 | Q3KQU3 | MAP7D1 |
| -0.3448 | -0.0331 | *-2.2812* | -0.8863 | 1.2180 | 0.3346 | H7BXZ6;Q8IXI2;K7EIQ7 | RHOT1 |
| 0.1239 | -1.3805 | *-0.4077* | -0.5548 | 0.7629 | 0.3349 | O14818;H0Y586;F5GY34 | PSMA7 |
| 0.0692 | 0.1857 | *1.5858* | 0.6136 | 0.8440 | 0.3350 | Q04727 | TLE4 |
| -1.3637 | 0.5132 | *-1.9623* | -0.9376 | 1.2916 | 0.3356 | Q8TE82;D6REB6;D6RI07;E7EQR1;Q6NVH2 | SH3TC1 |
| -0.0798 | 0.8804 | *0.2596* | 0.3534 | 0.4869 | 0.3357 | Q8N944;C9J4B8;C9JS07 | AMER3 |
| 0.0349 | 0.1580 | *1.1995* | 0.4641 | 0.6398 | 0.3358 | O43283 | MAP3K13 |
| 0.4475 | 1.4202 | *-0.1485* | 0.5731 | 0.7919 | 0.3367 | V9GYY6 | MCAT |
| 0.0537 | 2.7024 | *0.3795* | 1.0452 | 1.4444 | 0.3367 | Q8WZ64;D6RAD6 | ARAP2 |
| 0.1932 | -0.0595 | *0.6803* | 0.2714 | 0.3760 | 0.3377 | Q06323;H0YKK6 | PSME1 |
| -0.1166 | 0.5022 | *0.2966* | 0.2274 | 0.3152 | 0.3378 | G5E994;Q5VW38;U3KQD2 | GPR107 |
| 0.1454 | -0.0521 | *0.4211* | 0.1715 | 0.2377 | 0.3379 | P49902 | NT5C2 |
| 1.2577 | 0.1769 | *0.0222* | 0.4856 | 0.6731 | 0.3379 | A0A087WTW5;Q9UKL3;A0A096LP21 | CASP8AP2 |
| -0.3525 | -0.2988 | *-4.2781* | -1.6431 | 2.2821 | 0.3386 | A0A0C4DFV2;Q5TBQ0;Q5TBQ1;Q9BX40 | LSM14B |
| 2.8537 | -0.8733 | *2.4231* | 1.4678 | 2.0389 | 0.3386 | F5GYK2;Q9NRL3 | STRN4 |
| 0.2622 | -0.0051 | *1.6132* | 0.6234 | 0.8675 | 0.3393 | Q5T035 | C9orf129 |
| -0.3380 | -0.3068 | *-4.2760* | -1.6403 | 2.2826 | 0.3393 | Q9Y2Z0 | SUGT1 |
| -0.0181 | -0.4690 | *-0.0531* | -0.1801 | 0.2509 | 0.3397 | E5RHG8;Q15369;R4GMY8 | ELOC |
| 2.4897 | 4.0614 | *-0.9880* | 1.8544 | 2.5840 | 0.3398 | Q6NV74;C9JK00;C9JXD6;C9JFM2 | KIAA1211L |
| 0.2156 | -0.0352 | *1.0914* | 0.4239 | 0.5915 | 0.3403 | A0A1B0GW77;P49419;A0A1B0GTJ4;A0A1B0GUA | ALDH7A1 |

| L100/ CTRL1 | L100/ CTRL2 | L100/ CTRL3 | Mean | SD | T-test p-value | Accession | Gene Symbol |
|---|---|---|---|---|---|---|---|
| | | | | | | 1;A0A1B0GW82;A0A1B0GV49;A0A1B0GTG2;A0A1B0GUY0;A0A1B0GTY9;A0A0J9YWF7;H0YHM6;A0A0J9YWK1;A0A0J9YWM6;A0A1B0GW65;F8VVF2;F8WD33;F8WDY6 | |
| -1.0989 | 0.3287 | *-4.2165* | -1.6622 | 2.3244 | 0.3411 | A0A087WSY9;Q16881;A0A087WSW9;A0A182DWI3;E9PIR7;F8W809;E9PKD3;E2QRB9;A0A0B4J225;E9PIZ5;E9PKI4;E9PLT3;E9PQI3 | TXNRD1 |
| -0.1756 | 0.6110 | *0.4597* | 0.2983 | 0.4174 | 0.3413 | P28331;B4DJ81;C9JPQ5;F8WDL5 | NDUFS1 |
| -0.0971 | 0.2468 | *0.6272* | 0.2589 | 0.3623 | 0.3413 | Q86TG7;A0A087WUL4;A0A087WX23;A0A087WXK2;A0A087WZG9;B4DSP0;A0A087WYS2 | PEG10 |
| 1.1866 | 0.4336 | *-0.1664* | 0.4846 | 0.6780 | 0.3413 | O15516 | CLOCK |
| -0.1660 | -0.6987 | *-5.8948* | -2.2531 | 3.1650 | 0.3428 | Q7L591;D6RAM3;D6RC22 | DOK3 |
| 0.6431 | 0.8184 | *-0.2446* | 0.4056 | 0.5699 | 0.3429 | Q9UI46;A0A087WWV9;Q5T8G8 | DNAI1 |
| -0.2550 | -0.4598 | *0.1057* | -0.2030 | 0.2863 | 0.3443 | Q99733;C9JZI7;A8MXH2;C9J6D1;E9PNJ7;H0YCI4;E9PJJ2;E9PKT8;E9PNW0;E9PP22;E9PS34;E9PKI2 | NAP1L4 |
| -0.1789 | -0.0338 | *-1.4765* | -0.5631 | 0.7943 | 0.3444 | Q9NQC3;A0A0U1RQR6;H7C106;C9J685;F8W914 | RTN4 |
| 0.1476 | -1.2087 | *-0.3905* | -0.4839 | 0.6829 | 0.3446 | P52758;H0YB34;H0YBX3 | RIDA |
| -0.0842 | 0.2065 | *0.5273* | 0.2166 | 0.3059 | 0.3449 | Q01780;K7EJ37 | EXOSC10 |
| -0.2950 | 0.0365 | *-1.7123* | -0.6569 | 0.9289 | 0.3453 | Q9BX66 | SORBS1 |
| 0.0776 | -0.2110 | *-0.7032* | -0.2789 | 0.3948 | 0.3457 | Q9C0C2;A0A2R8Y5C4;E9PKK0 | TNKS1BP1 |
| -0.0975 | 0.4493 | *0.2314* | 0.1944 | 0.2753 | 0.3458 | Q32P51 | HNRNPA1L2 |
| 0.1849 | 1.0753 | *-0.0255* | 0.4116 | 0.5843 | 0.3468 | Q9BT09;A0A0C4DFY8 | CNPY3 |
| 0.7553 | -0.2365 | *0.6147* | 0.3779 | 0.5366 | 0.3469 | Q8IUR6;E5RI19 | CREBRF |
| 0.0724 | -0.3697 | *-1.9663* | -0.7545 | 1.0724 | 0.3472 | J3KNX4;Q2TAK8;A0A0D9SGJ8 | MUM1 |
| -0.7388 | -0.3308 | *0.1407* | -0.3097 | 0.4401 | 0.3472 | C9J1P0;C9JZB1;Q96HH4 | TMEM169 |

| L100/ CTRL1 | L100/ CTRL2 | L100/ CTRL3 | Mean | SD | T-test p-value | Accession | Gene Symbol |
|---|---|---|---|---|---|---|---|
| 0.5138 | -1.4899 | *-1.4237* | -0.7999 | 1.1382 | 0.3476 | P22748;K7EKY5;K7ENI8 | CA4 |
| -0.2419 | -0.1219 | *0.0521* | -0.1039 | 0.1478 | 0.3477 | P50990;H7C4C8;H7C2U0 | CCT8 |
| 0.3258 | -1.7605 | *-0.7638* | -0.7328 | 1.0435 | 0.3479 | M0QY97;Q9UPT8 | ZC3H4 |
| -0.1562 | -0.3175 | *-3.5970* | -1.3569 | 1.9416 | 0.3497 | O00178;F5H716;F5H257 | GTPBP1 |
| -0.8505 | -0.3358 | *0.1439* | -0.3475 | 0.4973 | 0.3498 | Q13404;I3L0A0;G3V2F7;A0A0A0MSL3;D6RG00;E5RIF1 | UBE2V1 |
| -0.6897 | -1.9286 | *0.2894* | -0.7763 | 1.1115 | 0.3500 | P50570;K7EPK9 | DNM2 |
| 0.1948 | -0.6125 | *-2.6588* | -1.0255 | 1.4710 | 0.3506 | Q9Y6K8 | AK5 |
| -0.1700 | 0.0149 | *-1.1313* | -0.4288 | 0.6154 | 0.3508 | A0A0B4J2C3;P13693;Q5W0H4;E9PJF7;J3KPG2;H0YCX0;Q56UQ5 | TPT1 |
| 0.0345 | 0.0433 | *0.6032* | 0.2270 | 0.3258 | 0.3509 | P04275 | VWF |
| 0.1579 | -0.0246 | *0.9541* | 0.3625 | 0.5204 | 0.3510 | L8E898 | MUC12 |
| -1.8120 | 0.7739 | *-4.9698* | -2.0026 | 2.8766 | 0.3512 | Q5JSH3 | WDR44 |
| -0.1946 | 0.4657 | *1.3806* | 0.5505 | 0.7910 | 0.3513 | Q9Y263;E5RIM3;H0YBW4 | PLAA |
| -0.2594 | 0.0303 | *-1.6732* | -0.6341 | 0.9115 | 0.3515 | Q9NZZ3 | CHMP5 |
| -2.1161 | 0.5827 | *-1.3746* | -0.9694 | 1.3943 | 0.3517 | C9JXG5;F8WCP5;H7C441;Q13769 | THOC5 |
| -0.9156 | -1.3898 | *0.3870* | -0.6395 | 0.9200 | 0.3518 | P48643;B7ZAR1;E9PCA1;E7ENZ3;D6RIZ7;H0Y914 | CCT5 |
| -0.1310 | -0.5959 | *0.0405* | -0.2288 | 0.3293 | 0.3519 | Q16181;E7EPK1;E7ES33;G3V1Q4;A0A0U1RRM2;A0A0U1RRE1;Q5JXL7;A0A0U1RRH9;A0A0U1RRD1 | SEPT7 |
| -1.2590 | 0.5245 | *-1.7946* | -0.8430 | 1.2142 | 0.3522 | O95995;H3BP65 | GAS8 |
| -0.2939 | 0.6761 | *1.8357* | 0.7393 | 1.0662 | 0.3527 | Q8NCQ5;J3QLF9;J3KRT3 | FBXO15 |
| -0.5741 | -0.6360 | *0.2197* | -0.3302 | 0.4772 | 0.3535 | P08581 | MET |
| -0.1196 | 0.6153 | *0.2673* | 0.2543 | 0.3676 | 0.3535 | P54652 | HSPA2 |
| -0.6496 | -0.3372 | *0.1503* | -0.2789 | 0.4031 | 0.3536 | Q8IWB6 | TEX14 |
| -0.7045 | 0.0990 | *-0.2323* | -0.2793 | 0.4038 | 0.3537 | P38919 | EIF4A3 |
| 2.2819 | 0.7525 | *-0.3210* | 0.9045 | 1.3081 | 0.3538 | Q86VV8 | RTTN |
| 0.4269 | -1.7923 | *-0.9599* | -0.7751 | 1.1211 | 0.3538 | A0A0D9SEV0;A0A0D9SFL2;M0QYH2;M0QYI1;M0R000;M0R3C8;Q96T60 | PNKP |
| 0.2995 | -1.1247 | *-6.0559* | -2.2937 | 3.3351 | 0.3558 | Q86WA8;H3BPF7 | LONP2 |
| 0.1181 | -0.7888 | *-0.2668* | -0.3125 | 0.4552 | 0.3565 | O76013 | KRT36 |
| -0.7778 | -0.5695 | *0.2402* | -0.3691 | 0.5378 | 0.3566 | Q9NVH0;C9JLF4 | EXD2 |

| L100/ CTRL1 | L100/ CTRL2 | L100/ CTRL3 | Mean | SD | T-test p-value | Accession | Gene Symbol |
|---|---|---|---|---|---|---|---|
| 0.1206 | 0.4603 | *-0.0481* | 0.1776 | 0.2589 | 0.3568 | Q13535;H0Y9K2; H0Y8Y6;H0Y8R8 | ATR |
| -0.2237 | -0.4945 | *0.1040* | -0.2047 | 0.2997 | 0.3583 | P36871 | PGM1 |
| -0.1665 | 0.4288 | *1.7677* | 0.6767 | 0.9906 | 0.3583 | Q5XKE5 | KRT79 |
| 0.3811 | -0.1659 | *1.2712* | 0.4955 | 0.7254 | 0.3584 | Q5TA45;C9J979; Q96HV7;A0A087 WYI0;C9IYS7;E9 PNS4;A0A087W XT8;E9PKA4;E9 PI75;E9PIG1;E9 PIL7;E9PQF0;E9 PNH9;H0YCE0;H 0YDB1 | INTS11 |
| -0.2942 | 0.0603 | *-1.8607* | -0.6982 | 1.0222 | 0.3584 | Q92747;A0A1W2 PNV4;E9PF58;F8 WFD3 | ARPC1A |
| -0.3210 | -0.4709 | *0.1399* | -0.2173 | 0.3183 | 0.3585 | Q7Z7A1;Q5JVD3 ;Q5JVD6;Q5JVD 5 | CNTRL |
| 0.5760 | 1.1175 | *-0.2662* | 0.4758 | 0.6973 | 0.3587 | H0Y7W5 | RFX3 |
| -0.8424 | -0.4074 | *0.1900* | -0.3533 | 0.5183 | 0.3592 | P48668;P02538 | KRT6C |
| 0.9098 | 5.1087 | *-0.2535* | 1.9217 | 2.8206 | 0.3593 | Q96M27 | PRRC1 |
| -0.0009 | 0.1398 | *1.1877* | 0.4422 | 0.6494 | 0.3595 | O15381;E9PH71; E7EWK7;E9PGD 8;F8WF01 | NVL |
| -0.0412 | 0.1515 | *0.8730* | 0.3278 | 0.4819 | 0.3600 | Q12756;F8W8V9 ;H7C0K6;C9JBH 1 | KIF1A |
| 1.1485 | 1.6877 | *-0.5050* | 0.7771 | 1.1426 | 0.3600 | O00754;M0QZ24 ;M0QYZ1;M0QZ G6;M0R2P5 | MAN2B1 |
| -0.4214 | 0.1345 | *-2.1960* | -0.8276 | 1.2172 | 0.3601 | Q69YQ0;C9J8U1 ;F8WAN1;C9JLY 8 | SPECC1L |
| 2.0147 | 0.3053 | *-0.0610* | 0.7530 | 1.1079 | 0.3602 | Q70Z35 | PREX2 |
| 0.6986 | -0.0598 | *0.1573* | 0.2653 | 0.3906 | 0.3604 | P28072;A0A087X 2I4;I3L3X7 | PSMB6 |
| 0.2813 | -0.1015 | *1.3319* | 0.5039 | 0.7422 | 0.3606 | P63128;P62684; P62685;P63126; P63130;P63145; P87889;Q7LDI9; Q9YNA8;P62683; Q9HDB9 | ERVK-9 |
| 0.2960 | -0.6503 | *-2.0648* | -0.8063 | 1.1881 | 0.3608 | Q6UVK1 | CSPG4 |
| -0.1041 | -0.0018 | *-0.9304* | -0.3454 | 0.5092 | 0.3609 | Q92785;J3KMZ8 | DPF2 |
| 0.0964 | -0.0449 | *0.2824* | 0.1113 | 0.1641 | 0.3610 | P04075;J3KPS3; H3BPS8;H3BUH 7;H3BR04;H3BM Q8;H3BR68;H3B U78 | ALDOA |
| 0.2314 | 1.6609 | *-0.0359* | 0.6188 | 0.9123 | 0.3610 | O95202 | LETM1 |
| 0.9365 | 0.8938 | *-0.3470* | 0.4944 | 0.7290 | 0.3610 | M0R2J8;B6ZDN3 ;H7C298;Q6ZRR 9 | DCDC1 |

| L100/ CTRL1 | L100/ CTRL2 | L100/ CTRL3 | Mean | SD | T-test p-value | Accession | Gene Symbol |
|---|---|---|---|---|---|---|---|
| 0.1326 | -0.0404 | *0.7315* | 0.2746 | 0.4051 | 0.3612 | Q8TAQ5;K7ELF6;K7EQC9;K7ERS3 | ZNF420 |
| -0.0218 | 0.1003 | *0.6593* | 0.2459 | 0.3631 | 0.3615 | P14324;A0A087WVN4;A0A087X1D8;A0A087X090;A0A087WTP2 | FDPS |
| -0.1224 | 0.5470 | *0.2561* | 0.2269 | 0.3356 | 0.3623 | P60763;J3KSC4;J3QLK0 | RAC3 |
| -8.0655 | -2.2565 | *1.0070* | -3.1050 | 4.5954 | 0.3625 | E9PGC0;P20936 | RASA1 |
| 0.5001 | -0.2377 | *1.4174* | 0.5600 | 0.8292 | 0.3626 | Q9UPV9;A0A0D9SFL5;C9JC32 | TRAK1 |
| -0.0162 | -1.1052 | *-0.1025* | -0.4080 | 0.6054 | 0.3634 | Q8NCN4 | RNF169 |
| 0.0071 | -0.1815 | *-1.5939* | -0.5894 | 0.8750 | 0.3636 | Q6P5Z2 | PKN3 |
| 0.4281 | 1.4574 | *-0.1971* | 0.5628 | 0.8355 | 0.3636 | Q8WVS4;H7C1E8 | WDR60 |
| -0.8850 | 0.1999 | *-5.9669* | -2.2173 | 3.2922 | 0.3637 | Q9P2Q2;Q5T376;A0A1W2PQE7 | FRMD4A |
| -0.0703 | 2.2561 | *0.3275* | 0.8378 | 1.2443 | 0.3638 | P29400 | COL4A5 |
| 0.0816 | 0.0169 | *0.9260* | 0.3415 | 0.5072 | 0.3638 | Q9ULB1;A0A0R4J2G7;A0A1D5RMU6;E7ERL8;A0A0D9SEP4;A0A0U1RRK7;F5GYC7;F8WB18;A0A1B0GTL0;A0A0D9SEM5;A0A0D9SF36;A0A0D9SFY6;H0Y568;H7BYC7;P58400;Q08AH0;A0A0D9SEQ7;A0A0D9SFF4;A0A0D9SG60;A0A1B0GU94;A0A1B0GVF4;E7EQN4 | NRXN1 |
| 1.6588 | 0.8819 | *-0.4219* | 0.7063 | 1.0514 | 0.3647 | Q9BWS9 | CHID1 |
| 0.8880 | -0.0713 | *0.1841* | 0.3336 | 0.4968 | 0.3649 | P16402 | HIST1H1D |
| -3.3985 | -0.9012 | *0.4064* | -1.2977 | 1.9332 | 0.3649 | B7ZC32 | KIF28P |
| -0.2877 | -0.3238 | *0.1179* | -0.1645 | 0.2453 | 0.3652 | Q07157;G3V1L9;A0A087X0K9;G5E9E7;H0Y3R8 | TJP1 |
| 1.9480 | 0.0905 | *0.1091* | 0.7158 | 1.0671 | 0.3652 | Q6P597;K7EL76;K7ENJ3 | KLC3 |
| 0.7289 | -0.3587 | *1.7071* | 0.6924 | 1.0334 | 0.3656 | A0A087WTH5;A0A087WU88;A0A087WWU3;P15382 | KCNE1B |
| -1.2559 | 0.3128 | *-0.6463* | -0.5298 | 0.7908 | 0.3657 | P25787;A0A024RA52;H3BT36;H7C402 | PSMA2 |
| -1.2263 | 0.3182 | *-8.1164* | -3.0082 | 4.4908 | 0.3657 | H7C3P7;P11233;C9JPE8;C9JQB3;C9JYR1 | RALA |
| 0.9340 | -0.2866 | *0.6289* | 0.4255 | 0.6352 | 0.3658 | O15066 | KIF3B |

| L100/ CTRL1 | L100/ CTRL2 | L100/ CTRL3 | Mean | SD | T-test p-value | Accession | Gene Symbol |
|---|---|---|---|---|---|---|---|
| -1.4472 | 0.2445 | *-0.4999* | -0.5675 | 0.8479 | 0.3660 | Q8TEP8;A0A0A0MR42;K7ENP4;H0Y966;K7EPA2;C9JT09;K7ELX0;K7ERF9 | CEP192 |
| -1.7215 | -1.2913 | *0.5691* | -0.8146 | 1.2175 | 0.3662 | P14174 | MIF |
| -1.2639 | 0.3196 | *-0.6536* | -0.5326 | 0.7987 | 0.3674 | P13797;A0A0A0MSQ0 | PLS3 |
| -2.6841 | 1.1433 | *-3.2194* | -1.5867 | 2.3794 | 0.3674 | A2RRP1;H0Y5G7;H7C007;H7C1U4 | NBAS |
| -0.1872 | 0.3950 | *1.4191* | 0.5423 | 0.8132 | 0.3674 | Q9HD67;A0A0A0MQX1;D6RGD1;E9PCN3 | MYO10 |
| 0.0665 | 2.4562 | *0.1759* | 0.8995 | 1.3493 | 0.3675 | P10253 | GAA |
| 0.2468 | -2.0475 | *-0.5314* | -0.7774 | 1.1668 | 0.3678 | Q9UPU5 | USP24 |
| 0.1040 | -0.2377 | *-0.3030* | -0.1456 | 0.2186 | 0.3679 | P05165;A0A1B0GU58;A0A1B0GUX9;A0A1B0GWI4;A0A1B0GWA1;A0A1B0GTR1;A0A2R8Y725 | PCCA |
| -0.0310 | 0.0770 | *0.3858* | 0.1439 | 0.2163 | 0.3682 | Q5THJ4;H3BLS7;A0A2R8Y876;A0A2R8YD87;F5GX56;E9PRM3;Q6ZVT0;R4GMW1 | VPS13D |
| -0.1315 | -0.0339 | *0.0159* | -0.0499 | 0.0750 | 0.3684 | P61018;M0R0X1;Q6PIK3 | RAB4B |
| -0.9410 | 0.4487 | *-3.5171* | -1.3365 | 2.0123 | 0.3690 | C9JEZ4;Q9UKI2 | CDC42EP3 |
| -0.4806 | -0.7532 | *0.2264* | -0.3358 | 0.5056 | 0.3690 | Q02224;A0A087X0P0;D6RBW0 | CENPE |
| -0.1822 | 0.8938 | *0.3588* | 0.3568 | 0.5380 | 0.3695 | Q6ZN16 | MAP3K15 |
| -0.1907 | 0.5594 | *0.4279* | 0.2655 | 0.4006 | 0.3697 | Q15056 | EIF4H |
| -0.3499 | 0.8648 | *0.8980* | 0.4710 | 0.7111 | 0.3700 | Q9H8V3;C9J0L6;C9J1C4;C9JDB4;C9JDV9;C9JTI2 | ECT2 |
| 0.8602 | 3.5171 | *-0.4051* | 1.3241 | 2.0018 | 0.3705 | Q6UB99;X5D778;A0A2R8Y438;A0A087WTN8;A0A2R8YE03;A0A2R8YEI0;A0A2R8Y4T9;A0A2R8Y5V1;A0A2R8Y7Z1;H0Y2U4;H3BNU4;A0A2R8Y728;H0Y3E3 | ANKRD11 |
| -0.6124 | 0.3117 | *-1.3792* | -0.5600 | 0.8467 | 0.3706 | Q15102;M0R389;M0QZT2;M0QXS6;M0R323 | PAFAH1B3 |
| 3.3541 | 3.2056 | *-1.3055* | 1.7514 | 2.6484 | 0.3706 | Q12929;A0A2R8Y4W2;F5H0R8 | EPS8 |
| -0.0521 | -0.6908 | *-0.0120* | -0.2516 | 0.3809 | 0.3710 | E9PNZ4;E9PLY0 | MACF1 |
| -0.0411 | -0.6260 | *-0.0165* | -0.2279 | 0.3450 | 0.3711 | Q6UXK2;Q24JQ5 | ISLR2 |

| L100/ CTRL1 | L100/ CTRL2 | L100/ CTRL3 | Mean | SD | T-test p-value | Accession | Gene Symbol |
|---|---|---|---|---|---|---|---|
| -0.0239 | -0.7431 | *-0.0441* | -0.2704 | 0.4096 | 0.3713 | Q9H582;A0A087WZL9 | ZNF644 |
| 0.2118 | -0.1089 | *0.4951* | 0.1993 | 0.3022 | 0.3715 | P26599;A0A0U1RRM4;A6NLN1;A0A0D9SF20;K7EK45;K7ELW5;A0A087WU68;A0A087WUW5 | PTBP1 |
| -3.7867 | 0.1100 | *-0.4772* | -1.3846 | 2.1009 | 0.3719 | P63241;I3L397;I3L504;Q6IS14;C9J4W5 | EIF5A |
| -0.1778 | 0.5405 | *0.3840* | 0.2489 | 0.3777 | 0.3720 | Q12888;A6NNK5;C9JXV0;H7BZY0;M0R142;H7C3N7 | TP53BP1 |
| 0.0821 | 0.0368 | *1.3246* | 0.4811 | 0.7308 | 0.3723 | A0A087WUI6;Q8WXW3 | PIBF1 |
| -0.2073 | 0.5476 | *0.4890* | 0.2764 | 0.4199 | 0.3723 | P05026;V9GYR2 | ATP1B1 |
| -0.1534 | -0.0065 | *-1.7525* | -0.6375 | 0.9684 | 0.3724 | P49841;B5BUC0 | GSK3B |
| -0.1773 | -1.1847 | *0.0586* | -0.4345 | 0.6603 | 0.3725 | O75592;H7C3U4 | MYCBP2 |
| -0.6058 | -0.5931 | *0.2413* | -0.3192 | 0.4854 | 0.3727 | P61604;B8ZZL8;B8ZZ54 | HSPE1 |
| 0.0591 | 1.9023 | *0.1082* | 0.6899 | 1.0503 | 0.3732 | Q99622;U3KQ85;F5GXW5 | C12orf57 |
| -0.2354 | 0.4523 | *1.0457* | 0.4209 | 0.6411 | 0.3734 | P53992;G5EA31 | SEC24C |
| -0.3817 | 0.7303 | *1.7743* | 0.7076 | 1.0782 | 0.3735 | P61313;A0A2R8YEM3;E7EQV9;A0A2R8Y738;E7EX53;E7ENU7;E7ERA2 | RPL15 |
| -0.0584 | 0.1256 | *0.5894* | 0.2189 | 0.3338 | 0.3739 | P11216;H0Y4Z6 | PYGB |
| 0.1802 | -0.3956 | *-1.9597* | -0.7251 | 1.1074 | 0.3744 | P98196;E9PEJ6 | ATP11A |
| -0.1243 | -0.5651 | *0.0592* | -0.2100 | 0.3208 | 0.3745 | Q5H9L2 | TCEAL5 |
| -0.3969 | -2.3970 | *0.1589* | -0.8784 | 1.3443 | 0.3752 | P50395;Q6IAT1;Q5SX87;Q5SX86;Q5SX90;V9GYF8;V9GYJ7 | GDI2 |
| -0.0467 | 0.2661 | *2.5018* | 0.9070 | 1.3899 | 0.3757 | Q9NUJ1 | ABHD10 |
| -0.7054 | 0.2841 | *-4.3241* | -1.5818 | 2.4259 | 0.3760 | Q9BRZ2;C9JI91 | TRIM56 |
| -1.4282 | 0.2553 | *-0.4768* | -0.5499 | 0.8442 | 0.3764 | Q9UJZ1;A0A087WYB4 | STOML2 |
| -0.6406 | -3.1921 | *0.3023* | -1.1768 | 1.8078 | 0.3766 | P13639 | EEF2 |
| -0.4628 | 0.8964 | *1.6912* | 0.7083 | 1.0893 | 0.3770 | H3BSG2;O75309 | CDH16 |
| 1.0183 | -0.0677 | *0.1640* | 0.3715 | 0.5720 | 0.3774 | P49368;B4DUR8;E9PRC8;Q5SZX9;E9PM09;E9PQ35;A0A1B0GTR8;Q6P4Q7;Q6ZUB1;Q8N957;Q9H8M5 | CCT3 |
| -0.2320 | 3.6425 | *0.5734* | 1.3280 | 2.0445 | 0.3775 | O60271;A0A087X2D8;H0YBE9 | SPAG9 |
| -0.2179 | 0.6256 | *0.4635* | 0.2904 | 0.4476 | 0.3779 | E7EU13;Q96P48 | ARAP1 |
| -0.9257 | 0.3747 | *-5.9647* | -2.1719 | 3.3484 | 0.3780 | E9PIZ2;Q8N3Y3 | LARGE2 |

| L100/ CTRL1 | L100/ CTRL2 | L100/ CTRL3 | Mean | SD | T-test p-value | Accession | Gene Symbol |
|---|---|---|---|---|---|---|---|
| -0.9460 | -2.0934 | *0.5070* | -0.8441 | 1.3032 | 0.3785 | Q9UKK3 | PARP4 |
| -0.2778 | 0.1400 | *-1.2786* | -0.4721 | 0.7290 | 0.3785 | C9JAV2;H7C216; Q6PH85;C9J2J1 | DCUN1D2 |
| 0.5367 | 0.1301 | *-0.0679* | 0.1996 | 0.3082 | 0.3785 | K7EM11;O14908 | GIPC1 |
| 0.1113 | 1.5055 | *0.0068* | 0.5412 | 0.8368 | 0.3791 | Q13905 | RAPGEF1 |
| -2.0706 | -0.8284 | *0.4519* | -0.8157 | 1.2613 | 0.3791 | O75489 | NDUFS3 |
| -0.3425 | -0.0334 | *0.0058* | -0.1234 | 0.1908 | 0.3792 | P0CJ79 | ZNF888 |
| -0.2402 | 0.1186 | *-1.1923* | -0.4380 | 0.6774 | 0.3792 | P05204 | HMGN2 |
| 0.4622 | -1.0512 | *-6.4189* | -2.3360 | 3.6160 | 0.3795 | Q4J6C6;H7C0M0 | PREPL |
| -0.0235 | 0.1346 | *1.3966* | 0.5026 | 0.7783 | 0.3797 | Q6ZU52;E9PQS0 | KIAA0408 |
| -0.0587 | 0.2012 | *1.7466* | 0.6297 | 0.9759 | 0.3800 | P02768;A0A0C4 DGB6;B7WNR0; H0YA55;C9JKR2; D6RHD5;A0A087 WWT3;H7C013; Q9UMX5 | ALB |
| -0.5638 | 0.2423 | *-0.5989* | -0.3068 | 0.4759 | 0.3803 | P04181 | OAT |
| 0.0463 | -0.2811 | *-3.0585* | -1.0978 | 1.7059 | 0.3810 | A0A0D9SGJ6;C9 JFZ1;C9JW66;J3 KPK1;J3KQV8;O 43426;C9J1Z6 | SYNJ1 |
| 0.0735 | -0.2755 | *-2.5763* | -0.9261 | 1.4397 | 0.3811 | Q6ZS81 | WDFY4 |
| 0.0860 | -0.4257 | *-4.4226* | -1.5874 | 2.4686 | 0.3813 | P49815;A0A2R8 Y7C8;A0A2R8YD R3;A0A2R8YGD 6;A0A2R8Y5F1;A 0A2R8Y7X5;A0A 2R8YDZ2;A0A2R 8YGU4;H3BMQ0; X5D2U8;A0A2R8 YEJ8;A0A2R8Y6 C9;H3BQY7 | TSC2 |
| 2.6851 | 3.0296 | *-1.1931* | 1.5072 | 2.3449 | 0.3815 | Q9UNA4;X6R2I3; J3KSW2;J3KQ09 ;J3QR36;J3KTN3 | POLI |
| -0.6978 | 0.3536 | *-3.6579* | -1.3341 | 2.0800 | 0.3823 | Q13362;Q96B13; H0YJ75 | PPP2R5C |
| 1.1834 | 0.5294 | *-0.2919* | 0.4736 | 0.7392 | 0.3827 | A0A0G2JR66;Q2 KJY2;B7WPD9 | KIF26B |
| -0.2276 | 0.4155 | *0.8610* | 0.3496 | 0.5473 | 0.3838 | Q13813;A0A0D9 SFF6;A0A0D9SF H4;A0A1B0GTB7 | SPTAN1 |
| 0.2972 | 0.6573 | *-0.1647* | 0.2632 | 0.4121 | 0.3838 | O60293 | ZFC3H1 |
| 0.3821 | -0.6734 | *-2.6720* | -0.9878 | 1.5511 | 0.3850 | P23381;H0YJP3 | WARS |
| -0.2493 | 0.7356 | *0.4938* | 0.3267 | 0.5133 | 0.3852 | Q6AI08;K7EIX2; K7EKW7;K7ELR 8;K7ESP5 | HEATR6 |
| 0.1907 | -2.1999 | *-0.3755* | -0.7949 | 1.2493 | 0.3853 | Q9UHD8;K7EL40 ;K7EIE4;K7EK18; K7EQD7;K7ER52 ;K7ELJ9;K7ER14 ;K7EIR4;K7EJ51; K7EJZ2;K7EKN4; K7EN52;K7ENQ5 | SEPT9 |

| L100/ CTRL1 | L100/ CTRL2 | L100/ CTRL3 | Mean | SD | T-test p-value | Accession | Gene Symbol |
|---|---|---|---|---|---|---|---|
| | | | | | | ;K7EJL9;K7ERG 1 | |
| 0.0039 | 0.0698 | *1.1354* | 0.4031 | 0.6351 | 0.3863 | H0Y684 | OBSL1 |
| 0.0512 | -0.0049 | *0.7164* | 0.2542 | 0.4012 | 0.3869 | Q8N2N9;A0A087 X1R3 | ANKRD36B |
| -0.4860 | 0.2843 | *-1.9780* | -0.7266 | 1.1501 | 0.3881 | Q8NFI3;F8W925 | ENGASE |
| 0.1396 | -0.0018 | *2.2246* | 0.7875 | 1.2466 | 0.3881 | P04080;A0A1W2 PS52 | CSTB |
| -0.6841 | -1.3814 | *0.3823* | -0.5611 | 0.8883 | 0.3881 | Q9NR09;H7C094 ;H7C3P0 | BIRC6 |
| 0.5510 | 1.6550 | *-0.3255* | 0.6268 | 0.9924 | 0.3881 | O15021;E7EX28; H7C0U3;H7C2V7 ;H7C3S4 | MAST4 |
| -0.1802 | 1.0702 | *0.3041* | 0.3980 | 0.6305 | 0.3883 | F8VXY3;P00973; H0YI20;H0YIB8 | OAS1 |
| -0.3665 | -0.9585 | *0.2152* | -0.3699 | 0.5869 | 0.3889 | P17812 | CTPS1 |
| -0.2341 | 0.8741 | *0.4124* | 0.3508 | 0.5567 | 0.3890 | P62745 | RHOB |
| -0.1264 | 1.6299 | *0.2454* | 0.5830 | 0.9256 | 0.3892 | Q96FW1;F5GYJ8 ;F5GYN4;J3KR4 4;F5H6Q1;F5H3F 0 | OTUB1 |
| 0.7445 | -0.4473 | *2.5495* | 0.9489 | 1.5088 | 0.3898 | P14136;A0A1W2 PR46;A0A1X7SB R3;K7EMP8;A0A 1X7SCE1;K7EJU 1;A0A1W2PRT3; K7EKH9;K7ELP4 ;B4DIR1 | GFAP |
| 0.4917 | -0.2023 | *4.5368* | 1.6088 | 2.5594 | 0.3900 | A6H900;B1AM31 ;Q5T1B0;D6RDY 4;D6REE1 | AXDND1 |
| -2.1134 | 0.2145 | *-0.3810* | -0.7599 | 1.2094 | 0.3901 | Q5SSJ5;B0QZK4 ;X6RGJ2;Q5SW C8 | HP1BP3 |
| 0.0020 | -0.9405 | *-0.0568* | -0.3317 | 0.5280 | 0.3902 | Q96JE7;E9PK14; H0YE70 | SEC16B |
| -0.2635 | 0.0676 | *-3.2693* | -1.1551 | 1.8384 | 0.3902 | O94813;X6R3P0; E9PCX4 | SLIT2 |
| -0.0448 | 0.0112 | *-0.5654* | -0.1997 | 0.3180 | 0.3904 | Q8TF40 | FNIP1 |
| -2.2171 | 1.2974 | *-5.1921* | -2.0373 | 3.2485 | 0.3909 | G5EA30;Q92879; E9PKA1;E9PKU1 ;E9PQK4;E9PSH 0;F5H0D8;F5H4Y 5 | CELF1 |
| -0.1866 | 0.3954 | *0.4631* | 0.2240 | 0.3571 | 0.3909 | O60282;A0A0G2 JMZ6;C9JWB9 | KIF5C |
| 0.2596 | -0.6090 | *-0.5772* | -0.3088 | 0.4925 | 0.3909 | Q9UHB9 | SRP68 |
| -0.0752 | -0.9624 | *0.0191* | -0.3395 | 0.5415 | 0.3910 | O60336 | MAPKBP1 |
| 6.7383 | 1.0997 | *-0.6107* | 2.4091 | 3.8455 | 0.3913 | C9JIF3;H7C3D5; Q8NC44 | RETREG2 |
| 0.4209 | 0.6198 | *-0.2182* | 0.2742 | 0.4378 | 0.3914 | Q08043;A0A087 WSZ2;D6RH00 | ACTN3 |
| 0.5887 | -0.3596 | *2.0859* | 0.7717 | 1.2330 | 0.3916 | Q8NCX0;E9PCV 3 | CCDC150 |

| L100/ CTRL1 | L100/ CTRL2 | L100/ CTRL3 | Mean | SD | T-test p-value | Accession | Gene Symbol |
|---|---|---|---|---|---|---|---|
| -1.3964 | 0.6036 | *-1.3472* | -0.7133 | 1.1407 | 0.3920 | O43488;H3BLU7;H7C5H7;Q8NHP1;H7C4Q7 | AKR7A2 |
| -0.6939 | 0.0100 | *-0.0484* | -0.2441 | 0.3906 | 0.3922 | P50454;E9PPV6;E9PR70;E9PK86;E9PMI5;E9PNX1;E9PIG2;E9PRS3;E9PJH8;E9PQ34;E9PKH2;E9PLA6 | SERPINH1 |
| 1.6927 | 0.2651 | *-0.1482* | 0.6032 | 0.9659 | 0.3925 | Q9ULT8;A0A087X2H1;H0YJP0 | HECTD1 |
| -0.7796 | 3.3368 | *1.2969* | 1.2847 | 2.0583 | 0.3927 | Q6P2I3 | FAHD2B |
| -0.4673 | 0.7770 | *1.9577* | 0.7558 | 1.2126 | 0.3932 | E9PM04;Q9BTE7;E9PLH8;E9PLS2;E9PM78;E9PQV9;H0YD80 | DCUN1D5 |
| -2.9091 | 1.0966 | *-2.1715* | -1.3280 | 2.1319 | 0.3935 | P49916;K7ERZ5;K7EJR4;K7EQB6 | LIG3 |
| -0.1004 | 0.1734 | *0.3557* | 0.1429 | 0.2295 | 0.3937 | P61081;M0QX69 | UBE2M |
| 0.0987 | -0.5591 | *-8.8404* | -3.1003 | 4.9819 | 0.3938 | Q7RTS9 | DYM |
| -1.0346 | 0.6400 | *-3.1120* | -1.1689 | 1.8796 | 0.3941 | F8VU56;P23467;F8VSD5;Q6ZR19;H0YHE8 | PTPRB |
| -0.0188 | 0.0375 | *0.3364* | 0.1184 | 0.1909 | 0.3953 | O00311;B1AMW7 | CDC7 |
| -0.3707 | 0.5900 | *1.8905* | 0.7033 | 1.1349 | 0.3954 | H7BYX7;Q6P1J6 | PLB1 |
| 0.2514 | -0.4169 | *-0.9696* | -0.3784 | 0.6114 | 0.3960 | Q02641 | CACNB1 |
| 3.6821 | -1.7853 | *4.3407* | 2.0792 | 3.3628 | 0.3963 | Q86XK2 | FBXO11 |
| 0.2754 | -0.5431 | *-5.0296* | -1.7657 | 2.8560 | 0.3963 | Q92556 | ELMO1 |
| 0.4951 | -0.7757 | *-3.4575* | -1.2461 | 2.0179 | 0.3968 | Q9BVV6;A0A087WYM5 | KIAA0586 |
| 0.6183 | -1.2745 | *-1.4963* | -0.7175 | 1.1622 | 0.3969 | O00222;A0A0A0MT06 | GRM8 |
| -0.4720 | -0.3626 | *0.1839* | -0.2169 | 0.3514 | 0.3969 | H0YK42;Q13596 | SNX1 |
| 0.0749 | -0.2615 | *-0.1267* | -0.1044 | 0.1693 | 0.3973 | Q8TF20 | ZNF721 |
| -0.3474 | 1.0815 | *0.6124* | 0.4488 | 0.7284 | 0.3976 | Q8NFP7;Q96G61;A0A024RBG1;A0A0C4DGJ4;F8VRL4;F8VRR0 | NUDT10 |
| -4.2969 | 0.6649 | *-1.0277* | -1.5533 | 2.5223 | 0.3979 | P20340;F5H3K7 | RAB6A |
| -0.3476 | -0.7528 | *0.2090* | -0.2971 | 0.4829 | 0.3982 | P31948;F5H783;F5GXD8;H0YGI8;G5EA25;Q9P217 | STIP1 |
| 3.1418 | 0.7244 | *-0.4730* | 1.1311 | 1.8414 | 0.3988 | Q13564;H3BQW6;J3KRK3 | NAE1 |
| 0.2143 | -0.5165 | *-0.4359* | -0.2460 | 0.4007 | 0.3990 | P50502;Q3KNR6;Q8NFI4;H7C3I1;F6VDH7;F8WAQ7 | ST13 |
| -0.4332 | 0.2087 | *-0.4849* | -0.2365 | 0.3864 | 0.4002 | H3BS70;P42126;Q96DC0 | ECI1 |
| 0.3914 | -0.2492 | *2.6264* | 0.9229 | 1.5097 | 0.4007 | Q8TET4;H3BN99 | GANC |

| L100/ CTRL1 | L100/ CTRL2 | L100/ CTRL3 | Mean | SD | T-test p-value | Accession | Gene Symbol |
|---|---|---|---|---|---|---|---|
| -1.2453 | 0.0030 | *-0.0508* | -0.4311 | 0.7057 | 0.4010 | Q4G0P3;F8WD03;J3QQJ7;J3QL79;A0A087WVK9;H0Y7Y5 | HYDIN |
| 0.8961 | -0.5507 | *1.9777* | 0.7744 | 1.2686 | 0.4012 | R4GMW8 | BIVM-ERCC5 |
| 0.1703 | -0.2562 | *-1.3417* | -0.4758 | 0.7796 | 0.4012 | A0A1W2PPT5;C9J2Y9;P30876;C9J4M6 | POLR2B |
| -0.1702 | 0.2983 | *0.5040* | 0.2107 | 0.3456 | 0.4017 | A0A1B0GXI6;A0A2R8YD95;A0A2R8YE10;A0A2R8YF01;Q9NRW7;Q5T4Q0;A0A087WU65;B7Z7G7;A0A2R8Y7W9 | VPS45 |
| -1.0164 | 0.6776 | *-3.5219* | -1.2869 | 2.1128 | 0.4021 | E9PBD5;G3V1X8 | ADAMTS20 |
| -3.5035 | 0.0364 | *-0.1624* | -1.2098 | 1.9889 | 0.4026 | P52179;A8MX12;J3KRK2 | MYOM1 |
| -0.1419 | -0.6525 | *0.0965* | -0.2326 | 0.3827 | 0.4028 | A0A0G2JRV3;G5E975;Q12824;A0A0G2JSE9;B5MCL5;C9JTA6 | SMARCB1 |
| -1.1852 | 0.7339 | *-10.6369* | -3.6961 | 6.0871 | 0.4033 | F2Z2K5;Q96N16 | JAKMIP1 |
| -0.1859 | 0.0121 | *-5.2096* | -1.7945 | 2.9592 | 0.4038 | P43686 | PSMC4 |
| 0.0602 | -1.2829 | *-0.1078* | -0.4435 | 0.7318 | 0.4040 | Q8N8L6 | ARL10 |
| 12.7293 | 9.9163 | *-5.1622* | 5.8278 | 9.6210 | 0.4042 | Q6P179 | ERAP2 |
| -2.7510 | 1.6204 | *-4.8918* | -2.0075 | 3.3191 | 0.4048 | Q8NA31;J3KSG9 | TERB1 |
| -0.1600 | 0.3594 | *0.3297* | 0.1763 | 0.2917 | 0.4049 | Q96GC6;A0A0A0MR47;M0QXW4;M0QY30 | ZNF274 |
| -1.1439 | -1.3857 | *0.5830* | -0.6489 | 1.0736 | 0.4051 | Q99832;F8WAM2;F8WBP8;A0A0D9SG95 | CCT7 |
| -0.0888 | 0.1754 | *0.2087* | 0.0985 | 0.1630 | 0.4053 | P14866;M0QXS5;B4DVF8;M0R1W6;M0QYL7;M0R076 | HNRNPL |
| -0.3157 | 0.2137 | *-2.3496* | -0.8172 | 1.3532 | 0.4054 | Q9NWH9;H7BXE3;H0YL55;H0YLE6;H0YMR6;H0YLW7;H0YMW8;H0YKU6 | SLTM |
| -0.3006 | -1.5511 | *0.2105* | -0.5471 | 0.9063 | 0.4055 | Q12789 | GTF3C1 |
| -0.2140 | 0.7959 | *0.3335* | 0.3051 | 0.5055 | 0.4056 | O00429;G8JLD5;F8VZ52;F8VUJ9;F8W1W3;B4DPZ9;B4DDQ3;F8VR28;F8VYL3;H0Y7D7;H0YI79;O76038 | DNM1L |
| 0.2023 | -0.3472 | *-0.5900* | -0.2450 | 0.4059 | 0.4056 | Q9UER7 | DAXX |
| 0.7877 | 2.2345 | *-0.5243* | 0.8327 | 1.3799 | 0.4057 | P31153 | MAT2A |

| L100/ CTRL1 | L100/ CTRL2 | L100/ CTRL3 | Mean | SD | T-test p-value | Accession | Gene Symbol |
|---|---|---|---|---|---|---|---|
| -0.0841 | -0.0096 | *-3.1969* | -1.0969 | 1.8191 | 0.4059 | A0A2R8Y549;Q8NI35;A0A0U1RQT2;A0A2R8Y5I3;B4DE90 | PATJ |
| -0.3721 | 0.5314 | *2.1855* | 0.7816 | 1.2970 | 0.4062 | F8VQS4;F8W1M4 | SLC16A7 |
| 0.2810 | -0.1979 | *1.2623* | 0.4485 | 0.7443 | 0.4062 | Q8IVM0 | CCDC50 |
| 2.4483 | -0.8690 | *1.5106* | 1.0300 | 1.7101 | 0.4064 | Q14511 | NEDD9 |
| -1.4783 | 0.3875 | *-0.5945* | -0.5618 | 0.9333 | 0.4066 | P42338;H0Y871;H7C565 | PIK3CB |
| -0.1960 | 0.2764 | *1.4670* | 0.5158 | 0.8570 | 0.4066 | Q92896;H3BM42;H3BQU9;H3BQT1;H3BS09;H3BSW9;J3KRR7;J3KTF7;J3QLS4;Q9NQC7 | GLG1 |
| -0.2940 | 2.8675 | *0.4179* | 0.9971 | 1.6584 | 0.4070 | P63096;C9JPP4 | GNAI1 |
| -0.6270 | 0.8910 | *6.1494* | 2.1378 | 3.5561 | 0.4071 | Q8WVE0 | EEF1AKMT1 |
| 3.3986 | 2.5395 | *-1.3626* | 1.5252 | 2.5375 | 0.4072 | E9PKF6;H7BXH2;Q5H9R7;E9PKG4;E9PNN8;E9PQP7;H0YEN2 | PPP6R3 |
| 0.2317 | -0.3303 | *-1.2288* | -0.4424 | 0.7367 | 0.4074 | Q9H2D6;F6WYE2;H0Y5J9;F6TR96 | TRIOBP |
| -0.4532 | 0.3210 | *-1.8477* | -0.6600 | 1.0991 | 0.4075 | P20337 | RAB3B |
| -0.3055 | -1.6850 | *0.2194* | -0.5904 | 0.9836 | 0.4077 | O14964;I3L1P5;I3L165;I3L2H4 | HGS |
| 0.2597 | 1.4791 | *-0.1865* | 0.5175 | 0.8622 | 0.4077 | Q9H0B6;A8MZ87;E9PQ02;C9JHT2;E9PI24;E9PM83;E9PP09;A8MX29 | KLC2 |
| 0.4982 | -0.7069 | *-2.5480* | -0.9189 | 1.5341 | 0.4085 | Q5TB80 | CEP162 |
| -0.0514 | -0.8622 | *0.0282* | -0.2951 | 0.4927 | 0.4085 | Q9H0K6 | PUS7L |
| -0.1220 | -0.1075 | *-10.1732* | -3.4676 | 5.8073 | 0.4097 | Q9BR84;A0A0A0MTT2;S4R3U0;A0A0A0MTS4;K7EKU6;K7EN17;K7EQI4;K7ERK9;S4R342;S4R3A1 | ZNF559 |
| 0.3020 | -0.1798 | *0.5105* | 0.2109 | 0.3541 | 0.4106 | Q6PKG0;A0A0B4J210 | LARP1 |
| 0.3779 | -0.5258 | *-1.9205* | -0.6895 | 1.1579 | 0.4108 | O43396;K7EML9;K7ER96;K7EKG2;K7EPB7 | TXNL1 |
| 0.0241 | -0.2067 | *-0.0320* | -0.0716 | 0.1203 | 0.4113 | Q6ZTR5;A0A0D9SEI5 | CFAP47 |
| 0.8041 | 4.8055 | *-0.6065* | 1.6677 | 2.8074 | 0.4117 | E9PJ95;Q9P000 | COMMD9 |
| -0.0402 | -0.0210 | *-3.2254* | -1.0955 | 1.8445 | 0.4118 | Q3ZCQ8;M0R047 | TIMM50 |
| -0.3890 | -0.3095 | *0.1651* | -0.1778 | 0.2996 | 0.4120 | P53990;H3BMU1;H3BUI0;H3BQF7;F5GXM3;H3BPP | IST1 |

| L100/ CTRL1 | L100/ CTRL2 | L100/ CTRL3 | Mean | SD | T-test p-value | Accession | Gene Symbol |
|---|---|---|---|---|---|---|---|
| | | | | | | 6;H3BRE2;J3KR23 | |
| -0.7402 | 0.3599 | *-0.7613* | -0.3805 | 0.6413 | 0.4121 | Q96AX9;D6RAZ0;E9PD12;F2Z2L2;D6RED3;D6RE96;D6RFJ2 | MIB2 |
| -1.4627 | 0.2349 | *-0.3135* | -0.5138 | 0.8663 | 0.4123 | P60900;G3V5Z7;G3V295;G3V3I1;G3V3U4;G3V2S7;G3V4S5;H0YJC4 | PSMA6 |
| -0.8286 | 1.5261 | *1.9932* | 0.8969 | 1.5125 | 0.4124 | O95395;H0YM40;H0YMW7;H0YNA3 | GCNT3 |
| -0.0665 | 0.0063 | *-3.3984* | -1.1529 | 1.9450 | 0.4125 | Q9Y4G6 | TLN2 |
| -0.2302 | 0.3099 | *1.2748* | 0.4515 | 0.7624 | 0.4129 | Q96BW9;A0A0G2JQ92 | TAMM41 |
| 1.2021 | 1.0071 | *-0.5269* | 0.5608 | 0.9470 | 0.4129 | Q9NQR4;F8WF70 | NIT2 |
| -0.5361 | 0.1684 | *-0.2631* | -0.2102 | 0.3552 | 0.4131 | P46926;D6R9P4;D6RAY7;D6RB13;D6RFF8;D6R917 | GNPDA1 |
| 0.0003 | 0.2038 | *0.0029* | 0.0690 | 0.1167 | 0.4136 | O00487 | PSMD14 |
| -0.0121 | -0.2727 | *0.0077* | -0.0924 | 0.1565 | 0.4140 | Q02252;G3V4Z4 | ALDH6A1 |
| -0.0279 | 0.0519 | *1.5357* | 0.5199 | 0.8806 | 0.4140 | A6NGW2;Q7RTU9 | STRCP1 |
| 0.1995 | -0.2668 | *-1.0680* | -0.3784 | 0.6411 | 0.4141 | Q8IXJ9;A0A2R8Y5U1;Q76L82 | ASXL1 |
| 1.1611 | -0.4213 | *0.7007* | 0.4802 | 0.8139 | 0.4143 | Q13488 | TCIRG1 |
| -0.3503 | 0.2598 | *-4.9510* | -1.6805 | 2.8487 | 0.4144 | P52788;H7C2R7 | SMS |
| -0.0098 | -0.6423 | *0.0006* | -0.2172 | 0.3682 | 0.4145 | F5H658;Q14562;K7EQH7 | DHX8 |
| -0.1749 | 0.2363 | *0.8260* | 0.2958 | 0.5031 | 0.4157 | Q12904;D6R937 | AIMP1 |
| -0.5584 | -2.6933 | *0.4337* | -0.9393 | 1.5980 | 0.4157 | Q7KZN9 | COX15 |
| -0.4830 | 0.7722 | *1.3948* | 0.5613 | 0.9565 | 0.4164 | P62136;E9PMD7;F5H037;F5H1L6;F8W1A0;F8WE71;H0Y3Y6 | PPP1CA |
| -0.4341 | 0.7154 | *1.1845* | 0.4886 | 0.8328 | 0.4165 | P12271 | RLBP1 |
| -0.3905 | -0.4995 | *0.2139* | -0.2254 | 0.3843 | 0.4166 | Q13442;F8WBW6 | PDAP1 |
| 0.5204 | -0.4166 | *2.9232* | 1.0090 | 1.7227 | 0.4171 | Q5BJF6;S4R411;S4R462 | ODF2 |
| -1.3056 | -1.6525 | *0.7135* | -0.7482 | 1.2777 | 0.4172 | J3KNN5;Q9UJV9 | DDX41 |
| -0.3555 | -0.1886 | *0.1202* | -0.1413 | 0.2414 | 0.4173 | Q96R06;J3KTQ0;K7ELC8;J3QS12 | SPAG5 |
| -0.7407 | 0.4489 | *-1.2068* | -0.4995 | 0.8538 | 0.4175 | O43242;H0YGV8 | PSMD3 |
| -0.4771 | 0.7482 | *1.4048* | 0.5586 | 0.9552 | 0.4177 | P62140;E7ETD8;C9JP48;C9J9S3 | PPP1CB |
| -0.4771 | 0.7482 | *1.4048* | 0.5586 | 0.9552 | 0.4177 | P36873;F8W0W8;F8VYE8;F8VR82;A0A087WYY5 | PPP1CC |
| 1.1964 | -1.6832 | *-4.5628* | -1.6832 | 2.8796 | 0.4179 | A0A0U1RRB9;F2Z2V1;Q96C11 | FGGY |

| L100/ CTRL1 | L100/ CTRL2 | L100/ CTRL3 | Mean | SD | T-test p-value | Accession | Gene Symbol |
|---|---|---|---|---|---|---|---|
| 0.0521 | 0.6909 | *-0.0432* | 0.2333 | 0.3992 | 0.4180 | Q13907 | IDI1 |
| -0.2464 | -0.2279 | *0.1163* | -0.1193 | 0.2042 | 0.4180 | P08670;B0YJC4;B0YJC5;A0A1B0GTT5;A0A1B0GVG8;Q8N5S1;A0A087X106;O43790;P12035;P78385;Q14533;U3KPR1 | VIM |
| -0.4593 | 2.0669 | *0.6131* | 0.7402 | 1.2679 | 0.4183 | Q5THN1;Q9Y5Z4 | HEBP2 |
| 3.8822 | 4.0644 | *-1.9548* | 1.9972 | 3.4238 | 0.4187 | S4R3L9 | KCTD10 |
| 0.3298 | -4.6013 | *-0.3885* | -1.5533 | 2.6639 | 0.4188 | M0QZ12;Q96CP6 | GRAMD1A |
| 0.6533 | 0.6126 | *-0.3115* | 0.3181 | 0.5456 | 0.4189 | Q5T200;E5RHF4;F5H3S6 | ZC3H13 |
| -0.1496 | 0.2273 | *0.4597* | 0.1791 | 0.3075 | 0.4192 | P16403;Q02539;P22492 | HIST1H1C |
| -0.2520 | 1.3605 | *0.3187* | 0.4757 | 0.8177 | 0.4197 | Q14108;A0A1W2PQB7;A0A1W2PQR6;A0A1W2PRS1;A0A1W2PS43;A0A1W2PPU6;A0A1W2PPX5;A0A1W2PRF6;A0A1W2PSE4;A0A1W2PNX7;A0A1W2PS70;D6RDG0;A0A1W2PPX6;A0A1W2PQL5 | SCARB2 |
| 0.1838 | -0.4842 | *-0.3023* | -0.2009 | 0.3453 | 0.4197 | P54136;E5RJM9 | RARS |
| 0.1543 | -0.1249 | *5.0269* | 1.6855 | 2.8972 | 0.4197 | Q92508 | PIEZO1 |
| -0.0219 | 1.2966 | *0.0288* | 0.4345 | 0.7471 | 0.4198 | Q6TFL3 | CCDC171 |
| 0.1789 | -0.5799 | *-0.2641* | -0.2217 | 0.3812 | 0.4199 | O00139;D6R9M0 | KIF2A |
| 0.0096 | -0.0138 | *-1.0390* | -0.3478 | 0.5988 | 0.4204 | Q6YP21 | KYAT3 |
| -0.0266 | 0.0227 | *-1.0659* | -0.3566 | 0.6147 | 0.4208 | O14936;A0A2R8YE77;A0A2U3TZM1;A0A2U3TZM4;A0A2U3TZN6;Q5JS74;A0A2R8Y3B3;A0A2R8YEJ7;Q5JS72;Q5JS79;A0A2R8Y4K4;A0A2R8YGH2;A0A2R8Y472;A0A2R8Y6D8;A0A2R8YEK3;A0A2R8Y6F8;A0A2R8YFN5 | CASK |
| 0.2784 | 0.9433 | *-0.2125* | 0.3364 | 0.5801 | 0.4210 | Q00796;H0YKB3 | SORD |
| -0.0187 | 0.0215 | *0.8565* | 0.2864 | 0.4941 | 0.4211 | P35612;C9J080;C9JTM0;A0A1C7CYY0 | ADD2 |
| 0.1409 | -0.1128 | *0.6005* | 0.2095 | 0.3615 | 0.4212 | P09651;F8W6I7;A0A2R8Y4L2;H0YH80;F8VTQ5;F8VYN5;F8W646 | HNRNPA1 |

| L100/ CTRL1 | L100/ CTRL2 | L100/ CTRL3 | Mean | SD | T-test p-value | Accession | Gene Symbol |
|---|---|---|---|---|---|---|---|
| -0.0415 | 0.0459 | *0.5994* | 0.2013 | 0.3476 | 0.4215 | A0A0A0MRX1;Q12926 | ELAVL2 |
| -2.0639 | 0.9293 | *-1.7034* | -0.9460 | 1.6340 | 0.4216 | Q02978;I3L1P8 | SLC25A11 |
| -0.0654 | 0.0578 | *-4.8639* | -1.6238 | 2.8066 | 0.4218 | J3KN36;P69849;Q5JPE7 | NOMO3 |
| 0.2527 | 2.2144 | *-0.2247* | 0.7475 | 1.2926 | 0.4220 | Q14929 | ZNF169 |
| 0.0055 | 0.0849 | *-0.0052* | 0.0284 | 0.0492 | 0.4223 | P22695;H3BRG4;H3BSJ9;H3BP04;H3BUE4;H3BUI9;A0A087WVZ4 | UQCRC2 |
| 0.4181 | -0.4817 | *-3.1734* | -1.0790 | 1.8687 | 0.4226 | Q9HBL0;E9PF55;E9PGF5;E7ERH1;E7EMG1;C9J8K5;C9JFT7;C9JI43;H7C3Z4 | TNS1 |
| -0.1453 | 0.1540 | *-9.8101* | -3.2671 | 5.6684 | 0.4233 | Q6ZMV9;H0Y718 | KIF6 |
| 0.0170 | -0.0183 | *1.0864* | 0.3617 | 0.6279 | 0.4235 | O00264 | PGRMC1 |
| 0.0590 | -0.0616 | *2.1547* | 0.7174 | 1.2462 | 0.4238 | Q8NHW5 | RPLP0P6 |
| -0.4104 | -1.7930 | *0.3389* | -0.6215 | 1.0815 | 0.4244 | Q99623;J3KPX7;F5GY37;F5GWA7;F5H0C5;F5H2D2 | PHB2 |
| -0.2712 | 1.4502 | *0.3267* | 0.5019 | 0.8740 | 0.4247 | P56537;B7ZBH1;A0A0U1RQV5 | EIF6 |
| -0.3728 | 0.6452 | *0.8652* | 0.3792 | 0.6605 | 0.4248 | P35580;E7ERA5;A0A087WV07;A0A2R8YFN9;C9J269;C9J911;D6RCE7;D6RED9;F5H1X6;H0YE44;H3BME1;H3BVB1;H7BZ91;H7C2E2;H7C4D8;K7EM46;O00463;Q13402;Q5VXN0;Q6AHZ1;Q8IV76;Q92835;Q9H7B2;Q9UN88;Q9Y283 | MYH10 |
| 0.3188 | -0.2581 | *1.2384* | 0.4330 | 0.7547 | 0.4251 | O15083;H7C4G9 | ERC2 |
| -0.5796 | -0.4246 | *0.2480* | -0.2521 | 0.4399 | 0.4256 | Q9NX40;D6RBN5;D6RDK6;D6R918;D6RA54;D6RBC5;D6RC55;D6RDI5;D6RF07;D6RG39;D6RIT9 | OCIAD1 |
| -1.2778 | 0.7675 | *-1.8620* | -0.7908 | 1.3807 | 0.4258 | Q5VZ46;A0A1W2PPL5 | KIAA1614 |
| 0.4165 | -0.4692 | *-2.8208* | -0.9578 | 1.6731 | 0.4259 | O75131;A0A087WYQ3;E5RHZ0;A0A087WXR6;H0YB26;A0A087WUS8;E5RFT7;Q9HCH3;Q9UBL6 | CPNE3 |
| 0.5339 | -0.5059 | *-8.8275* | -2.9332 | 5.1310 | 0.4265 | Q9H313;A0A0G2JPK6;E7ET67 | TTYH1 |
| -0.2513 | 1.8974 | *0.2716* | 0.6392 | 1.1205 | 0.4273 | P00505 | GOT2 |

| L100/ CTRL1 | L100/ CTRL2 | L100/ CTRL3 | Mean | SD | T-test p-value | Accession | Gene Symbol |
|---|---|---|---|---|---|---|---|
| 0.8665 | 2.8997 | *-0.6911* | 1.0250 | 1.8007 | 0.4281 | P46939 | UTRN |
| -0.2022 | 0.2775 | *0.6678* | 0.2477 | 0.4358 | 0.4286 | P31943;G8JLB6;E9PCY7;D6RBM0;D6RIT2;D6RIU0;D6R9T0;D6RAM1;D6RDU3;D6RIH9;D6RJ04;E5RGV0;H0YB39;E7EQJ0;E5RGH4;D6RDL0;H0YAQ2;H0YBD7;D6RF17;D6R9D3;H0YBG7;E5RJ94;E7EN40 | HNRNPH1 |
| 1.1499 | -2.3199 | *-2.1699* | -1.1133 | 1.9614 | 0.4292 | E9PPP6 | RRP8 |
| -0.3677 | -0.2742 | *0.1617* | -0.1601 | 0.2825 | 0.4299 | Q9H9B1;A0A1B0GV09;A0A1B0GU48;A0A0D9SEQ1;A0A1B0GUD1;A0A1B0GVZ8;A0A0D9SFX4;A0A1B0GW79;A0A0C4DGF8;A0A1B0GV89;A0A0D9SEY2;A0A0D9SFD7;A0A0D9SFM6;A0A1B0GTP4;A0A1B0GWF6;A0A0D9SER3;A0A0D9SFG7;A0A0D9SFS4 | EHMT1 |
| -0.5345 | 0.5038 | *-3.4728* | -1.1678 | 2.0625 | 0.4302 | P61163;R4GMT0;A0A1B0GVS3 | ACTR1A |
| -0.0077 | -3.0760 | *0.0518* | -1.0107 | 1.7889 | 0.4310 | A0A2R8YEQ5;Q5QGS0;H7C2N8 | NEXMIF |
| 0.0320 | 0.0117 | *-2.9799* | -0.9788 | 1.7331 | 0.4311 | Q9NUP9;G3V1D4;J3KN23 | LIN7C |
| -0.1927 | 3.0728 | *0.1537* | 1.0113 | 1.7937 | 0.4318 | P46100;A0A096LNW1;A0A096LNL9;A0A087WWG0;A0A096LNL6;A0A096LNN3 | ATRX |
| -0.1911 | 0.2516 | *0.6540* | 0.2381 | 0.4227 | 0.4321 | Q13043;F5H5B4;Q13188;A0A087WZ06;E5RFQ9;E5RIM6;Q8NBU1 | STK4 |
| 0.4791 | 4.0630 | *-0.4893* | 1.3509 | 2.3981 | 0.4321 | Q9H9E3;J3KNI1;A0A0A0MS45;H3BMV9;J3KRB5;E9PRT5;H3BSD2;J3QLW1 | COG4 |
| 0.1361 | -0.1455 | *1.3840* | 0.4582 | 0.8140 | 0.4324 | Q9BUJ2;A0A0A0MRA5;B7Z4B8;M0R3F1;M0QYZ0;M0QYI8;M0QYM5;M0R0K8 | HNRNPUL1 |
| -1.2883 | -0.8700 | *0.5407* | -0.5392 | 0.9583 | 0.4326 | H8Y6P7;P0CAP1;H3BTD5 | GCOM1 |

| L100/ CTRL1 | L100/ CTRL2 | L100/ CTRL3 | Mean | SD | T-test p-value | Accession | Gene Symbol |
|---|---|---|---|---|---|---|---|
| 0.1830 | -0.1798 | *1.2628* | 0.4220 | 0.7504 | 0.4328 | P25789;H0YL69;H0YMZ1;H0YMA1;H0YMI6;H0YN18;H0YLC2;H0YLS6;H0YKT8 | PSMA4 |
| 0.8207 | 1.2586 | *-0.5184* | 0.5203 | 0.9258 | 0.4330 | Q9P2R7;A0A2R8Y6Y7;A0A2R8YDQ9;Q5T9Q5;A0A2R8Y5P6;A0A2R8Y6E6;A0A0U1RQF8;A0A0U1RQL1;A0A0U1RRI1;Q5T9Q8 | SUCLA2 |
| 0.5860 | -0.5797 | *-4.3901* | -1.4613 | 2.6025 | 0.4334 | A0A1W2PPZ5;P23193;A0A1W2PRL9 | TCEA1 |
| -0.2484 | -0.8653 | *0.2075* | -0.3021 | 0.5384 | 0.4337 | P50993;B1AKY9;H0Y7C1 | ATP1A2 |
| 0.3550 | 0.5392 | *-0.2241* | 0.2233 | 0.3983 | 0.4339 | P30154;H0YDG7 | PPP2R1B |
| -0.1338 | 0.1100 | *-0.4342* | -0.1527 | 0.2726 | 0.4343 | Q14141;B1AMS2 | SEPT6 |
| 0.3666 | -0.3735 | *2.7366* | 0.9099 | 1.6247 | 0.4344 | K7N7B3;Q5T4T6;H7C4Q1 | SYCP2L |
| 0.0684 | -0.8335 | *-0.0558* | -0.2736 | 0.4888 | 0.4345 | Q53GS7 | GLE1 |
| 0.1192 | -0.5980 | *-0.1322* | -0.2037 | 0.3639 | 0.4346 | O15344;C9J453;A0A087X255;A0A087X0X0;C9JZJ7 | MID1 |
| -1.2407 | -3.2565 | *0.9584* | -1.1796 | 2.1081 | 0.4347 | Q9Y5Q6 | INSL5 |
| -0.1243 | -0.2557 | *0.0885* | -0.0972 | 0.1737 | 0.4348 | Q08380 | LGALS3BP |
| 0.1708 | -0.2560 | *4.5305* | 1.4818 | 2.6489 | 0.4348 | Q7L1Q6;C9IZ80;C9JFN4;C9J188;C9JWF5;C9JV57 | BZW1 |
| -0.4856 | 0.5148 | *-4.1426* | -1.3712 | 2.4517 | 0.4349 | Q9UIV1 | CNOT7 |
| -0.2014 | 0.4739 | *0.3214* | 0.1980 | 0.3542 | 0.4351 | Q15772;B9ZVR7 | SPEG |
| 0.4466 | -0.9475 | *-0.7721* | -0.4243 | 0.7593 | 0.4352 | Q96NL3 | ZNF599 |
| 0.5240 | -1.1272 | *-0.8922* | -0.4985 | 0.8932 | 0.4357 | Q9ULV0 | MYO5B |
| 0.7334 | 0.2157 | *-0.1814* | 0.2559 | 0.4587 | 0.4359 | P51570;K7EII7;K7ERJ9 | GALK1 |
| -0.0169 | 0.0733 | *-2.3863* | -0.7766 | 1.3948 | 0.4366 | Q92618;A0A087WUJ4;F5H2K2;H0YH41 | ZNF516 |
| 0.3592 | -0.2328 | *-5.8488* | -1.9075 | 3.4261 | 0.4366 | P29728;A0A087X0V5 | OAS2 |
| 0.7442 | 0.0051 | *-0.0229* | 0.2421 | 0.4350 | 0.4368 | P32942;K7EQC7 | ICAM3 |
| -0.1048 | 0.1441 | *-1.8561* | -0.6056 | 1.0901 | 0.4375 | Q9BT78;D6RAX7;D6RFN0;D6RD63;D6REK7 | COPS4 |
| -0.0367 | -2.9042 | *0.1105* | -0.9434 | 1.6996 | 0.4378 | O14744;G3V580;G3V2X6;H0YJX6;C9JSX3;G3V2F5;G3V2L6;G3V507;G3V5L5;H0YJ77;H0YJY6 | PRMT5 |
| 0.1494 | 1.4869 | *-0.1719* | 0.4881 | 0.8797 | 0.4379 | Q9Y6V0;A0A087X1H5;H7C114;H7 | PCLO |

| L100/ CTRL1 | L100/ CTRL2 | L100/ CTRL3 | Mean | SD | T-test p-value | Accession | Gene Symbol |
|---|---|---|---|---|---|---|---|
| | | | | | | C261;H7C440;I3L3H7;J3QRE1;Q8IYB7;Q9NSB8 | |
| 2.1430 | -0.2661 | *0.2362* | 0.7044 | 1.2709 | 0.4384 | O94956;A0A024R5I4;E9PI53;E9PIU9;E9PN87;A0A1B0GX35 | SLCO2B1 |
| -0.3057 | 0.4592 | *-6.4626* | -2.1030 | 3.7948 | 0.4384 | Q14940;J3KSE2;J3QRV1 | SLC9A5 |
| -2.3883 | -0.6594 | *0.5745* | -0.8244 | 1.4883 | 0.4386 | Q9P2S2;G5E9G7;P58401 | NRXN2 |
| -0.1184 | 0.4263 | *0.1445* | 0.1508 | 0.2724 | 0.4389 | Q14738;E9PFR3;H0Y8C4;H0YN84;H7C5Q9 | PPP2R5D |
| -0.1729 | 0.2669 | *-3.6657* | -1.1906 | 2.1548 | 0.4396 | A0A2R8Y8C6;Q8WXX7;A0A024RDL5;A0A087WVB5;A0A2R8Y522;A0A2R8Y568;Q75MD7;H7C090;H7C2P0 | AUTS2 |
| -2.4816 | 1.6513 | *-4.0469* | -1.6257 | 2.9439 | 0.4398 | P05386 | RPLP1 |
| 0.0251 | -0.0357 | *0.4252* | 0.1382 | 0.2504 | 0.4400 | Q01082;A0A087WUZ3;F8W6C1;B1ANM7;F5H6I6;F8VNZ0;Q9UNN5 | SPTBN1 |
| 0.1841 | -0.2357 | *-0.5885* | -0.2134 | 0.3868 | 0.4402 | Q13243;B4DJK0;B4DUA4;G3V5K8 | SRSF5 |
| -0.2975 | -0.6946 | *0.2271* | -0.2550 | 0.4623 | 0.4403 | Q9NQ89;F5GXX6;H0YGS6 | C12orf4 |
| -2.5704 | 0.0096 | *0.0700* | -0.8303 | 1.5073 | 0.4407 | P46063;F8WD97;F5GYB7;F5H2L2;F5H3W0;F5H4P4;F8WA66 | RECQL |
| -0.3859 | 0.3001 | *-0.9335* | -0.3398 | 0.6181 | 0.4415 | A0A1B0GV52;A0A1B0GVU4;A0A1B0GTY0 | CLCN2 |
| -0.1967 | 2.5660 | *0.1194* | 0.8296 | 1.5121 | 0.4423 | Q9Y281;F8WDN3;G3V2U0 | CFL2 |
| 0.1920 | -0.2890 | *3.3539* | 1.0856 | 1.9790 | 0.4423 | P62314;J3QLI9 | SNRPD1 |
| -0.3658 | 0.3903 | *-2.3882* | -0.7879 | 1.4365 | 0.4424 | F5GWN5;O00750;Q5SW97;Q5SW98 | PIK3C2B |
| -0.4337 | 0.3747 | *-1.4173* | -0.4921 | 0.8974 | 0.4425 | Q9NZJ4 | SACS |
| -1.0119 | 1.0712 | *-6.4020* | -2.1142 | 3.8566 | 0.4426 | Q9BXL6;I3L414 | CARD14 |
| -0.2391 | -1.4207 | *0.2489* | -0.4703 | 0.8585 | 0.4428 | Q16543;K7EQA9;K7EKQ2;K7EL68;K7EIU0 | CDC37 |
| -1.4975 | 1.4681 | *-7.2298* | -2.4197 | 4.4217 | 0.4433 | A0A087WYL5;Q6UXD5 | SEZ6L2 |
| 0.1687 | 0.9625 | *-0.1744* | 0.3189 | 0.5831 | 0.4435 | P26373;J3QSB4;J3KS98 | RPL13 |
| -0.0938 | 0.0671 | *-0.1756* | -0.0675 | 0.1235 | 0.4439 | Q96QK1;A0A1W2PP10;I3L4P4;I3L4S0 | VPS35 |
| 0.0711 | -0.0846 | *-0.2481* | -0.0872 | 0.1596 | 0.4439 | O94855 | SEC24D |

| L100/ CTRL1 | L100/ CTRL2 | L100/ CTRL3 | Mean | SD | T-test p-value | Accession | Gene Symbol |
|---|---|---|---|---|---|---|---|
| 0.3280 | -0.0332 | *-8.0422* | -2.5825 | 4.7317 | 0.4443 | Q9P107;K7EQR5 | GMIP |
| 0.5044 | 0.5188 | *-0.2774* | 0.2486 | 0.4556 | 0.4443 | Q9UJW0;H9KVE0;E5RGG1 | DCTN4 |
| -0.1336 | 0.2259 | *-2.7597* | -0.8891 | 1.6299 | 0.4445 | Q8N1G4;J3KRP5 | LRRC47 |
| 1.2678 | 0.6489 | *-0.4728* | 0.4813 | 0.8823 | 0.4445 | Q9Y536;A0A075B767;A0A0B4J2A2;P0DN37;A0A0H2UH34 | PPIAL4A |
| 0.0029 | -0.0388 | *0.9385* | 0.3009 | 0.5526 | 0.4452 | Q7Z406;M0QY43 | MYH14 |
| -0.0990 | 0.1218 | *-0.9019* | -0.2930 | 0.5387 | 0.4456 | O15357;A0A0A0MTP6 | INPPL1 |
| -0.4086 | -0.6873 | *0.2820* | -0.2713 | 0.4990 | 0.4458 | P15153;B1AH77;B1AH80;B1AH78 | RAC2 |
| 2.4347 | 1.0903 | *-0.8416* | 0.8945 | 1.6469 | 0.4462 | Q9Y4F1;C9JME2;M0R262;H0Y783;A0A1B0GV68 | FARP1 |
| -0.3738 | -0.2504 | *0.1647* | -0.1532 | 0.2821 | 0.4463 | H0Y3K2;Q6ZT12;H7C481 | UBR3 |
| -0.8742 | -3.8156 | *0.8485* | -1.2804 | 2.3584 | 0.4463 | Q6AWC2;D6R9P8 | WWC2 |
| -0.9810 | 2.5474 | *1.3568* | 0.9744 | 1.7950 | 0.4464 | Q9NYQ7 | CELSR3 |
| 1.4944 | -0.1268 | *0.0720* | 0.4799 | 0.8842 | 0.4464 | Q8IX12;A0A0C4DGG8;F5H1H2;F5H2E6;F5H3E1 | CCAR1 |
| -0.3597 | 0.3330 | *-1.3520* | -0.4596 | 0.8469 | 0.4465 | P62701 | RPS4X |
| -0.0127 | 0.1222 | *-2.6490* | -0.8465 | 1.5624 | 0.4471 | D6RBT3;O75380 | NDUFS6 |
| -0.1569 | 0.1078 | *1.4052* | 0.4520 | 0.8360 | 0.4478 | Q5SW79;H0Y2V6;H0YB92;H0Y4T4;E7EWM2;Q96L14;E5RG47;E5RGW7 | CEP170 |
| 0.3276 | -0.4422 | *-0.8683* | -0.3276 | 0.6061 | 0.4480 | Q9C0E4;A0A087WX15;A0A1B0GWJ3;A0A1B0GUL8 | GRIP2 |
| -0.2363 | 0.3223 | *-2.6068* | -0.8403 | 1.5552 | 0.4481 | Q14257;H0YL43;A8MXP8 | RCN2 |
| -0.7309 | -0.3916 | *0.2854* | -0.2790 | 0.5174 | 0.4489 | Q9UM54;E7EW20;A0A0D9SGC1;A0A0A0MRM8 | MYO6 |
| 0.0365 | -0.0098 | *-0.6102* | -0.1945 | 0.3608 | 0.4490 | Q15365 | PCBP1 |
| 0.1561 | -0.3528 | *4.6049* | 1.4694 | 2.7273 | 0.4492 | A4FU69;H0Y843 | EFCAB5 |
| -0.2204 | 1.1741 | *0.2017* | 0.3851 | 0.7151 | 0.4494 | P19013 | KRT4 |
| -2.0754 | -0.2092 | *0.2808* | -0.6680 | 1.2433 | 0.4503 | A0A1W2PRB8 | MED17 |
| -1.3432 | 0.9796 | *-2.4663* | -0.9433 | 1.7574 | 0.4507 | B1AKV2;B1AKV4;B1AKV6;F8WCR2;Q9NVA1;B1AKV3;H7BZM2 | UQCC1 |
| 0.3305 | -0.1915 | *-3.1870* | -1.0160 | 1.8982 | 0.4518 | B7WNT5;P36402 | TCF7 |
| -0.0595 | 0.0983 | *-0.8848* | -0.2820 | 0.5279 | 0.4525 | Q16851;A0A087WYS1;E7EUC7;C9JNZ1;C9JQU9;C9JTZ5;C9JUW1;C9JVG3;C9JW | UGP2 |

| L100/ CTRL1 | L100/ CTRL2 | L100/ CTRL3 | Mean | SD | T-test p-value | Accession | Gene Symbol |
|---|---|---|---|---|---|---|---|
| | | | | | | G0;Q9BW85;F2Z3H1 | |
| -1.0900 | 1.3707 | *3.0533* | 1.1113 | 2.0838 | 0.4531 | A0A087WT12;A0A087X2I2;P36969 | GPX4 |
| -0.3231 | -0.9366 | *0.2836* | -0.3254 | 0.6101 | 0.4532 | Q14008;H0YDX5;E9PQH5;H0YCF6;H0YEK7 | CKAP5 |
| 2.0396 | 5.0546 | *-1.6919* | 1.8008 | 3.3796 | 0.4535 | O00560;G5EA09;B4DHN5;E9PBU7 | SDCBP |
| 0.3172 | -0.1753 | *0.3080* | 0.1500 | 0.2817 | 0.4539 | P31644 | GABRA5 |
| -0.3903 | -0.0183 | *0.0374* | -0.1238 | 0.2325 | 0.4539 | P18669 | PGAM1 |
| 2.0626 | 2.7250 | *-1.3230* | 1.1548 | 2.1713 | 0.4542 | Q9Y4G8;A0A2R8YGD3;A0A2R8Y661 | RAPGEF2 |
| 3.6996 | -9.3718 | *-4.9312* | -3.5345 | 6.6467 | 0.4543 | P01023;F5H1E8;F8W7L3 | A2M |
| -0.2840 | 0.4124 | *-3.0308* | -0.9675 | 1.8205 | 0.4545 | P16435;H0Y4R2;E7EMD0;E7EWU0 | POR |
| 0.2266 | -0.2757 | *-0.6557* | -0.2349 | 0.4425 | 0.4549 | P14550;Q5T621;V9GYG2;V9GYP9 | AKR1A1 |
| 0.5989 | -1.0034 | *-1.1288* | -0.5111 | 0.9633 | 0.4551 | Q9NRR5 | UBQLN4 |
| 0.3052 | -0.3823 | *-0.8409* | -0.3060 | 0.5768 | 0.4552 | Q9Y570 | PPME1 |
| 0.1439 | -0.2183 | *-0.3034* | -0.1260 | 0.2375 | 0.4553 | Q16566;D6RCD6;D6RE65;D6REY7 | CAMK4 |
| 0.0765 | -0.7861 | *-0.0364* | -0.2487 | 0.4689 | 0.4553 | A6NMY6 | ANXA2P2 |
| 0.7663 | -0.4501 | *0.8322* | 0.3828 | 0.7221 | 0.4554 | P11215 | ITGAM |
| -0.2519 | 4.2290 | *0.0099* | 1.3290 | 2.5149 | 0.4566 | A0A1W2PQS6;A0A2R8Y6L3;A0A2R8Y7H1;A0A2R8YFH6;F6U211;S4R435 | RPS10-NUDT3 |
| -0.2434 | 1.1050 | *0.2235* | 0.3617 | 0.6847 | 0.4568 | Q05519;Q5T760 | SRSF11 |
| -0.2103 | 0.3423 | *0.4039* | 0.1786 | 0.3382 | 0.4569 | P47914 | RPL29 |
| -0.2471 | 0.2434 | *-0.9298* | -0.3112 | 0.5892 | 0.4569 | P59998;F8WCF6;A0A0A6YYG9;F8WDD7;H7C0A3;F8WE39;F8WDW3;R4GN08 | ARPC4 |
| -0.6839 | 0.7395 | *2.3241* | 0.7932 | 1.5048 | 0.4576 | Q08AF3;B4E128 | SLFN5 |
| -0.2268 | 0.2792 | *0.6236* | 0.2253 | 0.4277 | 0.4578 | Q96MU7;J3KRG5;J3QR07 | YTHDC1 |
| -0.1361 | 0.4698 | *-5.7872* | -1.8179 | 3.4509 | 0.4579 | Q9NZN3 | EHD3 |
| 4.1967 | 6.1219 | *-2.8539* | 2.4883 | 4.7255 | 0.4580 | O43347 | MSI1 |
| 0.2214 | -0.2241 | *0.8763* | 0.2912 | 0.5535 | 0.4584 | P51116;I3L1Z2 | FXR2 |
| 0.4579 | -1.0998 | *-0.6160* | -0.4193 | 0.7973 | 0.4585 | A0A0U1RQX8;A0A0U1RR39;A0A1B0GW38;P22681 | CBL |
| -0.2599 | -0.9730 | *0.2582* | -0.3249 | 0.6182 | 0.4588 | Q86YS6;C9JFM7 | RAB43 |

| L100/ CTRL1 | L100/ CTRL2 | L100/ CTRL3 | Mean | SD | T-test p-value | Accession | Gene Symbol |
|---|---|---|---|---|---|---|---|
| -0.1261 | 0.1227 | *0.5021* | 0.1662 | 0.3163 | 0.4589 | P55036;Q5VWC4;A6PVX3;H0Y3Y9 | PSMD4 |
| -0.4279 | 0.1776 | *-0.2375* | -0.1626 | 0.3096 | 0.4590 | Q99996;A0A0A0MRF6;H7BYL6;A0A087WX84;A0A0A0MTM1;A0A2R8Y590;A4D1F6;F2Z2X7;F5H7W1;J3KRS8;J3KTM8;J3QQZ8;P17181;Q16478;Q5H9S7;Q5JVX6;Q5JVX7;Q9C0J9;S4R3Z8;A0A0A0MRE9;H0Y6Q0 | AKAP9 |
| 0.6678 | -3.2932 | *-0.5680* | -1.0645 | 2.0266 | 0.4590 | P04114;A8MUN2;E3W980;Q8TDG4;Q9C0D6 | APOB |
| -0.2714 | 0.2902 | *-1.2261* | -0.4024 | 0.7666 | 0.4592 | P46060 | RANGAP1 |
| 0.1372 | -0.7624 | *-0.1069* | -0.2441 | 0.4652 | 0.4594 | P07949 | RET |
| -0.0467 | -0.0723 | *1.8848* | 0.5886 | 1.1226 | 0.4597 | Q8NI08 | NCOA7 |
| -0.2761 | -0.0327 | *0.0454* | -0.0878 | 0.1677 | 0.4603 | Q96EK9 | KTI12 |
| 0.8286 | 0.2988 | *-0.2671* | 0.2868 | 0.5479 | 0.4604 | P19827 | ITIH1 |
| 1.6965 | 0.4341 | *-0.4425* | 0.5627 | 1.0753 | 0.4604 | B4DYB8;F5GX71;H3BMZ9;H3BN01;H3BNY8;H3BP57;H3BPB8;H3BPM5;H3BPP3;H3BPU7;H3BT46;H3BT48;H3BUZ9;P34949 | MPI |
| -0.1983 | 0.3225 | *-2.2932* | -0.7230 | 1.3845 | 0.4612 | P00558;F5H0F9 | PGK1 |
| -2.4696 | 0.0007 | *0.1604* | -0.7695 | 1.4745 | 0.4615 | P09132 | SRP19 |
| 1.1249 | -0.8866 | *2.2402* | 0.8262 | 1.5847 | 0.4618 | Q00839;A0A1W2PPS1;A0A1X7SBS1;A0A1W2PP35;A0A1W2PQL0;A0A1W2PRZ7;A0A1W2PQD4;A0A1W2PRI6;A0A1W2PP22;A0A1W2PPE9 | HNRNPU |
| -0.0647 | 1.2341 | *-0.0189* | 0.3835 | 0.7370 | 0.4625 | Q8N7K0;C9JQA6;F8VTV7;F8VU36;F8W652 | ZNF433 |
| 0.0476 | -0.0267 | *0.0450* | 0.0219 | 0.0422 | 0.4628 | Q9Y678;H0Y8X7 | COPG1 |
| 1.1284 | -2.5197 | *-1.5535* | -0.9816 | 1.8901 | 0.4633 | Q9ULJ3;E7EVF9;Q5KS07 | ZBTB21 |
| -0.2770 | 0.1294 | *2.3437* | 0.7320 | 1.4104 | 0.4635 | P62195;J3QQM1;J3QSA9;J3KRP2;J3QLH6;J3QSE0;J3QRW1;J3QRR3 | PSMC5 |
| 0.1635 | -0.0057 | *-2.2427* | -0.6950 | 1.3430 | 0.4647 | P09960;B4DEH5 | LTA4H |

| L100/ CTRL1 | L100/ CTRL2 | L100/ CTRL3 | Mean | SD | T-test p-value | Accession | Gene Symbol |
|---|---|---|---|---|---|---|---|
| -0.3469 | 0.3774 | *1.0653* | 0.3652 | 0.7062 | 0.4649 | H0Y786;H7BZD5 | NEB |
| -0.0824 | 0.0988 | *-0.4511* | -0.1449 | 0.2802 | 0.4649 | O95140;Q5JXC5 | MFN2 |
| 1.0352 | 2.6505 | *-0.9158* | 0.9233 | 1.7858 | 0.4650 | O15090;K7EKT4;K7EJP8 | ZNF536 |
| 0.8442 | 1.1762 | *-0.5771* | 0.4811 | 0.9314 | 0.4654 | Q8IVF4;A0A1C7CYW8;A0A0J9YY17;A0A096LNK1;A0A0J9YWH2 | DNAH10 |
| -0.0965 | 0.1701 | *0.1608* | 0.0781 | 0.1513 | 0.4654 | P16298;Q5F2F8;Q5F2G0;P48454 | PPP3CB |
| -7.1958 | -1.3152 | *1.5880* | -2.3077 | 4.4752 | 0.4660 | G3XAJ6;Q14699 | RFTN1 |
| -0.0416 | 0.8840 | *-0.0244* | 0.2727 | 0.5295 | 0.4665 | Q9HC52 | CBX8 |
| 0.2294 | 2.1069 | *-0.3521* | 0.6614 | 1.2852 | 0.4668 | Q9UBD5 | ORC3 |
| 0.0159 | -0.0449 | *0.4091* | 0.1267 | 0.2464 | 0.4672 | P62899;B7Z4C8;B7Z4E3;C9JU56;B8ZZK4;H7C2W9 | RPL31 |
| -0.0261 | 1.6344 | *-0.0976* | 0.5036 | 0.9799 | 0.4673 | P42025 | ACTR1B |
| 0.2988 | 0.9548 | *-0.2925* | 0.3204 | 0.6239 | 0.4676 | P29083;C9IYL4 | GTF2E1 |
| -0.8166 | 0.7322 | *-2.0981* | -0.7275 | 1.4173 | 0.4678 | F5H2A4;F5H2U8;F5H6H0;P52926 | HMGA2 |
| -0.2688 | 0.1390 | *1.9372* | 0.6024 | 1.1738 | 0.4678 | Q8N653;H7BZQ9;H7C0X1;H7C305 | LZTR1 |
| 0.1970 | -0.1233 | *-1.1999* | -0.3754 | 0.7318 | 0.4680 | E9PC15;Q53H12;E9PG39;A0A0G2JLD0;A0A0G2JLG5 | AGK |
| -0.3967 | 0.0066 | *0.0239* | -0.1221 | 0.2380 | 0.4681 | E5RG77;O94903;E5RFX7 | PLPBP |
| -0.1511 | 0.1538 | *-0.5254* | -0.1742 | 0.3402 | 0.4686 | Q96T51;J3KPP6;H0YA47 | RUFY1 |
| 0.3197 | 2.8939 | *-0.4961* | 0.9059 | 1.7694 | 0.4688 | Q5T4S7;X6R960 | UBR4 |
| -0.5718 | 1.0296 | *0.9114* | 0.4564 | 0.8924 | 0.4692 | Q9NTJ4;B4DH23;H3BQH9;H3BRI3 | MAN2C1 |
| 4.4651 | 0.7375 | *-0.9582* | 1.4148 | 2.7744 | 0.4703 | P17040 | ZSCAN20 |
| -0.3324 | -1.5292 | *0.3848* | -0.4922 | 0.9669 | 0.4709 | Q96G03;E7ENQ8;E9PD70 | PGM2 |
| -0.1388 | 0.1511 | *0.3957* | 0.1360 | 0.2676 | 0.4716 | A0A087WZQ7;Q9H115 | NAPB |
| -0.3889 | -0.1700 | *6.6951* | 2.0454 | 4.0283 | 0.4719 | Q5GLZ8 | HERC4 |
| -0.0177 | -0.0268 | *0.5284* | 0.1613 | 0.3179 | 0.4723 | Q15008;C9J0E9;C9J7B7;H7C531 | PSMD6 |
| 0.0213 | -0.9780 | *0.0609* | -0.2986 | 0.5887 | 0.4723 | P39748;I3L3E9;F5H1Y3 | FEN1 |
| 0.4570 | -0.4468 | *1.3732* | 0.4611 | 0.9100 | 0.4727 | O60240;H0YM16 | PLIN1 |
| -0.1479 | 0.6565 | *-6.1578* | -1.8831 | 3.7238 | 0.4735 | Q01658 | DR1 |
| -0.2249 | 0.4421 | *-2.8847* | -0.8892 | 1.7601 | 0.4738 | P51513;F8VWX1;F8W659;I3L2B5;J3KQU3;F8VW64;G8JLA5 | NOVA1 |
| 0.8362 | 0.8577 | *-0.5082* | 0.3952 | 0.7825 | 0.4739 | P50914;E7EPB3 | RPL14 |

| L100/ CTRL1 | L100/ CTRL2 | L100/ CTRL3 | Mean | SD | T-test p-value | Accession | Gene Symbol |
|---|---|---|---|---|---|---|---|
| -0.4068 | -0.0049 | *0.0393* | -0.1241 | 0.2458 | 0.4739 | A0A2R8YFB8;G3V196;P59190;G3V562 | RAB15 |
| 0.2025 | -0.1926 | *0.5572* | 0.1891 | 0.3751 | 0.4747 | P28838;H0Y9Q1;H0Y983 | LAP3 |
| 0.8328 | 1.1026 | *-0.5756* | 0.4533 | 0.9012 | 0.4755 | Q8NB16;I3L2U3 | MLKL |
| 0.2499 | 0.1797 | *-0.1271* | 0.1008 | 0.2005 | 0.4757 | A0A087WW77;Q15334 | LLGL1 |
| -0.3172 | -0.5648 | *0.2514* | -0.2102 | 0.4185 | 0.4760 | P05455;B5BUB5;E7ERC4;E9PFL9;E9PGX9 | SSB |
| -0.5098 | 1.7831 | *-14.8823* | -4.5363 | 9.0329 | 0.4761 | P35789 | ZNF93 |
| 0.3349 | 0.0849 | *-0.0948* | 0.1083 | 0.2158 | 0.4763 | P49736;H0Y8E6;F8WDM3;C9J013;C9JZ21;H7C4N9 | MCM2 |
| 0.2289 | 0.5439 | *-8.4017* | -2.5429 | 5.0762 | 0.4770 | O95292;E5RK64 | VAPB |
| 0.0726 | 0.3756 | *-4.8900* | -1.4806 | 2.9565 | 0.4772 | A0A087WWT6;A0A0A0MSU1;Q06455 | RUNX1T1 |
| 0.1454 | -0.5501 | *4.5713* | 1.3889 | 2.7779 | 0.4778 | Q13509;G3V2N6;G3V2R8;G3V3R4;G3V5W4;G3V3J6;G3V3W7;G3V2A3;G3V542 | TUBB3 |
| -0.2718 | 0.9064 | *0.2505* | 0.2950 | 0.5903 | 0.4780 | P17987;E7ERF2;E7EQR6;F5H282;F5GZ03;F5H136;F5H676;F5H726;F5GYL4;F5H7Y1;F5GZI8 | TCP1 |
| -0.0076 | 0.2113 | *-2.2175* | -0.6713 | 1.3436 | 0.4781 | B7ZAQ6;P0CG08 | GPR89A |
| 0.0195 | 0.1212 | *-1.5120* | -0.4571 | 0.9150 | 0.4781 | Q15006;E5RGJ2 | EMC2 |
| -0.2285 | 0.3008 | *0.4794* | 0.1839 | 0.3682 | 0.4781 | O75390;B4DJV2;A0A0C4DGI3;F8W4S1;F8VTT8;F8VPA1;F8VWQ5;F8W1S4;F8VRI6;H0YH82;F8VPF9;F8VRP1;F8VX68;F8VR34;F8VX07;F8VZK9;H0YIC4;F8W642;F8VU34;F8W0J2 | CS |
| -0.1505 | 0.3020 | *-1.8522* | -0.5669 | 1.1359 | 0.4785 | A0A0A0MSK3;A0A0C4DH47;B7Z6W1;C9JS68;P24557;Q53F23 | TBXAS1 |
| 0.8851 | 1.4429 | *-0.6805* | 0.5492 | 1.1008 | 0.4786 | Q00610;A0A087WVQ6;J3KS13;K7EJJ5;J3KRF5;J3KSQ2;J3QL20 | CLTC |
| -0.6360 | 0.8533 | *1.2945* | 0.5039 | 1.0116 | 0.4792 | P13591;A0A087WX77;A0A087WVU1;A0A087WZS4;A0A087WTR3 | NCAM1 |

| L100/ CTRL1 | L100/ CTRL2 | L100/ CTRL3 | Mean | SD | T-test p-value | Accession | Gene Symbol |
|---|---|---|---|---|---|---|---|
| | | | | | | ;A0A0C4DGS4;A0A0A0MT39;A0A0C4DG82;A0A1W2PR94;C9J6Q2;C9J7M5;C9JA15;C9JAW0;D6RB49;E9PG18;E9PHB6;H7C064;H9KVD2;K4DIA1;Q14524;Q6ZMT9;Q86V90;Q9BSK4;Q9NY84;Q9UI33;A0A0D9SF30;H7BYX6;A0A087WWJ5 | |
| -0.3659 | -5.7999 | *0.8715* | -1.7648 | 3.5489 | 0.4798 | O14976 | GAK |
| 0.5910 | -0.1914 | *-4.4176* | -1.3393 | 2.6944 | 0.4800 | P26583;D6R9A6 | HMGB2 |
| -0.1410 | 0.1373 | *0.4269* | 0.1411 | 0.2839 | 0.4802 | P11142;E9PKE3;E9PNE6;A8K7Q2;E9PN89;E9PS65;E9PLF4;E9PK54;E9PQK7;E9PQQ4;E9PPY6;E9PI65;E9PN25;E9PM13 | HSPA8 |
| -0.5051 | 0.6840 | *-2.8874* | -0.9028 | 1.8186 | 0.4805 | Q8TD23 | ZNF675 |
| -0.0927 | 0.1830 | *-1.0726* | -0.3275 | 0.6599 | 0.4807 | Q14558;B4DP31;C9J168;C9JKT9;C9JNQ3;C9JUN4 | PRPSAP1 |
| -4.4740 | 2.6538 | *-4.1794* | -1.9999 | 4.0329 | 0.4809 | Q7L311 | ARMCX2 |
| -0.1476 | -0.1562 | *3.0567* | 0.9176 | 1.8525 | 0.4813 | P35052;H7C410;H7C024 | GPC1 |
| 0.8373 | 0.1793 | *-0.2222* | 0.2648 | 0.5349 | 0.4815 | A8MQ14 | ZNF850 |
| 0.2184 | -0.1768 | *0.3888* | 0.1435 | 0.2902 | 0.4820 | Q86UL8;A0A0D9SGI1;A0A0E3D6M1;E7EWI0;A0A1B0GTC0;A0A0D9SEY4;A0A0D9SGF8;A0A0D9SFP3;A0A1B0GVS6 | MAGI2 |
| -0.8322 | -0.5587 | *0.4174* | -0.3245 | 0.6569 | 0.4823 | Q01968 | OCRL |
| -0.2082 | 0.3037 | *-1.3376* | -0.4140 | 0.8398 | 0.4831 | A0A087WT99;Q9H0W9;E9PQS1;E9PPB5;E9PIP1;E9PJU8;E9PLB3;E9PLC5;E9PR95;E9PSC3 | C11orf54 |
| 0.7783 | -0.6842 | *1.6491* | 0.5810 | 1.1791 | 0.4833 | P68371;M0R2D3;A0A075B724 | TUBB4B |
| 0.4977 | -0.2982 | *-2.4332* | -0.7446 | 1.5156 | 0.4844 | Q8WXG6;A0A0A0MRB5;C9JLZ9;C9K0L0;C9JM97 | MADD |
| -0.0091 | -0.0480 | *0.5507* | 0.1645 | 0.3350 | 0.4846 | H0Y2S9;H0Y7E2;H7C3G6;J3QRL2;K7EL39;Q6WCQ1;J3KSW8 | MPRIP |

| L100/ CTRL1 | L100/ CTRL2 | L100/ CTRL3 | Mean | SD | T-test p-value | Accession | Gene Symbol |
|---|---|---|---|---|---|---|---|
| -0.2693 | 0.3016 | *-0.9707* | -0.3128 | 0.6372 | 0.4848 | P55809;E9PDW2 | OXCT1 |
| 1.6294 | -0.3771 | *0.2572* | 0.5032 | 1.0256 | 0.4849 | Q8N823;A0A1W2PQH1;C9IZE3;C9J871;C9JVX0;M0QXE4;M0QXQ6;M0QYR0;M0QYZ5;M0R1T1;M0R2W3 | ZNF611 |
| -1.2598 | 0.0710 | *0.0620* | -0.3756 | 0.7658 | 0.4851 | P11137;A8MZ31;E7EV03 | MAP2 |
| -0.5512 | 0.6005 | *-1.8583* | -0.6030 | 1.2302 | 0.4853 | P51843;A6NNU8 | NR0B1 |
| -0.6115 | -1.5728 | *0.5929* | -0.5304 | 1.0851 | 0.4863 | Q14204;A0A2R8Y706;A0A2R8Y5T0;A0A2R8YFZ7;A0A2R8Y6H2;A0A2R8Y6I5;A0A2R8YGC7;D6RIG7;M0R044;M0R0F2;M0R189;O75298;Q7RTN0;Q8N594;Q8WWL2;Q9Y5H4;W4VSR2;X6RC15;A0A2R8Y542 | DYNC1H1 |
| -0.6295 | -1.2377 | *0.5417* | -0.4419 | 0.9044 | 0.4865 | Q9Y6D9;C9JKI7;C9JP81;C9JTA2 | MAD1L1 |
| -0.5382 | 0.3699 | *2.2575* | 0.6964 | 1.4262 | 0.4868 | Q14667;K7EQ86;Q08E86;K7EQR3 | KIAA0100 |
| 0.3227 | -0.4648 | *-0.5730* | -0.2384 | 0.4889 | 0.4873 | Q92688 | ANP32B |
| -0.0282 | -0.1655 | *0.0406* | -0.0510 | 0.1049 | 0.4881 | Q9UL46;A0A087X1Z3;H0YM70 | PSME2 |
| 0.1408 | -0.2295 | *1.0355* | 0.3156 | 0.6504 | 0.4891 | P35222;A0A2R8Y543;A0A2R8Y5A3;A0A2R8Y804;A0A2R8YCH5;B4DGU4;A0A2R8Y5Z1;A0A2R8Y7Z0;A0A2R8Y750;B5BU28;A0A2R8Y5C3;A0A2R8Y6G0;E7EMJ5;A0A2R8Y815;A0A2R8YG06;E7EV28;E9PDF9 | CTNNB1 |
| 0.2012 | -0.3026 | *-0.3372* | -0.1462 | 0.3014 | 0.4892 | P09471;A0A1W2PRJ7;A0A1W2PS82;H3BTM2;A0A1W2PP38;A0A1W2PP87;A0A1W2PQK2;A0A1W2PRE1;A0A1W2PQ24;H3BNR5 | GNAO1 |
| -2.6412 | 0.7292 | *-0.5629* | -0.8250 | 1.7004 | 0.4892 | Q9ULK5 | VANGL2 |
| 0.9112 | -0.5693 | *0.8948* | 0.4123 | 0.8501 | 0.4893 | Q03113 | GNA12 |
| -0.0552 | -0.0277 | *0.7369* | 0.2180 | 0.4496 | 0.4894 | P62269;J3JS69;A0A0G2JQH2;Q5GGW2 | RPS18 |

| L100/ CTRL1 | L100/ CTRL2 | L100/ CTRL3 | Mean | SD | T-test p-value | Accession | Gene Symbol |
|---|---|---|---|---|---|---|---|
| -0.1530 | -0.8480 | *0.2161* | -0.2616 | 0.5403 | 0.4899 | P17252;J3KRN5 | PRKCA |
| -0.0183 | -0.4788 | *0.0700* | -0.1424 | 0.2947 | 0.4908 | O60888;C9IZG4; C9IZQ5 | CUTA |
| -2.3754 | -1.1992 | *1.0518* | -0.8410 | 1.7415 | 0.4909 | Q9Y5G2 | PCDHGB2 |
| -0.2000 | 0.1578 | *-0.3149* | -0.1190 | 0.2465 | 0.4911 | O60256;E7EPA1; I3L0S1;C9JJS3;C9K0K7;C9JDU5; C9JDH0;E7EW35;I3L4G9;I3L164; I3L331;I3L1T1;I3L2J4;I3L3B8 | PRPSAP2 |
| -0.1900 | -0.2715 | *0.1433* | -0.1061 | 0.2198 | 0.4911 | Q9Y383;A0A0A6YYJ8 | LUC7L2 |
| 0.3797 | -0.0225 | *-0.0220* | 0.1117 | 0.2321 | 0.4921 | Q08211 | DHX9 |
| -0.2133 | 0.1298 | *0.9331* | 0.2832 | 0.5884 | 0.4922 | Q9H4A3;F5GWT4;F6UYG0;H0YH68 | WNK1 |
| -0.2608 | 4.3171 | *-0.2501* | 1.2688 | 2.6400 | 0.4927 | P54819;F8W1A4; F8VY04;F8VZG5; G3V213;F8VPP1 | AK2 |
| -0.8308 | 0.7062 | *-1.5164* | -0.5470 | 1.1382 | 0.4928 | O43776;K7EIU7; K7EPK2;K7EMQ6;K7EQ35 | NARS |
| 0.6572 | 2.2184 | *-0.7442* | 0.7104 | 1.4820 | 0.4937 | A0A0G2JMX5;A0A0G2JNP3;A0A0G2JNY7;A0A0G2JPM8;O75023 | LILRB5 |
| -0.0109 | 0.1394 | *-1.1039* | -0.3251 | 0.6786 | 0.4939 | Q5T8A7 | PPP1R26 |
| -0.0668 | -0.1115 | *1.4611* | 0.4276 | 0.8953 | 0.4951 | Q9H936;E9PJH7; A0A0D9SEI9;A0A0D9SFE1;E9PS95;K4DIA8 | SLC25A22 |
| -0.4955 | -0.5507 | *0.3354* | -0.2369 | 0.4964 | 0.4953 | Q14157;F8W726; Q5VU77;Q5VU79;Q5VU80;Q5VU81 | UBAP2L |
| -0.3204 | 1.7155 | *0.1324* | 0.5092 | 1.0690 | 0.4961 | P55196;J3KN01; A8MQ02;Q5TIG5;H0Y7R8;H0Y8L4;H0Y9I0;H0Y948; H0YA98 | AFDN |
| -0.6757 | 0.7569 | *1.4956* | 0.5256 | 1.1040 | 0.4963 | P51153 | RAB13 |
| -0.2252 | 0.2708 | *-0.8365* | -0.2636 | 0.5546 | 0.4969 | Q9BYK8 | HELZ2 |
| -1.2310 | -0.2587 | *0.3521* | -0.3792 | 0.7984 | 0.4972 | C9JFP8 | SHANK2 |
| -0.1865 | 0.2178 | *-0.6413* | -0.2033 | 0.4298 | 0.4986 | P55287 | CDH11 |
| 1.3228 | -5.7499 | *-0.7354* | -1.7208 | 3.6379 | 0.4987 | Q16853 | AOC3 |
| -0.2813 | -1.7690 | *0.4477* | -0.5342 | 1.1298 | 0.4989 | Q53FP2 | TMEM35A |
| 0.3368 | -0.9288 | *-0.3046* | -0.2989 | 0.6328 | 0.4993 | O95450 | ADAMTS2 |
| -0.0138 | 0.0565 | *-0.3507* | -0.1027 | 0.2177 | 0.4998 | P78310 | CXADR |
| -0.9276 | -0.0358 | *0.1492* | -0.2714 | 0.5757 | 0.5000 | P23528;E9PP50; E9PK25;G3V1A4;E9PQB7;E9PS23;E9PLJ3 | CFL1 |

| L100/ CTRL1 | L100/ CTRL2 | L100/ CTRL3 | Mean | SD | T-test p-value | Accession | Gene Symbol |
|---|---|---|---|---|---|---|---|
| -0.6888 | 0.2651 | *-0.2509* | -0.2249 | 0.4775 | 0.5003 | P07202;A0A0G2JNN7;A0A0G2JP53;A0A0G2JR31;A0A0G2JRZ1;A0A0G2JR90;A0A0G2JR79;H0Y6H4;C9J511;E9PFM6;A0A0G2JS21;H7C5B6 | TPO |
| 0.2497 | 0.1660 | *-3.1773* | -0.9205 | 1.9548 | 0.5004 | Q9H0R4;K7ER15 | HDHD2 |
| 0.6042 | -0.0697 | *-4.2300* | -1.2319 | 2.6183 | 0.5007 | P51114;B4DXZ6;E7EU85;E9PFF5 | FXR1 |
| -0.2426 | 0.2802 | *-0.7966* | -0.2530 | 0.5385 | 0.5012 | Q7L2E3;F6R0H4 | DHX30 |
| 0.2752 | 0.3795 | *-4.9464* | -1.4306 | 3.0452 | 0.5013 | O43290 | SART1 |
| 0.0323 | -0.1936 | *1.2777* | 0.3721 | 0.7923 | 0.5014 | Q9Y6E2;E7ETZ4;B5MCH7;B5MCE7;Q75MG1;E9PFD4;E7EMS9;C9JF98;E9PFE3;F8WDX8 | BZW2 |
| 1.3723 | -0.5152 | *0.4719* | 0.4430 | 0.9441 | 0.5017 | E9PB61;Q86V81 | ALYREF |
| -0.2178 | 0.1950 | *0.5930* | 0.1900 | 0.4054 | 0.5021 | Q9H2U2;D6RGV9;H0Y9D8;D6RGI1;H0YAK2;D6R967 | PPA2 |
| 0.2272 | -0.3450 | *-0.3478* | -0.1552 | 0.3312 | 0.5022 | Q15637;C9J792;H7C561 | SF1 |
| 0.5642 | -1.0466 | *4.4614* | 1.3264 | 2.8320 | 0.5024 | B7ZM87;O75044;A0A075B7B5;A0A286YEY3;P0DMP2;P0DJJ0;A0A075B7C8 | SRGAP2 |
| -1.5384 | -1.1303 | *0.8658* | -0.6010 | 1.2865 | 0.5034 | Q5T890 | ERCC6L2 |
| 0.0925 | -0.4771 | *-0.0340* | -0.1395 | 0.2991 | 0.5039 | P09496;C9J8P9;F8WF69 | CLTA |
| 0.7917 | -0.8139 | *-1.8225* | -0.6149 | 1.3184 | 0.5040 | Q6ULP2 | AFTPH |
| -0.6261 | -0.1510 | *0.1982* | -0.1930 | 0.4138 | 0.5040 | P08754 | GNAI3 |
| -0.7047 | -2.6458 | *0.8805* | -0.8233 | 1.7662 | 0.5042 | C9J3G2;F2Z2T4;Q8IUF1;Q9BRT8;A0A087WZQ3;A0A087X140;F8WEG4;F8WEU0 | CBWD2 |
| 0.2154 | -0.2710 | *0.8182* | 0.2542 | 0.5456 | 0.5044 | P12235;V9GYG0 | SLC25A4 |
| -0.4309 | 0.1742 | *2.1038* | 0.6157 | 1.3238 | 0.5050 | O43237;J3KRI4;J3KRZ2;B4E2E0;J3KSD2 | DYNC1LI2 |
| 0.3727 | -0.5438 | *1.8656* | 0.5649 | 1.2162 | 0.5056 | Q9UMD9;H0Y420 | COL17A1 |
| -0.1085 | 0.8484 | *-0.0078* | 0.2440 | 0.5258 | 0.5058 | Q13315;E9PIN0;A0A087X0E9;M0QXY8 | ATM |
| 0.0688 | -0.3442 | *-0.0258* | -0.1004 | 0.2164 | 0.5059 | P78563;A0A0A0MSG8 | ADARB1 |
| -0.0338 | 0.0000 | *0.2468* | 0.0710 | 0.1531 | 0.5062 | Q96E17 | RAB3C |

| L100/ CTRL1 | L100/ CTRL2 | L100/ CTRL3 | Mean | SD | T-test p-value | Accession | Gene Symbol |
|---|---|---|---|---|---|---|---|
| -0.1996 | 0.3630 | *-1.4661* | -0.4342 | 0.9368 | 0.5063 | O75914;A0A087X294 | PAK3 |
| -0.0580 | -1.2458 | *0.2214* | -0.3608 | 0.7791 | 0.5066 | Q6FI81;H3BV90;H3BT65;H3BUG4;H3BPG7 | CIAPIN1 |
| -3.9666 | -0.3754 | *0.8620* | -1.1600 | 2.5081 | 0.5071 | H3BR42;H3BST1;Q9UL45;H3BRA4 | BLOC1S6 |
| -0.5226 | -0.9513 | *0.4656* | -0.3361 | 0.7267 | 0.5071 | P35609;F6THM6 | ACTN2 |
| 0.3899 | -0.1827 | *0.1958* | 0.1343 | 0.2912 | 0.5081 | Q8N782;J3KR51;J3KR62 | ZNF525 |
| -0.7262 | -0.0087 | *0.1092* | -0.2086 | 0.4522 | 0.5082 | F5H3U9;F5H6N1;F5H6P7;P61326;Q96A72 | MAGOHB |
| 0.7402 | 1.2408 | *-0.6372* | 0.4479 | 0.9725 | 0.5087 | Q70J99;K7EIH3;K7EN29;K7EN81;K7EQ37 | UNC13D |
| 2.0883 | 0.5591 | *-0.7119* | 0.6452 | 1.4021 | 0.5090 | A0A0A0MSX2;Q96M02 | C10orf90 |
| -0.6104 | -0.2582 | *0.2624* | -0.2020 | 0.4391 | 0.5091 | Q5THR3 | EFCAB6 |
| -0.3520 | -0.2148 | *0.1832* | -0.1279 | 0.2780 | 0.5092 | P62826;B5MDF5;J3KQE5;F5H018;H0YFC6;B4DV51 | RAN |
| -0.7321 | 4.4051 | *0.1239* | 1.2656 | 2.7523 | 0.5093 | Q96DA2 | RAB39B |
| 0.5363 | 0.5661 | *-7.5566* | -2.1514 | 4.6811 | 0.5095 | Q99729;D6R9P3;D6RBZ0;D6RD18 | HNRNPAB |
| -0.7568 | -0.1769 | *0.2421* | -0.2305 | 0.5016 | 0.5095 | P62913;Q5VVC8 | RPL11 |
| -0.6441 | 0.6393 | *1.4789* | 0.4913 | 1.0692 | 0.5095 | P24588 | AKAP5 |
| 0.5543 | 2.5769 | *-0.7959* | 0.7784 | 1.6976 | 0.5103 | Q9HBR0;H0YF92;I3L0Q8 | SLC38A10 |
| 0.3412 | 0.5013 | *-0.2774* | 0.1884 | 0.4112 | 0.5107 | Q8WU90;H7C466 | ZC3H15 |
| -0.3032 | -0.2907 | *0.1998* | -0.1314 | 0.2868 | 0.5108 | B4E2Q0;H0Y9V7;P98194;H0Y9S7 | ATP2C1 |
| -0.3016 | -0.0094 | *2.1540* | 0.6143 | 1.3413 | 0.5108 | P61019;H7C125;H0YD31;A0A1D5RMT4;H0YDL5;E9PKL7 | RAB2A |
| -0.3236 | 0.0828 | *-0.0444* | -0.0951 | 0.2079 | 0.5113 | A0A087WZG8;A0A2Q2TCK2;Q6XZB0 | LIPI |
| 0.6762 | 1.1082 | *-0.5808* | 0.4012 | 0.8774 | 0.5114 | P12036 | NEFH |
| -0.9300 | 0.1490 | *-0.0161* | -0.2657 | 0.5812 | 0.5114 | Q63ZY3 | KANK2 |
| 1.7907 | -0.8942 | *0.9922* | 0.6296 | 1.3787 | 0.5119 | Q86YS7;H0YFC4 | C2CD5 |
| 1.0933 | 3.5867 | *-1.3229* | 1.1191 | 2.4549 | 0.5125 | Q502W6;F8WBX4;F8WD48;H0YEM4;H0YF54;D6RBA5;F8VZA8;H0YIW7;Q92911 | VWA3B |
| 1.1707 | -2.2297 | *-1.3542* | -0.8044 | 1.7656 | 0.5127 | Q9NYB9;A0A0C4DG21;B7Z836;E7EP65;E7EW77;F8WAL6;H7C3Q7;E9PEZ7;H0Y6B5 | ABI2 |
| -0.4145 | -1.3981 | *0.5092* | -0.4345 | 0.9538 | 0.5128 | Q8IXI1 | RHOT2 |

| L100/ CTRL1 | L100/ CTRL2 | L100/ CTRL3 | Mean | SD | T-test p-value | Accession | Gene Symbol |
|---|---|---|---|---|---|---|---|
| 0.1942 | -0.2478 | *-0.3333* | -0.1290 | 0.2831 | 0.5128 | P52742;M0R0N7 | ZNF135 |
| 0.1748 | 0.6654 | *-5.6025* | -1.5874 | 3.4858 | 0.5129 | A0A2R8Y5C5;O43520;K7ERI0 | ATP8B1 |
| 0.1855 | -0.3587 | *-0.2112* | -0.1281 | 0.2814 | 0.5130 | P25205;J3KQ69 | MCM3 |
| -0.3682 | -0.5270 | *0.2976* | -0.1992 | 0.4375 | 0.5130 | P06748;E5RI98; E5RGW4 | NPM1 |
| -0.3889 | -0.2760 | *0.2220* | -0.1476 | 0.3250 | 0.5139 | P06576;H0YH81; H0YI37;F8W079 | ATP5F1B |
| 0.2391 | -0.3473 | *-0.3562* | -0.1548 | 0.3411 | 0.5143 | P09497;H0Y9Q6 | CLTB |
| -0.4493 | -0.7693 | *0.3979* | -0.2735 | 0.6031 | 0.5144 | P48047;H7C086; H7C068 | ATP5PO |
| 0.2262 | 0.3702 | *-3.8730* | -1.0922 | 2.4093 | 0.5146 | Q99250;A0A1B0GWA6;A0A1B0GW40;A0A1B0GW67;A0A1B0GTS6; F6U291;A0A1B0GTX0;A0A1W2PQ58;A0A1W2PSB2;C9JBM7 | SCN2A |
| 1.9444 | -1.8372 | *-4.5238* | -1.4722 | 3.2495 | 0.5148 | Q9NZR2 | LRP1B |
| 0.2024 | 0.0799 | *-1.8349* | -0.5176 | 1.1425 | 0.5149 | Q9P0M6;Q5SQT3 | H2AFY2 |
| -0.5061 | 0.8383 | *-2.9186* | -0.8621 | 1.9035 | 0.5149 | O43390;B4DT28; A0A286YEZ8 | HNRNPR |
| -0.2625 | -0.2983 | *0.1906* | -0.1234 | 0.2725 | 0.5150 | P25705;K7EK77; K7ENJ4;K7ERX7;K7EJP1;K7EQH4;K7ESA0;K7EM08;Q9UPU3 | ATP5F1A |
| -0.3830 | 0.4488 | *-1.1474* | -0.3605 | 0.7983 | 0.5160 | Q13347;Q5TFK1 | EIF3I |
| -0.3130 | 0.3582 | *0.5903* | 0.2118 | 0.4691 | 0.5160 | O95714;A0A0J9YVP0;A0A0J9YXQ8;H3BRG9 | HERC2 |
| -0.4219 | 1.2432 | *0.3078* | 0.3764 | 0.8347 | 0.5165 | Q96MZ0;A0A087WWT8;H0UIB3;H0Y6A7;Q5TE61 | GDAP1L1 |
| -0.3827 | 0.3057 | *1.0379* | 0.3203 | 0.7104 | 0.5166 | Q9NQT8;E7ERX9 | KIF13B |
| 0.4766 | -0.6789 | *-0.7134* | -0.3052 | 0.6773 | 0.5168 | Q5QJ74 | TBCEL |
| 1.0352 | -0.0605 | *-0.1029* | 0.2906 | 0.6452 | 0.5170 | P34931;Q53FA3 | HSPA1L |
| 0.6260 | -0.6688 | *1.5446* | 0.5006 | 1.1120 | 0.5172 | Q86W92;A0A0A0MTP2;H0YFE4 | PPFIBP1 |
| -0.4314 | 0.0702 | *2.4021* | 0.6803 | 1.5121 | 0.5174 | Q9Y6N5;H3BNX3;A0A1B0GXB4; H3BV36;H3BUD7 | SQOR |
| -3.2309 | 4.2767 | *5.1815* | 2.0758 | 4.6179 | 0.5177 | P41440 | SLC19A1 |
| 0.7900 | -0.6767 | *-1.9831* | -0.6233 | 1.3874 | 0.5179 | Q92616 | GCN1 |
| -0.3904 | 0.3568 | *0.9177* | 0.2947 | 0.6563 | 0.5181 | Q92626;H7C1W1;H7C300 | PXDN |
| -0.2286 | 0.2104 | *0.5324* | 0.1714 | 0.3820 | 0.5184 | Q03252 | LMNB2 |
| -0.4660 | 0.2663 | *1.6354* | 0.4786 | 1.0667 | 0.5184 | Q8WXF7;G3V321 | ATL1 |
| -0.6220 | 0.1238 | *-0.0309* | -0.1764 | 0.3936 | 0.5189 | A5PLK6;H3BU64;A0A0U1RRF6;A | RGSL1 |

| L100/ CTRL1 | L100/ CTRL2 | L100/ CTRL3 | Mean | SD | T-test p-value | Accession | Gene Symbol |
|---|---|---|---|---|---|---|---|
| | | | | | | 0A0U1RQD8;A0 A0U1RRG0 | |
| -0.4578 | -0.9144 | *0.4435* | -0.3096 | 0.6910 | 0.5189 | P14625;A0A1W2 PRR1;Q96GW1; H0YIV0;F8W026; F8VWL7 | HSP90B1 |
| -0.3253 | 0.3990 | *-1.0395* | -0.3219 | 0.7193 | 0.5193 | K7ERF1;Q9UBQ 5;K7ES31 | EIF3K |
| -0.2186 | -0.0143 | *0.0465* | -0.0621 | 0.1388 | 0.5194 | P11908;H7C540; A6NMS2;D3YTJ7 | PRPS2 |
| 0.2295 | -0.0579 | *-1.1429* | -0.3238 | 0.7238 | 0.5196 | Q13724;C9J8D4 | MOGS |
| 0.7119 | -1.4778 | *-0.7268* | -0.4975 | 1.1127 | 0.5197 | O14529 | CUX2 |
| 0.8325 | -0.1317 | *-4.5491* | -1.2828 | 2.8696 | 0.5198 | Q15643;H0YJ97 | TRIP11 |
| 1.1339 | -0.4230 | *0.3313* | 0.3474 | 0.7786 | 0.5204 | Q14584 | ZNF266 |
| 0.1180 | 0.8105 | *-5.7308* | -1.6008 | 3.5935 | 0.5211 | Q12851;C9JCU6; F2Z2B3;F8WCP4 | MAP4K2 |
| -0.2545 | -4.6525 | *0.9610* | -1.3153 | 2.9533 | 0.5211 | Q3T906;H0YIE6 | GNPTAB |
| -0.6937 | 0.4705 | *-0.6633* | -0.2955 | 0.6635 | 0.5212 | E7EWN3;Q9C0A 6;C9JLA7 | SETD5 |
| -0.9336 | -0.2159 | *0.3135* | -0.2787 | 0.6259 | 0.5212 | Q9BXM9;A0A0A 0MRS5;A0A0C4 DG97;F8W946;Q 8N450;C9JD05 | FSD1L |
| 1.0025 | -1.2424 | *3.2183* | 0.9928 | 2.2304 | 0.5213 | Q96CS2 | HAUS1 |
| 0.4493 | -0.2799 | *-1.4409* | -0.4238 | 0.9533 | 0.5218 | Q9P219 | CCDC88C |
| -0.8817 | 0.3455 | *-0.2822* | -0.2728 | 0.6136 | 0.5218 | P61224;A8KAH9; E7ESV4;F5GX62 ;F5H7Y6;P62834; A6NIZ1;F5GYB5; F5H004;F5H6R7; F5H0B7;F5H500; A0A075B6Q0;F5 GWU8;F5H077;F 5H491;B7ZB78;F 5H823;F5GYH7; F5H4H0;F8WBC 0;F5GZG1 | RAP1B |
| 0.3657 | -0.8741 | *-0.3198* | -0.2761 | 0.6211 | 0.5219 | P17677 | GAP43 |
| -0.2762 | 0.4860 | *0.3258* | 0.1785 | 0.4019 | 0.5220 | Q6ZU15 | SEPT14 |
| 0.5356 | -0.0042 | *-3.2811* | -0.9166 | 2.0654 | 0.5225 | Q99873;E9PKG1 ;E9PIX6;E9PQ98 ;H0YDE4;E9PNR 9 | PRMT1 |
| -1.2427 | -0.8119 | *0.6982* | -0.4521 | 1.0193 | 0.5226 | Q86UV5 | USP48 |
| 0.4078 | -0.1404 | *0.0983* | 0.1219 | 0.2749 | 0.5227 | B1AK53;A0A1B0 GUN9;A0A2R8Y 6D3 | ESPN |
| -0.5748 | -0.4390 | *0.3499* | -0.2213 | 0.4993 | 0.5230 | P11021;O95399; Q5H8X8 | HSPA5 |
| 1.3054 | 2.3401 | *-1.2149* | 0.8102 | 1.8285 | 0.5230 | J3QLE5;P14678; P63162;J3KRY3; S4R3P3 | SNRPN |
| -0.3115 | 0.0273 | *0.0252* | -0.0863 | 0.1950 | 0.5233 | Q13867;K7ES02; K7ESE8;J3KSD8 ;J3KS79;K7ENH5 | BLMH |

| L100/ CTRL1 | L100/ CTRL2 | L100/ CTRL3 | Mean | SD | T-test p-value | Accession | Gene Symbol |
|---|---|---|---|---|---|---|---|
| 0.2403 | -0.2953 | *-0.4015* | -0.1522 | 0.3440 | 0.5237 | Q9UNA1 | ARHGAP26 |
| -0.0964 | 0.4926 | *-2.4916* | -0.6985 | 1.5806 | 0.5240 | Q9Y5B6 | PAXBP1 |
| -0.0991 | -1.5882 | *0.3452* | -0.4474 | 1.0127 | 0.5241 | P17600;A0A1W2PS00 | SYN1 |
| 1.0077 | 0.5544 | *-0.5216* | 0.3468 | 0.7855 | 0.5244 | Q15276;A0A087WZZ3 | RABEP1 |
| -0.7978 | 0.9669 | *-2.3944* | -0.7418 | 1.6813 | 0.5246 | Q99714;Q5H928 | HSD17B10 |
| -0.2292 | -0.7575 | *0.2923* | -0.2314 | 0.5249 | 0.5248 | C9JVB6;Q7L0Y3 | TRMT10C |
| -0.4954 | 0.4995 | *1.0063* | 0.3368 | 0.7639 | 0.5249 | Q9BVS4 | RIOK2 |
| -0.1469 | 0.0562 | *0.6037* | 0.1710 | 0.3883 | 0.5252 | P0DMV8;A0A0G2JIW1;P0DMV9;V9GZ37 | HSPA1A |
| -0.2680 | 0.4299 | *-1.3398* | -0.3926 | 0.8914 | 0.5252 | P00451 | F8 |
| -1.1862 | -0.4973 | *0.5378* | -0.3819 | 0.8678 | 0.5255 | Q99259 | GAD1 |
| 0.1140 | 0.1637 | *-1.5962* | -0.4395 | 1.0020 | 0.5268 | P35527;K7EQQ3 | KRT9 |
| -0.2206 | -0.8073 | *0.2994* | -0.2428 | 0.5537 | 0.5268 | P11413;E7EUI8;E7EM57;E9PD92 | G6PD |
| -0.2093 | -0.0632 | *1.5721* | 0.4332 | 0.9890 | 0.5273 | P01112 | HRAS |
| 0.0819 | 0.0045 | *-0.5037* | -0.1391 | 0.3181 | 0.5279 | P67809;H0Y449;C9J5V9;A0A0D9SEI8 | YBX1 |
| -0.2404 | -0.6527 | *0.2801* | -0.2043 | 0.4674 | 0.5280 | Q13283;E5RIZ6;E5RH42;E5RI46;E5RIF8;E5RJU8 | G3BP1 |
| 1.0833 | -1.2383 | *-1.8949* | -0.6833 | 1.5648 | 0.5284 | Q13042 | CDC16 |
| -0.2928 | 0.6009 | *-2.1314* | -0.6077 | 1.3931 | 0.5288 | P53621 | COPA |
| -0.3423 | 0.0328 | *1.8109* | 0.5005 | 1.1503 | 0.5297 | O75820 | ZNF189 |
| -0.3049 | 0.1858 | *-0.2242* | -0.1144 | 0.2631 | 0.5299 | D3YTB5;P51617 | IRAK1 |
| -0.2919 | -0.5445 | *0.2832* | -0.1844 | 0.4242 | 0.5301 | F8VP71;F8W1Z2;Q6P1Q0 | LETMD1 |
| -0.3380 | 0.2398 | *0.9177* | 0.2732 | 0.6285 | 0.5301 | O95487 | SEC24B |
| 0.6516 | -0.0937 | *-3.2449* | -0.8957 | 2.0683 | 0.5314 | J3KPQ0;P22455;H0Y9P2;F8W9L4;H0YE20;P22607 | FGFR4 |
| -0.2375 | -0.3618 | *0.2095* | -0.1300 | 0.3004 | 0.5319 | P49321;Q5T624;E9PRH9;E9PPR5;H0YF33;H0YDS9;E9PI86;E9PNB5;E9PPQ8;E9PAU3;E9PJQ2;E9PKR5;E9PQG5 | NASP |
| 0.5603 | 1.1171 | *-0.5653* | 0.3707 | 0.8571 | 0.5319 | Q86VW0;H7BZR8 | SESTD1 |
| -0.3740 | -0.4645 | *0.2982* | -0.1801 | 0.4167 | 0.5321 | P22314;Q5JRR6;Q5JRR9;Q5JRS2;Q5JRS3;C9IYS6;C9J1V2;C9J4H5;J3QT59;K7EMJ7;Q13275;Q8IZL9 | UBA1 |
| -0.3961 | -0.7664 | *0.3942* | -0.2561 | 0.5928 | 0.5324 | E9PDF1;Q8TF17 | SH3TC2 |
| 0.6635 | -1.1293 | *3.4688* | 1.0010 | 2.3175 | 0.5324 | O94986 | CEP152 |
| 0.8704 | -1.5607 | *-0.9478* | -0.5460 | 1.2644 | 0.5324 | Q12931;I3L0K7;I3L239;I3L2D5 | TRAP1 |

| L100/ CTRL1 | L100/ CTRL2 | L100/ CTRL3 | Mean | SD | T-test p-value | Accession | Gene Symbol |
|---|---|---|---|---|---|---|---|
| 0.1713 | -0.3925 | *-0.1448* | -0.1220 | 0.2826 | 0.5326 | P61764;A0A1B0GVQ5;A0A1B0GWF2;A0A0D9SG72;A0A1B0GTP9;A0A2R8Y5D4;A0A1B0GW76;A0A0D9SFQ7;A0A0D9SFW6;A0A0D9SEH5;A0A0D9SEP9;A0A096LP33;A0A096LP52 | STXBP1 |
| 0.3732 | 1.5634 | *-10.6535* | -2.9056 | 6.7362 | 0.5329 | H3BQZ9;P07741;H3BQB1;H3BQF1;H3BSW3 | APRT |
| -0.1094 | 0.0239 | *0.5003* | 0.1382 | 0.3206 | 0.5330 | A0A0A0MRE6;O94967;E9PN15;E9PNF6;E9PR96 | WDR47 |
| -0.0617 | 0.0726 | *0.1018* | 0.0376 | 0.0872 | 0.5330 | Q9ULL1 | PLEKHG1 |
| 0.5612 | 0.2533 | *-4.4369* | -1.2075 | 2.8010 | 0.5331 | Q14126;J3KSI6 | DSG2 |
| -0.5989 | 0.4188 | *-0.5684* | -0.2495 | 0.5789 | 0.5332 | Q9BZZ5;H0YER7;E9PQK6;G3V1C3 | API5 |
| 0.0952 | 0.2183 | *-0.1039* | 0.0699 | 0.1626 | 0.5342 | P51003;G3XAH6;A0A0C4DGK1;G3V3I9;H0YJ00 | PAPOLA |
| -1.0880 | 0.0027 | *6.0200* | 1.6449 | 3.8280 | 0.5343 | Q9NPI1 | BRD7 |
| -0.1957 | -0.0233 | *1.1985* | 0.3265 | 0.7600 | 0.5344 | P63267;P62736;C9JFL5;F6QUT6;F6UVQ4;B8ZZJ2;F8WB63;F8WCH0 | ACTG2 |
| 0.0911 | -0.1697 | *0.5438* | 0.1551 | 0.3610 | 0.5344 | Q9UBQ7 | GRHPR |
| 1.0778 | -0.2766 | *0.0992* | 0.3001 | 0.6992 | 0.5347 | P48506;A0A0C4DGB2;A0A2R8YEL6;B4E2I4;E1CEI4;D6RGF8 | GCLC |
| -0.7721 | -0.0034 | *4.2690* | 1.1645 | 2.7160 | 0.5351 | Q9UBK8;H0Y963 | MTRR |
| 1.4302 | 0.2096 | *-0.4264* | 0.4045 | 0.9435 | 0.5351 | Q5T7W7;Q5T7X0 | TSTD2 |
| 0.4673 | 4.7240 | *-1.2420* | 1.3164 | 3.0723 | 0.5353 | H0YME5;Q9P2K8 | EIF2AK4 |
| -0.6044 | -2.1219 | *0.8300* | -0.6321 | 1.4761 | 0.5356 | Q9H7Z3;G3V338;H0YJT6 | NRDE2 |
| 0.2101 | -0.6983 | *2.8689* | 0.7936 | 1.8538 | 0.5357 | O95352;C9J415;C9JE55;C9JFF4;C9JGL2;C9JKA3;C9JNU2;F8WDY9 | ATG7 |
| 0.2483 | -0.9295 | *-0.0961* | -0.2591 | 0.6056 | 0.5358 | O60229;H7BXZ5;J3QSW6;B1AKS2;C9IZQ6;C9J3U7;C9JR33;F8WCF4;F8WDM4;F8WF57 | KALRN |
| -0.0298 | -0.0652 | *0.5050* | 0.1367 | 0.3194 | 0.5359 | O00299 | CLIC1 |

| L100/ CTRL1 | L100/ CTRL2 | L100/ CTRL3 | Mean | SD | T-test p-value | Accession | Gene Symbol |
|---|---|---|---|---|---|---|---|
| 0.0622 | 0.0800 | *-0.7492* | -0.2023 | 0.4737 | 0.5365 | Q9Y265;B5BUB1;E7ETR0;H7C4G5;J3QLR1;H7C4I3 | RUVBL1 |
| -0.0895 | 0.1290 | *-0.3383* | -0.0996 | 0.2338 | 0.5375 | P04264 | KRT1 |
| -0.2323 | -0.1170 | *1.8221* | 0.4909 | 1.1543 | 0.5380 | P20073 | ANXA7 |
| 0.7605 | -1.2785 | *3.7167* | 1.0662 | 2.5116 | 0.5387 | P48147 | PREP |
| 7.4291 | 1.4011 | *-2.4751* | 2.1184 | 4.9909 | 0.5388 | P58546 | MTPN |
| -0.5968 | -0.2242 | *0.2684* | -0.1842 | 0.4340 | 0.5388 | Q8WUM4;C9IZF9;F8WBR8;F8WEQ7 | PDCD6IP |
| 0.4426 | -1.0786 | *-0.3324* | -0.3228 | 0.7607 | 0.5388 | O60763 | USO1 |
| 0.3854 | 0.1747 | *-2.8988* | -0.7796 | 1.8383 | 0.5391 | Q14118 | DAG1 |
| -0.1983 | 0.2178 | *-0.4453* | -0.1420 | 0.3351 | 0.5395 | Q14103;H0Y8G5;D6RAF8;H0YA96;D6RF44;D6RBQ9;D6RD83 | HNRNPD |
| -2.3723 | 3.0131 | *-7.0130* | -2.1241 | 5.0177 | 0.5397 | Q71F56;H0YIQ9 | MED13L |
| 0.6111 | -1.8496 | *-0.3373* | -0.5253 | 1.2411 | 0.5398 | Q96FZ7;I3L4A1;I3L4G8;I3L3E4 | CHMP6 |
| -0.1771 | -1.8441 | *0.4937* | -0.5092 | 1.2038 | 0.5400 | Q9NP73;A0A087WTT9;A0A087WX43;A0A096LNJ4;A0A096LP10;A0A1B0GVW6;D6RE84 | ALG13 |
| -0.3350 | -0.2576 | *0.2156* | -0.1257 | 0.2981 | 0.5412 | P62310 | LSM3 |
| -0.4447 | 0.4289 | *0.8510* | 0.2784 | 0.6608 | 0.5415 | Q86UW7;A0A087X1P3;C9IYE1;F8W8P5 | CADPS2 |
| 1.9245 | 1.5347 | *-1.2629* | 0.7321 | 1.7387 | 0.5417 | A0A1B0GUS3;K7EQ95;A0A1B0GUM7;K7EKF7;A0A087WW63;A0A1B0GTI4;A0A1B0GTN7;A0A1B0GU74;A0A1B0GU81;A0A1B0GVQ9;A0A1C7CYY9;A0A384DVW2;B5TYJ1;O00555;A0A1B0GTW2 | CACNA1A |
| -0.1027 | -0.0240 | *0.6446* | 0.1726 | 0.4107 | 0.5423 | P54296 | MYOM2 |
| 0.2640 | 0.1287 | *-1.9684* | -0.5253 | 1.2516 | 0.5429 | P49961;A0A0U1RQZ5;A0A1W2PQK8 | ENTPD1 |
| 0.2862 | -0.3975 | *0.9752* | 0.2880 | 0.6864 | 0.5430 | Q96AE7 | TTC17 |
| -0.1906 | 0.7399 | *-2.9776* | -0.8094 | 1.9345 | 0.5439 | Q08945;E9PMD4 | SSRP1 |
| -0.0133 | -1.1193 | *0.2300* | -0.3009 | 0.7192 | 0.5440 | Q8TER5;G3V3N2;G3V5C1 | ARHGEF40 |
| -0.5566 | -1.1817 | *0.6027* | -0.3785 | 0.9054 | 0.5443 | Q96J66;H3BRJ2 | ABCC11 |
| 0.1819 | 0.3710 | *-2.7412* | -0.7294 | 1.7448 | 0.5443 | Q16720 | ATP2B3 |

| L100/ CTRL1 | L100/ CTRL2 | L100/ CTRL3 | Mean | SD | T-test p-value | Accession | Gene Symbol |
|---|---|---|---|---|---|---|---|
| 0.2056 | 1.0724 | *-0.3689* | 0.3030 | 0.7256 | 0.5446 | Q6P158;H7C109; A0A087WZ11;H7 BZ23 | DHX57 |
| -1.2585 | 1.0304 | *-1.5359* | -0.5880 | 1.4084 | 0.5447 | Q86X76 | NIT1 |
| 0.6684 | 0.9999 | *-0.6074* | 0.3536 | 0.8486 | 0.5454 | P00352 | ALDH1A1 |
| 0.1845 | 3.0664 | *-0.7608* | 0.8300 | 1.9936 | 0.5457 | J3KTH2;Q92750 | TAF4B |
| 1.8186 | -0.1165 | *-0.2544* | 0.4826 | 1.1591 | 0.5457 | Q9NTJ3;E9PD53 ;C9JR83;C9IYK2; C9JJ64;C9JWF0 | SMC4 |
| -0.0877 | 0.2139 | *-0.7237* | -0.1991 | 0.4786 | 0.5460 | Q9H078;A0A2U3 TZY2;A0A2R8Y6 R5;F5GX99;H0Y GM0;A0A2R8Y7 E8;A0A2R8Y602; F5H7A5;A0A2R8 YDH5 | CLPB |
| -0.1958 | 1.3111 | *-0.0747* | 0.3469 | 0.8372 | 0.5475 | O94887;H7C210 | FARP2 |
| -0.2642 | -0.9165 | *0.3770* | -0.2679 | 0.6468 | 0.5476 | P02545;Q5TCI8; A0A0C4DGC5;Q 3BDU5 | LMNA |
| 0.1517 | -0.1950 | *0.4344* | 0.1304 | 0.3152 | 0.5482 | Q7KZF4;H7C597 | SND1 |
| -0.5692 | 0.5598 | *-0.9782* | -0.3292 | 0.7966 | 0.5484 | Q86US8;I3L176;I 3L1W8;I3L355;I3 L3A9;I3L421 | SMG6 |
| -0.2925 | 0.5030 | *0.3022* | 0.1709 | 0.4137 | 0.5485 | K7EM38 | ACTG1 |
| -0.1782 | 0.7322 | *0.0358* | 0.1966 | 0.4760 | 0.5487 | P52565;J3KTF8; J3KRY1;J3KRE2; J3KS60 | ARHGDIA |
| -0.3460 | 0.3262 | *0.6466* | 0.2089 | 0.5066 | 0.5492 | P17980;R4GNH3 ;E9PM69;E9PKD 5;E9PMD8;E9PN 50;E9PLG2 | PSMC3 |
| 0.6430 | 0.4657 | *-0.4110* | 0.2325 | 0.5643 | 0.5494 | Q9UBE0;M0QZS 6;B3KNJ4;M0QX 65;M0QYM8;M0 R054;M0R375 | SAE1 |
| -1.0907 | -0.1274 | *0.3245* | -0.2978 | 0.7228 | 0.5495 | Q5JU85;A0A1W2 PR28;A0A1W2P PE7;A0A1W2PP U7;A0A1W2PQS 2;A0A1W2PQ34; A0A1W2PRJ5 | IQSEC2 |
| -0.1550 | -1.0733 | *0.3412* | -0.2957 | 0.7177 | 0.5495 | E7EUS2;Q8TBZ0 ;D6RDN4;D6RB1 8;D6RCR3 | CCDC110 |
| 0.0686 | 0.3974 | *-2.2422* | -0.5921 | 1.4385 | 0.5499 | Q14674 | ESPL1 |
| -0.2682 | 0.7346 | *-2.5269* | -0.6868 | 1.6706 | 0.5503 | Q5TDH0 | DDI2 |
| 0.1254 | -0.1917 | *-0.1427* | -0.0696 | 0.1707 | 0.5531 | Q9BV73;E7ETF9 ;H0Y5R2;E9PHT 2;H7C0D6 | CEP250 |
| 1.4211 | -1.4828 | *-2.3569* | -0.8062 | 1.9778 | 0.5533 | Q2TAL8;C9JAL2; C9JIA8 | QRICH1 |
| -0.3820 | 0.2938 | *-0.3917* | -0.1600 | 0.3930 | 0.5538 | Q4V328;A0A075 B793;A0A087WT 45;A0A087WXA6 | GRIPAP1 |

| L100/ CTRL1 | L100/ CTRL2 | L100/ CTRL3 | Mean | SD | T-test p-value | Accession | Gene Symbol |
|---|---|---|---|---|---|---|---|
| | | | | | | ;A0A087WW92;A0A087WZF5 | |
| -0.0028 | -0.9035 | *0.1927* | -0.2379 | 0.5847 | 0.5540 | A6QL64;A0A1W2PNQ7 | ANKRD36 |
| -0.4866 | 0.4777 | *-0.8053* | -0.2714 | 0.6680 | 0.5545 | P41252;A0A0A0MSX9;J3KR24;Q5TCD1;Q5TCC6 | IARS |
| -1.1511 | 1.1405 | *-1.9363* | -0.6490 | 1.5987 | 0.5548 | P50402;Q5HY57 | EMD |
| 0.1563 | -0.2158 | *0.4883* | 0.1429 | 0.3522 | 0.5550 | P62263;A0A2R8Y811;E5RH77 | RPS14 |
| 0.0414 | 0.3391 | *-1.7668* | -0.4621 | 1.1396 | 0.5552 | Q02790;F5H1U3;H0YFG2;F5H120 | FKBP4 |
| 0.7712 | -0.0007 | *-0.1634* | 0.2024 | 0.4993 | 0.5554 | Q9UBB9;F6SQZ1;F6UKU9;F6XM96 | TFIP11 |
| 0.9900 | 0.8833 | *-8.4763* | -2.2010 | 5.4348 | 0.5557 | Q8N8A6 | DDX51 |
| 0.8163 | -0.5773 | *-1.8729* | -0.5446 | 1.3449 | 0.5557 | Q7Z4H8;H0YEM3 | KDELC2 |
| -0.1121 | 1.2007 | *-0.1532* | 0.3118 | 0.7701 | 0.5557 | Q8N668;H0Y7J4;H7C377 | COMMD1 |
| 0.2325 | -0.3934 | *1.0225* | 0.2872 | 0.7095 | 0.5558 | Q9NXZ1;F5H2Z8 | SAGE1 |
| 0.5556 | 0.1021 | *-3.0111* | -0.7844 | 1.9416 | 0.5565 | P50851;H0YA17;H0YAC6 | LRBA |
| -0.7092 | 2.8632 | *0.1134* | 0.7558 | 1.8709 | 0.5565 | A0A1B0GUA3;Q96EK5;A0A1B0GUH6;A0A0D9SFK7;A0A1B0GTE6;A0A1B0GV79;A0A1B0GW21 | KIF1BP |
| 0.0372 | -0.3378 | *0.0380* | -0.0875 | 0.2167 | 0.5566 | P42285;H0Y8U3 | MTREX |
| -0.5172 | -0.2221 | *0.2628* | -0.1588 | 0.3938 | 0.5571 | Q06830;A0A0A0MSI0;A0A0A0MRQ5 | PRDX1 |
| 0.6579 | -0.1079 | *-2.6343* | -0.6948 | 1.7228 | 0.5572 | Q8WVY7 | UBLCP1 |
| -0.1957 | 0.3849 | *0.1646* | 0.1180 | 0.2931 | 0.5579 | Q8WWI1;J3KP06;F8WD26;A0A0A0MTE2;E9PMT2;E9PMS6;E9PMP7;H0Y424;A0A140T9Y7;H0YDG6;H0YDQ3;H0YE95 | LMO7 |
| 2.8460 | 6.4429 | *-3.3331* | 1.9853 | 4.9445 | 0.5587 | H0YJ82 | ABCD4 |
| 0.2920 | -0.6111 | *1.7583* | 0.4797 | 1.1958 | 0.5590 | Q14BN4;H7BZK0;B7Z964;H7C3M8;C9JA20;H7C5G9 | SLMAP |
| 0.0113 | -0.7328 | *0.1504* | -0.1904 | 0.4749 | 0.5593 | Q460N5 | PARP14 |
| 0.0016 | -0.2152 | *0.0459* | -0.0559 | 0.1397 | 0.5599 | P07355;H0YMD0;H0YMU9;H0YN42;H0YM50;H0YKS4;H0YL33;H0YMM1;H0YN28;H0YNP5;H0YKL9;H0YKV8;H0YKX9;H0YKZ7;H0YLV6; | ANXA2 |

| L100/ CTRL1 | L100/ CTRL2 | L100/ CTRL3 | Mean | SD | T-test p-value | Accession | Gene Symbol |
|---|---|---|---|---|---|---|---|
| | | | | | | H0YMT9;H0YMW4;H0YNA0;H0YMD9;H0YN52;H0YKN4;H0YLE2;H0YNB8 | |
| -0.8794 | -0.0423 | *0.2286* | -0.2310 | 0.5776 | 0.5601 | Q9NRF8 | CTPS2 |
| 0.0131 | -0.1160 | *0.4761* | 0.1244 | 0.3113 | 0.5605 | Q5T011 | SZT2 |
| -0.1549 | 0.3828 | *-1.1762* | -0.3161 | 0.7919 | 0.5608 | Q96H40 | ZNF486 |
| 0.2946 | -0.2924 | *0.4801* | 0.1608 | 0.4032 | 0.5612 | Q8N8N7;G3V3Y1;G3V2R9 | PTGR2 |
| 0.3209 | -0.2968 | *0.4553* | 0.1598 | 0.4011 | 0.5615 | Q5THK1;C9J9V0 | PRR14L |
| 0.1293 | 2.4323 | *-0.6482* | 0.6378 | 1.6020 | 0.5617 | Q9NWQ4;G3V553;G3V290;G3V5D0 | GPATCH2L |
| -0.2635 | 0.5704 | *-1.6386* | -0.4439 | 1.1155 | 0.5619 | Q14974;J3KTM9;J3QR48;J3QRG4 | KPNB1 |
| -0.3939 | 0.5665 | *-1.2652* | -0.3642 | 0.9163 | 0.5623 | Q8NBT2 | SPC24 |
| 1.1142 | 0.6805 | *-0.6808* | 0.3713 | 0.9366 | 0.5632 | P00390;A0A2R8YF59;H0YC68 | GSR |
| 1.5269 | 0.0685 | *-0.4020* | 0.3978 | 1.0057 | 0.5641 | H3BT97 | MMP15 |
| -0.1740 | 0.7808 | *-2.8087* | -0.7340 | 1.8591 | 0.5647 | Q15020;F8VV04;F8VVK9;F8VZM2;F8W667;H0YHU8 | SART3 |
| -0.1750 | -1.2692 | *0.4262* | -0.3393 | 0.8596 | 0.5647 | Q9H4A5;Q5T5I6 | GOLPH3L |
| -0.6422 | 0.0117 | *0.1356* | -0.1649 | 0.4179 | 0.5648 | B8ZZM6;Q9NQX7 | ITM2C |
| -0.3609 | -0.7228 | *0.4037* | -0.2267 | 0.5752 | 0.5653 | O75964;E9PN17 | ATP5MG |
| 0.8349 | -0.5736 | *-1.8470* | -0.5285 | 1.3415 | 0.5654 | Q9BTU6 | PI4K2A |
| 0.1369 | -0.2214 | *-0.1366* | -0.0737 | 0.1872 | 0.5655 | Q9NYC9;E7EP17;J3QQK8;K7EMX3 | DNAH9 |
| 0.6361 | -0.4208 | *0.4472* | 0.2208 | 0.5636 | 0.5674 | Q9H2J7;F8VX16;F8WJN6 | SLC6A15 |
| -0.3271 | 0.1153 | *1.0148* | 0.2677 | 0.6838 | 0.5677 | P21266;A0A0A0MTN3;Q5T8R1;B9ZVX7;E7EWW9;H3BQT3;P09488 | GSTM3 |
| 0.1860 | -0.8792 | *0.0208* | -0.2241 | 0.5733 | 0.5681 | Q96JJ6;A0A087WWT0;F5H1L9 | JPH4 |
| 0.0280 | 0.1170 | *-0.6068* | -0.1539 | 0.3947 | 0.5691 | F5H4N6;Q9Y3R0;F5H3F9;F5H4P8;F5H4Q7;H0YGF1 | GRIP1 |
| 0.6754 | -0.2623 | *0.1370* | 0.1834 | 0.4706 | 0.5693 | F8W6N3;Q92560 | BAP1 |
| -0.5558 | 0.8095 | *0.6058* | 0.2865 | 0.7365 | 0.5699 | Q9H251;A0A0A0MQS6;A0A087X097;A0A087WYR8;A0A0A0MS94 | CDH23 |
| -0.5701 | -0.5142 | *0.4303* | -0.2180 | 0.5621 | 0.5710 | Q9UBT7 | CTNNAL1 |
| 0.6658 | -0.0567 | *-2.5999* | -0.6636 | 1.7153 | 0.5718 | Q9Y4D1 | DAAM1 |
| -0.6091 | 0.2376 | *-0.1213* | -0.1643 | 0.4250 | 0.5721 | P63244;J3KPE3;D6RAC2;D6RHH4;H0YAF8;D6RE | RACK1 |

| L100/ CTRL1 | L100/ CTRL2 | L100/ CTRL3 | Mean | SD | T-test p-value | Accession | Gene Symbol |
|---|---|---|---|---|---|---|---|
| | | | | | | E5;D6R9Z1;D6RFZ9;D6RAU2;D6RFX4;E9PD14;D6RBD0;D6R909;D6RF23;D6RGK8;D6RHJ5;H0Y9P0;H0Y8W2;H0YAM7;D6R9L0;H0Y8R5 | |
| 0.2794 | -0.0290 | *-1.0697* | -0.2731 | 0.7069 | 0.5723 | P98171;E7EQN5;E9PCM6;H7C2Z8;A0A0B4J1X7 | ARHGAP4 |
| -2.9920 | 0.5379 | *0.1981* | -0.7520 | 1.9473 | 0.5724 | E5RHN2;E5RK20;O96020;Q8WUE3 | CCNE2 |
| -1.1822 | 1.5413 | *-3.0150* | -0.8853 | 2.2926 | 0.5725 | Q13884 | SNTB1 |
| -0.0887 | 0.3081 | *0.0183* | 0.0792 | 0.2053 | 0.5728 | Q96AX1;A0A2R8YF87;H3BMM5;F5H2X5;A0A2R8Y5U3 | VPS33A |
| -0.2947 | 0.0778 | *0.9675* | 0.2502 | 0.6485 | 0.5728 | Q9Y3F4;H0YH33 | STRAP |
| 0.2040 | 0.9721 | *-0.3875* | 0.2629 | 0.6817 | 0.5729 | Q06210 | GFPT1 |
| 2.8228 | -3.9635 | *-3.1375* | -1.4261 | 3.7027 | 0.5734 | P04049 | RAF1 |
| -0.2363 | -0.8145 | *0.3684* | -0.2275 | 0.5915 | 0.5739 | P61964;V9GYQ5;V9GZ59 | WDR5 |
| -1.0044 | 0.4966 | *-0.3609* | -0.2896 | 0.7531 | 0.5740 | Q9GZX5 | ZNF350 |
| -0.5769 | 0.7144 | *-1.3300* | -0.3975 | 1.0339 | 0.5740 | P08183;E7EWT8 | ABCB1 |
| 0.0729 | -1.5177 | *0.3025* | -0.3808 | 0.9913 | 0.5743 | Q9Y3L5;F6U784;P10114 | RAP2C |
| -0.9574 | -1.1446 | *0.8397* | -0.4207 | 1.0956 | 0.5744 | D6RAR6;D6RHY4;Q14CZ7;D6RC07 | FASTKD3 |
| -0.1962 | -0.2041 | *1.5873* | 0.3957 | 1.0320 | 0.5750 | Q10571;H7C105 | MN1 |
| -1.6358 | 1.7175 | *2.4015* | 0.8277 | 2.1607 | 0.5753 | Q9ULV4;B4E3S0;H0YHL7;F8VRE9;F8VSA4;F8VTT6;F8VUX3;F8VVB7;F8W1H8 | CORO1C |
| 1.2473 | 1.8209 | *-1.2153* | 0.6176 | 1.6131 | 0.5754 | A0A1W2PPJ3;A0A286YEQ8;A0A286YF26;P35498;A0A1B0GVX7;A0A1B0GUX7 | SCN1A |
| 0.1652 | -0.1670 | *0.2587* | 0.0857 | 0.2237 | 0.5755 | Q9Y5S2;A0A0U1RRC3 | CDC42BPB |
| -0.1025 | -0.0815 | *0.7248* | 0.1802 | 0.4717 | 0.5761 | P04406;E7EUT5;Q8N7A1;G3V5V5;Q9UIQ6 | GAPDH |
| 0.0964 | -0.7249 | *2.6360* | 0.6691 | 1.7521 | 0.5763 | P62081;A0A2R8Y623;B5MCP9 | RPS7 |
| 0.5961 | -0.3426 | *0.2951* | 0.1829 | 0.4793 | 0.5766 | Q92830 | KAT2A |
| 0.0576 | -0.4583 | *0.0593* | -0.1138 | 0.2983 | 0.5767 | Q9P2D3 | HEATR5B |
| -0.6334 | -0.4450 | *0.4295* | -0.2163 | 0.5672 | 0.5768 | Q6R327 | RICTOR |
| -0.1987 | -0.0001 | *0.8101* | 0.2038 | 0.5344 | 0.5769 | P27695;G3V3M6;G3V5Q1;G3V359 | APEX1 |

| L100/ CTRL1 | L100/ CTRL2 | L100/ CTRL3 | Mean | SD | T-test p-value | Accession | Gene Symbol |
|---|---|---|---|---|---|---|---|
| | | | | | | ;G3V5M0;A0A0C4DGK8;G3V3C7;H7C4A8;G3V5D9;G3V3Y6 | |
| 1.0046 | 0.4780 | *-0.5674* | 0.3051 | 0.8001 | 0.5769 | Q76I76;F5H527;K7EKN7 | SSH2 |
| -0.0203 | 1.0412 | *-0.2396* | 0.2604 | 0.6850 | 0.5779 | Q96G01;A8MVZ6;F8VZX7;F8W056 | BICD1 |
| -0.0645 | 0.9124 | *-3.4522* | -0.8681 | 2.2906 | 0.5790 | E7ERS3;Q86VM9;H3BRN6 | ZC3H18 |
| -0.6139 | -0.4747 | *0.4386* | -0.2166 | 0.5717 | 0.5790 | P35268;K7EMH1;K7ERI7;K7EJT5;K7EKS7;K7ELC4;K7EP65 | RPL22 |
| 0.1713 | -0.0077 | *-0.6619* | -0.1661 | 0.4386 | 0.5792 | O14513;A0A0A0MS79;A0A0A0MSE4;H7C187;C9JYL7 | NCKAP5 |
| -0.1246 | 0.2920 | *-0.7824* | -0.2050 | 0.5417 | 0.5795 | Q13576;F5H7S7;E7EWC2;D6R939;E9PDT6;H0YA28 | IQGAP2 |
| 1.0352 | -0.5508 | *-2.4802* | -0.6653 | 1.7605 | 0.5800 | Q9C0D4;D6RDM9 | ZNF518B |
| -2.1310 | -0.5058 | *0.9122* | -0.5748 | 1.5228 | 0.5803 | Q12792;F8VS81 | TWF1 |
| 2.5836 | 2.0921 | *-1.8949* | 0.9269 | 2.4561 | 0.5804 | Q9NXV6 | CDKN2AIP |
| -1.0387 | -0.5952 | *0.6455* | -0.3295 | 0.8730 | 0.5804 | P52597;A0A1B0GW42 | HNRNPF |
| -1.0600 | 1.6969 | *0.9826* | 0.5398 | 1.4308 | 0.5805 | F8VW96;Q16527;F8VQR7 | CSRP2 |
| 1.8818 | 1.5289 | *-1.3836* | 0.6757 | 1.7921 | 0.5808 | Q03938 | ZNF90 |
| -0.2014 | 0.5249 | *0.0894* | 0.1376 | 0.3656 | 0.5813 | Q9BZ95 | NSD3 |
| 1.6244 | -2.3153 | *4.6166* | 1.3086 | 3.4767 | 0.5814 | Q16666;H3BM18;X6RHM1 | IFI16 |
| 0.0881 | 0.2030 | *-1.1115* | -0.2735 | 0.7280 | 0.5820 | Q9UF12;K7EJK5;S4R3D8 | PRODH2 |
| -0.1536 | -0.1164 | *1.0235* | 0.2512 | 0.6691 | 0.5823 | Q71F23;Q09GN1 | CENPU |
| -0.4067 | -0.0514 | *1.7781* | 0.4400 | 1.1724 | 0.5824 | Q9BUF5;K7EL29;K7ESM5 | TUBB6 |
| -0.4557 | 0.2531 | *-0.2009* | -0.1345 | 0.3590 | 0.5830 | P09417;B7Z415;D6RGG7;D6RHJ7;H0Y8F7 | QDPR |
| -0.1676 | -0.5828 | *0.2709* | -0.1598 | 0.4269 | 0.5832 | O15075;Q5VZY9 | DCLK1 |
| -0.2491 | 0.7317 | *-2.0923* | -0.5366 | 1.4338 | 0.5833 | Q9UG01;H0YAI8 | IFT172 |
| 0.3538 | -0.6512 | *-0.2723* | -0.1899 | 0.5075 | 0.5834 | O95834;K7EIK7;A0A0C4DGQ7;C9JRL6;K7EIM1;K7EKG3;K7EKU5;K7EQR0;K7ERR2;K7ERY9 | EML2 |
| -0.2727 | 0.2876 | *0.3822* | 0.1324 | 0.3540 | 0.5836 | O43761;Q96L30 | SYNGR3 |
| 0.7314 | 0.7971 | *-5.7431* | -1.4049 | 3.7572 | 0.5836 | G3XAG1;Q96ME7 | ZNF512 |
| 0.5288 | -0.4392 | *-0.9148* | -0.2751 | 0.7357 | 0.5837 | P12955 | PEPD |

| L100/ CTRL1 | L100/ CTRL2 | L100/ CTRL3 | Mean | SD | T-test p-value | Accession | Gene Symbol |
|---|---|---|---|---|---|---|---|
| -0.1622 | -0.0224 | *0.7084* | 0.1746 | 0.4676 | 0.5841 | Q12901;K7ELA0; K7ENV6;K7EJJ6 | ZNF155 |
| -0.0102 | 0.7527 | *-2.9010* | -0.7195 | 1.9274 | 0.5842 | Q6PUV4;D6R960 ;D6RGY3 | CPLX2 |
| 0.0658 | -0.7177 | *0.1255* | -0.1755 | 0.4705 | 0.5846 | O95757;E9PDE8; D6RJ96 | HSPA4L |
| -0.1995 | 0.2753 | *0.2124* | 0.0961 | 0.2579 | 0.5849 | O43175;A0A286 YF22;A0A2C9F2 M7;A0A286YFA2; A0A286YFM8;A0 A286YER3;A0A2 86YFB2;A0A286 YFC8;A0A286YF 34;A0A286YFF3; A0A286YFE1;A0 A286YFL2;A0A2 86YFK5 | PHGDH |
| 0.3985 | -0.2360 | *-0.8681* | -0.2352 | 0.6333 | 0.5859 | P25786;F5GX11; B4DEV8;F5H112 | PSMA1 |
| -0.8434 | -1.2635 | *0.8577* | -0.4164 | 1.1232 | 0.5866 | F8W7A7;Q5BJE1 ;J3KSU3;A0A0A0 MS97 | CCDC178 |
| 1.8122 | 0.5549 | *-8.8377* | -2.1569 | 5.8198 | 0.5867 | O60884;A0A087 WT48;I3L320 | DNAJA2 |
| -0.2733 | 0.4349 | *-0.9081* | -0.2488 | 0.6718 | 0.5869 | P83731;C9JNW5 ;C9JXB8 | RPL24 |
| 0.1442 | 0.1891 | *-1.2285* | -0.2984 | 0.8058 | 0.5870 | O60488 | ACSL4 |
| -0.2995 | -0.0365 | *0.1076* | -0.0761 | 0.2065 | 0.5884 | P07814;V9GYZ6 | EPRS |
| 0.4084 | -0.2164 | *-0.9318* | -0.2466 | 0.6706 | 0.5893 | Q9UBU7;B4DXK 0 | DBF4 |
| -1.0724 | 1.9283 | *-4.2708* | -1.1383 | 3.1001 | 0.5899 | G3V0E4;O75439 | PMPCB |
| -0.6764 | 1.4159 | *-3.3982* | -0.8862 | 2.4139 | 0.5899 | P51826;C9J622; C9J847;C9JC67; C9JZ66;C9JMS1; C9JUC4 | AFF3 |
| -2.6164 | 1.9055 | *-1.9662* | -0.8924 | 2.4447 | 0.5919 | Q2WGJ9 | FER1L6 |
| -1.7605 | -1.0530 | *1.1523* | -0.5537 | 1.5192 | 0.5924 | Q9P2D1 | CHD7 |
| 0.7708 | -0.0605 | *-0.1540* | 0.1854 | 0.5091 | 0.5926 | E5RFR8;E5RIU9; E5RJI3;Q8WUX9 | CHMP7 |
| -0.1152 | 0.0534 | *0.2753* | 0.0712 | 0.1959 | 0.5933 | Q9NVU0;A0A0C 4DH01;H3BNJ0 | POLR3E |
| -0.9441 | 0.4691 | *2.1742* | 0.5664 | 1.5614 | 0.5940 | Q16891;B9A067; C9J406;H7C463; A0A087WYS0 | IMMT |
| 0.7699 | -1.3348 | *2.8288* | 0.7546 | 2.0818 | 0.5942 | F8W7R3;Q9NVI1 ;H3BMG4;H3BN3 5;H3BT54 | FANCI |
| -0.4580 | 3.8686 | *-0.6412* | 0.9231 | 2.5525 | 0.5950 | Q86U06;G3XAP0 | RBM23 |
| 0.9893 | -0.5788 | *0.4532* | 0.2879 | 0.7970 | 0.5954 | P02533 | KRT14 |
| 0.3991 | 0.3867 | *-2.7485* | -0.6542 | 1.8137 | 0.5959 | Q9BX82 | ZNF471 |
| 1.5248 | 0.4527 | *-0.7476* | 0.4100 | 1.1368 | 0.5960 | P35637;H3BPE7 | FUS |
| -0.0376 | -0.2784 | *1.1318* | 0.2719 | 0.7543 | 0.5961 | C9J6P4;Q7Z2W4 ;H7C5K1 | ZC3HAV1 |
| -0.1965 | -0.0283 | *0.0762* | -0.0495 | 0.1376 | 0.5967 | Q9Y617 | PSAT1 |

| L100/ CTRL1 | L100/ CTRL2 | L100/ CTRL3 | Mean | SD | T-test p-value | Accession | Gene Symbol |
|---|---|---|---|---|---|---|---|
| 0.5497 | -0.2034 | *0.0656* | 0.1373 | 0.3817 | 0.5968 | Q01814;A0A2U3TZI3;A0A2U3U055;H0Y7S3;A0A2R8Y4R4;A0A2R8Y535 | ATP2B2 |
| -0.3236 | 0.0554 | *0.0381* | -0.0767 | 0.2140 | 0.5980 | Q9H2T7 | RANBP17 |
| -0.0113 | 0.2566 | *-0.8900* | -0.2149 | 0.5998 | 0.5982 | M0QXZ5;Q96B54 | ZNF428 |
| 0.3023 | -0.8806 | *2.3136* | 0.5785 | 1.6149 | 0.5983 | P46783 | RPS10 |
| 0.3428 | -0.0696 | *-1.0292* | -0.2520 | 0.7040 | 0.5985 | Q9P225;H7C3C3;Q5TCY1 | DNAH2 |
| -0.1023 | -0.3095 | *1.4343* | 0.3409 | 0.9526 | 0.5986 | P39023;H7C422;G5E9G0;H7C3M2;B5MCW2;F8WCR1 | RPL3 |
| 0.3353 | -0.2553 | *0.2656* | 0.1152 | 0.3228 | 0.5995 | Q9HCG1;M0QZI7 | ZNF160 |
| 0.9606 | 0.0397 | *-0.3028* | 0.2325 | 0.6534 | 0.6005 | O14647;B7Z3I4;A0A0D9SGK0;A0A1B0GTU9;A0A1B0GU59 | CHD2 |
| -0.6748 | -0.4679 | *0.4835* | -0.2197 | 0.6177 | 0.6006 | A2RUB6;C9JYB8;F8WCY0 | CCDC66 |
| 0.2809 | -0.5126 | *1.0831* | 0.2838 | 0.7978 | 0.6006 | Q08495 | DMTN |
| 0.1748 | 0.4228 | *-2.0340* | -0.4788 | 1.3526 | 0.6022 | Q9Y623 | MYH4 |
| -1.5218 | -0.2223 | *0.6058* | -0.3794 | 1.0725 | 0.6024 | Q04637;E7EUU4;E7EX73;E9PGM1;H7C044;H7C0V6 | EIF4G1 |
| -0.7812 | 1.1469 | *0.7066* | 0.3574 | 1.0104 | 0.6024 | Q3V6T2;A0A2R8Y4W8;A0A2R8Y4X4;A0A2R8YG73;A0A2R8Y820;A0A2R8Y885;H0Y470;H7C2C6;A0A087WXD9;A0A2R8YCU9;A0A2R8YG52;H0Y7U8;A0A2R8Y5D7;A0A2R8YGU1;A0A2R8YD81;A0A2R8Y6B2;A0A2R8Y7D9;A0A2R8Y7L3;A0A2R8Y846;A0A2R8YD99;A0A2R8Y7B1 | CCDC88A |
| 0.7349 | 0.1021 | *-2.8837* | -0.6822 | 1.9326 | 0.6031 | O60518;A0A096LPA6;A0A096LNS2 | RANBP6 |
| -0.1435 | 4.6282 | *-1.1976* | 1.0957 | 3.1043 | 0.6032 | Q92503;K7EJ08;K7ELM0;K7EPE4 | SEC14L1 |
| -0.4716 | -0.1033 | *0.2125* | -0.1208 | 0.3424 | 0.6032 | P22102;F8WD69;C9JBJ1;C9JKQ7;C9JZG2;H7C366 | GART |
| -0.5126 | -0.4820 | *3.3334* | 0.7796 | 2.2117 | 0.6036 | Q06136;K7ERC8;K7EQS7 | KDSR |
| 0.7458 | 0.5516 | *-4.3511* | -1.0179 | 2.8883 | 0.6037 | Q49AJ0;J3QSR3 | FAM135B |

| L100/ CTRL1 | L100/ CTRL2 | L100/ CTRL3 | Mean | SD | T-test p-value | Accession | Gene Symbol |
|---|---|---|---|---|---|---|---|
| 1.3251 | 0.7351 | *-0.8635* | 0.3989 | 1.1324 | 0.6039 | Q9BYP7;A0A087WYK2 | WNK3 |
| -0.9952 | -1.0579 | *0.8867* | -0.3888 | 1.1051 | 0.6043 | Q93084 | ATP2A3 |
| -0.6478 | -0.3633 | *3.3786* | 0.7892 | 2.2470 | 0.6049 | Q14576;K7EPB5 | ELAVL3 |
| -0.0222 | -2.6374 | *0.7795* | -0.6267 | 1.7869 | 0.6053 | A0A0G2JH68;A0A140T8Z0;A0A2R8Y5N1;H9KV28;O60610;A0A2R8YEF8;B4E2I7 | DIAPH1 |
| 0.4463 | -0.9956 | *2.2698* | 0.5735 | 1.6364 | 0.6056 | Q86WZ6;K7EIL7;K7ELS1 | ZNF227 |
| -0.1827 | 0.1374 | *-0.1365* | -0.0606 | 0.1730 | 0.6057 | Q10567;C9J1E7 | AP1B1 |
| 3.4597 | -3.2738 | *-4.8019* | -1.5387 | 4.3956 | 0.6060 | Q9C0E8;C9JM95;C9JL94 | LNPK |
| -0.3643 | 0.1555 | *0.8466* | 0.2126 | 0.6075 | 0.6060 | Q16537;B5BTZ7 | PPP2R5E |
| 1.0528 | 0.0612 | *-0.3566* | 0.2525 | 0.7239 | 0.6072 | Q5JWF2;P63092;A0A0A0MR13;S4R3E3;H0Y7F4;Q5JWE9;H0Y7E8;S4R3V9;Q5JWD1;A2A2R6 | GNAS |
| -0.3104 | 0.0376 | *0.9584* | 0.2286 | 0.6556 | 0.6073 | P21333;Q60FE5;A0A087WWY3;F8WE98;H0Y5C6;H0Y5F3;H7C2E7 | FLNA |
| 2.5747 | -0.7961 | *0.0539* | 0.6108 | 1.7531 | 0.6075 | Q05DH4 | FAM160A1 |
| -1.0910 | 0.8489 | *-0.8611* | -0.3677 | 1.0599 | 0.6089 | Q9NPH2;J3QS51;J3KRH4 | ISYNA1 |
| -0.4784 | 1.3577 | *-3.3456* | -0.8221 | 2.3704 | 0.6090 | H0YDU8;P53041;A8MU39 | PPP5C |
| -0.0046 | -0.2636 | *0.0814* | -0.0622 | 0.1796 | 0.6093 | Q4FZB7 | KMT5B |
| 0.0951 | -0.5030 | *0.0592* | -0.1162 | 0.3354 | 0.6094 | O00273;K7ERT1 | DFFA |
| -1.1485 | 0.1629 | *0.1904* | -0.2651 | 0.7652 | 0.6094 | P62244;I3L3P7;H3BVC7;H3BT37;H3BV27;I3L246;I3L303 | RPS15A |
| -0.2603 | -0.5957 | *0.3554* | -0.1669 | 0.4824 | 0.6099 | Q99666 | RGPD5 |
| -0.3617 | -0.4063 | *0.3366* | -0.1438 | 0.4167 | 0.6107 | Q14344 | GNA13 |
| -0.3172 | 0.8241 | *0.0914* | 0.1994 | 0.5783 | 0.6109 | Q9HBD1 | RC3H2 |
| 1.8318 | -1.9909 | *2.7627* | 0.8679 | 2.5192 | 0.6113 | Q9UJX5;D6RAP6 | ANAPC4 |
| 0.5040 | -0.1773 | *0.0333* | 0.1200 | 0.3488 | 0.6117 | P15880;E9PQD7;H0YEN5;E9PMM9;E9PM36;I3L404;E9PPT0;H0YE27;H3BNG3 | RPS2 |
| 0.4628 | -1.1060 | *-0.1700* | -0.2711 | 0.7893 | 0.6123 | Q8ND30;E9PP16;H0YDJ2;E9PK17;E9PMH3;E9PMN3;H0YDM0 | PPFIBP2 |
| 1.2418 | -0.7392 | *0.5313* | 0.3446 | 1.0036 | 0.6124 | Q8NEN0 | ARMC2 |
| -0.5704 | 1.6723 | *0.0856* | 0.3958 | 1.1531 | 0.6124 | P42574;A8MVM1;C9JXR7 | CASP3 |
| -0.6764 | 1.5143 | *0.2925* | 0.3768 | 1.0978 | 0.6125 | Q68CZ1;A0A087WX34;H3BS47;H | RPGRIP1L |

| L100/ CTRL1 | L100/ CTRL2 | L100/ CTRL3 | Mean | SD | T-test p-value | Accession | Gene Symbol |
|---|---|---|---|---|---|---|---|
| | | | | | | 3BV03;H3BPF5;H3BPS4;I3L1B5;I3L2P2;J3QLR9 | |
| -0.3441 | -0.2844 | *0.2770* | -0.1172 | 0.3427 | 0.6138 | Q15751 | HERC1 |
| 0.7054 | -0.4549 | *-1.2669* | -0.3388 | 0.9912 | 0.6139 | D6RCF1;P29973;D6R978 | CNGA1 |
| 0.2628 | -2.1087 | *0.4038* | -0.4807 | 1.4117 | 0.6151 | P78524;B4DDL8;E9PJC3 | ST5 |
| -0.2015 | -0.1410 | *1.0810* | 0.2461 | 0.7236 | 0.6154 | O15173 | PGRMC2 |
| 0.0094 | 1.9061 | *-6.1979* | -1.4275 | 4.2387 | 0.6187 | O75061;S4R305 | DNAJC6 |
| -0.8065 | 0.5749 | *-0.5015* | -0.2443 | 0.7257 | 0.6188 | Q14498;G3XAC6;H0Y4X3;A0A0U1RQH7;A0A0U1RQW2;Q5QP22;Q5QP23 | RBM39 |
| -0.7694 | -1.4322 | *0.9604* | -0.4138 | 1.2353 | 0.6205 | H3BQX2;H3BSA5;H3BTU5;Q9P2J9;H3BV50;H3BRB7 | PDP2 |
| -0.1951 | 0.3412 | *-0.6370* | -0.1636 | 0.4899 | 0.6214 | P53814;A0A087WVP4;A0A087X1R1;C9JGQ0;H7C372 | SMTN |
| 0.8388 | -1.1792 | *1.9129* | 0.5242 | 1.5699 | 0.6215 | Q01518;Q5T0R1;Q5T0R2;Q5T0R3;Q5T0R4;Q5T0R5;Q5T0R6;Q5T0R7;Q5T0S3;Q5T0R8;Q5T0R9 | CAP1 |
| 2.2020 | -6.1485 | *-0.3302* | -1.4256 | 4.2817 | 0.6224 | Q9NVR2;E5RHN1;E5RJN5 | INTS10 |
| -0.5210 | 0.2133 | *1.1337* | 0.2754 | 0.8291 | 0.6232 | Q86YW9;F8WAE6;H7C4I9 | MED12L |
| -0.4415 | 0.1550 | *-0.0185* | -0.1017 | 0.3068 | 0.6240 | Q15154;H0YBA1 | PCM1 |
| -0.0725 | 0.0407 | *0.1339* | 0.0340 | 0.1034 | 0.6259 | Q8NEZ4;H0Y765;H7BY37;H0YMU7;H7C2V8 | KMT2C |
| -0.0736 | -0.6460 | *0.2650* | -0.1515 | 0.4605 | 0.6262 | M0R2Z9;M0R3F6;Q8IX01 | SUGP2 |
| -0.7820 | -0.3940 | *0.5180* | -0.2193 | 0.6674 | 0.6266 | P11766;H0YAG8;D6RFE4;D6R9G2;D6RAY0 | ADH5 |
| -0.1847 | 1.7660 | *-0.4039* | 0.3925 | 1.1946 | 0.6267 | Q9H0Q3 | FXYD6 |
| -0.6090 | -1.6280 | *0.9554* | -0.4272 | 1.3013 | 0.6270 | K7EMM8;Q49A26 | GLYR1 |
| -1.7371 | -0.7949 | *1.1060* | -0.4753 | 1.4482 | 0.6270 | A0A1X7SC74;Q6ZTR7;A0A087WU51 | FAM92B |
| 0.1377 | -0.1097 | *-0.2000* | -0.0573 | 0.1748 | 0.6272 | P55060 | CSE1L |
| -0.5875 | 0.8541 | *0.4677* | 0.2448 | 0.7462 | 0.6273 | P0C0S5;Q71UI9;C9J0D1;C9J386 | H2AFZ |
| 1.2964 | 4.7387 | *-2.4819* | 1.1844 | 3.6116 | 0.6273 | E9PJT4 | DNHD1 |
| -0.1963 | 0.2283 | *-0.3116* | -0.0932 | 0.2843 | 0.6273 | Q8IUG5 | MYO18B |
| -0.8761 | 0.1516 | *0.1432* | -0.1937 | 0.5909 | 0.6274 | Q92870;G5E9Y1;H0YAJ5;D6RB00 | APBB2 |

| L100/ CTRL1 | L100/ CTRL2 | L100/ CTRL3 | Mean | SD | T-test p-value | Accession | Gene Symbol |
|---|---|---|---|---|---|---|---|
| -0.4684 | 0.3453 | *0.7203* | 0.1991 | 0.6077 | 0.6276 | Q6BDS2;H7C1J4 | UHRF1BP1 |
| -1.5916 | -1.0020 | *1.1665* | -0.4757 | 1.4524 | 0.6277 | Q15027;I3L0K9;I3L268 | ACAP1 |
| 0.1189 | 0.7503 | *-2.6271* | -0.5859 | 1.7956 | 0.6289 | Q68DK2;A0A2H2FF08 | ZFYVE26 |
| -0.6755 | 0.9963 | *0.5224* | 0.2811 | 0.8616 | 0.6290 | Q9NQ76;D6RAC8 | MEPE |
| -1.0060 | -0.3398 | *0.5713* | -0.2582 | 0.7918 | 0.6291 | M0R268;P09012 | SNRPA |
| -0.6151 | -0.0156 | *0.2126* | -0.1394 | 0.4275 | 0.6291 | Q8WXX5 | DNAJC9 |
| -0.4346 | -0.0273 | *0.1636* | -0.0994 | 0.3056 | 0.6299 | P30041 | PRDX6 |
| -0.6908 | -0.1694 | *0.3518* | -0.1695 | 0.5213 | 0.6301 | P47897;A0A1B0GVU9;B4DDN1;H7C0R3;A0A0U1RQX5;A0A0U1RQX9;A0A0U1RQT0;C9J165;A0A0U1RQE9;A0A0U1RRC8;A0A0U1RRI9;A0A0U1RQL2;A0A0U1RQM8;A0A0U1RQU2;A0A0U1RR66;A0A0U1RQJ6;J3QLM2 | QARS |
| -0.0340 | -0.3234 | *1.0788* | 0.2405 | 0.7403 | 0.6303 | A4D0V7;E7ENG7;H0YB19 | CPED1 |
| -0.2631 | -0.5761 | *0.3699* | -0.1564 | 0.4820 | 0.6306 | P04843;B7Z4L4;F8WF32;P04279 | RPN1 |
| 0.4887 | -0.2850 | *-0.8623* | -0.2195 | 0.6779 | 0.6313 | Q8N7H5;M0QYC7 | PAF1 |
| 2.4832 | 1.9534 | *-2.0393* | 0.7991 | 2.4724 | 0.6319 | Q9P2R3 | ANKFY1 |
| 0.0600 | -0.5786 | *0.1376* | -0.1270 | 0.3930 | 0.6320 | Q8IWJ2;B8ZZW2;B8ZZA5;U3KPU4;U3KQE0;E9PLS0;E9PNU3;E9PPL0;E9PQC8;E9PR23;E9PRA4;H7C010;Q71H61 | GCC2 |
| -0.5335 | 1.5979 | *-3.5872* | -0.8409 | 2.6062 | 0.6325 | Q9H972;G3V2P0;G3V4W6;J3KPV9;G3V2U2;G3V396;G3V3W0;G3V4L9 | C14orf93 |
| 1.8231 | -1.4583 | *-2.5769* | -0.7373 | 2.2869 | 0.6327 | Q6P0N0;G5E9K5;H7C1S6;H0YJM1 | MIS18BP1 |
| 0.3962 | -1.1715 | *2.6133* | 0.6127 | 1.9017 | 0.6330 | A0A2R8YD58;A0A2R8YF43;E7EU96;P68400;Q5U5J2;A0A2R8Y5A0;B5BUH5;A0A2R8Y4D6;A0A2R8Y4H0;A0A2R8YDP2;A0A2R8YFU2;V9GYA2;A0A2R8YEW1;A0A2R8YCK2;A0A2R8YDY7;V9GY80;A0A2R | CSNK2A1 |

| L100/ CTRL1 | L100/ CTRL2 | L100/ CTRL3 | Mean | SD | T-test p-value | Accession | Gene Symbol |
|---|---|---|---|---|---|---|---|
| | | | | | | 8Y3W6;A0A2R8Y797;A0A2R8Y7T1;A0A2R8YEL7;A0A2R8YF47 | |
| 1.5293 | -0.3086 | *-0.2193* | 0.3338 | 1.0363 | 0.6330 | O75030 | MITF |
| -0.0015 | -0.6821 | *0.2269* | -0.1522 | 0.4728 | 0.6332 | Q70CQ2;H7C183 | USP34 |
| 1.0139 | 0.2234 | *-0.5044* | 0.2443 | 0.7594 | 0.6334 | Q9HCJ0 | TNRC6C |
| 1.0006 | 2.8768 | *-1.6722* | 0.7351 | 2.2861 | 0.6336 | Q8TF32;Q96N22;M0R1H8;Q5JVG8 | ZNF431 |
| -0.2070 | -0.5823 | *0.3415* | -0.1492 | 0.4646 | 0.6339 | O95197;F5H617 | RTN3 |
| -0.3605 | 0.1886 | *-0.0926* | -0.0882 | 0.2746 | 0.6340 | Q96SZ5 | ADO |
| 0.0630 | 0.1206 | *-0.5309* | -0.1158 | 0.3606 | 0.6341 | Q13151 | HNRNPA0 |
| -4.4053 | 1.3305 | *0.1624* | -0.9708 | 3.0311 | 0.6348 | F5H7N0;P28472 | GABRB3 |
| -2.4471 | 1.3280 | *-0.6960* | -0.6051 | 1.8892 | 0.6348 | Q96T23;H0YCN2 | RSF1 |
| -0.1079 | -0.0584 | *0.0754* | -0.0303 | 0.0948 | 0.6355 | P62937;C9J5S7;F8WE65;E5RIZ5 | PPIA |
| -0.0820 | 1.6673 | *-0.4877* | 0.3658 | 1.1452 | 0.6356 | Q5SQI0 | ATAT1 |
| 1.1425 | -1.7030 | *-0.8361* | -0.4656 | 1.4585 | 0.6359 | P30038;Q5TF55 | ALDH4A1 |
| -0.2763 | 1.6674 | *-0.3080* | 0.3611 | 1.1315 | 0.6360 | Q9Y597 | KCTD3 |
| -0.1966 | 0.5600 | *0.0107* | 0.1247 | 0.3910 | 0.6361 | P46821;D6RA40;D6RGJ3;D6RCL2 | MAP1B |
| -0.5928 | -0.9926 | *0.7256* | -0.2866 | 0.8991 | 0.6363 | Q96FA7 | ZBED6CL |
| -0.2107 | -0.6110 | *0.3574* | -0.1547 | 0.4866 | 0.6371 | Q6ZMI0;F8WE40;H7C1Y1 | PPP1R21 |
| -0.1076 | 0.0589 | *0.1919* | 0.0477 | 0.1501 | 0.6371 | Q9NZI8 | IGF2BP1 |
| -1.5037 | 0.2013 | *0.3277* | -0.3249 | 1.0228 | 0.6374 | Q96BY6;A0A2R8YD85;H0YFC5 | DOCK10 |
| 1.6425 | 0.2770 | *-0.7678* | 0.3839 | 1.2087 | 0.6375 | A0A0U1RQC5;A0A0U1RRJ0;Q9Y4C0;A0A0A0MR89;G3V247;G3V4R9 | NRXN3 |
| 0.1869 | 0.8977 | *-0.4458* | 0.2129 | 0.6722 | 0.6383 | Q92526;J3KRI6 | CCT6B |
| -1.7523 | -0.3191 | *0.8433* | -0.4094 | 1.3001 | 0.6402 | Q9Y6X9;H7C1V1 | MORC2 |
| -0.3799 | -0.4762 | *0.4016* | -0.1515 | 0.4814 | 0.6403 | Q04206;A0A087X0W8;Q2TAM5;E9PKH5;A0A087WVP0;E9PI38;E9PKV4;E9PQS6;E9PMD5;E9PRX2;E9PJR1;E9PNV4;E9PJZ9;E9PM47;E9PN69;E9PNK5 | RELA |
| 0.3353 | 0.0701 | *-1.1563* | -0.2503 | 0.7957 | 0.6405 | E7EVD6;Q9HAR2;H0Y9K5 | ADGRL3 |
| 0.2485 | -1.1098 | *0.1473* | -0.2380 | 0.7567 | 0.6406 | O00443;A0A0C4DGF9;E9PPP3 | PIK3C2A |
| -1.0735 | -1.3893 | *1.1570* | -0.4353 | 1.3880 | 0.6415 | Q86YM6;Q86YM7 | HOMER1 |
| 0.9482 | 3.2392 | *-1.8095* | 0.7926 | 2.5279 | 0.6415 | B7WPN9;Q4VNC1;H0Y4Z2;H7C1P5 | ATP13A4 |

| L100/ CTRL1 | L100/ CTRL2 | L100/ CTRL3 | Mean | SD | T-test p-value | Accession | Gene Symbol |
|---|---|---|---|---|---|---|---|
| -0.2300 | -0.0886 | *0.8942* | 0.1919 | 0.6123 | 0.6417 | P10586;A2A437 | PTPRF |
| 0.3372 | 0.4223 | *-0.3575* | 0.1340 | 0.4278 | 0.6418 | P40123;E9PDI2; A0A087WZ15;A0A087X0J3;F8WDB9 | CAP2 |
| -0.3206 | 0.5454 | *-0.9131* | -0.2294 | 0.7335 | 0.6423 | A0A0A0MRE5;H0YBF7;Q9ULH1; E5RFD9 | ASAP1 |
| -0.3131 | -0.1586 | *0.2165* | -0.0851 | 0.2723 | 0.6427 | P52272;A0A087X0X3;M0R019;M0R2T0;M0R0N3;M0R2I7;M0QZM1; M0QYQ7;M0QY96;M0QYL3;M0R0Y6 | HNRNPM |
| 3.6130 | -0.7456 | *-0.5629* | 0.7682 | 2.4654 | 0.6435 | Q9Y6I9;C9J7N0; C9JHH5;C9JXQ7;A0A087WTU3;C9JL43;C9JUC3 | TEX264 |
| 0.1918 | 0.2466 | *-0.2070* | 0.0771 | 0.2476 | 0.6435 | P26232;A0A0A0MRI5;B9A010;A0A0A0MTJ6 | CTNNA2 |
| 0.7136 | 1.6601 | *-1.0758* | 0.4327 | 1.3894 | 0.6437 | Q969P6;E5RIC7; E5RFS0;E5RJ95 | TOP1MT |
| 0.5080 | -0.6793 | *0.9617* | 0.2635 | 0.8474 | 0.6441 | P33993 | MCM7 |
| -0.2369 | -0.9283 | *0.4999* | -0.2218 | 0.7142 | 0.6445 | P17483 | HOXB4 |
| 2.8987 | -1.2841 | *0.3490* | 0.6545 | 2.1081 | 0.6446 | Q9HBT8;A8MTT8;J3KRF9;J3KSJ1;J3KSW0;K7ENW2;K7EQ88 | ZNF286A |
| 0.5423 | -1.6789 | *3.6041* | 0.8225 | 2.6527 | 0.6450 | Q7Z6G8;F8VR14;H0YI72;H0YJZ1; R4GN78 | ANKS1B |
| 0.1221 | 0.2276 | *-0.9565* | -0.2023 | 0.6553 | 0.6464 | Q96PC5;G3V599;O15320;G3V5K6;A0A2R8Y569;A0A2R8YE69;A4D2H0;A4FU28;P0CG41;Q86UF2;Q8IX94;Q8IX95 | MIA2 |
| -0.3956 | -0.1276 | *0.2319* | -0.0971 | 0.3149 | 0.6466 | Q15293 | RCN1 |
| -0.2734 | 0.1055 | *0.5482* | 0.1268 | 0.4112 | 0.6467 | U3KQ37 | ZNF318 |
| 0.8475 | 1.1596 | *-0.9530* | 0.3514 | 1.1403 | 0.6469 | P30837 | ALDH1B1 |
| -0.2943 | 0.1408 | *0.5381* | 0.1282 | 0.4163 | 0.6471 | Q5T2S8;Q5T2S9 | ARMC4 |
| -2.1383 | -0.0332 | *0.7806* | -0.4636 | 1.5063 | 0.6473 | Q8NF50;A2A369; E9PDJ4;C9J7A3; E9PHZ4;F8WC95 | DOCK8 |
| -0.8716 | -0.8124 | *0.8058* | -0.2927 | 0.9518 | 0.6475 | P51784;G5E9A6; Q5JXD3;C9JBP8 | USP11 |
| 0.4313 | -0.0925 | *-1.0121* | -0.2244 | 0.7307 | 0.6479 | O60313;C9JMB8; A0A2R8YE78;A0A2R8YDM2;E5KLJ9;A0A2R8Y3X5;A0A2R8YGE5; A0A2R8YD53;A0 | OPA1 |

| L100/ CTRL1 | L100/ CTRL2 | L100/ CTRL3 | Mean | SD | T-test p-value | Accession | Gene Symbol |
|---|---|---|---|---|---|---|---|
| | | | | | | A2R8YFD1;A0A2R8Y4Q3;A0A2R8YE54;A0A2R8Y4G4;A0A2R8Y5G3;H7C3G2;C9JY58;H7C141 | |
| 0.3469 | 0.3907 | *-0.3531* | 0.1282 | 0.4174 | 0.6479 | Q9H2K8;G3V1Q8;F5GWV8;F5H3L7;F5H5E0 | TAOK3 |
| -0.1493 | 2.1296 | *-0.6245* | 0.4519 | 1.4722 | 0.6481 | O75113 | N4BP1 |
| 0.1274 | -0.2495 | *0.4391* | 0.1057 | 0.3448 | 0.6486 | P25054;E9PFT7;A0A2Q2SV78;E7EMH9;D6RFL6 | APC |
| -0.2745 | -0.3905 | *0.3167* | -0.1161 | 0.3793 | 0.6489 | P61978;Q5T6W2;S4R457;S4R359 | HNRNPK |
| -0.1824 | -1.4511 | *0.6600* | -0.3245 | 1.0627 | 0.6497 | Q14831 | GRM7 |
| -0.6953 | 1.2235 | *-2.0206* | -0.4974 | 1.6311 | 0.6501 | Q8TB72 | PUM2 |
| -0.2228 | 0.0127 | *0.5944* | 0.1281 | 0.4206 | 0.6506 | Q14568 | HSP90AA2P |
| 0.3731 | 0.4149 | *-2.1045* | -0.4388 | 1.4426 | 0.6509 | Q07092 | COL16A1 |
| 1.5929 | 4.9732 | *-2.9437* | 1.2075 | 3.9725 | 0.6511 | Q5VZ89 | DENND4C |
| 0.3838 | -1.5299 | *3.4158* | 0.7566 | 2.4938 | 0.6517 | Q9NQX4;H0YMK3;H0YM96 | MYO5C |
| -0.2779 | -0.5544 | *0.3905* | -0.1473 | 0.4858 | 0.6520 | P33176 | KIF5B |
| 0.8562 | -2.5820 | *0.0863* | -0.5465 | 1.8043 | 0.6522 | A0A0A0MR67;A0A0A0MSA1;E9PDI6;F5GY28;F5H522;Q13936;A0A087WZV3;F5H638 | CACNA1C |
| -0.1269 | -1.4736 | *0.6318* | -0.3229 | 1.0663 | 0.6523 | P05141 | SLC25A5 |
| -1.1083 | 2.2351 | *0.3913* | 0.5060 | 1.6747 | 0.6529 | Q96Q35 | ALS2CR12 |
| 1.4110 | 0.2379 | *-4.4553* | -0.9355 | 3.1042 | 0.6537 | Q9UHY7 | ENOPH1 |
| 1.1388 | -0.7404 | *-1.7063* | -0.4360 | 1.4468 | 0.6538 | P21399 | ACO1 |
| 1.3956 | -2.0661 | *-0.9232* | -0.5313 | 1.7638 | 0.6539 | P16870;C9JE88;D6R930;D6RF88;H0YAM0 | CPE |
| 0.7924 | -1.4935 | *2.5041* | 0.6010 | 2.0057 | 0.6555 | P06744;A0A0J9YXP8;A0A2R8Y6C7;K7EPY4;K7ERC6;K7EP41;A0A2R8YF08;K7ELR7;K7ENA0;A0A0J9YXM3;A0A0J9YXH9;A0A0J9YYI8;K7EIL4;K7ERK8 | GPI |
| -0.1962 | -0.7968 | *0.4385* | -0.1849 | 0.6178 | 0.6559 | Q9C0B0;K7EQ93 | UNK |
| -0.0639 | -0.2659 | *0.8775* | 0.1826 | 0.6102 | 0.6559 | P61026 | RAB10 |
| -1.9268 | 0.1251 | *0.5983* | -0.4011 | 1.3423 | 0.6563 | Q5SNV9;H7BXQ2;H0Y5F2 | C1orf167 |
| -0.3931 | -0.7229 | *0.5335* | -0.1942 | 0.6514 | 0.6571 | P55072;C9IZA5;C9JUP7;Q5JRK0 | VCP |
| -0.0599 | -3.7606 | *1.4335* | -0.7957 | 2.6741 | 0.6576 | E9PQR7;Q9UK41;E9PM90;E9PLM9;E9PR04 | VPS28 |

| L100/ CTRL1 | L100/ CTRL2 | L100/ CTRL3 | Mean | SD | T-test p-value | Accession | Gene Symbol |
|---|---|---|---|---|---|---|---|
| 0.0571 | -1.9455 | *5.1502* | 1.0873 | 3.6583 | 0.6579 | H3BUX2;O43169;J3KNF8;D6RFH4 | CYB5B |
| 0.4894 | -0.6896 | *0.9526* | 0.2508 | 0.8467 | 0.6590 | P49821;G3V0I5;B4DE93;E9PMX3;E9PLC6;E9PQP1;E9PPR0;E9PPS5;H0YE81;H0YD04;E9PJL9;E9PPD6 | NDUFV1 |
| -0.3512 | -0.0169 | *0.1438* | -0.0748 | 0.2526 | 0.6591 | P22307;H0YD06;H0YF61;E9PLD1;H0YCB0 | SCP2 |
| -0.6864 | -0.1832 | *0.3909* | -0.1595 | 0.5390 | 0.6592 | Q96PU5;A0A1B0GVY1;K7ERN1 | NEDD4L |
| 0.4035 | -0.4293 | *0.4707* | 0.1483 | 0.5013 | 0.6594 | O75962;E7EPJ7;E7EWP2;F5H228 | TRIO |
| 0.0368 | -0.4202 | *0.1241* | -0.0864 | 0.2923 | 0.6596 | Q86SQ7;A0A0G2JQM2;A0A0G2JR50;A0A0C4DG71;A0A0G2JR20 | SDCCAG8 |
| 0.5188 | -0.5039 | *0.5054* | 0.1734 | 0.5866 | 0.6596 | O60716;C9JZR2;E9PRE2;H0YC95;E9PKY0;E9PKL1 | CTNND1 |
| -5.8529 | 4.4946 | *-3.4427* | -1.6003 | 5.4142 | 0.6596 | Q86UK0 | ABCA12 |
| 0.5406 | -3.2929 | *0.7386* | -0.6712 | 2.2726 | 0.6598 | Q15436;F5H365;G3V2R6;G3V1W4;G3V3G5;G3V4V1;G3V4Q2;G3V5X8;G3V5K1 | SEC23A |
| -0.2690 | -0.0800 | *0.9100* | 0.1870 | 0.6332 | 0.6598 | P31040;D6RFM5;A0A087X1I3;H0Y8X1 | SDHA |
| 0.0222 | -0.4735 | *0.1571* | -0.0980 | 0.3321 | 0.6599 | P46108;I3L297 | CRK |
| -0.1811 | -0.2573 | *1.1237* | 0.2284 | 0.7763 | 0.6609 | Q8WXI2;A0A2R8Y700;A0A2R8Y7A1;A0A2R8YED7;A0A2R8Y5S6;A0A2R8Y622;A0A2R8Y7K8;A0A2R8YGJ7;A0A2U3TZH5;A0A2R8YFM1;A0A2R8Y5R2;A0A2R8YDB6 | CNKSR2 |
| 0.5172 | 0.3314 | *-0.4141* | 0.1448 | 0.4929 | 0.6614 | Q9C0D5;A0A087WWL2;A0A087X1Z6;E9PH21 | TANC1 |
| -0.4217 | -0.4497 | *0.4316* | -0.1466 | 0.5009 | 0.6626 | P01111 | NRAS |
| 0.3411 | -0.6883 | *-0.1056* | -0.1509 | 0.5162 | 0.6628 | Q8N7X0;H0Y334;F8W7W4;H0Y333 | ADGB |
| -0.5287 | -1.8624 | *1.0936* | -0.4325 | 1.4804 | 0.6631 | Q6P9B6 | TLDC1 |
| 0.9191 | -0.4951 | *0.1946* | 0.2062 | 0.7072 | 0.6637 | O15397;F5GXT5;F5H009;F5H292 | IPO8 |
| 0.4337 | -0.4673 | *0.5084* | 0.1583 | 0.5430 | 0.6638 | Q9BXW7;A8MYZ9 | HDHD5 |

| L100/ CTRL1 | L100/ CTRL2 | L100/ CTRL3 | Mean | SD | T-test p-value | Accession | Gene Symbol |
|---|---|---|---|---|---|---|---|
| -0.6096 | -1.1882 | *0.8699* | -0.3093 | 1.0614 | 0.6639 | P46782;M0R0F0;M0R0R2;M0QZN2 | RPS5 |
| -0.2852 | -0.5035 | *0.3845* | -0.1347 | 0.4628 | 0.6641 | P38606;C9JVW8;C9JA17 | ATP6V1A |
| 0.1005 | -1.2490 | *0.3865* | -0.2540 | 0.8735 | 0.6645 | P20042;B5BU01 | EIF2S2 |
| -0.1767 | -0.8625 | *2.6715* | 0.5441 | 1.8740 | 0.6650 | Q99615 | DNAJC7 |
| 0.5630 | -0.2717 | *0.0731* | 0.1215 | 0.4194 | 0.6657 | C9JRZ6;Q9NX63;F8WAR4;A0A286YEX5 | CHCHD3 |
| -0.0649 | 1.2817 | *-3.2246* | -0.6693 | 2.3131 | 0.6660 | P62987;B4DV12;F5GXK7;F5GYU3;F5H265;F5H2Z3;F5H388;F5H6Q2;F5H747;J3QKN0;J3QS39;P0CG47;P0CG48;Q5PY61;Q96C32;M0R1V7;Q49A90;F5GZ39;J3QSA3;M0R2S1;M0R1M6 | UBA52 |
| 1.5937 | -1.9496 | *-1.2759* | -0.5440 | 1.8816 | 0.6662 | H0YC56 | MGAT4B |
| -0.6485 | -0.2945 | *2.3657* | 0.4742 | 1.6475 | 0.6675 | Q8NFZ4 | NLGN2 |
| 0.5888 | -0.2485 | *-1.0461* | -0.2353 | 0.8175 | 0.6676 | Q2M243;J3QKX2 | CCDC27 |
| 0.0170 | -0.4628 | *0.1637* | -0.0941 | 0.3277 | 0.6683 | Q8IWS0;A0A0D9SGE8;Q5JRC6 | PHF6 |
| -0.4046 | 0.4822 | *-0.5615* | -0.1613 | 0.5628 | 0.6688 | Q9UDY2;A0A1B0GTW1;A0A2R8YDH4 | TJP2 |
| 0.1552 | 0.3567 | *-1.2743* | -0.2541 | 0.8892 | 0.6696 | P00568;Q5T9B7;H0YID2 | AK1 |
| -0.6320 | -0.1003 | *0.3226* | -0.1366 | 0.4783 | 0.6699 | O00571;A0A0D9SF53;A0A0D9SFB3;A0A0D9SG12;A0A2R8Y4A4;A0A2R8YF78;A0A2R8YFS5;B5BTY4;A0A2R8Y5G6;A0A2R8Y645;A0A2R8Y7T2;A0A2R8YCW1;A0A2R8YDT5;A0A2R8YFR4;A0A2U3TZJ9;F6S8Q4;A0A2R8YDH3;O15523;A0A0J9YVQ7;A0A087WVZ1 | DDX3X |
| -0.2767 | -0.4513 | *1.8007* | 0.3576 | 1.2528 | 0.6700 | A0A087WTR6;A0A087WUA8;A0A087WZ37;A0A087WZN9;A0A087WZW3;A0A087X1T6;A0A087X250;A0A2R8Y6C0;A2A3D8;A2A3E3;A2A3E6;A2A3E7;A2A3E8;A9Z1W1;C9 | PCDH15 |

| L100/ CTRL1 | L100/ CTRL2 | L100/ CTRL3 | Mean | SD | T-test p-value | Accession | Gene Symbol |
|---|---|---|---|---|---|---|---|
| | | | | | | J4F3;E7EM53;E7EMG0;Q96QU1 | |
| -3.0127 | 0.9086 | *0.2986* | -0.6018 | 2.1100 | 0.6702 | Q9GZT4;V9GYE8;I3L4L3 | SRR |
| 1.8820 | 1.0319 | *-1.4397* | 0.4914 | 1.7255 | 0.6707 | P48637;A0A2R8Y5T7;A0A2R8Y430;A0A2R8Y7I7;A0A2R8Y6Q7 | GSS |
| 0.6449 | -1.0553 | *-0.3176* | -0.2426 | 0.8526 | 0.6709 | Q8IZY2 | ABCA7 |
| 0.6300 | -1.4227 | *-0.0953* | -0.2960 | 1.0410 | 0.6711 | A0A087WXS7;O43681;K7ERW9 | ASNA1 |
| -0.0758 | 1.4625 | *-3.5995* | -0.7376 | 2.5951 | 0.6712 | P0CJ78 | ZNF865 |
| -0.2519 | 0.9034 | *-1.8136* | -0.3874 | 1.3636 | 0.6714 | A0A0A0MSE1;Q15303 | ERBB4 |
| -0.0156 | 2.5507 | *-0.9807* | 0.5181 | 1.8252 | 0.6716 | C9IZE1;F8WAE5;Q9BY44;H7C5Q3 | EIF2A |
| 0.0227 | -0.3343 | *0.8080* | 0.1655 | 0.5844 | 0.6723 | P08758;D6RBL5;D6RBE9;E9PHT9;D6RCN3 | ANXA5 |
| 0.0334 | 0.9630 | *-0.4046* | 0.1972 | 0.6984 | 0.6731 | Q9P2I0 | CPSF2 |
| -1.1912 | 0.0319 | *0.4403* | -0.2397 | 0.8490 | 0.6732 | P30048 | PRDX3 |
| 0.4602 | -0.4798 | *-0.4265* | -0.1487 | 0.5280 | 0.6739 | Q15811;F8W7U0;A8CTZ0;C9JQZ7;C9J1A4;D6PAW0;C9JXS9;H7C429 | ITSN1 |
| 1.2991 | -0.5067 | *-2.3131* | -0.5069 | 1.8061 | 0.6749 | Q96MT7 | CFAP44 |
| 0.0348 | -0.2342 | *0.5213* | 0.1073 | 0.3830 | 0.6754 | P43304;F5GYK7;E7EM56;E9PDK4 | GPD2 |
| 0.6362 | 0.8670 | *-0.7623* | 0.2469 | 0.8816 | 0.6755 | Q9NR31;Q5SQT8;D6RAA2;D6RD69;D6RDB2;H0Y5E8;Q9Y6B6;X1WI22 | SAR1A |
| -1.1073 | 2.3689 | *-3.8935* | -0.8773 | 3.1375 | 0.6760 | Q2M2I8;E9PG46 | AAK1 |
| 0.5413 | 0.0305 | *-1.4265* | -0.2849 | 1.0211 | 0.6767 | P0C671 | C6orf222 |
| 1.2825 | 0.0008 | *-0.5111* | 0.2574 | 0.9239 | 0.6771 | Q6ZNG1 | ZNF600 |
| 0.8766 | -1.7840 | *-0.2099* | -0.3724 | 1.3377 | 0.6773 | Q9UBP0;A0A2R8YGN6;A0A2U3TZR0;A0A2R8Y4I8;A0A2R8YCL5;A0A2R8Y7W6;A0A2R8YFC9;A0A2R8YFW8 | SPAST |
| -0.6721 | -0.2000 | *0.4182* | -0.1513 | 0.5468 | 0.6791 | Q9UIA9;E7ESC6;H0YBE1 | XPO7 |
| 0.7469 | 0.4307 | *-0.5960* | 0.1939 | 0.7020 | 0.6796 | Q14687;H0Y613;H7C3B5 | GSE1 |
| 1.0088 | -0.4095 | *0.0047* | 0.2013 | 0.7293 | 0.6797 | Q9Y2H0;A0A0B4J2C2;F8WF49 | DLGAP4 |
| -0.3889 | 0.0650 | *0.0986* | -0.0751 | 0.2723 | 0.6799 | B3KUN1;P67775;E5RI56 | PPP2CA |
| -0.3889 | 0.0650 | *0.0986* | -0.0751 | 0.2723 | 0.6799 | P62714;E5RHC1;H0YC23;H0YBN9;E5RFI3;E5RHP4 | PPP2CB |

| L100/ CTRL1 | L100/ CTRL2 | L100/ CTRL3 | Mean | SD | T-test p-value | Accession | Gene Symbol |
|---|---|---|---|---|---|---|---|
| | | | | | | ;E5RJX4;E7ESG8 | |
| 1.8438 | 1.5009 | *-1.7218* | 0.5410 | 1.9671 | 0.6808 | E7ER45;O43451 | MGAM |
| 0.7396 | 2.6845 | *-1.6404* | 0.5946 | 2.1661 | 0.6813 | Q6ZU65 | UBN2 |
| -0.3163 | -0.2565 | *0.2954* | -0.0925 | 0.3372 | 0.6816 | P0DME0;A0A087X027 | SETSIP |
| -0.6773 | 0.5671 | *-0.4313* | -0.1805 | 0.6590 | 0.6820 | Q9UBT2;K7EPL2;U3KQ93;K7ES38;U3KQ55;K7ESK7 | UBA2 |
| 2.5530 | -0.2311 | *-0.8378* | 0.4947 | 1.8082 | 0.6823 | A0A0A0MRM9;Q14978 | NOLC1 |
| -1.2615 | 0.6450 | *1.9329* | 0.4388 | 1.6072 | 0.6829 | Q9P2D8 | UNC79 |
| 0.1441 | -0.9282 | *0.2507* | -0.1778 | 0.6520 | 0.6833 | F8VYC4;P0DJD1;A0A1W2PNU4;A0A286YES2;P0DJD0;A0A286YEQ5 | RGPD1 |
| -0.7395 | 2.4204 | *-4.5252* | -0.9481 | 3.4775 | 0.6833 | P13569;E7EPB6 | CFTR |
| -0.0551 | -0.3592 | *0.1895* | -0.0749 | 0.2749 | 0.6833 | Q99961;M0R0I3;M0QYE0 | SH3GL1 |
| 0.0116 | 0.5572 | *-1.3902* | -0.2738 | 1.0046 | 0.6833 | Q9HD45 | TM9SF3 |
| 2.2781 | 1.9418 | *-2.1888* | 0.6770 | 2.4876 | 0.6838 | P35908 | KRT2 |
| 0.1857 | -0.3560 | *0.5375* | 0.1224 | 0.4501 | 0.6839 | P10636;A0A0G2JPD5;A0A0G2JMX7;I3L170 | MAPT |
| 0.5072 | -0.4617 | *-0.5141* | -0.1562 | 0.5751 | 0.6844 | Q16799;Q2NKQ5;A8MT72 | RTN1 |
| -0.8411 | 0.7846 | *-0.6701* | -0.2422 | 0.8933 | 0.6849 | Q52M93;F2Z3L4;K7EL12;K7EPG8;Q8N7B4 | ZNF585B |
| -1.0150 | -1.4751 | *1.2888* | -0.4004 | 1.4809 | 0.6856 | Q9UQ16;E5RHK8;E5RIK2 | DNM3 |
| 0.5225 | 1.4717 | *-0.9897* | 0.3349 | 1.2414 | 0.6863 | Q96K76 | USP47 |
| -0.1517 | -0.5527 | *0.3420* | -0.1208 | 0.4482 | 0.6864 | A5A3E0 | POTEF |
| 0.6006 | -0.0860 | *-1.2848* | -0.2568 | 0.9542 | 0.6870 | J3QLH3;J3QQJ0;Q9UHR5;X6R3T8;F5H478;J3KRU5;J3KS14;J3KRR6 | SAP30BP |
| 0.2926 | 0.6010 | *-2.0744* | -0.3936 | 1.4638 | 0.6872 | P30084 | ECHS1 |
| -1.8338 | 2.5092 | *1.1125* | 0.5960 | 2.2171 | 0.6873 | G3V1V8;P10916 | MYL2 |
| 0.1921 | -0.1014 | *-0.2846* | -0.0646 | 0.2405 | 0.6874 | P62917;E9PKZ0;E9PKU4;E9PP36;G3V1A1 | RPL8 |
| -0.7400 | -1.8532 | *1.3071* | -0.4287 | 1.6030 | 0.6887 | P29373 | CRABP2 |
| 1.1702 | 0.7764 | *-4.4634* | -0.8389 | 3.1451 | 0.6895 | Q68DE3;C9JBW0 | USF3 |
| -0.1938 | -0.3238 | *1.1878* | 0.2234 | 0.8377 | 0.6896 | Q15738;C9JDR0 | NSDHL |
| -0.7696 | 0.2154 | *0.1201* | -0.1447 | 0.5433 | 0.6899 | P46781;A0A024R4M0;B5MCT8;C9JM19;A8MXK4;F2Z3C0 | RPS9 |
| 0.0407 | -0.3337 | *0.7191* | 0.1420 | 0.5337 | 0.6901 | P46013 | MKI67 |

| L100/ CTRL1 | L100/ CTRL2 | L100/ CTRL3 | Mean | SD | T-test p-value | Accession | Gene Symbol |
|---|---|---|---|---|---|---|---|
| -0.0323 | -0.1003 | *0.3063* | 0.0579 | 0.2178 | 0.6904 | Q5T3I3;Q5T3I4; Q8NCW5 | NAXE |
| 0.1154 | -0.1162 | *0.1050* | 0.0347 | 0.1308 | 0.6908 | Q12860;H0YIJ1 | CNTN1 |
| 0.0727 | 0.4130 | *-1.1304* | -0.2149 | 0.8109 | 0.6913 | Q9NPQ8;E9PI04 | RIC8A |
| -0.6181 | -0.3856 | *0.5242* | -0.1598 | 0.6037 | 0.6916 | Q15019;B5MCX3 ;C9J2Q4;C9J938; C9JB25;C9IY94; C9IZU3;H7C2Y0; H7C310;C9JQJ4; B5MD47;C9JZI2; A0A1B0GXJ2;C9 JFT1;F8WB65;C 9JSE7;C9JT15 | SEPT2 |
| -0.0124 | 0.8980 | *-0.3677* | 0.1726 | 0.6529 | 0.6919 | Q9H479 | FN3K |
| 0.1597 | 0.5019 | *-1.5153* | -0.2845 | 1.0795 | 0.6928 | Q8TD26;C9JFU2 ;H7C294 | CHD6 |
| -0.6645 | -0.8271 | *0.7885* | -0.2343 | 0.8895 | 0.6929 | Q92614;A0A0D9 SFK2 | MYO18A |
| -0.2931 | -0.0645 | *0.8235* | 0.1553 | 0.5898 | 0.6931 | A0A2U3TZV8;Q4 KWH8 | PLCH1 |
| -0.1371 | -1.9166 | *0.9254* | -0.3761 | 1.4360 | 0.6946 | Q6NUK1;J3KN42 | SLC25A24 |
| 1.1843 | 3.1848 | *-2.2249* | 0.7147 | 2.7352 | 0.6952 | Q7Z4W1;J3KS22 ;J3QS36;J3KRZ4 ;J3QL34;J3KSZ5; J3QS45 | DCXR |
| 0.0532 | -0.3898 | *0.8130* | 0.1588 | 0.6083 | 0.6954 | Q9Y618;C9J0Q5; C9JFD3;C9JE98; A0A384DVL6;C9 J7T7;C9JQE8;C9 J330 | NCOR2 |
| -0.1309 | -0.3699 | *0.2551* | -0.0819 | 0.3153 | 0.6968 | P12004 | PCNA |
| -0.5600 | 0.7189 | *0.3537* | 0.1709 | 0.6588 | 0.6972 | Q96KK5;P0C0S8 ;Q99878;A0A0U1 RR32;A0A0U1R RH7;P20671;Q93 077;Q9BTM1;P0 4908;Q7L7L0;H0 YFX9 | HIST1H2AH |
| 0.4196 | 0.5715 | *-0.5280* | 0.1544 | 0.5958 | 0.6975 | Q13148;A0A087 WV68;A0A087W XQ5;A0A087WY Y0;A0A087X260; G3V162;A0A087 WW61;A0A087W X29;A0A087WX6 7;K7EJM5;K7EN 94;A0A087WZC9 ;A0A087WZM1;A 0A1W2PNU8 | TARDBP |
| -0.4814 | -0.1643 | *1.4505* | 0.2682 | 1.0361 | 0.6977 | Q9ULI3 | HEG1 |
| 0.0954 | 0.8517 | *-0.4426* | 0.1682 | 0.6502 | 0.6980 | Q3KP31 | ZNF791 |
| -0.2927 | 0.4199 | *0.1517* | 0.0929 | 0.3599 | 0.6985 | A0A0G2JP87;B0 S7V6;Q8TD31;A 0A0G2JJ47;A2A BH1;A0A0G2JHN 4;A0A0G2JIL2;A | CCHCR1 |

| L100/ CTRL1 | L100/ CTRL2 | L100/ CTRL3 | Mean | SD | T-test p-value | Accession | Gene Symbol |
|---|---|---|---|---|---|---|---|
| | | | | | | 0A0G2JJZ1;A0A140T9J5;A2ABH3;A2ABH4;D6RAE7;E7EPK4;E7EQC5;E7EQE8;E9PGB6;E9PHV1 | |
| -0.3943 | -0.5179 | *2.0200* | 0.3692 | 1.4309 | 0.6987 | O75915;C9JQU6;F8WF33;F8WF90 | ARL6IP5 |
| 0.6754 | 0.8508 | *-3.3731* | -0.6156 | 2.3897 | 0.6991 | Q92599;A0A087X142;A6NFQ9;A6NMH6;F8W8I8;C9JV02 | SEPT8 |
| -0.4007 | 1.1715 | *-1.9937* | -0.4076 | 1.5826 | 0.6992 | A0A0A6YYH1;Q7Z6K5;H0YL03;H0YMP5 | C15orf38-AP3S2 |
| -0.1621 | -1.8623 | *0.9369* | -0.3625 | 1.4103 | 0.6997 | Q9UH03;A0A2R8Y4H2;B1AHR1;B1AHR2 | SEPT3 |
| -0.1431 | 0.1945 | *-0.2219* | -0.0569 | 0.2212 | 0.6997 | Q7Z3B4 | NUP54 |
| 1.6824 | -0.2203 | *-0.5390* | 0.3077 | 1.2012 | 0.7007 | O60299 | LZTS3 |
| -0.0130 | 0.6889 | *-0.2894* | 0.1289 | 0.5043 | 0.7013 | O60309 | LRRC37A3 |
| -1.4827 | -1.3135 | *1.5089* | -0.4291 | 1.6805 | 0.7015 | Q9Y2Q0;H0YAJ4 | ATP8A1 |
| 1.5484 | 0.3191 | *-4.1697* | -0.7674 | 3.0099 | 0.7019 | O14511 | NRG2 |
| -0.1630 | -1.0082 | *0.5684* | -0.2010 | 0.7890 | 0.7022 | Q13085;Q59FY4;A0A087X0W4;A0A0C4DGT1;A0A087WVR6;A0A087WWN5;A0A087WYK6;A0A087WYS8;A0A087X126;A0A087X2F8 | ACACA |
| -0.4962 | -0.1197 | *0.3085* | -0.1025 | 0.4026 | 0.7024 | O43314;A0A087WZV0;D6RBU4;H0Y9M0;A0A087WWN8;H0Y9S9 | PPIP5K2 |
| 0.3179 | -0.7209 | *1.0968* | 0.2313 | 0.9120 | 0.7034 | Q6UVM3;Q3SY61 | KCNT2 |
| 0.9362 | -1.1649 | *1.2185* | 0.3299 | 1.3023 | 0.7036 | P35555;F6U495;H0YN80 | FBN1 |
| -0.5251 | 0.1163 | *0.1257* | -0.0944 | 0.3730 | 0.7040 | Q75QN2;J3KNV5;H0YBS1;H0YC12 | INTS8 |
| -0.3455 | 0.3268 | *-0.2585* | -0.0924 | 0.3656 | 0.7043 | P22626;A0A087WUI2 | HNRNPA2B1 |
| 2.2030 | 1.3545 | *-1.9163* | 0.5471 | 2.1751 | 0.7056 | Q86UU1;A0A087WXY7 | PHLDB1 |
| 0.0333 | -0.9857 | *0.4090* | -0.1812 | 0.7217 | 0.7061 | P26641 | EEF1G |
| -0.2146 | -0.2277 | *0.9514* | 0.1697 | 0.6770 | 0.7065 | Q96AE4;E9PEB5;C9JSZ1;M0R3J3;M0R0C6;M0R251;M0R263 | FUBP1 |
| -0.5010 | 1.0731 | *0.0301* | 0.2007 | 0.8008 | 0.7065 | Q9H0E9;B5MCW3;H7C127;H7C128;H7C179 | BRD8 |

| L100/ CTRL1 | L100/ CTRL2 | L100/ CTRL3 | Mean | SD | T-test p-value | Accession | Gene Symbol |
|---|---|---|---|---|---|---|---|
| -0.0271 | -2.7195 | *1.2347* | -0.5040 | 2.0198 | 0.7077 | A0A1W2PP81;A0A1W2PPE2;A0A1W2PRV1 | hCG_1809904 |
| 0.0536 | 1.1382 | *-0.5522* | 0.2132 | 0.8564 | 0.7084 | Q86XP3;A0A0A0MSJ0;J3KRE3 | DDX42 |
| -1.7952 | -0.0872 | *0.8769* | -0.3352 | 1.3532 | 0.7097 | P07339;A0A1B0GV23;A0A1B0GVD5;A0A1B0GW44;A0A1B0GWE8;A0A1B0GU03;A0A1B0GU92;A0A1B0GVP3;H7C469;C9JH19;F8WD96;H7C1V0;F8W787 | CTSD |
| 0.1896 | -0.2072 | *0.1871* | 0.0565 | 0.2284 | 0.7100 | P05388;F8VWS0;F8VU65;F8VW21;G3V210;F8VPE8;F8VQY6;F8VRK7;F8VZS0;F8VS58;F8VWV4;F8W1K8;F8VYN4 | RPLP0 |
| -0.3947 | -0.2778 | *0.3679* | -0.1016 | 0.4107 | 0.7102 | M0QXL5;M0R0P1;M0R299;M0R2Q4;P22087;A6NHQ2 | FBL |
| 0.2470 | 0.7838 | *-2.2137* | -0.3943 | 1.5984 | 0.7108 | Q16775;H3BPK3;H3BPQ4;H3BQW8 | HAGH |
| -1.4263 | 1.7663 | *-1.7865* | -0.4822 | 1.9555 | 0.7109 | P62328 | TMSB4X |
| -0.0120 | 1.0906 | *-2.3963* | -0.4392 | 1.7823 | 0.7110 | P41223;C9JCD9;F8WCN7 | BUD31 |
| -0.0905 | -0.8378 | *0.4510* | -0.1591 | 0.6471 | 0.7116 | Q92878;A0A1W2PQ90;E7EN38;E7ESD9;H7C0V2;E9PM98 | RAD50 |
| -0.6659 | -2.2632 | *1.5264* | -0.4676 | 1.9026 | 0.7118 | Q8WXF1;X6RDA4 | PSPC1 |
| -1.2554 | 0.0161 | *0.5547* | -0.2282 | 0.9294 | 0.7120 | P63151;Q66LE6 | PPP2R2A |
| -0.4807 | 0.4637 | *0.4068* | 0.1299 | 0.5296 | 0.7122 | A0A1B0GUS7;B1AM27;F8W8M9;I6L9J0;A0A0U1RRB5;A0A0U1RRL2 | UNC13B |
| 0.6021 | 1.0640 | *-0.9075* | 0.2529 | 1.0311 | 0.7123 | Q9UBB4;A0A1W2PQD2;B1AHE4;B1AHE3 | ATXN10 |
| -0.9531 | 0.6117 | *-0.2337* | -0.1917 | 0.7832 | 0.7129 | O43809;H3BND3;H3BV41 | NUDT21 |
| -0.0845 | 0.4926 | *-0.9356* | -0.1758 | 0.7185 | 0.7129 | Q12840;J3KNA1 | KIF5A |
| -1.6341 | 0.7123 | *2.4077* | 0.4953 | 2.0296 | 0.7136 | Q9BS26 | ERP44 |
| 0.2769 | 1.1924 | *-3.1400* | -0.5569 | 2.2834 | 0.7138 | F5GZK2;Q96P44;H0YDH6 | COL21A1 |
| 0.5428 | -0.2513 | *-0.7776* | -0.1621 | 0.6647 | 0.7139 | P62191;Q53XL8 | PSMC1 |
| -0.4301 | -0.4811 | *0.5053* | -0.1353 | 0.5553 | 0.7140 | O95336;M0R261;M0R0U3;M0R1L2 | PGLS |

| L100/ CTRL1 | L100/ CTRL2 | L100/ CTRL3 | Mean | SD | T-test p-value | Accession | Gene Symbol |
|---|---|---|---|---|---|---|---|
| 0.2005 | -0.6992 | *1.1849* | 0.2287 | 0.9424 | 0.7151 | A0A0B4J1W0;F5GY88;O75448;J3KTL3 | MED24 |
| -0.1215 | -0.8889 | *2.1674* | 0.3857 | 1.5901 | 0.7152 | P62258;B4DJF2;I3L3T1;K7EIT4;K7EM20 | YWHAE |
| -0.3895 | 0.8542 | *-1.2240* | -0.2531 | 1.0458 | 0.7158 | Q9Y5H2 | PCDHGA11 |
| -0.8932 | 0.7057 | *-0.4069* | -0.1982 | 0.8197 | 0.7161 | A0A2R8Y793 | ACTB |
| 0.0787 | -0.3843 | *0.6992* | 0.1312 | 0.5436 | 0.7165 | Q96N67;A0A1B0GUE9;A0A1B0GVW2;A0A0C4DGY6;A0A0U1RR97 | DOCK7 |
| 0.4010 | -0.7546 | *0.9973* | 0.2146 | 0.8907 | 0.7170 | Q6P2D8 | XRRA1 |
| -0.1463 | -0.7571 | *0.4633* | -0.1467 | 0.6102 | 0.7175 | P15531 | NME1 |
| 0.2392 | 0.5309 | *-1.6038* | -0.2779 | 1.1575 | 0.7179 | Q7KZI7;E9PC69;E7ETY4;F5H4F6;H0YNV4 | MARK2 |
| 0.6746 | 1.4123 | *-4.3414* | -0.7515 | 3.1307 | 0.7180 | Q9NSC5;M0R2U7 | HOMER3 |
| -1.2078 | 0.4251 | *1.9015* | 0.3729 | 1.5553 | 0.7182 | Q96I99;E9PDQ8;H0Y852 | SUCLG2 |
| 0.0041 | -0.4714 | *0.2146* | -0.0842 | 0.3514 | 0.7184 | P34897;H0YIZ0;G3V5L0;G3V2Y4;G3V2W0;G3V4W5;G3V540;G3V241;G3V2E4;G3V3Y8;G3V4T0;G3V4X0;G3V2Y1;G3V3C6 | SHMT2 |
| -0.0886 | -0.7495 | *1.7729* | 0.3116 | 1.3079 | 0.7199 | P80404;H3BRN4;H3BNQ7;H3BMJ9;H3BPW8;H3BRJ1;H3BRT1 | ABAT |
| 0.3279 | 0.8407 | *-2.4183* | -0.4165 | 1.7524 | 0.7205 | J3KNC0;P52655 | GTF2A1 |
| 2.2457 | -0.1067 | *-4.6251* | -0.8287 | 3.4918 | 0.7209 | O94823;A0A2R8YDI5 | ATP10B |
| -0.6951 | 1.4012 | *0.0488* | 0.2516 | 1.0627 | 0.7215 | P62805;B2R4R0;B4E3H6;H0YBT8 | HIST1H4A |
| 4.8441 | 0.2092 | *-2.4457* | 0.8692 | 3.6895 | 0.7228 | Q14145 | KEAP1 |
| -0.3294 | 0.6718 | *-0.9067* | -0.1881 | 0.7987 | 0.7228 | Q14117 | DPYS |
| 0.8077 | 0.1517 | *-1.9936* | -0.3447 | 1.4652 | 0.7231 | Q68CZ2;E9PCX8;C9JHU5 | TNS3 |
| -1.8836 | 1.3014 | *2.0522* | 0.4900 | 2.0896 | 0.7240 | Q96FT9;G3V4X2 | IFT43 |
| -0.6742 | -1.0790 | *0.9843* | -0.2563 | 1.0933 | 0.7241 | Q9NPF5;Q5TG40;Q5TG39 | DMAP1 |
| -0.1513 | 0.3547 | *-0.0193* | 0.0613 | 0.2625 | 0.7248 | P07900 | HSP90AA1 |
| 0.4120 | 0.4629 | *-1.7649* | -0.2967 | 1.2718 | 0.7253 | Q96PE2 | ARHGEF17 |
| -0.3127 | 0.0623 | *0.0927* | -0.0526 | 0.2258 | 0.7258 | P06733;A0A2R8Y6G6;K7EM90;A0A2R8YEM5;A0A2R8Y879;K7ERS8;A0A2R8Y798;A0A2R8Y6I8;G5E9A7;Q9BZL6 | ENO1 |

| L100/ CTRL1 | L100/ CTRL2 | L100/ CTRL3 | Mean | SD | T-test p-value | Accession | Gene Symbol |
|---|---|---|---|---|---|---|---|
| 0.1612 | -2.9633 | *1.2815* | -0.5068 | 2.1998 | 0.7284 | B1AKL4;Q9NRA8 | EIF4ENIF1 |
| 2.1538 | 0.3782 | *-5.1586* | -0.8755 | 3.8140 | 0.7293 | Q5T7W0;B5MDS3 | ZNF618 |
| -0.6176 | -0.0499 | *1.3734* | 0.2353 | 1.0257 | 0.7295 | M0R2L6;A6NKH3;C9J4Z3;M0R0A1 | RPL37A |
| 0.1200 | -0.6057 | *0.1848* | -0.1003 | 0.4389 | 0.7305 | A0A2R8Y5Q8;Q15813;A0A2R8Y6Q1;A0A2R8Y7E7;A0A2R8YHI9;A0A2U3TZJ6;A0A2R8Y4V6;A0A2R8Y784;A0A2R8Y809;A0A2R8YER9;A0A2R8YFL4;A0A2R8Y4E7;A0A2R8Y4P2;A0A2R8Y5H6;A0A2R8Y6G3;A0A2R8Y6L2;A0A2R8Y787;A0A2R8Y7H8;A0A2R8Y897;A0A2R8YFM3 | TBCE |
| 1.0323 | 1.3219 | *-4.6649* | -0.7702 | 3.3760 | 0.7309 | Q8NBX0 | SCCPDH |
| 0.3847 | 0.4080 | *-1.5680* | -0.2584 | 1.1342 | 0.7312 | O95626 | ANP32D |
| -1.2828 | -0.8327 | *1.2086* | -0.3023 | 1.3277 | 0.7314 | Q7L576;A0A0G2JQT1;A0A0G2JR96;A0A087WU52;A0A087WWL1;A0A0G2JRV9;A0A0G2JRX2;H0YL50;A0A087WWY5;A0A0G2JRF5 | CYFIP1 |
| 0.8686 | 0.9025 | *-1.0199* | 0.2504 | 1.1002 | 0.7315 | Q9H4M9;A0A024R571;C9J2Z4;C9JC03 | EHD1 |
| -0.2619 | -0.3781 | *1.2616* | 0.2072 | 0.9150 | 0.7328 | P55884;C9JQN7;C9JZG1 | EIF3B |
| 0.0265 | -1.5044 | *0.7082* | -0.2566 | 1.1331 | 0.7328 | P15586;F6S8M0;H7C3P4;H0YFA9;F5H4C6 | GNS |
| 0.3914 | -0.2797 | *-0.4016* | -0.0966 | 0.4270 | 0.7329 | Q5T9S5;J3KP97;E9PFB9 | CCDC18 |
| 1.0071 | -0.7301 | *0.3167* | 0.1979 | 0.8747 | 0.7330 | P68363;C9JDS9 | TUBA1B |
| 2.6502 | 0.6198 | *-1.7693* | 0.5002 | 2.2122 | 0.7331 | A0A1B0GV93;Q53GQ0 | HSD17B12 |
| -0.4017 | 0.7483 | *-0.9275* | -0.1936 | 0.8571 | 0.7333 | P06730;D6RBW1 | EIF4E |
| 0.0074 | -0.0896 | *0.1715* | 0.0298 | 0.1320 | 0.7339 | P11940;E7EQV3;A0A087WTT1;E7ERJ7;H0YAR2;H0YBN4;H0YB86;E5RGH3;E5RJB9;H0YAP2;E5RH24;H0YAS6;H0YB75;E5RHG7;H0YAS7;E5RFD8;E5RGC4;H0YAW6;H0YC10 | PABPC1 |

| L100/ CTRL1 | L100/ CTRL2 | L100/ CTRL3 | Mean | SD | T-test p-value | Accession | Gene Symbol |
|---|---|---|---|---|---|---|---|
| 1.2355 | 0.8887 | *-1.2268* | 0.2991 | 1.3328 | 0.7350 | Q6PDA5 | WDR3 |
| -0.2329 | -0.1373 | *0.7256* | 0.1185 | 0.5280 | 0.7350 | Q13033;G3V340;G3V3G7;H0YJT2;H0YJZ4 | STRN3 |
| -0.3239 | 0.0210 | *0.6245* | 0.1072 | 0.4800 | 0.7362 | Q8IZ02;G3V115;H0YEZ4 | LRRC34 |
| 0.1894 | -1.6290 | *0.6363* | -0.2678 | 1.1999 | 0.7363 | Q2KHM9;F6SFD5;I3L341 | KIAA0753 |
| -0.0926 | 0.8391 | *-1.5537* | -0.2691 | 1.2061 | 0.7364 | Q9Y243;F8VS91 | AKT3 |
| -0.4068 | -0.5052 | *1.7733* | 0.2871 | 1.2880 | 0.7366 | A0A2R8YEI5;B3VCG5;P52757;B8ZZU1;C9JQ94;H0YG02 | CHN2 |
| 0.3193 | 0.3680 | *-0.4000* | 0.0957 | 0.4300 | 0.7369 | A0A2U3TZH1;Q9UQ90;A0A2R8Y3M4;A0A2R8Y4Y7;A0A2R8Y726;A0A2R8Y7B8;A0A2R8Y7E2;A0A2R8YDQ1;A0A2R8YEH4;A0A2R8YFW4;A0A2R8YGZ0;J3KRF6;A0A2R8Y6K2;H3BTR8;H3BTY6 | SPG7 |
| 0.9888 | -1.1277 | *0.9460* | 0.2690 | 1.2098 | 0.7372 | Q13561;F8VW18;H0YI98;F8VRV7;F8VX93;H0YHL1;F8W0U6 | DCTN2 |
| -0.2210 | -1.1610 | *0.7502* | -0.2106 | 0.9556 | 0.7394 | P03951;H0Y596 | F11 |
| -0.2857 | -0.5495 | *1.6163* | 0.2604 | 1.1817 | 0.7394 | Q8WUF5;K7EN03 | PPP1R13L |
| -0.4656 | -0.3679 | *1.6023* | 0.2563 | 1.1667 | 0.7402 | P10515;E9PEJ4;E9PKC7;H0YDD4 | DLAT |
| -0.0376 | 0.1969 | *-0.0647* | 0.0315 | 0.1438 | 0.7410 | Q9UHB7;C9JCE0 | AFF4 |
| -0.9583 | -0.6757 | *0.9555* | -0.2262 | 1.0330 | 0.7410 | Q15149;E9PMV1;E7ERX3;E7ESK0;Q9NV70;R4GMM7 | PLEC |
| -0.8582 | 0.0632 | *0.3743* | -0.1403 | 0.6409 | 0.7411 | P51665;H3BNT7;H3BTM8;H3BQV2 | PSMD7 |
| -0.2418 | -0.0026 | *0.1228* | -0.0405 | 0.1853 | 0.7411 | P09874;Q5VX84;Q5VX85 | PARP1 |
| 0.3265 | -0.4001 | *0.3542* | 0.0935 | 0.4277 | 0.7413 | Q5TCS8;J3KP89 | AK9 |
| 0.7141 | 0.5180 | *-2.3585* | -0.3755 | 1.7202 | 0.7417 | G3XAL9;P55011 | SLC12A2 |
| -0.3828 | 0.0382 | *0.7028* | 0.1194 | 0.5473 | 0.7419 | P42765;A0A0B4J2A4;K7EME0 | ACAA2 |
| 1.2942 | 1.3742 | *-5.0954* | -0.8090 | 3.7124 | 0.7421 | O14924 | RGS12 |
| 0.3174 | -0.0346 | *-0.1291* | 0.0513 | 0.2353 | 0.7422 | Q15084 | PDIA6 |
| -0.4651 | -0.0934 | *0.3068* | -0.0839 | 0.3861 | 0.7428 | O60264 | SMARCA5 |
| 0.6223 | -1.6026 | *0.2091* | -0.2571 | 1.1835 | 0.7429 | O00291;C9JMG5 | HIP1 |

| L100/ CTRL1 | L100/ CTRL2 | L100/ CTRL3 | Mean | SD | T-test p-value | Accession | Gene Symbol |
|---|---|---|---|---|---|---|---|
| -0.1973 | -0.6029 | *1.5384* | 0.2461 | 1.1375 | 0.7439 | P49411;H3BNU3;F8VZ96 | TUFM |
| 0.3725 | -0.2917 | *0.1378* | 0.0729 | 0.3369 | 0.7439 | Q9UL36;J9JID5 | ZNF236 |
| 0.0282 | -1.1854 | *0.5731* | -0.1947 | 0.9002 | 0.7439 | Q8IYY4;C9JRW2 | DZIP1L |
| 0.5680 | 2.1313 | *-5.2074* | -0.8360 | 3.8656 | 0.7440 | O95819;A0A0D9SEY1;E7EN19;E7ENQ1;G3XAA2;G5E948;H7C360;A0A0D9SG62;E7EX83;C9J840;H7C0P6;C9J338;E7ETN6 | MAP4K4 |
| 0.6026 | 0.3898 | *-0.5837* | 0.1362 | 0.6325 | 0.7450 | O75140;A0A2R8Y5K9;A0A2R8Y6H3;A0A2R8Y721;A0A2R8Y7U0;H0Y770;A0A2R8Y5E9;A0A2R8Y5P2;A0A2R8Y5T1;A0A2R8Y6H8;A0A2R8Y6V4;A0A2R8Y7U6;A0A2R8Y842;A0A2R8YEW8;A0A2R8YF50;A0A2R8YF97;A0A2R8YFS1 | DEPDC5 |
| 0.2455 | -2.3522 | *0.9820* | -0.3749 | 1.7515 | 0.7465 | Q9BXP5;H7C3A1;H7C1K0;A0A0A0MSP6 | SRRT |
| 1.5457 | 2.4714 | *-2.3681* | 0.5497 | 2.5689 | 0.7465 | Q9Y2X7;J3QRU8;A0A0C4DGN6;J3QLH1;K7EN79;J3QL89 | GIT1 |
| -0.6383 | 0.0981 | *1.1000* | 0.1866 | 0.8725 | 0.7467 | Q58FF6 | HSP90AB4P |
| -0.9744 | 0.5983 | *-0.1277* | -0.1679 | 0.7871 | 0.7472 | P30049 | ATP5F1D |
| 2.1594 | 1.0529 | *-6.0663* | -0.9514 | 4.4641 | 0.7475 | Q9Y2I8;A0A087WTQ2;E7EQ49;C9JGR9 | WDR37 |
| -0.1511 | 0.1030 | *0.1519* | 0.0346 | 0.1626 | 0.7479 | Q92932;A0A0J9YWV3 | PTPRN2 |
| -0.2644 | 1.3358 | *-2.2015* | -0.3767 | 1.7713 | 0.7480 | Q86W50 | METTL16 |
| 0.6838 | 0.4162 | *-0.6501* | 0.1500 | 0.7056 | 0.7481 | K7ELQ6 | MISP3 |
| 2.0852 | -4.3834 | *5.4909* | 1.0642 | 5.0157 | 0.7485 | Q7Z553;G3V5U0 | MDGA2 |
| 0.1009 | -0.0097 | *-0.1817* | -0.0302 | 0.1424 | 0.7489 | P13667;C9JAB0;C9JMN9;C9K0C4 | PDIA4 |
| -0.2424 | 0.2332 | *-0.1508* | -0.0533 | 0.2524 | 0.7496 | A0A2R8YEQ9;P51787 | KCNQ1 |
| 0.0482 | -0.3188 | *0.1216* | -0.0497 | 0.2360 | 0.7503 | Q3L8U1;H3BTW3 | CHD9 |
| -0.6685 | 0.4902 | *-0.1880* | -0.1221 | 0.5822 | 0.7512 | P31689 | DNAJA1 |
| 1.5199 | -4.0987 | *0.6747* | -0.6347 | 3.0295 | 0.7515 | P22735;H0YN27;H0YKI6;H0YLJ6;H0YMQ8;H0YNM4 | TGM1 |
| -0.6194 | -1.3713 | *1.1717* | -0.2730 | 1.3064 | 0.7520 | P11388 | TOP2A |

| L100/ CTRL1 | L100/ CTRL2 | L100/ CTRL3 | Mean | SD | T-test p-value | Accession | Gene Symbol |
|---|---|---|---|---|---|---|---|
| 3.0706 | -0.3468 | *-1.2873* | 0.4788 | 2.2933 | 0.7522 | Q9ULL4 | PLXNB3 |
| 2.4502 | -0.9137 | *-3.3612* | -0.6082 | 2.9177 | 0.7526 | Q9NSD4;A6NFS0 | ZNF275 |
| -0.6300 | -1.3785 | *3.7271* | 0.5729 | 2.7572 | 0.7534 | Q9H0B3;A0A087WXN0;V9GY12 | IQCN |
| -1.8303 | 1.6751 | *1.3648* | 0.4032 | 1.9405 | 0.7534 | Q9ULD4;E9PSF3;F6XDC4;H0Y7B4;A8WI61;A8WI62 | BRPF3 |
| -5.8307 | 3.1185 | *6.7375* | 1.3417 | 6.4697 | 0.7538 | Q8IVL0;A0A2R8YFX5 | NAV3 |
| -0.1601 | -0.4863 | *1.2031* | 0.1856 | 0.8962 | 0.7542 | P62333;A0A087X2I1;H0YJC0;H0YJS8;H0YJX2 | PSMC6 |
| -0.4701 | -0.5896 | *0.6395* | -0.1401 | 0.6777 | 0.7546 | Q9Y224;G3V4C6;G3V4E7;H0YJB9 | RTRAF |
| -0.4898 | 0.3464 | *-0.1158* | -0.0864 | 0.4189 | 0.7551 | P27635;X1WI28;A0A087WV22;F8W7C6;A6QRI9;B8A6G2;Q96L21;H7C123;H7C2C5;H7C2U2 | RPL10 |
| -5.5139 | -1.9359 | *4.3602* | -1.0299 | 4.9990 | 0.7553 | Q8WYQ5 | DGCR8 |
| 0.1572 | 0.2462 | *-0.7385* | -0.1117 | 0.5446 | 0.7564 | A0A0G2JQF4;A0A0G2JQF5;I3L233;A0A0G2JRW7;I3L243;A0A0G2JNB1;A0A0G2JNT7;Q7Z3B3;A0A0J9YVS1;A0A1W2PRB5;A0A1W2PPV8 | KANSL1 |
| -0.3629 | -0.4602 | *1.5038* | 0.2269 | 1.1069 | 0.7565 | A8TX70;E9PAL5 | COL6A5 |
| 1.1212 | -1.2031 | *-0.6640* | -0.2486 | 1.2165 | 0.7572 | Q5JXI8;Q13642;Q5JXH7;Q5JXH8;Q5JXI3;A0A0D9SFZ9;A0A0D9SFB0;A0A0D9SFI6;Q5JXH9;A0A0D9SGB2;Q5JXI0;A0A0D9SG53;A0A0D9SGD1;A0A0D9SEY7;A0A0D9SGC5 | FHL1 |
| -0.4081 | 0.0783 | *0.1448* | -0.0616 | 0.3019 | 0.7574 | P42695;G3V1A9;E9PKK4;E9PQA3;E9PLE0 | NCAPD3 |
| 0.6379 | -0.3328 | *-0.0038* | 0.1004 | 0.4937 | 0.7582 | P21281;H0YC04;E5RGH6;H0YC45 | ATP6V1B2 |
| 0.8416 | 0.5281 | *-2.4861* | -0.3721 | 1.8374 | 0.7593 | Q8IYU2;H0YC48 | HACE1 |
| -0.1918 | -0.2855 | *0.8649* | 0.1292 | 0.6389 | 0.7596 | P20645;F5GX30;H0YF90;H0YGT2 | M6PR |
| -0.8232 | 1.0201 | *0.3702* | 0.1890 | 0.9349 | 0.7597 | I1E4Y6;Q6Y7W6;E7ESB6;C9JHW1;C9JW88 | GIGYF2 |

| L100/ CTRL1 | L100/ CTRL2 | L100/ CTRL3 | Mean | SD | T-test p-value | Accession | Gene Symbol |
|---|---|---|---|---|---|---|---|
| -0.0942 | 0.3837 | *-0.1179* | 0.0572 | 0.2830 | 0.7598 | Q9UQB3;E7EPC8;B4DRK2;E9PHB5;D6R9A8;D6RC65;D6RBA8;D6RHE9;D6RF55 | CTNND2 |
| -0.4082 | 0.4515 | *-0.3297* | -0.0954 | 0.4753 | 0.7612 | A0A0C4DGK3;G3V4T3;A0A087X139;A0A1W2PRD7;G3V5N1 | SYNE2 |
| 0.5673 | -0.4190 | *0.1497* | 0.0994 | 0.4951 | 0.7613 | C9IZY1;C9J5P1;C9JA90;C9JNE2;Q9Y530 | OARD1 |
| -0.2515 | -2.5521 | *1.5654* | -0.4128 | 2.0635 | 0.7620 | Q8IWK6;D6RDX4 | ADGRA3 |
| -2.5763 | 1.6124 | *-0.2945* | -0.4194 | 2.0971 | 0.7621 | P54707 | ATP12A |
| 1.2667 | -0.8586 | *0.2293* | 0.2125 | 1.0628 | 0.7622 | Q9BY76 | ANGPTL4 |
| 0.2409 | 2.5522 | *-5.1651* | -0.7907 | 3.9607 | 0.7625 | A0A2R8YF49;A0A2R8YFR7;Q12774;A0A2R8Y614 | ARHGEF5 |
| -2.4294 | 1.4580 | *-0.1960* | -0.3891 | 1.9509 | 0.7627 | Q8N7M2 | ZNF283 |
| -0.0515 | 0.4711 | *-0.8026* | -0.1276 | 0.6402 | 0.7628 | Q9ULE0 | WWC3 |
| -0.1888 | -0.4271 | *0.3712* | -0.0816 | 0.4098 | 0.7631 | Q8N4C8 | MINK1 |
| 0.2071 | -0.2353 | *0.1755* | 0.0491 | 0.2468 | 0.7633 | Q13363;D6RAX2;H0Y8W7;E7EPF8;E7ESU7;E9PGB1;H0Y9M9;H0Y8U5;E7EUB3;A0A087WYL1 | CTBP1 |
| -0.2752 | 0.8783 | *-1.2324* | -0.2098 | 1.0569 | 0.7638 | Q8NCM8;H0YEX1 | DYNC2H1 |
| -0.4598 | 1.3568 | *-0.3002* | 0.1989 | 1.0059 | 0.7646 | P39656;A0A0C4DGS1;U3KQ84 | DDOST |
| 1.3205 | 0.8701 | *-3.9006* | -0.5700 | 2.8932 | 0.7654 | Q02809 | PLOD1 |
| 0.9670 | 1.0753 | *-1.2650* | 0.2591 | 1.3210 | 0.7664 | Q9Y653;H3BRH0;A0A1B0GX62;H3BM73;H3BMF8;H3BMY9;H3BNH4;H3BP67;H3BQ46;H3BQZ1;H3BRA1;H3BRI2;H3BRI7;H3BRZ4;H3BS94;H3BSB8;H3BSJ6;H3BSN7;H3BT88;H3BTH7;H3BUH2;H3BUI6;H3BUU6;H3BV52;H3BV72;H3BVA0;H3BVD3;H3BNN3;H3BPA6;H3BQJ9;H3BRB4;H3BSP5;H3BSR1;H3BTD2;H3BTK9;H3BVE9 | ADGRG1 |
| 0.5252 | -0.2745 | *-0.0111* | 0.0799 | 0.4075 | 0.7666 | P23497 | SP100 |
| 0.2780 | 2.8829 | *-5.7688* | -0.8693 | 4.4385 | 0.7668 | P57740;H0YG15 | NUP107 |

| L100/ CTRL1 | L100/ CTRL2 | L100/ CTRL3 | Mean | SD | T-test p-value | Accession | Gene Symbol |
|---|---|---|---|---|---|---|---|
| 0.4923 | 0.7041 | *-0.7429* | 0.1512 | 0.7815 | 0.7695 | Q9UIF8;C9JCA6; F6VJC3 | BAZ2B |
| -0.6529 | -0.3038 | *0.5862* | -0.1235 | 0.6389 | 0.7696 | O00425;F8WD15 | IGF2BP3 |
| -0.4354 | -0.0963 | *0.3143* | -0.0725 | 0.3754 | 0.7699 | P12236 | SLC25A6 |
| -4.9048 | -0.1319 | *2.7918* | -0.7483 | 3.8852 | 0.7704 | P50479;C9J542 | PDLIM4 |
| 0.7638 | 0.3307 | *-0.6700* | 0.1415 | 0.7354 | 0.7706 | Q9HCE3;K7EIU3 ;K7EQN0;K7ES4 7 | ZNF532 |
| 0.6454 | 0.8095 | *-2.5460* | -0.3637 | 1.8917 | 0.7708 | O94888;C9JAT7; F8WB69 | UBXN7 |
| 0.6322 | -0.4605 | *-0.5512* | -0.1265 | 0.6586 | 0.7710 | Q7L2R6;D6RF03 | ZNF765 |
| -0.4899 | -0.2707 | *0.4708* | -0.0966 | 0.5035 | 0.7713 | Q9NRN7;E9PLW 6 | AASDHPPT |
| -0.1150 | -0.1246 | *0.4185* | 0.0596 | 0.3108 | 0.7713 | Q9HAV7 | GRPEL1 |
| -1.3717 | 1.9569 | *-1.7568* | -0.3906 | 2.0420 | 0.7719 | P11488 | GNAT1 |
| 0.6743 | -0.3189 | *-0.7823* | -0.1423 | 0.7442 | 0.7720 | P15170;H3BR35; H3BSV8 | GSPT1 |
| 1.9761 | 0.0934 | *-1.1647* | 0.3016 | 1.5807 | 0.7724 | Q9BYV8 | CEP41 |
| 1.0303 | -0.1960 | *-0.3959* | 0.1461 | 0.7722 | 0.7742 | P16989 | YBX3 |
| 0.2905 | -0.1893 | *0.0350* | 0.0454 | 0.2400 | 0.7743 | P52907 | CAPZA1 |
| -0.1567 | -1.3738 | *0.8901* | -0.2135 | 1.1330 | 0.7751 | Q5W5X9 | TTC23 |
| 0.5069 | 1.2446 | *-1.0817* | 0.2233 | 1.1888 | 0.7758 | A6NK75;Q9P255 | ZNF98 |
| 0.5850 | 1.7308 | *-4.0319* | -0.5720 | 3.0506 | 0.7762 | Q9UHG3 | PCYOX1 |
| 0.6886 | -0.8230 | *-0.2959* | -0.1434 | 0.7673 | 0.7768 | P16152;A8MTM1 ;E9PQ63 | CBR1 |
| -1.2760 | 3.3904 | *-0.6910* | 0.4745 | 2.5422 | 0.7772 | Q14694;H3BNS8 ;H3BQC6;H3BQP 1 | USP10 |
| 2.5298 | 0.4020 | *-5.1435* | -0.7372 | 3.9615 | 0.7778 | O00472;D6RC27 | ELL2 |
| 0.0158 | -0.0351 | *0.0409* | 0.0072 | 0.0387 | 0.7783 | Q6ZVF9 | GPRIN3 |
| 0.0851 | -0.0924 | *0.0607* | 0.0178 | 0.0962 | 0.7791 | Q58FG0 | HSP90AA5P |
| 0.3117 | -0.0206 | *-0.5230* | -0.0773 | 0.4203 | 0.7802 | Q99715;D6RGG3 ;A0A087X0A8;H0 Y5N9;H0Y991 | COL12A1 |
| -0.4102 | 3.7377 | *-6.0312* | -0.9012 | 4.9029 | 0.7804 | Q8N8L2 | ZNF491 |
| 0.1875 | -0.8672 | *0.3209* | -0.1196 | 0.6509 | 0.7804 | Q96ER9;C9JSW 8 | CCDC51 |
| -0.1751 | -0.2417 | *0.7088* | 0.0973 | 0.5306 | 0.7808 | O75369;E7EN95; A0A0A0MT44;A0 A0C4DGA1 | FLNB |
| -0.2728 | -0.1052 | *0.6464* | 0.0894 | 0.4896 | 0.7816 | P36551 | CPOX |
| 1.4493 | -0.0605 | *-2.4717* | -0.3610 | 1.9777 | 0.7818 | Q7L1W4;E9PL08 ;E9PMF9;E9PJ89 ;E9PJS7;Q5VWA 0 | LRRC8D |
| -0.0369 | -0.1606 | *0.3402* | 0.0476 | 0.2608 | 0.7821 | A0A2U3TZI4;Q5 VVM6 | CCDC30 |
| -1.2214 | -0.2875 | *0.9230* | -0.1953 | 1.0752 | 0.7829 | O95477 | ABCA1 |
| -0.4702 | 0.3825 | *0.3509* | 0.0878 | 0.4834 | 0.7830 | Q14195;H0YBT4; D6RF19;H0YB87 ;Q8IYB8 | DPYSL3 |
| -0.2445 | 1.0095 | *-0.3532* | 0.1373 | 0.7573 | 0.7833 | P49588;H3BPK7 | AARS |

| L100/ CTRL1 | L100/ CTRL2 | L100/ CTRL3 | Mean | SD | T-test p-value | Accession | Gene Symbol |
|---|---|---|---|---|---|---|---|
| -0.1588 | -0.2988 | *0.2906* | -0.0557 | 0.3079 | 0.7838 | P62873;B3KVK2; F6UT28;F6X3N5; B1AKQ8 | GNB1 |
| 0.1910 | -1.2315 | *1.8869* | 0.2821 | 1.5612 | 0.7839 | Q92542;H0Y3Z4; H0Y6T7;Q5T205; Q5T209;Q5T211 | NCSTN |
| 0.1083 | -0.3440 | *0.4519* | 0.0721 | 0.3991 | 0.7841 | Q9NVA2;D6RER 5;D6RGI3;D6RD U5;H0Y961;D6R 9Y6;D6RDP1;H0 Y9G8 | SEPT11 |
| -1.0352 | 1.9017 | *-1.9557* | -0.3631 | 2.0146 | 0.7845 | Q5MIZ7 | PPP4R3B |
| 0.5205 | -1.1671 | *0.1672* | -0.1598 | 0.8900 | 0.7852 | Q8IUD2;X6RLX0; X6RM00;A0A0U1 RQM4;K7EPP6;A 0A0U1RQN0;K7 EPD6 | ERC1 |
| -2.4705 | 1.3574 | *0.0655* | -0.3492 | 1.9474 | 0.7855 | Q08378;A0A087 WV43 | GOLGA3 |
| -0.0980 | 0.7445 | *-0.3402* | 0.1021 | 0.5694 | 0.7855 | Q66K14 | TBC1D9B |
| -0.2377 | -0.4217 | *1.1077* | 0.1494 | 0.8349 | 0.7859 | C9JPM4;P18085; C9JAK5;F8WDB 3;C9J6P1 | ARF4 |
| -0.5631 | 2.0154 | *-2.7264* | -0.4247 | 2.3739 | 0.7860 | P48739;A0A0A0 MSW4;B3KYB6; B3KYB7 | PITPNB |
| 0.8856 | -0.3762 | *-1.0322* | -0.1743 | 0.9748 | 0.7861 | B2RPK0 | HMGB1P1 |
| 2.1646 | -0.1237 | *-1.1387* | 0.3008 | 1.6921 | 0.7873 | P13929;E5RGZ4; K7EKN2;K7EPM 1;E5RG95;E5RI0 9 | ENO3 |
| -0.1648 | 0.7177 | *-1.0114* | -0.1528 | 0.8646 | 0.7884 | Q9BVG8;F5H3M 2;B7Z896 | KIFC3 |
| 0.4910 | 1.2419 | *-2.9033* | -0.3902 | 2.2086 | 0.7885 | Q92608;E7ERW7 | DOCK2 |
| -0.2830 | 1.1126 | *-1.5287* | -0.2330 | 1.3213 | 0.7889 | P47985;P0C7P4 | UQCRFS1 |
| 0.5820 | -0.1250 | *-0.8303* | -0.1244 | 0.7061 | 0.7890 | Q8TB36 | GDAP1 |
| 0.4601 | 1.2226 | *-2.8171* | -0.3781 | 2.1463 | 0.7891 | O15056;H7BY56 | SYNJ2 |
| -0.2579 | -0.5873 | *0.5392* | -0.1020 | 0.5792 | 0.7892 | P23634;H7BY13; H7BZS8;H0YDG 5 | ATP2B4 |
| -0.1718 | -0.4830 | *0.4143* | -0.0802 | 0.4556 | 0.7893 | Q9BYX7 | POTEKP |
| 0.3922 | 0.9316 | *-0.8439* | 0.1600 | 0.9102 | 0.7896 | Q8NAP3;D6RBC 4;A0A1B0GV48 | ZBTB38 |
| -0.0623 | 0.4245 | *-0.6440* | -0.0939 | 0.5350 | 0.7898 | P35269;M0R0R9; M0QXD6;M0R0Z 3 | GTF2F1 |
| -0.2541 | -0.3387 | *0.9807* | 0.1293 | 0.7385 | 0.7903 | E9PJK2;O75530 | EED |
| 0.2904 | -2.1520 | *3.2886* | 0.4757 | 2.7251 | 0.7909 | Q6IN85;G3V5Z3; H0YIY8;M0R103 | PPP4R3A |
| -0.6164 | -0.2871 | *0.5803* | -0.1077 | 0.6182 | 0.7913 | Q9NQW1 | SEC31B |
| -0.4501 | -0.0987 | *0.9201* | 0.1238 | 0.7117 | 0.7917 | O43602;A8K340; A0A1B0GWD1;H 3BLV5;A0A1B0G UE1;E7EU50 | DCX |

| L100/ CTRL1 | L100/ CTRL2 | L100/ CTRL3 | Mean | SD | T-test p-value | Accession | Gene Symbol |
|---|---|---|---|---|---|---|---|
| 0.6010 | -2.3609 | *0.8326* | -0.3091 | 1.7807 | 0.7920 | Q9UBW7 | ZMYM2 |
| 1.0527 | 0.0607 | *-0.6646* | 0.1496 | 0.8621 | 0.7922 | Q15717 | ELAVL1 |
| 0.0821 | 1.4396 | *-0.9104* | 0.2038 | 1.1797 | 0.7930 | P31751;M0R0P9;C9JHS6;J3QLS6;A0A0A0MRF1;A0A1B0GXA2;A8MX96;C9JIJ1;E7EVP8;J3KRI8;M0QZK3;C9J258;C9JC83;C9JIF6;J3KSY8;M0R275;J3QKW1 | AKT2 |
| -0.0971 | 1.4474 | *-0.7630* | 0.1958 | 1.1339 | 0.7931 | Q9Y657 | SPIN1 |
| -0.4218 | 0.9466 | *-0.1498* | 0.1250 | 0.7244 | 0.7932 | O15347;E7EQU1;E7ES08;E9PES6 | HMGB3 |
| -0.3741 | -0.1564 | *0.3409* | -0.0632 | 0.3665 | 0.7934 | Q9HB07;F8VQQ3;F8VR84;H3BPH3 | C12orf10 |
| -0.1997 | 1.8973 | *-0.9370* | 0.2535 | 1.4705 | 0.7934 | P31323 | PRKAR2B |
| -0.3739 | -1.6470 | *1.2691* | -0.2506 | 1.4619 | 0.7945 | Q8N568;A0A0U1RR70;G5E9L9 | DCLK2 |
| 2.0184 | -0.7391 | *-0.4950* | 0.2614 | 1.5264 | 0.7947 | Q17RW3;Q8N2I2 | ZNF619 |
| 0.2384 | 0.0003 | *-0.4058* | -0.0557 | 0.3257 | 0.7950 | P13674 | P4HA1 |
| -0.7171 | 1.6833 | *-0.3077* | 0.2195 | 1.2841 | 0.7951 | Q2M389;A0A087X256 | WASHC4 |
| -0.4001 | -0.5419 | *1.5355* | 0.1978 | 1.1606 | 0.7956 | P26640;A0A024RCN6;A0A1U9X9A3;A0A1U9X9C4;A0A1U9X9C7;A0A1U9X9C8;A0A140T936;A0A140T954;H0Y426;A0A0G2JJT9;A2ABF4 | VARS |
| -0.5395 | -0.5153 | *0.6950* | -0.1199 | 0.7059 | 0.7962 | Q9UKG1 | APPL1 |
| 3.2650 | 3.0128 | *-4.1359* | 0.7140 | 4.2020 | 0.7963 | Q8IV35;H7C4Y4 | WDR49 |
| 0.9122 | -1.4111 | *1.2328* | 0.2446 | 1.4428 | 0.7967 | F5GWH7 | MDM2 |
| -0.7754 | 0.3562 | *0.8407* | 0.1405 | 0.8294 | 0.7968 | Q07343;E9PMG3 | PDE4B |
| 2.5173 | 1.6477 | *-2.7369* | 0.4760 | 2.8163 | 0.7973 | P05556 | ITGB1 |
| -0.2396 | -0.5932 | *0.5399* | -0.0976 | 0.5797 | 0.7980 | P57737;A0A0A6YYL4;I3L167;I3L1G9;I3L1M1;I3L2Y6;I3L359;I3L426;I3NI06 | CORO7 |
| 0.3635 | 2.4448 | *-4.6352* | -0.6090 | 3.6388 | 0.7992 | P07099 | EPHX1 |
| -0.1057 | -0.2778 | *0.6237* | 0.0801 | 0.4786 | 0.7992 | P61106;X6RFL8 | RAB14 |
| -0.0489 | -0.6204 | *1.1097* | 0.1468 | 0.8815 | 0.8001 | Q9H040;L8E708 | SPRTN |
| -0.9203 | -0.0119 | *1.5556* | 0.2078 | 1.2525 | 0.8009 | A0A087X0T8;A0A087X1W8;A0A0A0MTJ8;Q9BY67;X5DQS5 | CADM1 |

| L100/ CTRL1 | L100/ CTRL2 | L100/ CTRL3 | Mean | SD | T-test p-value | Accession | Gene Symbol |
|---|---|---|---|---|---|---|---|
| 0.0060 | 0.0470 | *-0.0871* | -0.0114 | 0.0687 | 0.8012 | Q12791;A0A1W2PP94;A0A1W2PPX7;A0A1W2PQU4;A0A1W2PR62;A0A1W2PRN5;A0A1W2PRV4;B7ZMF5;H0Y5Z6;Q5SVJ8;Q5SVJ9;A0A0A0MRR0;A0A1W2PNH9;A0A1W2PNQ3;A0A1W2PNW6;A0A1W2PNY9;A0A1W2PP06;A0A1W2PPY5;A0A1W2PPZ1;A0A1W2PQ61;A0A1W2PQ93;A0A1W2PQA0;A0A1W2PQK5;A0A1W2PR56;A0A1W2PRB0;A0A1W2PRG5;A0A1W2PSD3;Q5SVJ7;D5MRH1;A0A1W2PQR1;A0A087WZL8;A0A1W2PPT7;A0A1W2PQZ8;S4R453;A0A0A0MSE6;A0A1W2PNG1;A0A1W2PNY7;A0A1W2PP26;A0A1W2PPH9;A0A1W2PPQ3;A0A1W2PPS2;A0A1W2PQ39;A0A1W2PQJ9;A0A1W2PRE5;A0A1W2PRX6;A0A1W2PS97;H0Y379;H0Y382;Q5SVK0;Q5SVK5 | KCNMA1 |
| -0.5214 | -0.2062 | *0.4752* | -0.0841 | 0.5094 | 0.8017 | P36542 | ATP5F1C |
| -0.3586 | 0.5959 | *0.0013* | 0.0795 | 0.4821 | 0.8020 | Q14966 | ZNF638 |
| -2.1203 | 1.1426 | *0.1527* | -0.2750 | 1.6730 | 0.8026 | P15313;C9JL73;C9JNS9;C9JZ02 | ATP6V1B1 |
| -0.1397 | 1.8225 | *-0.9751* | 0.2359 | 1.4361 | 0.8028 | Q8NE09;A0A087WV61;G3V112;H0YBV2 | RGS22 |
| -0.3131 | 0.1010 | *0.3847* | 0.0576 | 0.3509 | 0.8031 | Q13459;M0R0P8;M0R300 | MYO9B |
| 0.5946 | 1.6698 | *-1.4778* | 0.2622 | 1.5999 | 0.8032 | Q9NSV4 | DIAPH3 |
| 0.3775 | 0.8261 | *-0.7931* | 0.1368 | 0.8360 | 0.8034 | P38398;Q5YLB2;A0A0U1RRA9;E7EWN5;E9PH68;Q3B891;E7EUM2;E7EQW4;B7ZA85;E7ENB7;H0Y8 | BRCA1 |

| L100/ CTRL1 | L100/ CTRL2 | L100/ CTRL3 | Mean | SD | T-test p-value | Accession | Gene Symbol |
|---|---|---|---|---|---|---|---|
| | | | | | | B8;A0A2R8Y7V5; H0Y881;K7EPC7 ;A0A2R8Y587;Q9 6S94 | |
| 1.9818 | -0.0790 | *-3.1775* | -0.4249 | 2.5970 | 0.8035 | Q92766 | RREB1 |
| -0.6408 | -0.0823 | *0.4545* | -0.0895 | 0.5476 | 0.8037 | Q16352 | INA |
| 0.3383 | -1.1693 | *0.3966* | -0.1448 | 0.8877 | 0.8041 | Q6IBS0;D6RG15; H7C5I2 | TWF2 |
| -0.0038 | 1.1002 | *-0.6613* | 0.1451 | 0.8901 | 0.8043 | Q15185;A0A087 WYT3 | PTGES3 |
| 1.1474 | -0.3755 | *-1.3963* | -0.2081 | 1.2801 | 0.8047 | Q96JE9 | MAP6 |
| -0.7908 | 0.9382 | *-0.6081* | -0.1535 | 0.9499 | 0.8058 | Q15746;F8WBL7 ;A0A1W2PPZ3;A 0A1W2PRF2;A0 A2R8Y4U5;B5BT Y5;B5MDL5;C9J WQ4;E7EX54;O1 5264;P53778;Q1 5759;Q5R3E6;Q8 6YV6;Q9H1R3 | MYLK |
| -1.6842 | 1.5284 | *0.9891* | 0.2778 | 1.7204 | 0.8060 | P27815 | PDE4A |
| 0.1462 | -0.0576 | *-0.1647* | -0.0254 | 0.1579 | 0.8071 | O75083;D6RD66 | WDR1 |
| 0.0112 | -0.1599 | *0.0881* | -0.0202 | 0.1270 | 0.8084 | O15020;A4QPE4 | SPTBN2 |
| 0.3199 | -0.5439 | *0.4885* | 0.0882 | 0.5538 | 0.8086 | P16885;A0A0A0 MRF9;H3BPZ3 | PLCG2 |
| -0.2681 | 0.8644 | *-0.2821* | 0.1047 | 0.6579 | 0.8086 | P16401 | HIST1H1B |
| -1.6034 | 0.8193 | *0.1861* | -0.1993 | 1.2565 | 0.8093 | Q7Z794 | KRT77 |
| -1.1289 | 0.2770 | *1.4706* | 0.2062 | 1.3012 | 0.8094 | Q9BV35 | SLC25A23 |
| 0.4425 | -0.6289 | *0.4865* | 0.1000 | 0.6317 | 0.8096 | Q52LW3;F8VWZ 8 | ARHGAP29 |
| 0.5783 | 0.4456 | *-1.6054* | -0.1938 | 1.2242 | 0.8096 | Q9Y5G1 | PCDHGB3 |
| 0.0429 | -0.9494 | *1.4872* | 0.1936 | 1.2253 | 0.8100 | P09104;F5H0C8; F5H1C3;U3KQP4 | ENO2 |
| 0.2528 | -1.9138 | *2.7707* | 0.3699 | 2.3444 | 0.8103 | P00918;E5RID5; E5RK37 | CA2 |
| -0.4711 | -0.2055 | *0.4525* | -0.0747 | 0.4755 | 0.8111 | P62241;Q5JR95; H0Y547;H0Y8F0 | RPS8 |
| -1.3588 | -0.5951 | *3.0607* | 0.3690 | 2.3622 | 0.8121 | P20701;I3L1D1 | ITGAL |
| 0.0629 | 1.0954 | *-1.8562* | -0.2326 | 1.4978 | 0.8131 | P42224;J3KPM9; D2KFR9;E7ENM 1;E7EPD2;H7BZ 88 | STAT1 |
| -0.0591 | -0.8184 | *0.5568* | -0.1069 | 0.6888 | 0.8132 | P49770 | EIF2B2 |
| -0.4024 | 0.3193 | *0.2705* | 0.0625 | 0.4033 | 0.8137 | O14773;A0A2R8 YD45;A0A2R8YG D1;A0A2R8Y7U1 | TPP1 |
| 0.2326 | -1.8173 | *0.9235* | -0.2204 | 1.4254 | 0.8139 | Q9NZC9;C9J6I8; C9J8F8;C9JP32; C9JS37 | SMARCAL1 |
| -0.1578 | 1.2945 | *-0.6656* | 0.1570 | 1.0172 | 0.8142 | O75367;B4DJC3; D6RCF2 | H2AFY |
| -2.5266 | -0.0599 | *1.6236* | -0.3210 | 2.0874 | 0.8149 | P07919 | UQCRH |
| 0.3956 | -0.5836 | *0.4566* | 0.0896 | 0.5838 | 0.8153 | Q9UK73;H3BT12 | FEM1B |

| L100/ CTRL1 | L100/ CTRL2 | L100/ CTRL3 | Mean | SD | T-test p-value | Accession | Gene Symbol |
|---|---|---|---|---|---|---|---|
| 0.0413 | -0.5304 | *0.7949* | 0.1019 | 0.6647 | 0.8154 | Q99613;B5ME19;H3BPE4;H3BPE3;H3BTY8 | EIF3C |
| -0.4174 | -1.8174 | *1.4748* | -0.2533 | 1.6522 | 0.8154 | Q96E40 | SPACA9 |
| 0.4845 | -0.2817 | *-0.4276* | -0.0749 | 0.4899 | 0.8159 | Q15147;A0A096LPH9;B1AJW2;B1AJW3;B1AJW4 | PLCB4 |
| -1.0063 | 0.1613 | *0.4858* | -0.1197 | 0.7848 | 0.8163 | P36578;H3BM89;H3BTP7;H3BU31;H0YLA4;K7ELG0 | RPL4 |
| 1.0293 | -0.3982 | *-0.2697* | 0.1205 | 0.7897 | 0.8163 | Q14789;E7EU81;H0Y867 | GOLGB1 |
| 0.3035 | 1.0392 | *-2.0877* | -0.2483 | 1.6348 | 0.8171 | Q6ZU80;H0YJE3 | CEP128 |
| 1.8033 | 0.8900 | *-4.1557* | -0.4875 | 3.2094 | 0.8171 | O00241;H3BV43;H3BML4;H3BRP9;H9KV29;Q5TFQ8;P78324 | SIRPB1 |
| -2.7828 | 0.1657 | *1.6005* | -0.3389 | 2.2348 | 0.8174 | Q0PNE2 | ELP6 |
| -0.2775 | 0.0406 | *0.1382* | -0.0329 | 0.2174 | 0.8177 | P08238 | HSP90AB1 |
| -0.3871 | -0.4745 | *1.3185* | 0.1523 | 1.0109 | 0.8186 | P61956;A8MU27;A8MUA9;P55854;Q6EEV6 | SUMO2 |
| -0.1881 | -0.5096 | *0.4718* | -0.0753 | 0.5004 | 0.8188 | O95071;E7EMW7;E7ET84;E5RFK7 | UBR5 |
| -1.2898 | 0.1673 | *0.6658* | -0.1522 | 1.0162 | 0.8196 | P07602;C9JIZ6;Q5BJH1;A0A0J9YXB8 | PSAP |
| 0.4264 | -3.3518 | *1.7442* | -0.3937 | 2.6451 | 0.8207 | A8MW92;A0A0A0MQS0;F8W9L8;A8MUE8;A8MXR8;E9PM57 | PHF20L1 |
| 0.7965 | 0.4893 | *-1.9602* | -0.2248 | 1.5107 | 0.8207 | O14977 | AZIN1 |
| 0.5835 | -0.5163 | *-0.3299* | -0.0875 | 0.5886 | 0.8208 | E9PG32;Q6ZR08;H7C5N3;J3QTM1 | DNAH12 |
| 0.1341 | 0.1296 | *-0.1828* | 0.0270 | 0.1817 | 0.8209 | Q3LXA3;H0YCY6;I3L252 | TKFC |
| 0.2858 | 0.8477 | *-0.7684* | 0.1217 | 0.8204 | 0.8212 | O00505 | KPNA3 |
| -0.2835 | -0.9860 | *1.9523* | 0.2276 | 1.5344 | 0.8213 | O94760;B4DYP1 | DDAH1 |
| -1.0433 | 0.8525 | *0.6545* | 0.1545 | 1.0421 | 0.8213 | Q96DB9;A8MVG1;F5H4X8;K7EJH6 | FXYD5 |
| 0.8508 | 0.1868 | *-0.6939* | 0.1146 | 0.7749 | 0.8218 | S4R3N1;Q9Y3A3 | HSPE1-MOB4 |
| 0.6310 | -0.1454 | *-0.2693* | 0.0721 | 0.4879 | 0.8220 | Q96DT5;A0A087WYC6;A0A0C4DFR0 | DNAH11 |
| -0.2078 | -1.2737 | *0.9842* | -0.1658 | 1.1295 | 0.8231 | O00764;F2Z2Y4 | PDXK |
| -0.3474 | 2.0725 | *-1.0116* | 0.2378 | 1.6232 | 0.8234 | Q9NQG5 | RPRD1B |
| -0.0159 | -1.3639 | *0.8828* | -0.1657 | 1.1308 | 0.8234 | Q04760 | GLO1 |
| -0.3708 | 0.2959 | *-0.0717* | -0.0488 | 0.3339 | 0.8237 | Q86UW6;H0YA93 | N4BP2 |

| L100/ CTRL1 | L100/ CTRL2 | L100/ CTRL3 | Mean | SD | T-test p-value | Accession | Gene Symbol |
|---|---|---|---|---|---|---|---|
| -0.1191 | -0.0858 | *0.3095* | 0.0349 | 0.2384 | 0.8237 | Q9BZ29;A0A0A0MSY4;A0A088AWN3;A0A0A0MT38;Q6ZSL5;A0A0D9SF41 | DOCK9 |
| 1.1244 | -1.0996 | *-0.5312* | -0.1688 | 1.1555 | 0.8239 | A0A087WZ84;C9JLX5;C9K0F2;C9JXQ5 | ZNF568 |
| 0.9491 | 1.9219 | *-4.3375* | -0.4888 | 3.3684 | 0.8250 | A6NK53;K7EN46 | ZNF233 |
| 2.2151 | -0.2436 | *-3.1339* | -0.3875 | 2.6774 | 0.8255 | Q13332;A0A0A0MR60;G8JL96 | PTPRS |
| -1.4174 | 1.1728 | *-0.3198* | -0.1881 | 1.3001 | 0.8255 | Q8TEK3;A0A087X1A7 | DOT1L |
| -0.0933 | -0.2009 | *0.4434* | 0.0497 | 0.3451 | 0.8262 | P07437;Q5JP53 | TUBB |
| -0.0119 | 0.2476 | *-0.3692* | -0.0445 | 0.3097 | 0.8266 | Q13948;A0A2R8Y852;A0A2R8YDI1;P39880 | CUX1 |
| 0.3671 | -0.0098 | *-0.5577* | -0.0668 | 0.4650 | 0.8268 | I3L119 | ZNF785 |
| 1.1907 | 1.8496 | *-2.1238* | 0.3055 | 2.1295 | 0.8269 | Q7RTS7;F8W1S1 | KRT74 |
| -0.3214 | 1.2571 | *-1.5314* | -0.1986 | 1.3983 | 0.8287 | A8MU46;E9PPJ3 | SMTNL1 |
| 0.1698 | -0.8927 | *0.4259* | -0.0990 | 0.6992 | 0.8291 | P30044 | PRDX5 |
| -0.2235 | -0.4886 | *0.4963* | -0.0719 | 0.5096 | 0.8297 | Q9BV86;S4R3J7 | NTMT1 |
| -2.8347 | -0.0789 | *1.9108* | -0.3343 | 2.3830 | 0.8307 | H0YBS9;Q9H0M0 | WWP1 |
| -0.4118 | 0.1261 | *0.4734* | 0.0625 | 0.4460 | 0.8307 | P78318 | IGBP1 |
| 0.0966 | -0.2018 | *0.1913* | 0.0287 | 0.2052 | 0.8310 | Q92804;A0A075B7D9 | TAF15 |
| -0.4784 | 0.4167 | *-0.1274* | -0.0630 | 0.4510 | 0.8313 | P56192;H0YHV5;H0YIP0;F5H2V6;F8VS26;F8VZZ9;F8W0M7;F8W0S4;H0YHL6;H0YI27;H0YI94 | MARS |
| -0.7545 | -0.1722 | *0.6349* | -0.0973 | 0.6977 | 0.8317 | P51812;B1AXG1;B4DG22;B7ZB17;A0A2R8Y7S9;D6R910;A0A2R8Y603;D6RHW7 | RPS6KA3 |
| 0.0468 | 2.9380 | *-1.9582* | 0.3422 | 2.4614 | 0.8321 | H0Y360;Q01433;E9PIJ1;E9PJF6;H0YCL9;H0YF16 | AMPD2 |
| 3.6663 | -2.7842 | *-2.3864* | -0.5014 | 3.6148 | 0.8325 | P32019 | INPP5B |
| -0.0684 | -0.1461 | *0.1507* | -0.0213 | 0.1539 | 0.8328 | P55786;E9PLK3;E9PJY4;E9PPD4;H0YCQ5;H0YDG0 | NPEPPS |
| -0.2465 | -0.1628 | *0.6032* | 0.0646 | 0.4683 | 0.8333 | F8VXC8;F8VZW6 | SMARCC2 |
| -0.2885 | -2.3473 | *1.7856* | -0.2834 | 2.0665 | 0.8344 | K7EM48;K7EN63;Q96SI1;K7EPF0;K7EQS3 | KCTD15 |
| 2.2330 | -3.3136 | *2.4161* | 0.4452 | 3.2564 | 0.8349 | Q96KR1;H0Y8W1 | ZFR |

| L100/ CTRL1 | L100/ CTRL2 | L100/ CTRL3 | Mean | SD | T-test p-value | Accession | Gene Symbol |
|---|---|---|---|---|---|---|---|
| 0.0695 | -0.4033 | *0.2032* | -0.0435 | 0.3187 | 0.8349 | P31946;A0A0J9YWE8;Q4VY19;Q4VY20;A0A0J9YWZ2;B5BU24 | YWHAB |
| 0.0266 | 0.2176 | *-0.3657* | -0.0405 | 0.2974 | 0.8355 | P01024;M0R0Q9 | C3 |
| 2.2631 | -0.2854 | *-1.2396* | 0.2460 | 1.8108 | 0.8359 | Q12972;A0A0A0MT09 | PPP1R8 |
| -2.6409 | -6.5816 | *6.4909* | -0.9105 | 6.7058 | 0.8360 | Q8N3L3 | TXLNB |
| -3.1367 | 8.3446 | *-2.5734* | 0.8782 | 6.4723 | 0.8361 | B4DTR8;Q9Y5X9 | LIPG |
| -0.7032 | 0.1589 | *0.8637* | 0.1065 | 0.7848 | 0.8361 | E9PIK4;E9PIN5;E9PKN9;E9PKZ4;E9PMW4;E9PPM1;E9PQ46;E9PS55;O14683;U3KQ32;E9PMY0;E9PN66;E9PNB3 | TP53I11 |
| 1.9549 | 0.1658 | *-3.1762* | -0.3518 | 2.6045 | 0.8368 | E5RFR7;H0YC42;H0YC44;P55327 | TPD52 |
| -0.1273 | 0.8061 | *-0.4195* | 0.0865 | 0.6401 | 0.8368 | A0A1B0GU45;F5H006;F8W9L6;Q12912;H0YJC6 | LRMP |
| -0.4474 | 0.0102 | *0.2871* | -0.0500 | 0.3710 | 0.8371 | M0QYS1;P40429;Q8J015;A0A096LPE0;M0QZU1 | RPL13A |
| -0.2227 | -0.7179 | *1.3842* | 0.1479 | 1.0990 | 0.8374 | Q8NDM7 | CFAP43 |
| -0.5306 | 0.7304 | *0.0545* | 0.0848 | 0.6310 | 0.8376 | Q8WWM7;H3BUF6;H3BSK9;H3BSQ5 | ATXN2L |
| 0.8931 | -0.7853 | *0.2324* | 0.1134 | 0.8455 | 0.8379 | Q8NFC6;Q13495 | BOD1L1 |
| 0.6432 | -0.6152 | *0.2270* | 0.0850 | 0.6411 | 0.8397 | P81605 | DCD |
| -0.2679 | -0.6531 | *1.3419* | 0.1403 | 1.0583 | 0.8398 | P78371;F8VQ14 | CCT2 |
| 0.1037 | -1.8638 | *2.6627* | 0.3009 | 2.2697 | 0.8398 | P60510;H3BTA2;H3BV22;I3L4X0 | PPP4C |
| 0.5417 | -0.4624 | *0.1212* | 0.0668 | 0.5042 | 0.8398 | Q9Y2J2;A0A0A0MRA8;A0A1B0GTF8;A0A0J9YY18;J3KRD1;J3KS70;J3QKK4;J3QKY2;J3QR33;J3QS55;J3QRQ6;J3KT37;J3QLU5;J3QS83;A0A0A0MSA4 | EPB41L3 |
| -0.2352 | -0.1856 | *0.3031* | -0.0392 | 0.2975 | 0.8406 | Q9P2J5;A0A087WXY1 | LARS |
| 2.0315 | 0.2654 | *-1.5830* | 0.2380 | 1.8074 | 0.8408 | Q8TES7;A0A0R4J2E4;K7ENL6;K7ESG2 | FBF1 |
| 0.0510 | 0.9573 | *-0.6839* | 0.1081 | 0.8221 | 0.8410 | P24752;H0YEL7;E9PKF3;E9PRQ6 | ACAT1 |
| 0.0162 | -0.3367 | *0.2120* | -0.0362 | 0.2781 | 0.8426 | Q9NSE4 | IARS2 |
| 0.1346 | -1.4791 | *0.8754* | -0.1564 | 1.2039 | 0.8429 | L8E6V6 | LAMA3 |
| -0.2466 | -0.7970 | *0.7432* | -0.1001 | 0.7804 | 0.8448 | Q9NYV4;J3QSD7 | CDK12 |
| 0.3230 | -1.0062 | *0.3827* | -0.1002 | 0.7852 | 0.8456 | Q9HB75 | PIDD1 |

| L100/ CTRL1 | L100/ CTRL2 | L100/ CTRL3 | Mean | SD | T-test p-value | Accession | Gene Symbol |
|---|---|---|---|---|---|---|---|
| -0.1010 | -0.5609 | *0.9605* | 0.0995 | 0.7803 | 0.8457 | P20807;F8W8F5 | CAPN3 |
| -0.6353 | 1.2129 | *-0.2077* | 0.1233 | 0.9676 | 0.8458 | Q9Y2E4;A0A0U1RQW6;E7EPU2 | DIP2C |
| -0.6158 | 1.8309 | *-0.6708* | 0.1815 | 1.4288 | 0.8463 | P35613;A0A087X2B5;A0A087WUV8;I3L192;R4GMX5;R4GN83 | BSG |
| -0.1139 | 1.9665 | *-2.7511* | -0.2995 | 2.3643 | 0.8467 | O14980;C9IZS4;C9J673;C9JKM9;C9JQ02;C9JV99;F8WF71;C9IYM2;C9JF49;H7BZC5 | XPO1 |
| -1.2993 | -0.1219 | *0.9876* | -0.1445 | 1.1437 | 0.8470 | M0R063;M0R106;P48664 | SLC1A6 |
| -0.6693 | 0.6142 | *-0.1901* | -0.0817 | 0.6486 | 0.8475 | B1AMU4;B1AMU7;Q9Y3B2;R4GMQ7;R4GNH9 | EXOSC1 |
| -1.3358 | 2.4156 | *-1.9712* | -0.2971 | 2.3707 | 0.8483 | Q5VT97 | SYDE2 |
| 0.2068 | 0.8429 | *-1.5057* | -0.1520 | 1.2147 | 0.8485 | Q96GD0;B1AHD3;Q6ZT62 | PDXP |
| -0.3290 | -0.1246 | *0.3279* | -0.0419 | 0.3362 | 0.8490 | P30622;J3KP58;F5H6A0;F6VGP8;F5H1T5;F5H270;F5H367 | CLIP1 |
| 1.4318 | 0.6053 | *-2.8958* | -0.2863 | 2.2974 | 0.8491 | Q6YN16 | HSDL2 |
| 2.6286 | 0.1764 | *-1.9493* | 0.2852 | 2.2909 | 0.8493 | Q9UI17;E5RG50;E5RK15 | DMGDH |
| 1.7751 | 1.3601 | *-2.2985* | 0.2789 | 2.2417 | 0.8494 | A0A2R8YGH5;H7C1E4;P61966 | AP1S1 |
| -3.0105 | -0.0728 | *2.1262* | -0.3190 | 2.5772 | 0.8501 | Q9BTE6;C9J5N1;L7N2F4;K7N799 | AARSD1 |
| 0.8863 | 1.5930 | *-3.5026* | -0.3411 | 2.7607 | 0.8504 | P36543;C9J8H1 | ATP6V1E1 |
| -0.4685 | 0.0750 | *0.2544* | -0.0464 | 0.3764 | 0.8507 | Q15819;G3V113;H0YBX6;H0YBP9 | UBE2V2 |
| -0.6670 | -0.6073 | *0.9396* | -0.1116 | 0.9108 | 0.8516 | Q14847;C9J9W2;F6S2S5;K7ESD6 | LASP1 |
| 0.0106 | -0.5048 | *0.7198* | 0.0752 | 0.6149 | 0.8518 | Q00325;F8VVM2;F8VWQ0 | SLC25A3 |
| -0.5112 | -0.4912 | *1.4055* | 0.1344 | 1.1009 | 0.8522 | Q14254;E7EMK3;J3QLD9;K7EKW9 | FLOT2 |
| 0.6375 | 1.4928 | *-3.0048* | -0.2915 | 2.3884 | 0.8522 | Q9BXU1;C9JPB1;C9JQW5 | STK31 |
| -0.6012 | -0.2522 | *1.2026* | 0.1164 | 0.9567 | 0.8526 | Q8NCU4;C9JA89 | CCDC191 |
| 2.3830 | 3.3782 | *-4.2517* | 0.5032 | 4.1478 | 0.8530 | A1A4V9;H3BTQ9;H0YGC5 | CCDC189 |
| 1.3731 | 1.7254 | *-2.2905* | 0.2693 | 2.2239 | 0.8533 | A0A1B0GW10;E9PHH0;Q9P267;A0A0D9SG23;A0A1B0GUJ9 | MBD5 |
| 1.3727 | -1.3385 | *0.4652* | 0.1665 | 1.3800 | 0.8538 | O43592;F8WDU6;F5GYW6;F5GZM3 | XPOT |
| -0.0767 | 0.3568 | *-0.4202* | -0.0467 | 0.3894 | 0.8547 | Q13136;A0A2R8Y7R9;E9PJZ7;H0YDW2 | PPFIA1 |

| L100/ CTRL1 | L100/ CTRL2 | L100/ CTRL3 | Mean | SD | T-test p-value | Accession | Gene Symbol |
|---|---|---|---|---|---|---|---|
| 0.0708 | -1.0741 | *0.6833* | -0.1067 | 0.8920 | 0.8551 | Q6UB98;F5GYX2;J3KSU9;J3QRX3 | ANKRD12 |
| -0.0447 | -0.2634 | *0.2215* | -0.0289 | 0.2429 | 0.8558 | O60934;A0A0C4DG07 | NBN |
| 0.3289 | 0.2949 | *-0.4642* | 0.0532 | 0.4484 | 0.8562 | P43487;F6WQW2;C9JJ34;C9JXG8;C9JGV6;C9JDM3;C9JIC6 | RANBP1 |
| -0.2745 | -0.1244 | *0.5557* | 0.0523 | 0.4424 | 0.8568 | P61247;D6RAT0;D6RG13;H0Y9Y4;D6RB09;E9PFI5;D6R9B6;H0Y8L7;D6RAS7;D6RED7;D6RI02;D6RGE0 | RPS3A |
| -0.1177 | -1.5036 | *1.1543* | -0.1557 | 1.3293 | 0.8580 | O15294 | OGT |
| 0.7252 | -0.0213 | *-0.4896* | 0.0714 | 0.6127 | 0.8587 | P19012 | KRT15 |
| 0.5559 | 0.7355 | *-0.9668* | 0.1082 | 0.9353 | 0.8597 | P17036;C9J5S8;C9JE35;C9JIW8;C9JK31 | ZNF3 |
| 0.6194 | -0.0045 | *-0.4322* | 0.0609 | 0.5288 | 0.8603 | Q9C0D2 | CEP295 |
| 0.1226 | 0.9959 | *-0.8075* | 0.1037 | 0.9018 | 0.8606 | M0QWZ7;Q9NP81;B4DJM9;M0R2H5;M0R1C0;M0R2C6 | SARS2 |
| 1.0736 | -1.0214 | *0.3117* | 0.1213 | 1.0604 | 0.8613 | Q8TDM6 | DLG5 |
| 1.3933 | -1.7128 | *0.8896* | 0.1900 | 1.6670 | 0.8617 | Q96DN5;E5RFG6;E7ERK7;E7EWW7 | TBC1D31 |
| 0.4625 | -0.2539 | *-0.3612* | -0.0509 | 0.4478 | 0.8622 | Q16822;H0YML5;A0A0A0MS74;H0YM31 | PCK2 |
| -0.0850 | -0.4015 | *0.3575* | -0.0430 | 0.3813 | 0.8632 | P07237;H7BZ94;H0Y3Z3;I3L398;I3L312;I3L3U6;I3NI03;I3L3P5;I3L0S0;I3L4M2;I3L514;I3L1Y5 | P4HB |
| -0.4226 | -0.0128 | *0.6112* | 0.0586 | 0.5206 | 0.8634 | P37840;E7EPV7;H6UYS7;D6RA31 | SNCA |
| 0.2051 | -0.1588 | *0.0151* | 0.0205 | 0.1820 | 0.8636 | Q9Y4L1;A0A087X054;A0A087WWI4;E9PJ21;K7EQK2;J3KTF1;J3QL06 | HYOU1 |
| -0.2688 | 1.3886 | *-0.7424* | 0.1258 | 1.1190 | 0.8636 | O43295;A0A087WZH4 | SRGAP3 |
| -0.4683 | 1.6971 | *-0.7753* | 0.1512 | 1.3476 | 0.8639 | Q6ZRK6;H0YDV2 | CCDC73 |
| -0.7177 | -0.5738 | *1.7478* | 0.1521 | 1.3838 | 0.8666 | P13010;C9JZ81 | XRCC5 |
| -1.2112 | 1.2563 | *0.3667* | 0.1373 | 1.2497 | 0.8667 | A0A0G2JNQ3;B2RXH8 | HNRNPCL2 |
| -0.3607 | -0.2215 | *0.7889* | 0.0689 | 0.6274 | 0.8667 | J3QRC4;J3QQQ9;J3QQV1;J3QRI7;J3KSS0;P61254 | RPL26 |

| L100/ CTRL1 | L100/ CTRL2 | L100/ CTRL3 | Mean | SD | T-test p-value | Accession | Gene Symbol |
|---|---|---|---|---|---|---|---|
| -0.8510 | 0.3433 | *0.7863* | 0.0928 | 0.8469 | 0.8669 | A6NI56;B7ZBA8 | CCDC154 |
| 0.1990 | -0.4150 | *0.1073* | -0.0362 | 0.3312 | 0.8673 | Q8WZA9;M0QZP8 | IRGQ |
| -1.1009 | 1.3238 | *-0.6432* | -0.1401 | 1.2883 | 0.8680 | Q5SRE5;H7C4K7 | NUP188 |
| -0.6696 | 1.9312 | *-0.7628* | 0.1662 | 1.5292 | 0.8680 | Q96KP4;J3QKT2;J3QRA8;J3QRH4;A0A087WYZ1;J3KSV5;J3QKQ0;J3QL02;J3QLU1;J3QQN6;J3QR27;J3QRD0;A0A087WVS2;J3KRD5 | CNDP2 |
| -0.2736 | 4.0899 | *-2.7013* | 0.3717 | 3.4412 | 0.8689 | A8MYR2 | MSRB1 |
| -0.4841 | -0.4930 | *1.3126* | 0.1119 | 1.0399 | 0.8694 | Q96MU6;A0A0A0MSW5;H3BSD4;H3BUU4 | ZNF778 |
| -0.4435 | -1.4879 | *1.4523* | -0.1597 | 1.4905 | 0.8699 | Q14590;K7EL19 | ZNF235 |
| 0.7178 | -1.6614 | *0.5193* | -0.1414 | 1.3201 | 0.8699 | P24821;F5H7V9;J3QSU6;E9PC84;H0YGZ3 | TNC |
| 0.8534 | 0.4444 | *-0.9875* | 0.1034 | 0.9667 | 0.8701 | P07197;E7ESP9;E7EMV2 | NEFM |
| -0.2514 | -1.5349 | *1.3274* | -0.1530 | 1.4337 | 0.8704 | Q9H0H5 | RACGAP1 |
| -3.1622 | 3.6001 | *0.6448* | 0.3609 | 3.3901 | 0.8707 | O75478 | TADA2A |
| 1.1979 | -2.2241 | *0.4515* | -0.1916 | 1.7994 | 0.8707 | Q5VTH9;H0YCX1;H0YES1;Q5TAD8 | WDR78 |
| 0.5782 | 0.0197 | *-0.8221* | -0.0747 | 0.7049 | 0.8712 | Q9NUU7;I3L0H8;H3BTB3 | DDX19A |
| -0.1416 | 0.2135 | *-0.1366* | -0.0215 | 0.2036 | 0.8714 | P35813;B5BUD5;E9PKB5 | PPM1A |
| 0.1129 | 0.3669 | *-0.3624* | 0.0391 | 0.3702 | 0.8716 | Q9H4B7 | TUBB1 |
| -0.9477 | 1.2017 | *0.0868* | 0.1136 | 1.0750 | 0.8716 | Q9UII2 | ATP5IF1 |
| 0.4131 | 3.1827 | *-2.6703* | 0.3085 | 2.9279 | 0.8720 | F8WEZ9;Q8WV37 | ZNF480 |
| -0.4595 | -1.2953 | *1.3321* | -0.1409 | 1.3424 | 0.8725 | D6RJC6 | DTNBP1 |
| 0.1686 | 0.0238 | *-0.2608* | -0.0228 | 0.2185 | 0.8732 | A0A087WTU9;Q8TCU4 | ALMS1 |
| 0.5506 | -0.2074 | *-0.5147* | -0.0572 | 0.5483 | 0.8733 | Q15691 | MAPRE1 |
| -0.9304 | 0.7634 | *0.4481* | 0.0937 | 0.9008 | 0.8736 | P0DP23;P0DP24;P0DP25;H0Y7A7;E7EMB3;E7ETZ0;G3V361;Q96HY3;G3V226;F8WBR5;G3V479;M0QZ52 | CALM1 |
| 0.2394 | 1.0603 | *-0.9797* | 0.1067 | 1.0264 | 0.8737 | O75334;G3V200;H0YHK3;H0YH95;F8VU88;F8W1Y8;H0YEF9;A0A1B0GVT3;A0A286YEX6;H0YHJ4;H0YIJ4;Q8NGP3 | PPFIA2 |
| -0.2410 | -0.4878 | *0.5589* | -0.0566 | 0.5472 | 0.8742 | Q8N5A5;V9GY48 | ZGPAT |

| L100/ CTRL1 | L100/ CTRL2 | L100/ CTRL3 | Mean | SD | T-test p-value | Accession | Gene Symbol |
|---|---|---|---|---|---|---|---|
| -0.0419 | -0.4704 | *0.3807* | -0.0439 | 0.4255 | 0.8747 | P0CG38 | POTEI |
| -0.2844 | -0.1827 | *0.3601* | -0.0357 | 0.3465 | 0.8749 | Q07955;J3KTL2;J3KSR8;J3KSW7;J3QQV5 | SRSF1 |
| -1.2559 | 0.3456 | *0.5997* | -0.1035 | 1.0060 | 0.8750 | P61006 | RAB8A |
| -0.4174 | -0.3339 | *0.5807* | -0.0569 | 0.5537 | 0.8752 | Q09428;A0A2R8Y4V0;A0A2R8Y4Z4;A0A2R8Y5D8;A0A2R8YDG0;A0A2R8Y6Q0;A0A2R8YEE5;A0A2R8YGQ6;A0A2R8YHG6 | ABCC8 |
| -0.1116 | -0.7677 | *0.6593* | -0.0733 | 0.7142 | 0.8752 | Q9P253;H0YMC9 | VPS18 |
| -0.5286 | 0.7952 | *-0.4994* | -0.0776 | 0.7561 | 0.8753 | A0A2R8Y5Z6;A0A2R8Y5G2;A0A2R8Y570;A0A2R8Y6D0;A0A2R8Y4N6 | EPB41 |
| 0.1585 | 0.4300 | *-0.7847* | -0.0654 | 0.6375 | 0.8754 | Q5M9N0 | CCDC158 |
| -1.2031 | 2.7911 | *-2.4231* | -0.2783 | 2.7273 | 0.8760 | E7EVZ1;Q86UP3 | ZFHX4 |
| -0.6214 | 0.1223 | *0.3450* | -0.0514 | 0.5061 | 0.8766 | F5GZQ3;P55084;B5MD38;C9JE81;C9JEY0 | HADHB |
| -1.0417 | -0.4919 | *1.1823* | -0.1171 | 1.1584 | 0.8771 | F8WAD8;Q9P0K1 | ADAM22 |
| -0.2449 | -0.1972 | *0.5828* | 0.0469 | 0.4647 | 0.8774 | P84077;F5H423;P61204;F5H0C7;C9J1Z8;P84085;F5H1V1;F5H6T5;P62330 | ARF1 |
| -0.2695 | 0.0637 | *0.2911* | 0.0284 | 0.2820 | 0.8774 | Q9UEY8 | ADD3 |
| -0.1269 | -0.3594 | *0.3735* | -0.0376 | 0.3745 | 0.8779 | E9PAV3;F8VZJ2;F8W0W4;H0YHX9;Q13765;F8VNW4;F8W1N5;A0A087X1U4;A0A1W2PRG7;C9J1X3;C9J235;H0Y5H7;O95405;P0C7T7;P45954;Q6ZNE9 | NACA |
| -1.0013 | 0.2385 | *1.0780* | 0.1051 | 1.0460 | 0.8779 | Q7L7X3;J3QS76 | TAOK1 |
| 0.1641 | -1.0263 | *0.6079* | -0.0848 | 0.8450 | 0.8781 | Q00169;F5GWE5;I3L471;I3L4H1;I3L4U7;I3L459;I3L4C0;I3L2X8;I3L3W1 | PITPNA |
| -2.1779 | 0.1901 | *1.4377* | -0.1834 | 1.8365 | 0.8786 | Q8N841 | TTLL6 |
| 0.9241 | -1.7031 | *1.2616* | 0.1609 | 1.6230 | 0.8795 | A7E2Y1;A0A087X0T3;Q5JW45;Q5JW46 | MYH7B |
| -0.3693 | 0.3692 | *0.1111* | 0.0370 | 0.3748 | 0.8799 | J3KR97;Q9BTW9;A0A0J9YVR1;A0A0J9YWI5;A0A0J | TBCD |

| L100/ CTRL1 | L100/ CTRL2 | L100/ CTRL3 | Mean | SD | T-test p-value | Accession | Gene Symbol |
|---|---|---|---|---|---|---|---|
| | | | | | | 9YXU4;I3L120;I3L0V3;P52630 | |
| 0.4654 | -0.8921 | *0.2129* | -0.0713 | 0.7220 | 0.8800 | P54886 | ALDH18A1 |
| 0.2356 | -0.5566 | *0.4808* | 0.0533 | 0.5422 | 0.8805 | A0A0A0MTH0;Q8N0W4;A6NMU8;B4DHI3;Q8NFZ3 | NLGN4X |
| -2.4763 | 0.6754 | *2.5478* | 0.2490 | 2.5391 | 0.8808 | J3QRH7 | MARCH10 |
| 0.4934 | -0.5885 | *-0.0639* | -0.0530 | 0.5410 | 0.8809 | P43034;I3L495;I3L3N5 | PAFAH1B1 |
| 0.0908 | 0.5398 | *-0.4809* | 0.0499 | 0.5116 | 0.8814 | O75533;B4DGZ4;F8WC19 | SF3B1 |
| -0.9309 | 0.4887 | *0.6984* | 0.0854 | 0.8863 | 0.8828 | E9PLG0;E9PM30;Q17RS7 | GEN1 |
| -0.7920 | 0.7913 | *-0.2310* | -0.0772 | 0.8028 | 0.8830 | P29317 | EPHA2 |
| -0.1961 | -1.5912 | *2.3643* | 0.1924 | 2.0062 | 0.8834 | A0A0J9YY01;A0A286YF23 | MYO15B |
| -2.5744 | 2.5406 | *0.7756* | 0.2473 | 2.5981 | 0.8842 | P56524;F5H0B1 | HDAC4 |
| 1.3652 | -0.4626 | *-1.2900* | -0.1291 | 1.3587 | 0.8844 | P43155;A6PVN3;B7ZBP5 | CRAT |
| -0.0639 | -0.9950 | *1.4032* | 0.1148 | 1.2091 | 0.8845 | Q01813;Q5VSR5;B1APP6;V9GY25;V9GYV7 | PFKP |
| 0.0032 | 3.1952 | *-4.2627* | -0.3548 | 3.7418 | 0.8847 | Q14839;A0A0C4DGG9;A0A2R8Y212;A0A2R8Y425;A0A2R8Y521;A0A2R8Y5J0;A0A2R8YFK9;A0A2U3TZM0;F5GWX5;A0A2R8YDJ9;A0A2R8YFD8;A0A2R8YDW2;A0A2R8YD40;A0A2R8Y4X2;A0A2R8Y8C1;A0A2R8YER1;A0A2R8Y5Z7;A0A2R8Y685;A0A2R8Y8B3;A0A2R8Y5M9;A0A2R8Y7I0;A0A2R8Y7M9;A0A2R8YE38;K7EMY3;A0A2R8Y445;F2Z2R5 | CHD4 |
| -0.2706 | -0.7810 | *0.8192* | -0.0775 | 0.8174 | 0.8847 | P07384;E9PRM1 | CAPN1 |
| -0.4920 | -2.0162 | *1.9500* | -0.1861 | 2.0007 | 0.8868 | Q5VU43;A0A0A0MRM1;E9PQG4;A0A087WYE4;E9PS60;H0YCY0;A0A087X229;A0A0C4DG90;A0A1B0GTF3;A4D0S4;E9PJU0;E9PNE0;K7EJC3;O43293;O43301;Q53EU6;Q9BVT6;A0A075B749;A0A087WX83;A0A087WVQ4 | PDE4DIP |

| L100/ CTRL1 | L100/ CTRL2 | L100/ CTRL3 | Mean | SD | T-test p-value | Accession | Gene Symbol |
|---|---|---|---|---|---|---|---|
| -0.9350 | 1.0898 | *-0.4496* | -0.0983 | 1.0571 | 0.8869 | A0A087X0K1;Q9Y376 | CAB39 |
| -1.2760 | 0.6459 | *0.9676* | 0.1125 | 1.2132 | 0.8871 | Q9NQ35;E9PIW3 | NRIP3 |
| 0.3329 | -0.5277 | *0.0744* | -0.0401 | 0.4416 | 0.8894 | Q96NB2;A0A0C4DGR6;A0A1B0GX61;R4GMR9;R4GMW0;R4GN63;R4GN74;R4GNC2 | SFXN2 |
| -0.6993 | -0.4210 | *0.8895* | -0.0769 | 0.8485 | 0.8896 | P09211;A8MX94;A0A087X2E9;A0A087X243 | GSTP1 |
| -0.8748 | -0.7094 | *1.2620* | -0.1074 | 1.1888 | 0.8900 | O60493 | SNX3 |
| 1.0786 | -0.5427 | *-0.2999* | 0.0786 | 0.8744 | 0.8905 | O43294 | TGFB1I1 |
| -0.7166 | 0.3867 | *0.5118* | 0.0607 | 0.6760 | 0.8908 | O75746 | SLC25A12 |
| -0.8654 | 1.1688 | *-0.5996* | -0.0988 | 1.1057 | 0.8913 | O75368 | SH3BGRL |
| -0.4452 | -0.0464 | *0.3815* | -0.0367 | 0.4134 | 0.8920 | P19367;B1AR63;B1AR62 | HK1 |
| 0.5150 | -1.3267 | *1.1542* | 0.1142 | 1.2881 | 0.8921 | Q12767;C9JL75;J3KTL5;J3QQW3;J3QRY7;J3KRN3;J3KRU7;J3QLM7;J3QS17 | TMEM94 |
| -0.3936 | -0.6953 | *1.3874* | 0.0995 | 1.1255 | 0.8923 | P56385 | ATP5ME |
| -0.2030 | -0.3097 | *0.4100* | -0.0342 | 0.3884 | 0.8927 | A0A2U3TZH3;Q05639;A0A2R8Y488;A0A2R8Y660;A0A2R8YDN5 | EEF1A2 |
| 2.0718 | -0.6662 | *-1.9467* | -0.1804 | 2.0528 | 0.8930 | P54277;Q3BDU3;Q5FBZ9;C9JF76;E9PC40;E9PC65;F8W8L1;Q5FBZ4;Q5XG96 | PMS1 |
| 0.6000 | 0.2539 | *-0.6799* | 0.0580 | 0.6621 | 0.8933 | A6NKB5;H0YBF4 | PCNX2 |
| -0.0727 | 1.0032 | *-0.7039* | 0.0755 | 0.8632 | 0.8934 | P28070 | PSMB4 |
| -0.2113 | 0.6208 | *-0.2782* | 0.0438 | 0.5008 | 0.8936 | Q96JI7;C4B7M2;H0YN34;H0YLK7;H0YLR8 | SPG11 |
| -0.6654 | -0.2972 | *1.2188* | 0.0854 | 0.9987 | 0.8958 | Q8NEV8;E9PPH6 | EXPH5 |
| 2.5195 | -1.3347 | *-0.6588* | 0.1753 | 2.0581 | 0.8962 | P13646;K7ERE3;K7EMD9;K7EMJ2;K7EQH6 | KRT13 |
| -0.8923 | 0.6645 | *0.0315* | -0.0654 | 0.7829 | 0.8982 | Q9P2F5;H0YA59 | STOX2 |
| -0.1050 | -0.2847 | *0.4910* | 0.0338 | 0.4061 | 0.8987 | P60842;J3KT12;J3KSZ0;J3QL43;J3KTB5;J3QS69;J3QR64;J3QLN6;J3KTN0;J3KS25;J3QKZ9;J3QL52 | EIF4A1 |
| -0.1199 | 1.0356 | *-0.6963* | 0.0731 | 0.8819 | 0.8989 | P48741 | HSPA7 |
| -1.5818 | 1.1456 | *0.8044* | 0.1227 | 1.4860 | 0.8994 | P53618;E9PP73;E9PKQ1;E9PP63 | COPB1 |
| -0.1140 | -0.6010 | *0.9025* | 0.0625 | 0.7671 | 0.9007 | Q7RTP6;E9PP85;C9J922 | MICAL3 |

| L100/ CTRL1 | L100/ CTRL2 | L100/ CTRL3 | Mean | SD | T-test p-value | Accession | Gene Symbol |
|---|---|---|---|---|---|---|---|
| 1.0516 | -0.0932 | *-0.7377* | 0.0736 | 0.9062 | 0.9011 | Q9GZS3;H0YL19;H0YM76;H0YMF9;H0YN81 | WDR61 |
| -0.4451 | -1.2462 | *1.3668* | -0.1082 | 1.3387 | 0.9015 | P13861;H7C1L0;H7C330;C9J830 | PRKAR2A |
| 0.2611 | 0.0172 | *-0.2207* | 0.0192 | 0.2409 | 0.9027 | Q9HC56;B7ZM79;Q5VT82 | PCDH9 |
| -0.1168 | -0.6356 | *0.6049* | -0.0491 | 0.6230 | 0.9039 | O00232 | PSMD12 |
| -0.3672 | 2.2193 | *-2.3923* | -0.1801 | 2.3114 | 0.9050 | P98082 | DAB2 |
| -0.3916 | -0.6892 | *0.8862* | -0.0649 | 0.8370 | 0.9055 | P08779;K7ENV3;K7ENW6 | KRT16 |
| -0.8818 | 0.2659 | *0.8173* | 0.0671 | 0.8668 | 0.9056 | Q52LJ0;H0YNA1 | FAM98B |
| 0.7204 | -1.0095 | *0.0866* | -0.0675 | 0.8752 | 0.9060 | O43683;C9IYH4;C9JQA4;C9JRC7 | BUB1 |
| 0.2022 | 1.8238 | *-1.6275* | 0.1328 | 1.7267 | 0.9062 | A0A0G2JM70;Q9NV79;V9GZ41 | PCMTD2 |
| -0.4608 | -0.7166 | *1.4409* | 0.0879 | 1.1788 | 0.9091 | O94973;A0A0G2JS82;A0A0G2JQT9;H0YEG0;A0A0G2JS17;H0YDE9;E9PPY8;E9PQP4;A0A0G2JRS3;M0R2D9;E9PNC4;E9PPZ3;E9PS94;F8WAT4;P16066;Q96MA6;Q9BWT3A0A0G2JQM1 | AP2A2 |
| -0.3558 | -0.1734 | *0.4368* | -0.0308 | 0.4151 | 0.9095 | P36776;K7EJE8;K7EKE6;K7ERR6;K7EQF8;K7ER27 | LONP1 |
| -0.1261 | 1.5139 | *-1.0949* | 0.0977 | 1.3187 | 0.9097 | H3BLV9;Q96SB4;D6RBM8 | SRPK1 |
| -0.1261 | 1.6138 | *-1.1868* | 0.1003 | 1.4140 | 0.9135 | Q16531;F5GY55;F5GWI0;F5GZY8;F5H2L3 | DDB1 |
| 0.2123 | 1.1805 | *-1.1456* | 0.0824 | 1.1685 | 0.9140 | P14923;C9J826;C9JK18;C9JKY1;C9JTX4;C9JPI2;K7ERP3 | JUP |
| -0.3066 | 0.1810 | *0.0719* | -0.0179 | 0.2559 | 0.9148 | A0A0A0MRG2;E9PG40;H7C0V9;P05067 | APP |
| -2.5729 | 2.6367 | *-0.6144* | -0.1835 | 2.6314 | 0.9149 | F8WEH6;P50549;C9J4P4 | ETV1 |
| -0.3529 | 0.6976 | *-0.4798* | -0.0450 | 0.6463 | 0.9150 | Q6JQN1;F8W1I9;F8W179;D6RFF6 | ACAD10 |
| 0.1009 | 1.3933 | *-1.8317* | -0.1125 | 1.6230 | 0.9154 | Q86XA9;F5H619;H7C5W6;H0YIW3 | HEATR5A |
| -0.4124 | -0.9360 | *1.1262* | -0.0741 | 1.0719 | 0.9156 | P49748;G3V1M7;J3QRJ8;J3KSR4 | ACADVL |
| 0.1508 | 0.4863 | *-0.7700* | -0.0443 | 0.6505 | 0.9168 | P23527;Q16778;P06899;Q8N257 | HIST1H2BO |
| -1.9752 | 1.1747 | *1.1710* | 0.1235 | 1.8175 | 0.9171 | B1AKZ5 | PEA15 |

| L100/ CTRL1 | L100/ CTRL2 | L100/ CTRL3 | Mean | SD | T-test p-value | Accession | Gene Symbol |
|---|---|---|---|---|---|---|---|
| -0.4525 | -0.5501 | *0.8450* | -0.0525 | 0.7788 | 0.9177 | E5RHU1;Q96MH6;Q49A68 | TMEM68 |
| -0.3941 | 0.2718 | *0.1958* | 0.0245 | 0.3645 | 0.9179 | Q9Y277;E5RFP6;E5RHZ6;E5RJN6;E5RK27 | VDAC3 |
| 0.4538 | 0.6031 | *-1.2653* | -0.0695 | 1.0383 | 0.9183 | K7EK35;P42229 | STAT5A |
| 0.7211 | 0.4492 | *-1.3988* | -0.0762 | 1.1535 | 0.9194 | P31749;A0A087WY56;G3V3X1;G3V2I6 | AKT1 |
| 0.2682 | -0.5261 | *0.1724* | -0.0285 | 0.4336 | 0.9198 | Q9BQS8;C9J2W6;H7BZ74 | FYCO1 |
| -0.1436 | 2.0499 | *-2.3351* | -0.1429 | 2.1925 | 0.9204 | Q96BD8 | SKA1 |
| -0.1131 | 1.4936 | *-1.6901* | -0.1032 | 1.5919 | 0.9209 | Q96QS3 | ARX |
| -0.7904 | 0.6826 | *0.2546* | 0.0489 | 0.7577 | 0.9211 | Q16777;Q6FI13 | HIST2H2AC |
| 0.2761 | 0.3326 | *-0.5167* | 0.0306 | 0.4749 | 0.9212 | P78386 | KRT85 |
| 0.2916 | -1.4566 | *1.4472* | 0.0941 | 1.4619 | 0.9214 | O43824 | GTPBP6 |
| -0.4982 | 0.0657 | *0.5294* | 0.0323 | 0.5146 | 0.9234 | Q9H9A6 | LRRC40 |
| 0.3797 | 0.7035 | *-0.9222* | 0.0537 | 0.8605 | 0.9238 | P61353;K7ELC7 | RPL27 |
| 2.1877 | -0.2601 | *-1.5746* | 0.1177 | 1.9094 | 0.9247 | P61225 | RAP2B |
| -1.1590 | 1.0377 | *-0.0816* | -0.0676 | 1.0984 | 0.9248 | Q96S59 | RANBP9 |
| 3.4001 | 6.6963 | *-11.9304* | -0.6113 | 9.9402 | 0.9249 | P19784;H3BNI9 | CSNK2A2 |
| 1.9257 | -0.0250 | *-2.2893* | -0.1295 | 2.1094 | 0.9250 | P16118 | PFKFB1 |
| -1.5994 | 2.9681 | *-1.8640* | -0.1651 | 2.7166 | 0.9258 | Q92945;A0A087WTP3;M0R0I5;M0QYG1 | KHSRP |
| 1.6455 | -1.7957 | *0.4666* | 0.1054 | 1.7488 | 0.9264 | P62837 | UBE2D2 |
| 0.0759 | -0.0035 | *-0.0871* | -0.0049 | 0.0815 | 0.9266 | Q96C90 | PPP1R14B |
| -0.5202 | 0.0008 | *0.6223* | 0.0343 | 0.5719 | 0.9267 | Q00688 | FKBP3 |
| 0.8044 | 0.0167 | *-0.9810* | -0.0533 | 0.8948 | 0.9272 | Q13393;C9IY79 | PLD1 |
| -0.1171 | 1.5896 | *-1.2211* | 0.0838 | 1.4161 | 0.9277 | Q92823;F8W775;C9JYY6;A0A087X2B3;C9JH43;C9J8B6;C9JF43 | NRCAM |
| 3.9219 | -2.7761 | *-0.5467* | 0.1997 | 3.4108 | 0.9285 | Q13011;M0R248;M0QZW4;M0QXS7;M0R280 | ECH1 |
| -1.0046 | 0.0346 | *1.1600* | 0.0634 | 1.0826 | 0.9285 | O95373;E9PLB2 | IPO7 |
| -1.0604 | -0.3256 | *1.1860* | -0.0667 | 1.1454 | 0.9289 | O94776 | MTA2 |
| -0.3043 | 0.2716 | *0.0839* | 0.0171 | 0.2937 | 0.9291 | E7EPT4;P19404 | NDUFV2 |
| 0.6092 | 0.1185 | *-0.8566* | -0.0430 | 0.7461 | 0.9297 | H7BZK9;Q96N11 | C7orf26 |
| -0.9312 | 2.0984 | *-1.5005* | -0.1111 | 1.9345 | 0.9298 | Q8NBI6;A0A140T9D0;F8WEN6 | XXYLT1 |
| -1.1968 | 1.0631 | *-0.0606* | -0.0648 | 1.1300 | 0.9300 | O60343 | TBC1D4 |
| -0.3412 | 0.1486 | *0.1442* | -0.0161 | 0.2816 | 0.9300 | E7EVA0;B5MEG9;H7C4C5;H7C456;A0A0J9YVV8;A0A0J9YW37;F8W9U4;P27816 | MAP4 |
| 1.2528 | 0.7735 | *-1.7508* | 0.0918 | 1.6136 | 0.9305 | Q96RY5;J3QT63 | CRAMP1 |

| L100/ CTRL1 | L100/ CTRL2 | L100/ CTRL3 | Mean | SD | T-test p-value | Accession | Gene Symbol |
|---|---|---|---|---|---|---|---|
| -0.3500 | 0.2220 | *0.1823* | 0.0181 | 0.3194 | 0.9308 | Q9Y5H0 | PCDHGA3 |
| -0.2996 | -0.4336 | *0.6350* | -0.0327 | 0.5822 | 0.9313 | K7EK33;K7EQ55;Q96EP5;K7EQ02 | DAZAP1 |
| -0.5772 | 0.1679 | *0.5025* | 0.0310 | 0.5527 | 0.9314 | Q8IYD8;H0YJS3;H0YJ14 | FANCM |
| 0.1652 | -0.5413 | *0.3003* | -0.0252 | 0.4519 | 0.9317 | O60333;A0A087WWA3;Q4R9M9 | KIF1B |
| 0.1589 | 0.1810 | *-0.3933* | -0.0178 | 0.3254 | 0.9331 | A6NCI4 | VWA3A |
| 0.2800 | 0.0132 | *-0.3421* | -0.0163 | 0.3121 | 0.9363 | Q9ULE4 | FAM184B |
| -0.5589 | 0.0169 | *0.4627* | -0.0264 | 0.5122 | 0.9370 | P78559 | MAP1A |
| 0.2688 | 0.1547 | *-0.4863* | -0.0209 | 0.4070 | 0.9372 | P52292;J3KS65;J3QLL0 | KPNA2 |
| 0.4526 | -0.4806 | *-0.0430* | -0.0237 | 0.4669 | 0.9380 | Q6ZMV8 | ZNF730 |
| -0.6659 | -0.9365 | *1.4105* | -0.0640 | 1.2841 | 0.9391 | P18858;F5GZ28;M0R0Q7;B4E135;M0QY71;M0R1G7;M0R1S4 | LIG1 |
| 0.5025 | -0.3365 | *-0.1015* | 0.0215 | 0.4328 | 0.9393 | Q9NVI7;Q5SV16 | ATAD3A |
| 0.1750 | 1.3671 | *-1.3414* | 0.0669 | 1.3575 | 0.9397 | Q9NRW1;J3KR73;C9JU14 | RAB6B |
| -0.2794 | 0.5389 | *-0.3308* | -0.0238 | 0.4880 | 0.9405 | P19338;H7BY16;C9J1H7;C9JLB1;C9JWL1;C9JYW2 | NCL |
| -0.2922 | 0.5093 | *-0.2839* | -0.0223 | 0.4604 | 0.9408 | Q01130;J3KP15;J3QL05 | SRSF2 |
| -1.3031 | -0.0036 | *1.5088* | 0.0674 | 1.4073 | 0.9415 | Q96PU8;F5GXS8;F5GYM3;F5H8C8;H0YG47;F5H5U6;F5GYT7;H0YGD6 | QKI |
| -0.2986 | 0.0282 | *0.3138* | 0.0145 | 0.3064 | 0.9423 | Q9H0X9;H0YCD7;E9PNH0;E9PIJ6;E9PJE6;E9PLN3;E9PPQ2;E9PQB4;E9PRA9 | OSBPL5 |
| -0.2833 | -1.0262 | *1.1531* | -0.0521 | 1.1079 | 0.9425 | A0A2R8YFH5;Q15437;Q5QPE2;A0A2R8Y633;A0A2R8Y7S7;A0A2R8YF30 | SEC23B |
| -1.5181 | 0.8813 | *0.8298* | 0.0643 | 1.3707 | 0.9426 | Q99941 | ATF6B |
| 0.0084 | -0.2163 | *0.1801* | -0.0093 | 0.1988 | 0.9429 | P17174 | GOT1 |
| 1.0041 | -1.2485 | *0.0868* | -0.0525 | 1.1328 | 0.9433 | Q9NQ66;A0A087WT80;A0A087WW73;A0A1B0GWB6;A0A0D9SF51;A0A0D9SG17;H0YCJ2;A0A0D9SFJ4;A0A0D9SGI7;A0A0D9SFE7;A0A1B0GVC1;B1AK73 | PLCB1 |
| 0.2793 | -0.1472 | *-0.1669* | -0.0116 | 0.2521 | 0.9437 | Q9H307 | PNN |

| L100/ CTRL1 | L100/ CTRL2 | L100/ CTRL3 | Mean | SD | T-test p-value | Accession | Gene Symbol |
|---|---|---|---|---|---|---|---|
| -0.6304 | -0.1767 | *0.9154* | 0.0361 | 0.7945 | 0.9445 | P41250;H7C443 | GARS |
| 1.7961 | -3.5087 | *1.3138* | -0.1329 | 2.9334 | 0.9446 | Q9BUY5 | ZNF426 |
| 0.0866 | -0.8628 | *0.8943* | 0.0393 | 0.8795 | 0.9453 | Q01105;A0A0C4DFV9 | SET |
| 0.8211 | 1.1348 | *-2.2018* | -0.0820 | 1.8425 | 0.9456 | Q9HC10;A0A2U3TZT7 | OTOF |
| -0.3985 | -0.7497 | *1.0246* | -0.0412 | 0.9396 | 0.9464 | A0A087WY71;E9PFW3;Q96CW1;C9JJ47;C9JTK4;C9JJD3;C9JPV8;C9JGT8 | AP2M1 |
| 0.6189 | 0.5747 | *-1.3399* | -0.0488 | 1.1184 | 0.9467 | P28827;E7EPS8 | PTPRM |
| 0.2589 | -0.6754 | *0.4975* | 0.0270 | 0.6199 | 0.9467 | K7EQJ5;P62841;K7EJ78;K7ELC2 | RPS15 |
| -1.1988 | 0.1232 | *1.2277* | 0.0507 | 1.2149 | 0.9489 | Q10570;E9PIM1 | CPSF1 |
| 0.1739 | 0.6293 | *-0.9010* | -0.0326 | 0.7858 | 0.9492 | Q8N4V2;H0YHK4 | SVOP |
| -0.3785 | -0.3469 | *0.8082* | 0.0276 | 0.6762 | 0.9501 | Q14324 | MYBPC2 |
| 0.2997 | -0.1642 | *-0.1680* | -0.0108 | 0.2689 | 0.9507 | O95573;C9JC11 | ACSL3 |
| 0.2194 | 0.1788 | *-0.4431* | -0.0150 | 0.3713 | 0.9507 | Q7Z2Z1;H0YN97 | TICRR |
| -0.4670 | -0.1269 | *0.6634* | 0.0232 | 0.5799 | 0.9511 | P29692;E9PK01;A0A087X1X7;E9PI39;E9PQ49;E9PQZ1;E9PL12;E9PL71;E9PPR1;E9PMW7;H0YCK7;E9PIZ1;E9PN91;H0YE72;E9PK06;H0YE58;E9PK72;E9PKK3;E9PQC9;E9PJD0;E9PNW6;E9PRY8 | EEF1D |
| -0.4549 | -0.5013 | *0.8643* | -0.0306 | 0.7754 | 0.9517 | A0A0G2JIR1;A0A0G2JIS2;A0A0G2JK64;A0A0G2JRN8;A0A0G2JRR0;A2ABF8;A2ABF9;Q96KQ7 | EHMT2 |
| -0.4991 | -0.1407 | *0.5754* | -0.0215 | 0.5471 | 0.9520 | P46776;E9PLL6;E9PLX7;E9PJD9;H0YF34;E9PKT2 | RPL27A |
| -0.1656 | -0.1341 | *0.3321* | 0.0108 | 0.2787 | 0.9526 | P49207 | RPL34 |
| -0.2945 | -0.1545 | *0.4059* | -0.0143 | 0.3706 | 0.9526 | P23246;H0Y9K7 | SFPQ |
| -0.6583 | -0.0573 | *0.7983* | 0.0276 | 0.7320 | 0.9539 | P60981;F6RFD5 | DSTN |
| -0.7927 | 0.3921 | *0.3264* | -0.0248 | 0.6659 | 0.9545 | Q13162;H7C3T4;A6NG45;A6NJJ0 | PRDX4 |
| 1.3431 | -2.9291 | *1.3148* | -0.0904 | 2.4584 | 0.9550 | B7Z637;Q8IUR7 | ARMC8 |
| 0.4098 | 0.2752 | *-0.7548* | -0.0232 | 0.6371 | 0.9554 | Q14151 | SAFB2 |
| -0.1657 | -0.5485 | *0.7892* | 0.0250 | 0.6890 | 0.9556 | Q99973;G3V5X7;G3V470;G3V2A4;G3V591;H0YJF6 | TEP1 |
| 1.0970 | -0.3880 | *-0.8169* | -0.0360 | 1.0043 | 0.9562 | E7END4 | LOXL3 |
| -0.1994 | -2.7365 | *3.2533* | 0.1058 | 3.0065 | 0.9569 | P51571;A6NLM8 | SSR4 |

| L100/ CTRL1 | L100/ CTRL2 | L100/ CTRL3 | Mean | SD | T-test p-value | Accession | Gene Symbol |
|---|---|---|---|---|---|---|---|
| -0.2399 | -0.3424 | *0.5321* | -0.0167 | 0.4780 | 0.9572 | Q8N163;H0YB24;G3V119;E5RHJ4;A0A087X2B6;E5RFJ3;E5RGU7;H0YC58;H0YC69 | CCAR2 |
| -0.5007 | 0.1933 | *0.3543* | 0.0157 | 0.4543 | 0.9578 | Q15942;H0Y2Y8;H7C3D3;C9IZ41 | ZYX |
| -0.4695 | -0.1125 | *0.5299* | -0.0174 | 0.5064 | 0.9580 | B7WPL9;F8W9A8;Q6PFW1;C9J5E6;C9JZX6 | PPIP5K1 |
| 0.2915 | -0.3541 | *0.0295* | -0.0111 | 0.3247 | 0.9583 | O00629;H7C4F6 | KPNA4 |
| -0.0740 | -0.0140 | *0.0802* | -0.0026 | 0.0777 | 0.9596 | O75874;C9J4N6;C9JLU6;C9JJE5 | IDH1 |
| 0.4242 | 1.6116 | *-2.2302* | -0.0648 | 1.9670 | 0.9597 | O76054 | SEC14L2 |
| 0.7201 | 0.4917 | *-1.1139* | 0.0326 | 0.9994 | 0.9601 | O43813;E9PHS0;F8WDS9;H7C2E3 | LANCL1 |
| -0.1096 | 0.6470 | *-0.4819* | 0.0185 | 0.5752 | 0.9606 | O60216 | RAD21 |
| 0.5104 | -2.3708 | *2.0755* | 0.0717 | 2.2554 | 0.9611 | Q9UL25 | RAB21 |
| -0.0922 | -0.5019 | *0.5453* | -0.0163 | 0.5277 | 0.9623 | A0A2R8YD50;E7EWE5;P51659;E7ER27;A0A2R8Y7L2;E7ET17;G5E9S2;A0A2R8YF39;E7EPL9 | HSD17B4 |
| 0.5366 | 0.2160 | *-0.6941* | 0.0195 | 0.6385 | 0.9626 | G5EA42;Q9NZR1;H0YKU1;H0YNJ8;Q9NYL9 | TMOD2 |
| 0.4424 | -0.7567 | *0.2555* | -0.0196 | 0.6451 | 0.9629 | A0A075B7G8;O00213;V9GY97 | APBB1 |
| -0.2701 | 2.8839 | *-2.8751* | -0.0871 | 2.8839 | 0.9630 | P49720;A0A087WUL2;A0A087WXQ8;A0A087WY10 | PSMB3 |
| -0.3630 | -0.4562 | *0.8867* | 0.0225 | 0.7499 | 0.9633 | Q9UMX0;H0YDS0 | UBQLN1 |
| -0.1014 | -0.0415 | *0.1549* | 0.0040 | 0.1341 | 0.9636 | A0A087WSW2;A0A0G2JQ90;P17038 | ZNF43 |
| 1.4964 | -0.7535 | *-0.8603* | -0.0392 | 1.3309 | 0.9640 | Q08AN1 | ZNF616 |
| -0.5756 | -0.0192 | *0.5466* | -0.0161 | 0.5611 | 0.9650 | P78344;D3DQV9;H0Y3P2;H0YCH5;H0YD77;H0YEC5;E9PKF8;H0YCF8;H0YD99;H0YDC0;H0YEN8 | EIF4G2 |
| 0.9492 | 0.4049 | *-1.2558* | 0.0328 | 1.1486 | 0.9651 | O15169;H0Y830 | AXIN1 |
| -1.0010 | 1.1281 | *-0.2186* | -0.0305 | 1.0769 | 0.9654 | Q13308 | PTK7 |
| 0.6709 | 3.6973 | *-4.7170* | -0.1163 | 4.2620 | 0.9666 | Q16587;F8WDT0 | ZNF74 |
| -0.6734 | 1.0814 | *-0.4867* | -0.0262 | 0.9638 | 0.9667 | Q8TBC5;M0R1U9;M0R364 | ZSCAN18 |
| 0.7967 | -0.6209 | *-0.1186* | 0.0191 | 0.7188 | 0.9675 | Q16623;A0A0C4DFZ1;A8MZ54 | STX1A |
| -0.5140 | 0.5315 | *-0.0579* | -0.0135 | 0.5242 | 0.9685 | O75323;H7C333;C9K068;F8WBI5 | NIPSNAP2 |

| L100/ CTRL1 | L100/ CTRL2 | L100/ CTRL3 | Mean | SD | T-test p-value | Accession | Gene Symbol |
|---|---|---|---|---|---|---|---|
| 0.5032 | 1.6113 | *-1.9755* | 0.0463 | 1.8365 | 0.9691 | Q68DQ2 | CRYBG3 |
| -0.1739 | -0.6025 | *0.8320* | 0.0185 | 0.7363 | 0.9692 | P12277;H0YJG0;G3V4N7;G3V461;H0YJK0;H0YJJ7 | CKB |
| 0.3404 | -0.9704 | *0.5693* | -0.0203 | 0.8308 | 0.9702 | O14646 | CHD1 |
| -1.9445 | 0.7762 | *1.0476* | -0.0402 | 1.6547 | 0.9702 | J3KQB2;Q9Y4R7;H0Y5E3 | TTLL3 |
| -0.3075 | -0.4993 | *0.7576* | -0.0164 | 0.6771 | 0.9704 | P62942;A0A087WZM5;A0A087WTS4;Q1JUQ3;Q5W0X3 | FKBP1A |
| 0.2718 | 0.8051 | *-1.0103* | 0.0222 | 0.9331 | 0.9709 | Q14160;A0A0G2JMS7;A0A0G2JNZ2;A0A0G2JPP5 | SCRIB |
| -0.4094 | -1.2026 | *1.7197* | 0.0359 | 1.5112 | 0.9709 | A6NHJ4 | ZNF860 |
| 0.1557 | -0.8882 | *0.7927* | 0.0201 | 0.8486 | 0.9711 | Q9P2E9;A0A087WVV2;F8W7S5;A0A087WU26 | RRBP1 |
| -0.4887 | 0.0077 | *0.5165* | 0.0119 | 0.5026 | 0.9711 | P51531;F6VDE0;A0A0U1RQZ9;A0A0U1RQX3;A0A0U1RR83;A0A0U1RRG6;A0A0A0MSS5;A0A0U1RQU0;A0A0U1RQW7;A0A0U1RR26;A0A0U1RRN2;A0A1W2PS06;B1ALF6;B1ALG2;B4DNT1;F6XE55;A0A0U1RR09;A0A0U1RRD6;A0A0U1RRF8;B1ALG1;F6T8Q0;F6UH26;A0A0U1RQE1 | SMARCA2 |
| -0.7792 | -2.0503 | *2.6588* | -0.0569 | 2.4362 | 0.9714 | O94819 | KBTBD11 |
| 0.4165 | -0.3350 | *-0.1083* | -0.0089 | 0.3855 | 0.9716 | A0A087WWF6;F8W8R3;P49005;C9IZD2;C9J8Z7;C9JLE1 | POLD2 |
| 0.7031 | -2.2177 | *1.6522* | 0.0459 | 2.0169 | 0.9722 | O95391 | SLU7 |
| -0.0341 | 1.3179 | *-1.1987* | 0.0284 | 1.2595 | 0.9724 | Q4AC99 | ACCSL |
| -0.1279 | -0.2306 | *0.3800* | 0.0072 | 0.3269 | 0.9732 | Q8IZT6;Q5VYL4 | ASPM |
| 0.2488 | 0.3505 | *-0.6334* | -0.0114 | 0.5411 | 0.9743 | Q16643;D6R9W4;D6R9Q9;D6RCR4;D6RFI1 | DBN1 |
| 0.2777 | -1.0700 | *0.7336* | -0.0196 | 0.9378 | 0.9745 | A0A2R8Y5P7;P49590;A0A2R8Y3N3;A0A2R8Y6I1;D6RJE6;A0A2R8Y5E8 | HARS2 |
| 0.9315 | 4.4351 | *-5.6730* | -0.1022 | 5.1327 | 0.9756 | Q8WVM8;G3V2M8;J3KNG4 | SCFD1 |
| -0.0933 | -0.3239 | *0.4407* | 0.0078 | 0.3922 | 0.9756 | Q6S8J3;A0A0A6YYL3;A0A0G2JMU2;A0JP26;B2R | POTEE |

| L100/ CTRL1 | L100/ CTRL2 | L100/ CTRL3 | Mean | SD | T-test p-value | Accession | Gene Symbol |
|---|---|---|---|---|---|---|---|
| | | | | | | U33;H3BUK9;A0A087WXQ7;A0A087X067;A0A087X092;A0A0C4DH93;A0A0D9SFE8;A0A0G2JQ08;A0A0G2JQB7;A0A0G2JRD4;A6NC16;D7UEQ8;Q495V5 | |
| 2.8991 | 1.8605 | *-5.0141* | -0.0848 | 4.3003 | 0.9758 | Q08AE8;J3KNG6 | SPIRE1 |
| -0.9344 | 0.6698 | *0.3144* | 0.0166 | 0.8425 | 0.9759 | Q9UJS0 | SLC25A13 |
| 0.0988 | 0.1525 | *-0.2390* | 0.0041 | 0.2123 | 0.9763 | Q15024 | EXOSC7 |
| -0.5177 | -0.8441 | *1.2969* | -0.0216 | 1.1535 | 0.9770 | C9IZY8;Q96PY5;F8W1F5;Q8IVF7 | FMNL2 |
| -0.1589 | -0.1820 | *0.3252* | -0.0053 | 0.2864 | 0.9775 | Q9NP71;H7C1V3 | MLXIPL |
| -2.8914 | 2.6934 | *0.3443* | 0.0488 | 2.8041 | 0.9787 | Q8NEZ2;E5RG91;E5RHB8;E5RJ10;E5RJX6 | VPS37A |
| 0.0654 | 1.1815 | *-1.3117* | -0.0216 | 1.2489 | 0.9788 | Q92805 | GOLGA1 |
| -0.0259 | -0.2131 | *0.2276* | -0.0038 | 0.2212 | 0.9790 | P30153;B3KQV6;C9J9C1;E9PH38;E9PHZ6;E9PPI5;E9PNM7 | PPP2R1A |
| -0.1563 | 0.4199 | *-0.2457* | 0.0060 | 0.3612 | 0.9798 | Q6V0I7 | FAT4 |
| -0.0526 | 0.3261 | *-0.2589* | 0.0049 | 0.2967 | 0.9799 | Q96JM2;H3BLX4;H0Y6H9 | ZNF462 |
| -0.4013 | 0.7688 | *-0.3353* | 0.0107 | 0.6573 | 0.9800 | A8MXL6;P55735;E7ERC8 | SEC13 |
| -0.3816 | 0.7412 | *-0.3292* | 0.0101 | 0.6337 | 0.9804 | Q86U42;B4DEH8;G3V4T2;H0YJH9 | PABPN1 |
| -0.4337 | 0.6058 | *-0.1971* | -0.0083 | 0.5448 | 0.9813 | O00567;H0Y653;H0YDU4 | NOP56 |
| -0.6206 | -1.1860 | *1.8789* | 0.0241 | 1.6310 | 0.9819 | O15067;J3QSG0;J3QSH6;J3KTL4;J3QL39;J3KTQ5;H0YGH1;J3KT98 | PFAS |
| -0.3287 | -0.6414 | *0.9347* | -0.0118 | 0.8345 | 0.9827 | P27348;E9PG15 | YWHAQ |
| -0.0501 | 2.2628 | *-2.3085* | -0.0319 | 2.2857 | 0.9829 | A2A3N6 | PIPSL |
| 1.6781 | -0.4793 | *-1.1405* | 0.0194 | 1.4740 | 0.9839 | P86790;P86791;F8WD66 | CCZ1B |
| 0.0990 | -0.3580 | *0.2717* | 0.0042 | 0.3254 | 0.9840 | P10155;G5E9R9;H0Y9N5;D6RDN1;D6RE09 | TROVE2 |
| -0.0422 | -0.8467 | *0.8560* | -0.0110 | 0.8518 | 0.9842 | Q93009;H3BND8;F5H2X1;H3BQD1;H3BMF6;H3BRA2;H3BTM1;H3BUV0 | USP7 |
| 0.5633 | 0.4286 | *-0.9594* | 0.0108 | 0.8429 | 0.9843 | Q9NZM3;A0A087WVF7;H7C3E2;H7BZD4 | ITSN2 |
| 0.8365 | -0.4895 | *-0.3749* | -0.0093 | 0.7347 | 0.9845 | Q9NR28;A0A024RBT2;A0A2U3TZ | DIABLO |

| L100/ CTRL1 | L100/ CTRL2 | L100/ CTRL3 | Mean | SD | T-test p-value | Accession | Gene Symbol |
|---|---|---|---|---|---|---|---|
| | | | | | | H2;H7BZK7;H7BZQ7 | |
| 0.7770 | 2.9235 | *-3.5777* | 0.0410 | 3.3125 | 0.9849 | Q1MSJ5 | CSPP1 |
| 0.0586 | 0.4759 | *-0.5169* | 0.0059 | 0.4985 | 0.9856 | Q86V48;E5RFK8;E5RHU7 | LUZP1 |
| -4.1602 | -1.8902 | *5.8727* | -0.0593 | 5.2611 | 0.9862 | Q9UEW8 | STK39 |
| -1.4784 | -1.6786 | *3.0695* | -0.0291 | 2.6854 | 0.9867 | Q8WWZ7;K7EPM3;K7EJW6;Q96M69 | ABCA5 |
| 0.1443 | 1.7444 | *-1.9490* | -0.0201 | 1.8522 | 0.9867 | O95861;A6NF51;F8VVW8;F8VRY7;F8W1J0 | BPNT1 |
| -0.4193 | 1.2309 | *-0.7782* | 0.0111 | 1.0715 | 0.9873 | Q96RU2;F5GZ74 | USP28 |
| 0.0906 | -0.1411 | *0.0543* | 0.0013 | 0.1246 | 0.9876 | Q8NC51 | SERBP1 |
| 1.3736 | -0.7777 | *-0.6312* | -0.0118 | 1.2020 | 0.9880 | Q9UBC2;M0R165;M0R2S2 | EPS15L1 |
| -0.6107 | -0.0392 | *0.6319* | -0.0060 | 0.6219 | 0.9882 | B2RTY4;H3BRD5;H3BV44;H3BMM1;H3BU05;H3BMS3;H3BTL9;H0YJE9;H0YJT1;H3BND9;H3BP49;H3BSU8 | MYO9A |
| -0.4344 | 0.6254 | *-0.1755* | 0.0052 | 0.5525 | 0.9886 | Q7LBC6 | KDM3B |
| 1.3464 | 1.5613 | *-2.9760* | -0.0228 | 2.5599 | 0.9891 | Q9NT99;M0R2G0 | LRRC4B |
| -0.6542 | 0.7548 | *-0.1192* | -0.0062 | 0.7112 | 0.9893 | Q15046;H3BVA8;J3KRL2;H3BPV7 | KARS |
| -0.0813 | -0.9216 | *0.9801* | -0.0076 | 0.9530 | 0.9902 | Q8IWG1 | WDR63 |
| -0.1498 | -1.1995 | *1.3198* | -0.0098 | 1.2655 | 0.9905 | Q5T5C0 | STXBP5 |
| 1.2110 | 1.7439 | *-2.9000* | 0.0183 | 2.5413 | 0.9912 | C9J4M0;Q9NVR7 | TBCCD1 |
| 0.9205 | -0.6148 | *-0.3230* | -0.0058 | 0.8153 | 0.9913 | P49750;H0YIQ2;F8VU51;H0YI23 | YLPM1 |
| 0.0972 | 3.4838 | *-3.5074* | 0.0245 | 3.4962 | 0.9914 | O75843;H0YJ08 | AP1G2 |
| 0.1450 | -0.5990 | *0.4457* | -0.0028 | 0.5378 | 0.9937 | P40227 | CCT6A |
| 1.1787 | -0.4877 | *-0.7065* | -0.0052 | 1.0311 | 0.9938 | H3BP42 | PYCARD |
| 0.3852 | -0.2401 | *-0.1412* | 0.0013 | 0.3361 | 0.9953 | Q96NX9;D6RFG7;D6REJ4;A0A087WTT5 | DACH2 |
| -1.4112 | 1.9660 | *-0.5746* | -0.0066 | 1.7588 | 0.9954 | Q9GZR1 | SENP6 |
| 0.2518 | 0.3229 | *-0.5800* | -0.0018 | 0.5020 | 0.9957 | A0A0A0MT58;Q5JRI7 | TCEA2 |
| 0.0093 | -1.4459 | *1.4214* | -0.0051 | 1.4337 | 0.9957 | O75970;F5H1U9;B7ZB24;H0YGQ3;H0YH70 | MPDZ |
| 0.3299 | -1.4424 | *1.1248* | 0.0041 | 1.3143 | 0.9962 | Q9UHF7;E5RJ97;H0YC29;C9J6L7;E5RFF3;E7EVN4;F8W8T0 | TRPS1 |
| 0.1635 | -0.7814 | *0.6126* | -0.0018 | 0.7116 | 0.9969 | O75436;S4R3Q6;S4R2Y3 | VPS26A |

| L100/ CTRL1 | L100/ CTRL2 | L100/ CTRL3 | Mean | SD | T-test p-value | Accession | Gene Symbol |
|---|---|---|---|---|---|---|---|
| -0.5037 | -0.0394 | *0.5470* | 0.0013 | 0.5266 | 0.9970 | Q9UQ80;F8VR77;F8W0A3;H0YIN7;F8VZ69 | PA2G4 |

**Table S2.** Gene symbols, protein names, accession numbers, fold changes, and p-values of DAPs. The upper section (red/white) corresponds to upregulated proteins, while the lower section (blue/white) refers to downregulated ones.

| Gene Symbol | Protein Name | Accession | $Log_2FC$ | p-value |
|---|---|---|---|---|
| ABRAXAS1 | Abraxas 1, BRCA1 A complex subunit | A0A087WXK1 | 4.6054 | 0.0350 |
| ACSBG2 | Acyl-CoA synthetase bubblegum family member 2 | Q5FVE4;K7EKE4;K7EL11;K7ERT0;K7ESC8;K7ESF1 | 1.6498 | 0.0364 |
| ACTBL2 | Actin beta like 2 | Q562R1 | 0.3981 | 0.0094 |
| AHCYL2 | Adenosylhomocysteinase like 2 | Q96HN2;H0Y8B3;C9K0S0 | 0.5938 | 0.0022 |
| ANAPC1 | Anaphase promoting complex subunit 1 | Q9H1A4;H0Y564;A0A2R8YF63 | 1.3163 | 0.0435 |
| ARPC2 | Actin related protein 2/3 complex subunit 2 | O15144;H7C3F9;G5E9J0 | 1.4045 | 0.0422 |
| BCOR | BCL6 corepressor | Q6W2J9;H7BYY2 | 0.2489 | 0.0197 |
| BTF3L4 | Basic transcription factor 3 like 4 | Q96K17;E9PL10 | 1.3351 | 0.0133 |
| CCDC40 | Coiled-coil domain containing 40 | Q4G0X9;I3L477 | 0.8677 | 0.0378 |
| CCDC8 | Coiled-coil domain containing 8 | Q9H0W5 | 0.7107 | 0.0101 |
| CHCHD2P9 | Coiled-coil-helix-coiled-coil-helix domain containing 2 pseudogene 9 | Q5T1J5;Q9Y6H1 | 1.0845 | 0.0134 |
| CLTCL1 | Clathrin heavy chain like 1 | P53675;A0A087WX41;F5H5N6;A0A087WXH4;H0YGJ9 | 1.2363 | 0.0394 |
| CPT2 | Carnitine palmitoyltransferase 2 | P23786;A0A1B0GTB8;A0A1B0GV75;A0A1B0GVF3;A0A1B0GWC0 | 1.1683 | 0.0406 |

| Gene Symbol | Protein Name | Accession | $Log_2FC$ | p-value |
|---|---|---|---|---|
| CTBP2 | C-terminal binding protein 2 | P56545;Q5SQP8 | 1.1785 | 0.0273 |
| DCAF7 | DDB1 and CUL4 associated factor 7 | A0A087WWI6;P61962 | 0.4101 | 0.0107 |
| DDIAS | DNA damage induced apoptosis suppressor | Q8IXT1;E9PMA7;E9PN94 | 1.0579 | 0.0220 |
| DDX39B | DExD-box helicase 39B | Q13838;F6WLT2;A0A0A0MT12;F6TRA5;F6S4E6;A0A140T9X9;A0A140TA18;F6UJC5;H0Y400;H0YCC6;A0A0G2JJL7;F6QYI9;F6R6M7;A0A0G2JHN7;F6S2B7;F6U6E2;A0A140T9N3;F6SXL5 | 0.9424 | 0.0086 |
| DDX47 | DEAD-box helicase 47 | Q9H0S4;F5H1N9 | 1.6669 | 0.0099 |
| DNAH5 | Dynein axonemal heavy chain 5 | Q8TE73 | 0.8183 | 0.0455 |
| DSE | Dermatan sulfate epimerase | Q9UL01;A0A2R8Y6J1;A0A2U3TZJ0 | 0.7275 | 0.0238 |
| DZANK1 | Double zinc ribbon and ankyrin repeat domains 1 | Q9NVP4 | 0.8208 | 0.0280 |
| EIF1AX | Eukaryotic translation initiation factor 1A X-linked | P47813;O14602;X6RAC9;A6NJH9 | 2.1173 | 0.0061 |
| EIF3M | Eukaryotic translation initiation factor 3 subunit M | Q7L2H7;J3KNJ2 | 0.5823 | 0.0232 |
| EIF5A2 | Eukaryotic translation initiation factor 5A2 | Q9GZV4;C9J7B5;F8WCJ1 | 2.6496 | 0.0024 |

| Gene Symbol | Protein Name | Accession | $Log_2FC$ | p-value |
|---|---|---|---|---|
| ELMSAN1 | Mitotic deacetylase associated SANT domain protein | A0A1C7CYX1;Q6PJG2;A0A0A0MSU2 | 0.1531 | 0.0146 |
| FBLN1 | Fibulin 1 | P23142;B1AHM7;B1AHM9;H7C1M6;B1AHM6;B1AHM8;B1AHN3 | 1.4360 | 0.0109 |
| GAPDHS | Glyceraldehyde-3-phosphate dehydrogenase, spermatogenic | K7EP73;O14556 | 0.4501 | 0.0158 |
| GGH | Gamma-glutamyl hydrolase | Q92820 | 1.4239 | 0.0405 |
| GOLGA6L2 | Golgin A6 family like 2 | Q8N9W4;F8WBV9 | 0.1671 | 0.0360 |
| GPS1 | G protein pathway suppressor 1 | Q13098;A0A096LP07;A0A096LPJ3;A8K070;C9JFE4;J3KRJ4;J3KSA5;J3KTB0;J3QLT0;J3QS88;J3KRE8;J3QL53;J3QLE8;J3QQX0;J3QS84 | 1.1481 | 0.0244 |
| GSK3A | Glycogen synthase kinase 3 alpha | P49840;A8MT37;M0QYV0 | 0.3970 | 0.0388 |
| HMGN3 | High mobility group nucleosomal binding domain 3 | A0A087WZE9;Q15651 | 0.5828 | 0.0185 |
| HSPBP1 | HSPA (Hsp70) binding protein 1 | Q9NZL4 | 0.5842 | 0.0389 |
| LHX6 | LIM homeobox 6 | Q9UPM6;H0YMY8 | 0.9622 | 0.0382 |
| LRRC36 | Leucine rich repeat containing 36 | Q1X8D7;J3QSG3;H3BNV0;H3BQI9;H3BRB2;H3BRP7;H3BSQ6 | 0.2797 | 0.0195 |
| LRRC37B | Leucine rich repeat containing 37B | F5H5K1;J3KTP0;J3QSU1;Q96QE4 | 1.0703 | 0.0259 |
| LRRK2 | Leucine rich repeat kinase 2 | Q5S007;E9PC85 | 0.3554 | 0.0259 |
| MARCH7 | Membrane associated ring-CH-type finger 7 | Q9H992 | 1.5892 | 0.0301 |

| Gene Symbol | Protein Name | Accession | $Log_2FC$ | p-value |
|---|---|---|---|---|
| MTDH | Metadherin | Q86UE4;E5RJU9;H0YBE0 | 1.0757 | 0.0170 |
| MYO1F | Myosin IF | O00160 | 2.5042 | 0.0490 |
| NLRP2 | NLR family pyrin domain containing 2 | A0A0G2JLQ8;A0A0G2JMG8;A0A0G2JNC8;A0A0G2JPQ2;J3KN39;Q9NX02;A0A0G2JPB6;A0A0G2JLX3;A0A0G2JP37;K7EMK2 | 1.4664 | 0.0473 |
| NUP93 | Nucleoporin 93 | Q8N1F7;H3BVG0;H3BV15;H3BM93;H3BP95;H3BRI8;H3BMX0;H3BNG7;H3BNN5;H3BPA9 | 0.4813 | 0.0152 |
| PABPC1L2A | Poly(A) binding protein cytoplasmic 1 like 2A | Q5JQF8 | 2.0866 | 0.0027 |
| PDE6B | Phosphodiesterase 6B | P35913;H7C4P9 | 0.6271 | 0.0293 |
| PDP1 | Pyruvate dehydrogenase phosphatase catalytic subunit 1 | Q9P0J1;E5RI96;E5RIE5;E5RIV4 | 2.5390 | 0.0221 |
| PKLR | Pyruvate kinase L/R | P30613 | 0.3700 | 0.0337 |
| POLRMT | RNA polymerase mitochondrial | O00411;K7EMH3 | 3.3261 | 0.0296 |
| PRAG1 | PEAK1 related, kinase-activating pseudokinase 1 | Q86YV5 | 1.3650 | 0.0001 |
| PRKDC | Protein kinase, DNA-activated, catalytic subunit | P78527;H0YG84 | 0.1972 | 0.0358 |
| PRRC2C | Proline rich coiled-coil 2C | Q9Y520;E7EPN9;A0A0A0MS30 | 0.4678 | 0.0349 |
| PTPN11 | Protein tyrosine phosphatase non-receptor type 11 | Q06124;A0A1W2PPU4;A0A0U1RRI0 | 1.4603 | 0.0444 |
| PXK | PX domain containing serine/threonine kinase like | Q7Z7A4;W5RWE6 | 0.8144 | 0.0452 |

| Gene Symbol | Protein Name | Accession | $Log_2FC$ | p-value |
|---|---|---|---|---|
| RAB3A | RAB3A, member RAS oncogene family | P20336;S4R3Q3 | 1.4259 | 0.0390 |
| RAB8B | RAB8B, member RAS oncogene family | Q92930;H0YNE9 | 0.3254 | 0.0384 |
| RBBP7 | RB binding protein 7, chromatin remodeling factor | Q16576;E9PC52;Q5JP02;Q5JNZ6;Q5JP01 | 1.3048 | 0.0387 |
| RPL35A | Ribosomal protein L35a | C9K025;P18077;F8WB72;F8WBS5 | 1.0879 | 0.0093 |
| SASH1 | SAM and SH3 domain containing 1 | O94885 | 1.7097 | 0.0168 |
| SCN4A | Sodium voltage-gated channel alpha subunit 4 | P35499 | 1.4184 | 0.0318 |
| SMARCC1 | SWI/SNF related, matrix associated, actin dependent regulator of chromatin subfamily c member 1 | Q92922 | 1.0876 | 0.0382 |
| SNAPC3 | Small nuclear RNA activating complex polypeptide 3 | Q5T282;Q92966 | 1.3180 | 0.0334 |
| STK10 | Serine/threonine kinase 10 | O94804 | 1.0190 | 0.0241 |
| SYT1 | Synaptotagmin 1 | P21579;J3KQA0;C9JX50;F8VXH0;F8VYH8;F8VZY3;F8W1U9 | 4.4408 | 0.0128 |
| TACC2 | Transforming acidic coiled-coil containing protein 2 | O95359;E7EMZ9;E9PBC6;D6RAA5;Q4VXL4;Q4VXL8;H0Y9Y7 | 0.8253 | 0.0160 |
| TAF1C | TATA-box binding protein associated factor, RNA polymerase I subunit C | Q15572 | 0.8216 | 0.0090 |
| TCF25 | Transcription factor 25 | Q9BQ70;H3BMJ8;H3BSP8;H3BTU3;Q9H7D3 | 1.0545 | 0.0369 |

| Gene Symbol | Protein Name | Accession | $Log_2FC$ | p-value |
|---|---|---|---|---|
| TMEM43 | Transmembrane protein 43 | Q9BTV4 | 0.6323 | 0.0171 |
| TOR1AIP1 | Torsin 1A interacting protein 1 | A0A0A0MSK5;J3KN66;Q5JTV8;H0Y4R4 | 1.1019 | 0.0213 |
| TPM4 | Tropomyosin 4 | P67936;A0A2R8YGX3;A0A2R8YE05;K7EP68;A0A2R8YHD2;K7EMU5;K7EPV9 | 1.5643 | 0.0160 |
| TTC7B | Tetratricopeptide repeat domain 7B | A0A0C4DGK5;H0YJS2;H0YK02;Q86TV6 | 2.5809 | 0.0011 |
| USP8 | Ubiquitin specific peptidase 8 | P40818 | 1.3639 | 0.0477 |
| VPS13B | Vacuolar protein sorting 13 homolog B | Q7Z7G8 | 0.9593 | 0.0021 |
| VPS4A | Vacuolar protein sorting 4 homolog A | Q9UN37;I3L4J1;O75351 | 0.3350 | 0.0465 |
| VRK3 | VRK serine/threonine kinase 3 | M0QYA8;Q8IV63;M0R073;M0QXD7;M0QYG0;M0QZ79;M0R025 | 0.8696 | 0.0042 |
| ZCCHC8 | Zinc finger CCHC-type containing 8 | Q6NZY4 | 0.3831 | 0.0104 |
| ZFPM1 | Zinc finger protein, FOG family member 1 | Q8IX07;A0A087WWQ0;A0A087WZP1 | 0.5219 | 0.0031 |
| ZNF136 | Zinc finger protein 136 | P52737;C9J2Q9;C9JJK8 | 1.4529 | 0.0189 |
| ZNF302 | Zinc finger protein 302 | E7EVR1;Q9NR11 | 2.3123 | 0.0119 |
| ZNF395 | Zinc finger protein 395 | E5RG59 | 0.9953 | 0.0127 |
| ZNF766 | Zinc finger protein 766 | G3XAE0;Q5HY98 | 2.1607 | 0.0403 |

| | | | | |
|---|---|---|---|---|
| ZNF836 | Zinc finger protein 836 | Q6ZNA1;A0A0A0MR57;A0A075B7G2;A0A075B7G3;A0A087WUU8;A0A087WV98;A0A087WWI3;A0A087X254;A0A087X2A5;A0A087X2B0;A0A0C4DGP9;A0A0U1RQK1;A0A1W2PNY2;A0A1W2PQL4;A0A1W2PRC0;A2RRD8;A6NP11;A8MTY0;A8MUV8;B1APK8;B4DU55;B4DX44;B9EG95;C9K0H3;E7EWC5;F5H032;F5H290;F8W6Y9;F8W889;H0Y892;H3BS42;H9KV89;K7EK80;K7EP55;M0QX96;M0QXU9;M0QY24;M0QYS4;M0R0F3;M0R1M8;O14709;O43309;O43345;O43361;O95600;O95780;P0CB33;P0DPD5;P10073;P17019;P17022;P17035;P17097;P51522;P51815;P52744;Q03924;Q09FC8;Q15929;Q15937;Q2M3W8;Q2M3X9;Q2VY69;Q3ZCX4;Q5JNZ3;Q5SXM1;Q5VIY5;Q6AZW8;Q6P9G9;Q6PDB4;Q6ZMW2;Q6ZN06;Q6ZN19;Q6ZN57;Q76KX8;Q7Z3V5;Q86TJ5;Q86UE3;Q86V71;Q86XN6;Q86Y25;Q8IW36;Q8N8C0;Q8N8J6;Q8N972;Q8N988;Q8N9F8;Q8N9M3;Q8NDQ6;Q8NEM1;Q8NHY6;Q8TB69;Q8TBZ5;Q8TF39;Q8WXB4;Q96CX3;Q96IR2;Q96LX8;Q96MR9;Q96PE6;Q99676;Q9H5H4;Q9H7R5;Q9H963;Q9HBT7;Q9HC | 0.5851 | 0.0410 |

| Gene Symbol | Protein Name | Accession | $Log_2FC$ | p-value |
|---|---|---|---|---|
| | | L3;Q9NQX6;Q9P0L1;Q9Y473 | | |
| ZSCAN9 | Zinc finger and SCAN domain containing 9 | A0A0B4J224;E9PLJ4;E9PQL7;O15535;U3KQB2;U3KQV4 | 0.4191 | 0.0301 |

| Gene Symbol | Protein Name | Accession | $Log_2FC$ | p-value |
|---|---|---|---|---|
| ABCA8 | ATP binding cassette subfamily A member 8 | A0A0A0MSU4;O94911 | -0.9918 | 0.0214 |
| ACTN1 | Actinin alpha 1 | P12814;H9KV75;G3V2W4;G3V2N5;H7C5W8;H0YJW3;G3V2X9;H0YJ11;G3V5M4;G3V2E8 | -0.5430 | 0.0492 |
| ACTR3 | Actin related protein 3 | P61158;B4DXW1;F8WDR7;F8WE84;F8WEW2 | -0.3952 | 0.0122 |
| AHI1 | Abelson helper integration site 1 | Q8N157 | -0.5476 | 0.0238 |
| AKAP13 | A-kinase anchoring protein 13 | Q12802;H0YMW2;A0A087WTD7 | -1.3788 | 0.0046 |
| ALDOB | Aldolase, fructose-bisphosphate B | P05062;A0A087WXX2 | -0.3893 | 0.0090 |
| AMOTL1 | Angiomotin like 1 | Q8IY63 | -0.9831 | 0.0200 |
| AP2B1 | Adaptor related protein complex 2 subunit beta 1 | P63010;A0A087X253;A0A087WU93;A0A087WZQ6;A0A087WYD1;K7ERB2;K7EN71;A0A087WXS3;K7EJX1;K7EKZ5 | -1.2089 | 0.0452 |
| ARHGAP21 | Rho GTPase activating protein 21 | Q5T5U3;A0A1B0GV73;E7ESW5;F8W9U9 | -0.6036 | 0.0264 |
| ARHGAP5 | Rho GTPase activating protein 5 | A0A0A0MSK6;Q13017;G3V444;G3V5I7 | -0.8517 | 0.0110 |
| ARMC3 | Armadillo repeat containing 3 | Q5W041 | -0.7417 | 0.0173 |

| Gene Symbol | Protein Name | Accession | Log2FC | p-value |
|---|---|---|---|---|
| ASAH1 | N-acylsphingosine amidohydrolase 1 | Q13510;A0A1B0GTM3;A0A1B0GUA4;A0A1B0GUH5;A0A1B0GW68;E7EMM4;A0A1B0GTP7;A0A1B0GTZ5;A0A1B0GUG1;A0A1B0GUE3;A0A1B0GUW4;A0A1B0GV06;A0A1B0GUB3;A0A1B0GU62;A0A1B0GV88;A0A1B0GVC9;A0A1B0GVG2;A0A1B0GVJ1;A0A1B0GW48;A0A1B0GTD4;A0A1B0GTQ7;A0A1B0GU06;A0A1B0GV95;A0A1B0GVE7;Q86WS4 | -0.5079 | 0.0345 |
| ATP5PB | ATP synthase peripheral stalk-membrane subunit b | P24539;Q5QNZ2 | -0.5838 | 0.0346 |
| BOLA2 | BolA family member 2 | Q9H3K6;A0A087WZT3;A0A0B4J295;H3BTW0;H3BV85 | -1.0132 | 0.0354 |
| BRAF | B-Raf proto-oncogene, serine/threonine kinase | A0A2R8Y8E0;A0A2U3TZI2;H7C560;P15056;A0A2R8Y467;A0A2R8YDP5 | -3.6956 | 0.0321 |
| C1QBP | Complement C1q binding protein | Q07021;I3L3B0;I3L3Q7 | -0.5220 | 0.0442 |
| CACNA1I | Calcium voltage-gated channel subunit alpha1 I | Q9P0X4 | -0.4818 | 0.0292 |
| CDC42 | Cell division cycle 42 | P60953;Q5JYX0 | -1.2715 | 0.0385 |
| CEP76 | Centrosomal protein 76 | Q8TAP6 | -1.4186 | 0.0381 |
| CLU | Clusterin | P10909;H0YC35;H0YAS8;E5RJZ5;E7ERK6;E7ETB4;H0YLK8 | -0.2220 | 0.0361 |
| COL14A1 | Collagen type XIV alpha 1 chain | Q05707;J3QT83;Q4G0W3 | -0.3323 | 0.0469 |
| COPB2 | COPI coat complex subunit beta 2 | P35606;D6R997;D6RBT6;D6RBG7;D6RBZ7;D6RCL6;H0YAC7 | -0.3609 | 0.0471 |

| Gene Symbol | Protein Name | Accession | $Log_2FC$ | p-value |
|---|---|---|---|---|
| COPS8 | COP9 signalosome subunit 8 | Q99627;E9PGT6;H7C3S9 | -0.3532 | 0.0135 |
| CPNE1 | Copine 1 | Q99829;B0QZ18;A6PVH9;F2Z2V0;E7ENH5;Q5JX45;Q5JX44;Q5JX56;Q5JX58;Q5JX59;Q5JX60;H0Y524;Q5JX52;Q5JX61;Q5JX55;E7EV27;Q5JX53;Q5JX57;Q5JX54 | -0.3548 | 0.0429 |
| CRABP1 | Cellular retinoic acid binding protein 1 | P29762;B5MCB5 | -0.5873 | 0.0240 |
| CRMP1 | Collapsin response mediator protein 1 | Q14194;E9PD68 | -0.9273 | 0.0041 |
| CROCC | Ciliary rootlet coiled-coil, rootletin | Q5TZA2;B1AKD8;A0A087WW81;A0A087WU09;Q86T23;B0QYQ9;B0QYR0;B0QYR1;H0YKS0;Q8IVE0;Q9Y2F9 | -0.3505 | 0.0336 |
| CTAG1A | Cancer/testis antigen 1A | A0A0A0MTT5 | -1.6122 | 0.0072 |
| CYC1 | Cytochrome c1 | P08574 | -0.9763 | 0.0196 |
| DAD1 | Defender against cell death 1 | A0A0B4J239;F5GXX5;F5H895;P61803 | -1.7048 | 0.0168 |
| DAPK1 | Death associated protein kinase 1 | P53355;F8WCQ3 | -0.5697 | 0.0217 |
| DLC1 | DLC1 Rho GTPase activating protein | Q96QB1;A0A0J9YWS8;R4GMP5 | -0.8319 | 0.0296 |
| DNAH3 | Dynein axonemal heavy chain 3 | Q8TD57;Q5T7N2 | -0.3587 | 0.0020 |
| DUSP3 | Dual specificity phosphatase 3 | B5BUI8;P51452 | -0.8605 | 0.0260 |
| DYNLL1 | Dynein light chain LC8-type 1 | P63167;F8VRV5;F8VXI7;F8VXL2 | -0.8801 | 0.0284 |
| DYNLL2 | Dynein light chain LC8-type 2 | Q96FJ2 | -0.7299 | 0.0298 |

| Gene Symbol | Protein Name | Accession | $Log_2FC$ | p-value |
|---|---|---|---|---|
| DYNLRB2 | Dynein light chain roadblock-type 2 | Q8TF09;H3BQI1;Q9NP97;B1AKR6;H3BNG9;H3BPA0;Q7Z4M1 | -0.6107 | 0.0391 |
| EEA1 | Early endosome antigen 1 | Q15075 | -0.6477 | 0.0482 |
| EEF1A1 | Eukaryotic translation elongation factor 1 alpha 1 | A0A087WVQ9 | -0.3953 | 0.0384 |
| EEF1A1P5 | Eukaryotic translation elongation factor 1 alpha 1 pseudogene 5 | Q5VTE0;P68104;A0A087WV01;Q5JR01;A6PW80;A0A0A0MST8;C9JRP1;E9PFN4;H0YBD9;H0YC27;H0YC67;H7C3C4;P12757;Q9Y6M7 | -0.2824 | 0.0101 |
| EFTUD2 | Elongation factor Tu GTP binding domain containing 2 | Q15029;K7EP67;K7EJ74;K7EIT3 | -0.6847 | 0.0076 |
| EIF4A2 | Eukaryotic translation initiation factor 4A2 | Q14240;E7EQG2;J3KSN7;E7EMV8;E9PBH4;F8WE11;J3KS93;J3KT04;I3L3H2;J3QQP0 | -0.2525 | 0.0446 |
| EIF5 | Eukaryotic translation initiation factor 5 | P55010;H0YLZ1;H0YN40;H0YM54;H0YMJ8 | -0.6126 | 0.0418 |
| FABP7 | Fatty acid binding protein 7 | O15540 | -0.5000 | 0.0209 |
| FADS1 | Fatty acid desaturase 1 | A0A0A0MR51;O60427 | -0.3520 | 0.0367 |
| FAM49A | CYFIP related Rac1 interactor A | Q9H0Q0;C9IYV6;C9JPE5 | -0.2483 | 0.0059 |
| FHAD1 | Forkhead associated phosphopeptide binding domain 1 | B1AJZ9;H0Y3C6;H0YEU3;Q5JYW1 | -0.6985 | 0.0102 |
| FIP1L1 | Factor interacting with PAPOLA and CPSF1 | Q6UN15;H0Y8P7 | -2.5224 | 0.0154 |
| FSCN1 | Fascin actin-bundling protein 1 | A0A0A0MSB2;C9JFC0;C9JPH9 | -0.8975 | 0.0380 |

| Gene Symbol | Protein Name | Accession | $Log_2FC$ | p-value |
|---|---|---|---|---|
| FTL | Ferritin light chain | P02792 | -0.5435 | 0.0183 |
| GABRA6 | Gamma-aminobutyric acid type A receptor subunit alpha6 | Q16445 | -1.2321 | 0.0388 |
| GNB4 | G protein subunit beta 4 | Q9HAV0;C9JD14;H7C5J5 | -0.5247 | 0.0034 |
| GOPC | Golgi associated PDZ and coiled-coil motif containing | F5H1Y4;Q9HD26;A0A0J9YVX5 | -1.7995 | 0.0383 |
| GPHN | Gephyrin | F5H039;Q9NQX3;G3V582;H0YJR5;H0YJ30 | -0.9649 | 0.0209 |
| HDAC6 | Histone deacetylase 6 | Q9UBN7;A0A2R8YDE6;A0A2R8Y4G2 | -0.2567 | 0.0198 |
| HEATR4 | HEAT repeat containing 4 | Q86WZ0 | -0.4232 | 0.0338 |
| HIST1H1E | H1.4 linker histone, cluster member | P10412 | -2.7870 | 0.0415 |
| HMGCS1 | 3-hydroxy-3-methylglutaryl-CoA synthase 1 | Q01581;D6RIW1 | -0.3943 | 0.0088 |
| HNRNPC | Heterogeneous nuclear ribonucleoprotein C | B2R5W2;B4DY08;G3V4C1;G3V4W0;G3V2Q1;G3V576;G3V555;G3V575;G3V251;B4DSU6;G3V3K6;G3V5X6;G3V2D6;A0A0G2JPF8;B7ZW38;G3V4M8;O60812;P0DMR1;G3V2H6;G3V5V7 | -1.0760 | 0.0355 |
| HSD17B2 | Hydroxysteroid 17-beta dehydrogenase 2 | P37059 | -1.2452 | 0.0451 |
| IDH3A | Isocitrate dehydrogenase (NAD(+)) 3 catalytic subunit alpha | P50213;H0YL72;H0YKD0;H0YLI6;H0YMU3;H0YM64;H0YNF5;H0YNF8;H0YM46 | -0.9857 | 0.0397 |
| IQCH | IQ motif containing H | Q86VS3;H3BU17 | -0.6093 | 0.0457 |
| KAT14 | Lysine acetyltransferase 14 | A0A075B6H4;Q9H8E8 | -1.8891 | 0.0106 |

| Gene Symbol | Protein Name | Accession | $Log_2FC$ | p-value |
|---|---|---|---|---|
| KCNJ16 | Potassium inwardly rectifying channel subfamily J member 16 | K7EJR9;K7EPW9;Q9NPI9;K7EKJ4;K7ELL5 | -0.3640 | 0.0466 |
| KHDRBS1 | KH RNA binding domain containing, signal transduction associated 1 | Q07666 | -0.6460 | 0.0097 |
| KIF20B | Kinesin family member 20B | Q96Q89;A0A0A0MSJ5 | -0.8020 | 0.0270 |
| KTN1 | Kinectin 1 | Q86UP2;G3V4Y7;B7Z6P3;G3V5G2;H0YJV5;G3V5P0 | -1.4776 | 0.0125 |
| LCP1 | Lymphocyte cytosolic protein 1 | P13796;Q5TBN3 | -0.2381 | 0.0474 |
| LRP2 | LDL receptor related protein 2 | P98164 | -1.2390 | 0.0054 |
| LRP5 | LDL receptor related protein 5 | O75197;E9PHY1 | -0.4471 | 0.0113 |
| LRRIQ1 | Leucine rich repeats and IQ motif containing 1 | Q96JM4;H0YCJ9 | -0.4138 | 0.0397 |
| LSS | Lanosterol synthase | P48449;A0A0G2JQD0;C9J315;A0A0G2JS81 | -0.3618 | 0.0356 |
| MAPK8 | Mitogen-activated protein kinase 8 | A1L4K2;B5BUB8;P45983;A6NF29;C9J762 | -0.1981 | 0.0218 |
| MARCKS | Myristoylated alanine rich protein kinase C substrate | P29966 | -0.2752 | 0.0003 |
| MAX | MYC associated factor X | P61244;G3V5L1 | -2.2387 | 0.0141 |
| MDM1 | Mdm1 nuclear protein | Q8TC05 | -0.5850 | 0.0345 |
| MRPS36 | Alpha-ketoglutarate dehydrogenase subunit 4 | P82909 | -0.4527 | 0.0277 |

| Gene Symbol | Protein Name | Accession | $Log_2FC$ | p-value |
|---|---|---|---|---|
| MSH2 | MutS homolog 2 | P43246;V9H019;A0A2R8Y7S8;V9H0B2;V9H015;C9J809;A0A2R8Y713 | -1.1631 | 0.0138 |
| MTCL1 | Microtubule crosslinking factor 1 | Q9Y4B5;J3QLE1 | -0.7737 | 0.0001 |
| MTMR10 | Myotubularin related protein 10 | Q9NXD2 | -0.3480 | 0.0459 |
| MYL12B | Myosin light chain 12B | O14950;J3QRS3;P19105 | -0.6760 | 0.0272 |
| NAPA | NSF attachment protein alpha | P54920;M0R0Y2;M0R2M1;M0R027;M0R0I4;M0R213;M0R058 | -0.3963 | 0.0271 |
| NDUFS2 | NADH:ubiquinone oxidoreductase core subunit S2 | O75306 | -0.4088 | 0.0448 |
| NF1 | Neurofibromin 1 | P21359;J3KSB5;H0Y465 | -0.4917 | 0.0089 |
| NPLOC4 | NPL4 homolog, ubiquitin recognition factor | Q8TAT6 | -1.1865 | 0.0390 |
| NUDT3 | Nudix hydrolase 3 | O95989 | -0.6734 | 0.0076 |
| OGA | O-GlcNAcase | O60502;H7C3X0 | -0.7672 | 0.0104 |
| PABPC4 | Poly(A) binding protein cytoplasmic 4 | Q13310;B1ANR0;H0Y5F5;B1ANR1;H0YCC8;H0YEQ8;H0YEU6 | -2.2467 | 0.0151 |
| PAFAH1B2 | Platelet activating factor acetylhydrolase 1b catalytic subunit 2 | P68402;J3KNE3 | -0.5167 | 0.0250 |
| PAK2 | p21 (RAC1) activated kinase 2 | Q13177;H7C1X3 | -0.5593 | 0.0339 |
| PFDN4 | Prefoldin subunit 4 | E9PQY2;Q9NQP4 | -0.9549 | 0.0405 |
| PFKFB3 | 6-phosphofructo-2-kinase/fructose-2,6-biphosphatase 3 | Q16875;A0A1W2PR17;Q5VX20;Q5W015;F2Z2I2;A0A1W2PNV9;H0Y483;I1Z9G3;Q16877;Q4VBA9;Q66S35 | -0.5417 | 0.0361 |

| Gene Symbol | Protein Name | Accession | $Log_2FC$ | p-value |
|---|---|---|---|---|
| PFKM | Phosphofructokinase, muscle | P08237;A0A2R8Y891;F8VNX2;F8VZQ1;F8VP00;F8VX13;F8VSL1;F8VW30 | -0.4705 | 0.0269 |
| PGD | Phosphogluconate dehydrogenase | P52209;K7ELN9;K7EM49;K7EMN2 | -0.4835 | 0.0037 |
| PHF8 | PHD finger protein 8 | Q9UPP1;H0Y3N9;H0Y589;B0QZE1;B0QZZ2;B0QZZ3;B0QZZ4;Q5JPR8 | -1.4339 | 0.0001 |
| PHPT1 | Phosphohistidine phosphatase 1 | Q9NRX4 | -2.6759 | 0.0201 |
| PKP4 | Plakophilin 4 | A0A0D9SF60;Q99569;E7EST6;E7EMY7;E7EP40;E9PHJ1 | -1.9383 | 0.0328 |
| PLIN3 | Perilipin 3 | O60664;K7ERZ3;K7EL96;K7ER39 | -1.1367 | 0.0429 |
| PMVK | Phosphomevalonate kinase | Q15126 | -0.5136 | 0.0159 |
| PNMT | Phenylethanolamine N-methyltransferase | A8MT87;P11086 | -1.1874 | 0.0152 |
| POLE | DNA polymerase epsilon, catalytic subunit | Q07864;F5H1D6;F5H7E4 | -0.3267 | 0.0254 |
| PPA1 | Inorganic pyrophosphatase 1 | Q15181;Q5SQT6 | -0.7851 | 0.0208 |
| PPFIA4 | PTPRF interacting protein alpha 4 | O75335;B1APN9;M0QZB5 | -0.4364 | 0.0480 |
| PRPS1L1 | Phosphoribosyl pyrophosphate synthetase 1 like 1 | A0A0B4J207;P21108 | -0.2594 | 0.0069 |
| PSMC2 | Proteasome 26S subunit, ATPase 2 | P35998;A0A1W2PQS1;C9JLS9 | -0.3735 | 0.0128 |
| PSMD10 | Proteasome 26S subunit, non-ATPase 10 | B1AJY5;B1AJY7;O75832 | -0.6062 | 0.0371 |
| PTN | Pleiotrophin | P21246;C9JR52 | -0.6357 | 0.0447 |

| Gene Symbol | Protein Name | Accession | $Log_2FC$ | p-value |
|---|---|---|---|---|
| PXN | Paxillin | A0A1B0GTU4;F5GZ78;P49023;A0A1B0GU60;A0A1B0GWE7;A0A1B0GV30 | -0.7995 | 0.0316 |
| RAB11FIP4 | RAB11 family interacting protein 4 | Q86YS3;K7EL58 | -1.3441 | 0.0015 |
| RAB5C | RAB5C, member RAS oncogene family | P51148;F8VVK3;K7ERI8;K7ERQ8;K7ENY4;F8VVZ0;K7EIP6 | -2.3936 | 0.0300 |
| RABGAP1 | RAB GTPase activating protein 1 | Q9Y3P9;B5MCD9;C9JGR5 | -1.7708 | 0.0002 |
| RBBP4 | RB binding protein 4, chromatin remodeling factor | Q09028;H0YF10;H0YCT5;C9JPP3 | -0.5560 | 0.0296 |
| RCN3 | Reticulocalbin 3 | Q96D15;M0QZH0 | -0.7584 | 0.0251 |
| RNF20 | Ring finger protein 20 | Q5VTR2;C9J0A5;C9JXC9 | -0.5355 | 0.0251 |
| RPL12 | Ribosomal protein L12 | P30050 | -0.5821 | 0.0311 |
| RPL9 | Ribosomal protein L9 | P32969;A0A2R8Y5Y7;D6RAN4;E7ESE0;H0Y9V9;H0Y9R4 | -0.6588 | 0.0225 |
| RSPH6A | Radial spoke head 6 homolog A | M0R2K1;Q9H0K4 | -1.3780 | 0.0231 |
| RUFY2 | RUN and FYVE domain containing 2 | Q8WXA3;H0YD93 | -0.2621 | 0.0056 |
| SBSN | Suprabasin | Q6UWP8;K7ESC4 | -1.6230 | 0.0267 |
| SCAPER | S-phase cyclin A associated protein in the ER | Q9BY12;H3BPM0;H3BS25;H3BU24;H3BR40;H3BT27;H3BTL8 | -0.9431 | 0.0446 |
| SDK2 | Sidekick cell adhesion molecule 2 | Q58EX2;H7C2P2 | -1.0466 | 0.0205 |
| SF3A3 | Splicing factor 3a subunit 3 | Q12874 | -0.1478 | 0.0194 |
| SFXN3 | Sideroflexin 3 | Q9BWM7;A0A0A0MS41;A0A1P0AYU5;S4R3N9 | -0.4955 | 0.0375 |
| SLC12A4 | Solute carrier family 12 member 4 | Q9UP95;I3L1N8 | -0.5602 | 0.0385 |

| Gene Symbol | Protein Name | Accession | $Log_2FC$ | p-value |
|---|---|---|---|---|
| SLFN11 | Schlafen family member 11 | Q7Z7L1;K7EIM3;C9J902;C9JDG6;C9JUT2;K7ER38;K7EKT7;K7ES87 | -1.4030 | 0.0221 |
| SRC | SRC proto-oncogene, non-receptor tyrosine kinase | P12931;J3QRU1;P07947 | -0.2961 | 0.0289 |
| STK11 | Serine/threonine kinase 11 | K7EP59;Q15831;K7EMR0;K7EQN8 | -1.0618 | 0.0363 |
| STMN1 | Stathmin 1 | P16949;B5BU83;A2A2D0 | -0.6550 | 0.0035 |
| STMN2 | Stathmin 2 | Q93045;E5RGX5;Q6ZRC1 | -0.7001 | 0.0041 |
| SYNM | Synemin | O15061;A0A075B7B1;C9JIE4 | -0.5607 | 0.0055 |
| TACC1 | Transforming acidic coiled-coil containing protein 1 | O75410;R4GMT7;E7ET87;E7EVI4;H0YAY0;E5RFM9;E5RJG6 | -1.3200 | 0.0046 |
| TBCA | Tubulin folding cofactor A | O75347;E5RHG6;E5RJD8;E5RIW3;E5RIX8 | -0.8134 | 0.0254 |
| TIAM2 | TIAM Rac1 associated GEF 2 | Q8IVF5;E9PMZ8;F5H6W6;E9PKT1;F5H6R0 | -0.8456 | 0.0404 |
| TLE3 | TLE family member 3, transcriptional corepressor | H0YKN8;H0YKT5;H0YL70;H0YNT2;Q04726;A0A0D9SES8;F5H7D6;H0YNI7;Q04724 | -3.4835 | 0.0431 |
| TMED10 | Transmembrane p24 trafficking protein 10 | P49755;G3V2K7 | -0.5017 | 0.0028 |
| TNIP1 | TNFAIP3 interacting protein 1 | A0A0A0MRZ4;Q15025 | -0.5649 | 0.0161 |
| TOMM70 | Translocase of outer mitochondrial membrane 70 | O94826 | -1.0572 | 0.0205 |
| TONSL | Tonsoku like, DNA repair protein | Q96HA7 | -1.3844 | 0.0384 |
| TPR | Translocated promoter region, nuclear basket protein | P12270 | -0.2257 | 0.0417 |

| Gene Symbol | Protein Name | Accession | $Log_2FC$ | p-value |
|---|---|---|---|---|
| TRAF7 | TNF receptor associated factor 7 | Q6Q0C0;H3BR17 | -0.9751 | 0.0235 |
| TRIP12 | Thyroid hormone receptor interactor 12 | Q14669;C9JLD7;C9JLJ5;C9JSX9;F8W9P3;G5E9G6 | -3.0528 | 0.0208 |
| TTC3 | Tetratricopeptide repeat domain 3 | E9PCE7;E9PMP8;P53804 | -1.5647 | 0.0420 |
| TUBA3E | Tubulin alpha 3e | Q6PEY2 | -2.7836 | 0.0251 |
| TUBB2B | Tubulin beta 2B class IIb | Q9BVA1;K7EK43;G3V4U2;M0R2T4;I3L0U9;I3L0V9;I3L4T6;I3L4U4;K7EJ64;K7EJZ4;K7EN98;K7EPE5;K7EQT3;K7ERA8;K7ESQ3;Q8TBP0 | -0.3510 | 0.0423 |
| TUBB8 | Tubulin beta 8 class VIII | Q3ZCM7;A0A075B736;Q5SQY0 | -0.3406 | 0.0031 |
| UACA | Uveal autoantigen with coiled-coil domains and ankyrin repeats | Q9BZF9;F5H2B9;H0YNH8 | -0.1426 | 0.0065 |
| UBE2O | Ubiquitin conjugating enzyme E2 O | Q9C0C9;K7ES11;K7EQ12 | -0.7069 | 0.0169 |
| UGGT1 | UDP-glucose glycoprotein glucosyltransferase 1 | Q9NYU2;H7BZG0 | -0.3363 | 0.0248 |
| WASF1 | WASP family member 1 | Q5SZK4;Q5SZK5;Q92558;Q5SZK3;Q9UPY6 | -1.0625 | 0.0346 |
| WDR19 | WD repeat domain 19 | Q8NEZ3;D6R9P6;D6RE75;D6RAI4 | -0.9689 | 0.0050 |
| XAF1 | XIAP associated factor 1 | Q6GPH4;C9J7Z8;I3L3D9;I3L509;I3L2B3;I3L3B3;I3L3Q2;I3L534 | -1.1979 | 0.0443 |
| XPNPEP1 | X-prolyl aminopeptidase 1 | Q9NQW7;Q5T6H7;Q5T6H2 | -1.1189 | 0.0079 |

| Gene Symbol | Protein Name | Accession | $Log_2FC$ | p-value |
|---|---|---|---|---|
| YWHAG | Tyrosine 3-monooxygenase /tryptophan 5-monooxygenase activation protein gamma | P61981 | -0.5250 | 0.0272 |
| YWHAH | Tyrosine 3-monooxygenase /tryptophan 5-monooxygenase activation protein eta | Q04917 | -2.0400 | 0.0269 |
| ZNF565 | Zinc finger protein 565 | Q8N9K5 | -0.6053 | 0.0299 |
| ZNF594 | Zinc finger protein 594 | Q96JF6;I3L508 | -1.7823 | 0.0364 |
| ZNF677 | Zinc finger protein 677 | Q86XU0;M0R297 | -0.6763 | 0.0498 |
| ZSCAN5A | Zinc finger and SCAN domain containing 5A | A0A0C4DGQ1;Q9BUG6 | -1.9495 | 0.0170 |

**Table S3.** Result of the pathway and process enrichment analysis conducted on DAPs induced by 100 nM LSD using Metascape. The table showcases clusters of enriched terms categorized under two key enrichment ontology categories: Reactome and KEGG terms.

| Term | Description | LogP | Log(q-value) | Gene Symbols |
|---|---|---|---|---|
| R-HSA-9716542 | Signaling by Rho GTPases, Miro GTPases and RHOBTB3 | -16.9685 | -13.5099 | ACTN1,ARHGAP5,C1QBP,CDC42,GPS1,HNRNPC,KTN1,PAK2,SRC,STK10,TPM4,YWHAG,YWHAH,DDX39B,PKP4,DYNLL1,WASF1,ACTR3,ARPC2,PLIN3,DLC1,STMN2,AKAP13,TOR1AIP1,TIAM2,UACA,RNF20,GOPC,ARHGAP21,MYL12B,TUBA3E,DYNLL2,PRAG1,TUBB8,TUBB2B |
| R-HSA-199991 | Membrane Trafficking | -12.6279 | -9.7702 | AP2B1,FTL,GPS1,LRP2,PAFAH1B2,RAB3A,RAB5C,SRC,SYT1,YWHAG,YWHAH,CLTCL1,DYNLL1,NAPA,COPB2,KIF20B,ACTR3,ARPC2,PLIN3,COPS8,TMED10,RABGAP1,VPS4A,RAB8B,TUBA3E,DYNLL2,TUBB8,TUBB2B |

| Term | Description | LogP | Log(q-value) | Gene Symbols |
|---|---|---|---|---|
| R-HSA-422475 | Axon guidance | -8.4132 | -5.7327 | AP2B1,CDC42,CRMP1,PAK2,MAPK8,PSMC2,PSMD10,PTPN11,RPL9,RPL12,RPL35A,SCN4A,SRC,CLTCL1,CACNA1I,ACTR3,ARPC2,MYL12B,TUBA3E,TUBB8,TUBB2B,MYO1F,WASF1,TMED10,KTN1,YWHAG,YWHAH,DYNLL1,GOPC,DYNLL2,ACTN1,STK11,RAB8B,AMOTL1,RAB5C,DYNLRB2,EEA1,USP8,VPS4A,RUFY2,RAB11FIP4,PXN,GSK3A,TNIP1,BRAF,TRIP12,UBE2O,ANAPC1,TRAF7,MARCKS,GNB4 |
| R-HSA-9663891 | Selective autophagy | -7.7747 | -5.3161 | SRC,DYNLL1,TOMM70,HDAC6,PLIN3,TUBA3E,DYNLL2,TUBB8,TUBB2B,PAFAH1B2,NAPA,COPB2,KIF20B,TMED10,MAX,POLE,PSMC2,PSMD10,RBBP4,RBBP7,TPR,YWHAG,YWHAH,NUP93,PHF8,VPS4A,ANAPC1,CEP76,ABRAXAS1,EEF1A1,AHI1,WDR19,DYNLRB2,ATP5PB,POLRMT,AP2B1,CYC1,DNAH5,NDUFS2,MAPK8,CLTCL1,DNAH3,PFDN4,TBCA,DCAF7,GAPDHS,GNB4,CDC42,BRAF,LRP5,LRRK2,DAD1,UGGT1,SCN4A,ZFPM1,ACTN1,C1QBP,CLU,PTPN11,RAB5C,EEA1 |

| Term | Description | LogP | Log(q-value) | Gene Symbols |
|---|---|---|---|---|
| R-HSA-109581 | Apoptosis | -7.6229 | -5.2056 | C1QBP,DAPK1,H1-4,PAK2,MAPK8,PSMC2,PSMD10,YWHAG,YWHAH,DYNLL1,UACA,DYNLL2,WDR19,DYNLRB2,TUBB2B,ANAPC1,ABRAXAS1,RAB5C |
| R-HSA-2262752 | Cellular responses to stress | -6.4608 | -4.2326 | EEF1A1,GSK3A,H1-4,PGD,MAPK8,PSMC2,PSMD10,RBBP4,RBBP7,RPL9,RPL12,RPL35A,TPR,DYNLL1,NUP93,HDAC6,NPLOC4,ANAPC1,TUBA3E,DYNLL2,TUBB8,TUBB2B |
| R-HSA-70171 | Glycolysis | -5.8098 | -3.8563 | ALDOB,PFKFB3,PFKM,PKLR,TPR,NUP93,GAPDHS,PGD,PRPS1L1,DSE,IDH3A |
| R-HSA-9013406 | RHOQ GTPase cycle | -5.1329 | -3.2763 | ARHGAP5,CDC42,PAK2,DLC1,GOPC,ARHGAP21,C1QBP,STK10,AKAP13,KTN1,WASF1,TIAM2,ACTN1,TOR1AIP1,PKP4,PRAG1,STMN2 |
| R-HSA-9706574 | RHOBTB GTPase Cycle | -5.0969 | -3.2510 | ACTN1,GPS1,HNRNPC,DDX39B,RNF20 |
| R-HSA-112315 | Transmission across Chemical Synapses | -4.9391 | -3.1240 | AP2B1,GABRA6,KCNJ16,RAB3A,SRC,SYT1,PPFIA4,GNB4,TUBA3E,TUBB8,TUBB2B,GPHN |
| R-HSA-445355 | Smooth Muscle Contraction | -4.5493 | -2.7925 | PAK2,PXN,TPM4,CACNA1I,MYL12B,SCN4A |
| R-HSA-72706 | GTP hydrolysis and joining of the 60S ribosomal subunit | -4.4939 | -2.7596 | EIF1AX,EIF4A2,EIF5,RPL9,RPL12,RPL35A,EIF3M,EEF1A1,PPA1,MRPS36,TPR,NUP93 |

| Term | Description | LogP | Log(q-value) | Gene Symbols |
|---|---|---|---|---|
| hsa04510 | Focal adhesion | -4.4578 | -2.7456 | ACTN1,ARHGAP5,BRAF,CDC42,PAK2,MAPK8,PXN,SRC,MYL12B,LCP1,BOLA2,PTPN11,DUSP3,STMN1,MAX,NF1,CACNA1I,WASF1,YWHAG,YWHAH,HDAC6,RAB5C,GNB4,TPM4,GSK3A,PSMC2,PSMD10 |
| R-HSA-5663202 | Diseases of signal transduction by growth factor receptors and second messengers | -4.3282 | -2.6517 | BRAF,CTBP2,GSK3A,LRP5,NF1,PSMC2,PSMD10,PTPN11,SRC,TPM4,TPR,HDAC6,FIP1L1 |
| R-HSA-9679506 | SARS-CoV Infections | -3.6490 | -2.1094 | AP2B1,DAD1,GSK3A,PTPN11,RBBP4,RBBP7,TPR,YWHAG,YWHAH,NUP93,TOMM70 |
| R-HSA-1295596 | Spry regulation of FGF signaling | -3.5939 | -2.0748 | BRAF,PTPN11,SRC,AP2B1,CDC42,DUSP3,STMN1,PAK2,PTN,PXN,WASF1,USP8,VRK3,MAPK8,ATP5PB,CYC1,NDUFS2,ACTN1,SDK2,DAPK1,CLTCL1 |
| R-HSA-264642 | Acetylcholine Neurotransmitter Release Cycle | -3.5121 | -2.0292 | RAB3A,SYT1,PPFIA4,AP2B1,CLTCL1,NAPA |
| R-HSA-376176 | Signaling by ROBO receptors | -3.4335 | -2.0003 | CDC42,PAK2,PSMC2,PSMD10,RPL9,RPL12,RPL35A,SRC,AP2B1,TPR,NUP93,GPS1,RBBP7,DCAF7,COPS8,NPLOC4,CCDC8,MAX,POLE,RBBP4,CTBP2,LRP5,PDE6B,TLE3,USP8,GNB4,ANAPC1,VPS4A,GSK3A,PTPN11,PGD |

| Term | Description | LogP | Log(q-value) | Gene Symbols |
|---|---|---|---|---|
| R-HSA-9613829 | Chaperone Mediated Autophagy | -3.1698 | -1.7975 | EEF1A1,HDAC6,PLIN3,TPR,NUP93 |
| R-HSA-1280215 | Cytokine Signaling in Immune system | -3.1381 | -1.7763 | CDC42,DUSP3,EIF4A2,LCP1,PAK2,MAPK8,PSMC2,PSMD10,PTPN11,FSCN1,TPR,NUP93,VRK3,XAF1,BOLA2 |

**Table S4.** Persistently modulated proteins across 10 and 100 nM LSD datasets, along with their respective modulations (upregulated [↑] or downregulated [↓]).

| Gene Symbol | Protein Name | 10 nM | 100 nM |
|---|---|---|---|
| ABCA8 | ATP binding cassette subfamily A member 8 | ↓ | ↓ |
| ABRAXAS1 | BRCA1 A complex subunit abraxas 1 | ↑ | ↑ |
| CLU | Clusterin | ↓ | ↓ |
| CPNE1 | Copine 1 | ↓ | ↓ |
| CPT2 | Carnitine palmitoyltransferase 2 | ↑ | ↑ |
| CTBP2 | C-terminal binding protein 2 | ↑ | ↑ |
| EIF5A2 | Eukaryotic translation initiation factor 5A2 | ↑ | ↑ |
| FADS1 | Fatty acid desaturase 1 | ↑ | ↓ |
| GABRA6 | Gamma-aminobutyric acid type A receptor subunit alpha6 | ↓ | ↓ |
| GOPC | Golgi associated PDZ and coiled-coil motif containing | ↓ | ↓ |
| HIST1H1E | H1.4 linker histone, cluster member | ↓ | ↓ |
| HMGCS1 | 3-hydroxy-3-methylglutaryl-CoA synthase 1 | ↓ | ↓ |
| LCP1 | Lymphocyte cytosolic protein 1 | ↓ | ↓ |
| LRRC37B | Leucine rich repeat containing 37B | ↑ | ↑ |
| MSH2 | MutS homolog 2 | ↓ | ↓ |
| MTMR10 | Myotubularin related protein 10 | ↓ | ↓ |
| OGA | O-GlcNAcase | ↓ | ↓ |
| PABPC4 | Poly(A) binding protein cytoplasmic 4 | ↓ | ↓ |
| POLRMT | RNA polymerase mitochondrial | ↑ | ↑ |
| RPL35A | Ribosomal protein L35a | ↑ | ↑ |
| TACC2 | Transforming acidic coiled-coil containing protein 2 | ↑ | ↑ |
| TCF25 | Transcription factor 25 | ↑ | ↑ |
| TNIP1 | TNFAIP3 interacting protein 1 | ↓ | ↓ |
| TTC7B | Tetratricopeptide repeat domain 7B | ↑ | ↑ |
| YWHAH | Tyrosine 3-monooxygenase/tryptophan 5-monooxygenase activation protein eta | ↓ | ↓ |
| ZNF302 | Zinc finger protein 302 | ↑ | ↑ |

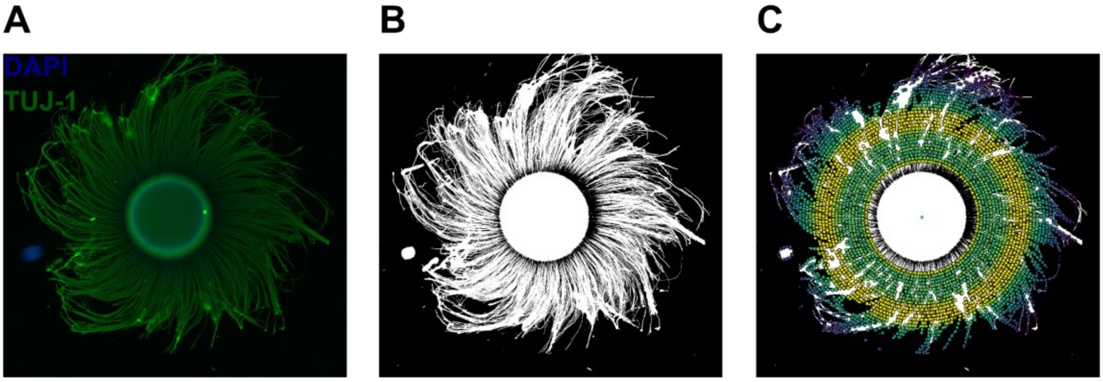


**Figure S1.** Sholl analysis in brain spheroids: (**A**) Representative image of a spheroid processed for immunofluorescence using antibody TUJ-1 against Beta-Tubulin III and 4',6-diamidino-2-phenylindole (DAPI) for nuclear staining; (**B**) Image of the spheroid of (**A**) post-processed with Fiji ImageJ. The process involves splitting channels and threshold adjustment for detailed visualization; (**C**) Representation of the Sholl analysis conducted on the image from (**B**), where each dot indicates an intersection of a neurite with a concentric circle.

**Table S5.** List of antibodies used for immunohistochemistry characterization of human cerebral organoids. The upper section corresponds to primary antibodies, while the lower section refers to secondary antibodies.

| Antibody | Origin | Source | Catalog | Dilution |
|---|---|---|---|---|
| Anti- β-Tubulin 3 (TUJ) | Chicken | Aves | TUJ | 1-400 |
| Anti-PAX6 IgG | Rabbit | Invitrogen | 42-6600 | 1-100 |
| Anti-GFAP IgG1 | Mouse | Neuromics | MO15052 | 1-100 |
| Anti-MAP2 IgG1 | Mouse | Sigma | M1406 | 1-300 |
| Anti-5-HT2A IgG | Rabbit | Invitrogen | PA5-103377 | 1-100 |
| Anti-β-Tubulin 3 IgG2A (TUJ1) | Mouse | Neuromics | MO15013 | 1-2000 |
| Anti-Mouse IgG (H+L), Alexa Fluor 594 | Goat | Invitrogen | A-11032 | 1-400 |
| Anti-Rabbit IgG (H+L), Alexa Fluor 488 | Goat | Invitrogen | A-11008 | 1-400 |
| Anti-Rabbit IgG (H+L), Alexa Fluor 568 | Goat | Invitrogen | A-11036 | 1-400 |
| Anti-Chicken IgY (H+L), Alexa Fluor 488 | Goat | Abcam | ab150173 | 1-250 |
